# The dynamic response to hypoosmotic stress reveals distinct stages of freshwater acclimation by a euryhaline diatom

**DOI:** 10.1101/2022.06.24.497401

**Authors:** Kala M. Downey, Kathryn J. Judy, Eveline Pinseel, Andrew J. Alverson, Jeffrey A. Lewis

**Affiliations:** University of Arkansas, Department of Biological Sciences, Fayetteville, AR 72701 USA

## Abstract

The salinity gradient separating marine and freshwater environments is a major ecological divide, and the mechanisms by which most organisms adapt to new salinity environments are poorly understood. Diatoms are a lineage of ancestrally marine microalgae that have repeatedly colonized and diversified in freshwaters. *Cyclotella cryptica* is a euryhaline diatom that naturally tolerates a broad range of salinities, thus providing a powerful system for understanding the genomic mechanisms for mitigating and acclimating to low salinity. To understand how diatoms mitigate acute hypoosmotic stress, we abruptly shifted *C. cryptica* from seawater to freshwater and performed transcriptional profiling during the first 10 hours. Freshwater shock dramatically remodeled the transcriptome, with ~50% of the genome differentially expressed in at least one time point. The peak response occurred within 1 hour, with strong repression of genes involved in cell growth and osmolyte production, and strong induction of specific stress defense genes. Transcripts largely returned to baseline levels within 4–10 hours, with growth resuming shortly thereafter, suggesting that gene expression dynamics may be useful for predicting acclimation. Moreover, comparison to a transcriptomics study of *C. cryptica* following months-long acclimation to freshwater revealed little overlap between the genes and processes differentially expressed in cells exposed to acute stress versus fully acclimated conditions. Altogether, this study highlights the power of time-resolved transcriptomics to reveal fundamental insights into how cells dynamically respond to an acute environmental shift and provides new insights into how diatoms mitigate natural salinity fluctuations and have successfully diversified across freshwater habitats worldwide.

## Introduction

The majority of aquatic organisms live exclusively in marine or freshwater environments, but some “euryhaline” species can tolerate a wide range of salt concentrations, allowing them to grow in environments with fluctuating salinities such as lagoons, estuaries, and coastal intertidal zones. The salinity in these habitats can vary in time from fresh to fully marine levels, and salinity shifts in these systems can occur gradually or within minutes to hours depending on rainfall, river flow, and tidal action (Balzano et al., 2015). Climate change is expected to increase the frequency and magnitude of these fluctuations. For example, the prevalence of large storms that inundate coastal habitats with freshwater from both precipitation and increased river flow is expected to increase in the future (Pörtner et al., 2019; Shu et al., 2018), and many marine and brackish environments will likely experience “freshening” due to melting ice caps and altered precipitation patterns (Lee et al., 2022). However, the specific mechanisms by which marine microbes mitigate the effects of short and long-term salinity changes remains poorly understood. Euryhaline species that have evolved to thrive across a wide range of salinities thus present an excellent opportunity to provide new insights into how these species will respond to increasingly dynamic salinity environments.

Diatoms are a diverse group of globally distributed unicellular algae that produce roughly one-fifth of the world’s oxygen and serve as the base for aquatic food webs (Armbrust, 2009). Diatoms are abundant in habitats across the entire marine to freshwater salinity gradient, and across evolutionary timescales, transitions between marine and fresh waters have been an important source of species diversification (Nakov et al., 2018). *Cyclotella cryptica* is one of many euryhaline diatoms that occurs naturally across a wide range of salinities and has served as a longstanding model for understanding the effects of salinity on diatom physiology and morphology. For example, salinity is known to impact the morphology of the siliceous cell wall of *C. cryptica*, including alterations to its thickness and ornamentation (Conley et al., 1989; Schultz, 1971). Acute salinity shifts can also induce gametogenesis in *C. cryptica* (Schultz & Trainor, 1970) and other diatoms (Godhe et al., 2014).

Hypoosmotic stress results in water influx that increases intracellular turgor pressure, which risks rupturing the plasma membrane (Kirst, 1990; Theseira et al., 2020; Van Bergeijk et al., 2003). Cells respond by modulating internal osmolyte concentrations, which can be accomplished either through transport of inorganic ions such as sodium or potassium (Krell et al., 2008), or by degrading organic osmolytes such as dimethylsulfoniopropionate (DMSP) (Lyon et al., 2016; Lyon et al., 2011), glycine betaine (Dickson & Kirst, 1987; Kageyama et al., 2018), or proline (Krell et al., 2007; M. S. Liu & Hellebust, 1974). In the early stages (i.e., 15 minutes to 3 hours) of hypoosmotic stress, *C. cryptica* reduces the concentration of osmolyte-functioning amino acids by incorporating them into proteins (M. S. Liu & Hellebust, 1974; Ming Sai Liu & Hellebust, 1976). However, transcriptional profiling showed that reduced proline levels do not appear to play an important role during long-term acclimation, where instead cells maintain reduced levels of glycine betaine, DMSP, and taurine (Nakov et al., 2020). This finding suggests that the processes diatoms rely upon to withstand short-term stress are different from those deployed during long-term acclimation. Moreover, it is clear that the response to acute hypoosmotic stress encompasses more than just strategies to maintain osmotic balance (Nakov et al., 2020; Pinseel et al., 2022). To address these gaps in our understanding, we exposed a brackish strain of *C. cryptica* to freshwater and used RNA sequencing (RNA-seq) to characterize changes in gene expression in the minutes to hours following a hypoosmotic shock. Comparisons to expression-level changes in fully acclimated cells of *C. cryptica* revealed important differences between short- and long-term responses to hyposalinity conditions by diatoms. Altogether, the results from this study highlight the power of time-resolved transcriptomics during an acute and dynamic environmental response, which we leveraged to provide new insights into how diatoms mitigate natural salinity fluctuations and successfully colonized and diversified in freshwater habitats worldwide.

## Materials and Methods

### Culture conditions

*Cyclotella cryptica* strain CCMP332 was obtained from the National Center for Marine Algae and Microbiota and maintained in artificial sea water at 24 parts per thousand salinity (ASW 24) (Nakov et al., 2020), which is within the range of the brackish salinity of the locality from which the strain was originally isolated (Reimann et al., 1963). To obtain sufficient biomass for subsequent experiments, cells were grown for 7 days in a 500 mL Erlenmeyer flask, placed in a Percival incubator at 15 °C and 22 μmol photons m^−2^ s^−1^ irradiance under a 16:8 h light:dark cycle. Cells were homogenized by agitating the flask, and 3 × 2-mL aliquots of cells were used to inoculate 3 × 1-L Erlenmeyer flasks containing 500 mL of ASW 24. Growth was monitored by counting cells with a Benchtop B3 Series FlowCAM^®^ cytometer (Fluid Imaging Technologies). Upon reaching exponential growth in ASW 24, cells from each triplicate culture were immediately exposed to freshwater conditions (ASW 0). To do this, we harvested 24 × 10^6^ cells from each 1-L flask for freshwater treatments, centrifuged the cells at 800 rcf for 3 min at 4 °C, decanted the supernatant, and resuspended the cells in 16 mL of ASW 0 to a final concentration of 1.5 × 10^6^ cells/ml. Two-mL aliquots of the resuspended cells were then immediately transferred into 8 tubes containing 38 mL of ASW 0, and the cells were incubated at 15 °C and 20 μmol photons m^−2^ s^−1^ irradiance with gentle agitation using a Boekel Scientific adjustable speed wave rocker.

Cells were collected for transcriptional profiling at seven different time points following inoculation in ASW 0: 15 min, 30 min, 1 h, 2 h, 4 h, 8 h, and 10 h. The 0-min sample was collected immediately following inoculation in ASW 0, and served as a control for comparing the freshwater response. Cells were harvested by centrifugation at 400 rcf for 3 min at 4 °C, flash-frozen in liquid nitrogen, and stored at −80 °C until processed.

Growth in ASW 24 and 0 (freshwater) was measured at each of the time points used for transcriptional profiling, as well as at 12 h, 24 h, and 7 days. Cells were handled identically as described above for time point collections, after which we transferred a 2-mL aliquot of concentrated cells to a 12-well plate. We treated the concentrated cells with 0.1% Lugol’s iodine solution as a fixative, and then counted cells with a Benchtop B3 Series FlowCAM^®^ cytometer (Fluid Imaging Technologies).

### RNA extraction and library preparation

To reduce potential batch effects, numbered samples were randomized using the ‘sample’ function in R v 4.05 prior to RNA extraction and library construction. RNA was extracted with an RNeasy Plant Mini Kit (QIAGEN, Netherlands) according to the manufacturer’s instructions and quantified with a Qubit 2.0 Fluorometer (Invitrogen, USA). RNA quality was assessed using an Agilent Technologies 2200 TapeStation (Agilent Technologies, USA). RNA concentrations and RNA integrity number (RIN) values can be found in Table S1. All samples had a RIN > 6. We constructed dual-indexed RNA libraries with the KAPA mRNA HyperPrep kit (KAPA Biosystems, USA), using half reaction volumes according to manufacturer’s instructions (a detailed protocol can be found on protocols.io under DOI: dx.doi.org/10.17504/protocols.io.uueewte). Libraries were synthesized using at least 150 ng total RNA, fragmentation time was optimized to generate 300-400-nt fragments, and libraries were amplified with 10 PCR cycles. Libraries were multiplexed and sequenced on a single lane of an Illumina HiSeq 4000 (2 × 100 bp paired-end reads) at the University of Chicago Genomics Facility. Additional details related to RNA and library quantity and/or quality are available in Table S1.

### RNA sequencing analysis

Quality of the raw reads was examined using FastQC v0.11.5 (Andrews, 2010). Low-quality reads were removed and adapter sequences trimmed (KAPA v3 dual indices) using kTrim v1.1.0 (Sun, 2020) with the following settings ‘-t 15 -p 33 -q 20 -s 36 -m 0.5’. Reads were mapped to the reference genome of *C. cryptica* (Roberts et al., 2020) using STAR v2.7.3a with settings ‘--alignIntronMin 1 --alignIntronMax 22618’ to account for the size distribution of annotated introns (Dobin & Gingeras, 2015). Mapping statistics are available in Table S1. Gene-level counts were quantified from uniquely mapped reads with HTSeq v0.11.3 in *union* mode (Anders et al., 2014), and can be found in Table S2.

Trimmed mean of M-values (TMM) normalization and differential expression analysis were conducted using the Bioconductor package edgeR v3.30.3 (Robinson et al., 2010). Only genes with at least one read count per million (CPM) in at least three samples were included in the analysis. Differential expression was performed in edgeR using the quasi-likelihood (QL) model (glmQLFit) with a group-model design that included each replicate timepoint combination and a Benjamini-Hochberg false discovery rate (FDR)-adjusted p-value cutoff of 5% (Lund et al., 2012). Each time point was compared to the control (0 min ASW 0 treatment). To take into account testing of multiple hypotheses for each gene across each comparison (i.e., significant differential expression across multiple time points), we used stageR v1.14.0, which allowed us to control the gene-level FDR across all contrasts with a 5 % cutoff (Heller et al., 2009; Van den Berge et al., 2017).

We compared previously published RNA-seq data from *C. cryptica* CCMP332 after long-term acclimation (120 days) to ASW 0 (Nakov et al., 2020). However, rather than having independent replicates of the same strain, the unit of replication by Nakov et al. 2020 included four different strains of *C. cryptica* which were all sampled once. Consequently, the Nakov et al. 2020 dataset included only one sample for *C. cryptica* strain CCMP332. To accommodate this, we followed recommendations by Robinson et al. 2010 for unreplicated data and used the edgeR exactTest function with an assigned dispersion of 0.16 to identify genes with differential expression when contrasting an unstressed control in ASW 24 versus the long-term (120 days) acclimated growth in ASW 0 measured by Nakov et al. 2020. To further compensate for the lack of replication, we restricted our analyses to genes with ≥ 1 log_2_-fold change. The outputs from differential expression analysis can be found in Table S3.

Global similarity of gene expression patterns for our dataset at each time point was assessed with metric multidimensional scaling of logCPM for the top 500 genes with largest standard deviations across samples (time points and replicates), using limma’s plotMDS function (Ritchie et al., 2015) (Figure S1). Hierarchical clustering was performed with Cluster v3.0 (Eisen et al., 1998) using uncentered Pearson’s correlation as the similarity metric and centroid linkage. Gene ontology (GO) enrichment analyses were conducted using the elim algorithm and Fisher’s exact test implemented in TopGO v2.36.0 (Alexa & Rahnenfuhrer, 2022), with Bonferroni-corrected p-values ≤ 0.05 considered significant.

Functional gene annotations were based primarily on the published annotation of the reference genome (Roberts et al., 2020) but were augmented in some cases with NCBI-BLASTP (Altschul et al., 1990) and searches against the Swissprot and Uniprot databases with an e-value cutoff 1e-6. KEGG pathway annotations were obtained from the KofamKOALA web server on 2020-09-17 (Aramaki et al., 2020). Related GO terms were condensed using the online server REVIGO (Supek et al., 2011) based on a 0.5 similarity threshold using the SimRel algorithm. All code used in this manuscript are provided as Markdown files (Appendices S1 and S2).

## Results

### Hypoosmotic stress causes transcriptional remodeling in *C. cryptica* that parallels transient growth arrest and acclimation

To understand the physiological and transcriptomic effects of acute hypoosmotic shock, we transferred *C. cryptica* strain CCMP332 from its native, brackish salinity (ASW 24) into freshwater (ASW 0), mimicking a dispersal event or rapid environmental fluctuation. Following the exposure to freshwater, we measured cell division in the first hours and up to 7 days, and used RNA-seq to characterize patterns of gene expression across 7 timepoints within the first 10 hours. Growth was temporarily halted in the 4 hours immediately following freshwater exposure (Figure 1). Transient arrest of the cell cycle during stress is common in eukaryotes (Nitta et al., 1997; Seaton & Krishnan, 2016; Skirycz et al., 2011; West et al., 2004), and is likely important for redirecting cellular resources away from growth and towards induction of stress defense genes (Ho et al., 2018). Consistent with this hypothesis, maximal changes in gene expression also peaked during this period of arrested growth (Figure 2). Once growth resumed, *C. cryptica* showed a reduced growth rate in freshwater over the course of 7 days compared to the ASW 24 brackish water control (doubling time of 147 hours in ASW 0 versus 86 hours in ASW 24, Figure 1). This is consistent with our previous observation of a reduced growth rate for long-term (120-day) freshwater acclimated *C. cryptica* CCMP332 compared to growth in the brackish (ASW 24) control (Nakov et al., 2020).

**FIGURE 1.**
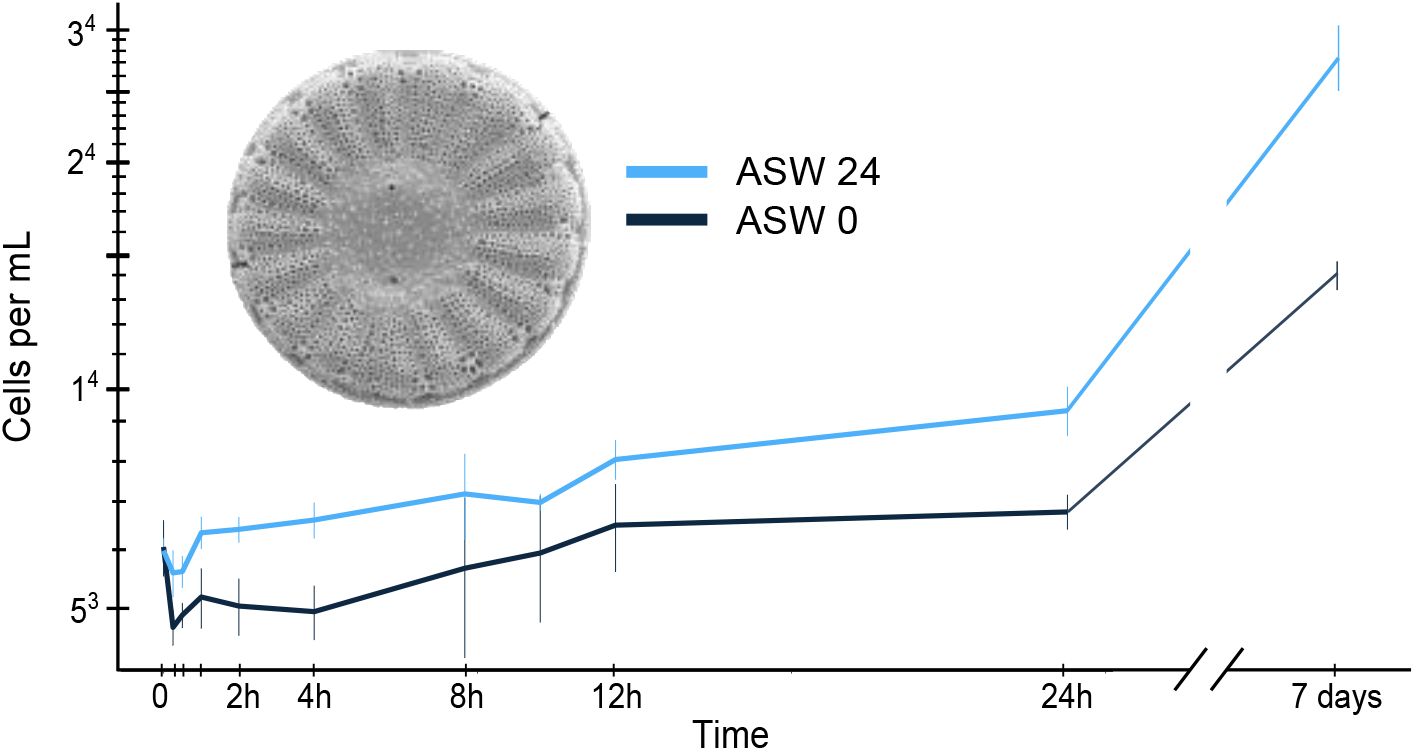
Hypoosmotic shock causes transient growth arrest in *C. cryptica* followed by acclimation to a reduced growth rate. Cell counts were obtained for *C. cryptica* transferred to freshwater (ASW 0) compared to its native brackish water control (ASW 24) over a 7-day time period. Error bars denote the standard deviation of biological triplicates. The inset image is a scanning electron micrograph of *C. cryptica* strain CCMP332 (cell diameter ≈ 10 μm). Raw count data is available in Table S4.

Transcriptional profiling revealed a substantial remodeling of global gene expression during acute hypoosmotic stress. Of the 21,250 predicted genes in the *C. cryptica* genome, 12,939 (61%) were expressed under the conditions of the experiment (defined as counts per million > 1 in at least 3 samples). Among these expressed genes, 10,566 (82%) were differentially expressed in at least one time point compared to the unstressed control (FDR < 0.05). This corresponds to roughly half of all genes in the genome, highlighting the profound effects of hypoosmotic stress on the ability of *C. cryptica* to maintain homeostasis. A large fraction (31%) of differentially expressed genes were induced or repressed less than 1.5-fold, which we interpret as hypoosmotic stress causing subtle, yet reproducible, downstream effects on many aspects of *C. cryptica* physiology. For example, at 60 minutes there was slight upregulation of genes encoding proteins involved in redox functions and regulation of potassium channels, and a slight downregulation of many genes associated with macromolecule localization and metal ion transport (Table S5).

With the rationale that the genes most critical to the hypoosmotic response should have consistent patterns of differential expression across time, we narrowed our focus to genes with significant differential expression in the same direction for at least 2 consecutive time points. A total of 4,298 genes met this criterion and were included in the downstream analyses. The largest numbers of differentially expressed genes, and the largest magnitude of expression changes, occurred at the 30- and 60-minute time points (Figure 2a), indicating that the peak of the transcriptional response occurred within this time frame. An MDS plot comparing the overall similarity of expression profiles at each time point showed that the 30- and 60-minute samples were the most dissimilar to the control (Figure S1). Genes encompassing the peak response gradually returned to near baseline levels by 4–10 hours (Figure 2b and Figure S1). Notably, that 4–10 hour time period coincided with the resumption of growth (Figure 1 and Figure 2b), suggesting that the short-term acute transcriptional response to hyposaline stress potentiates growth acclimation (see Discussion).

**FIGURE 2.**
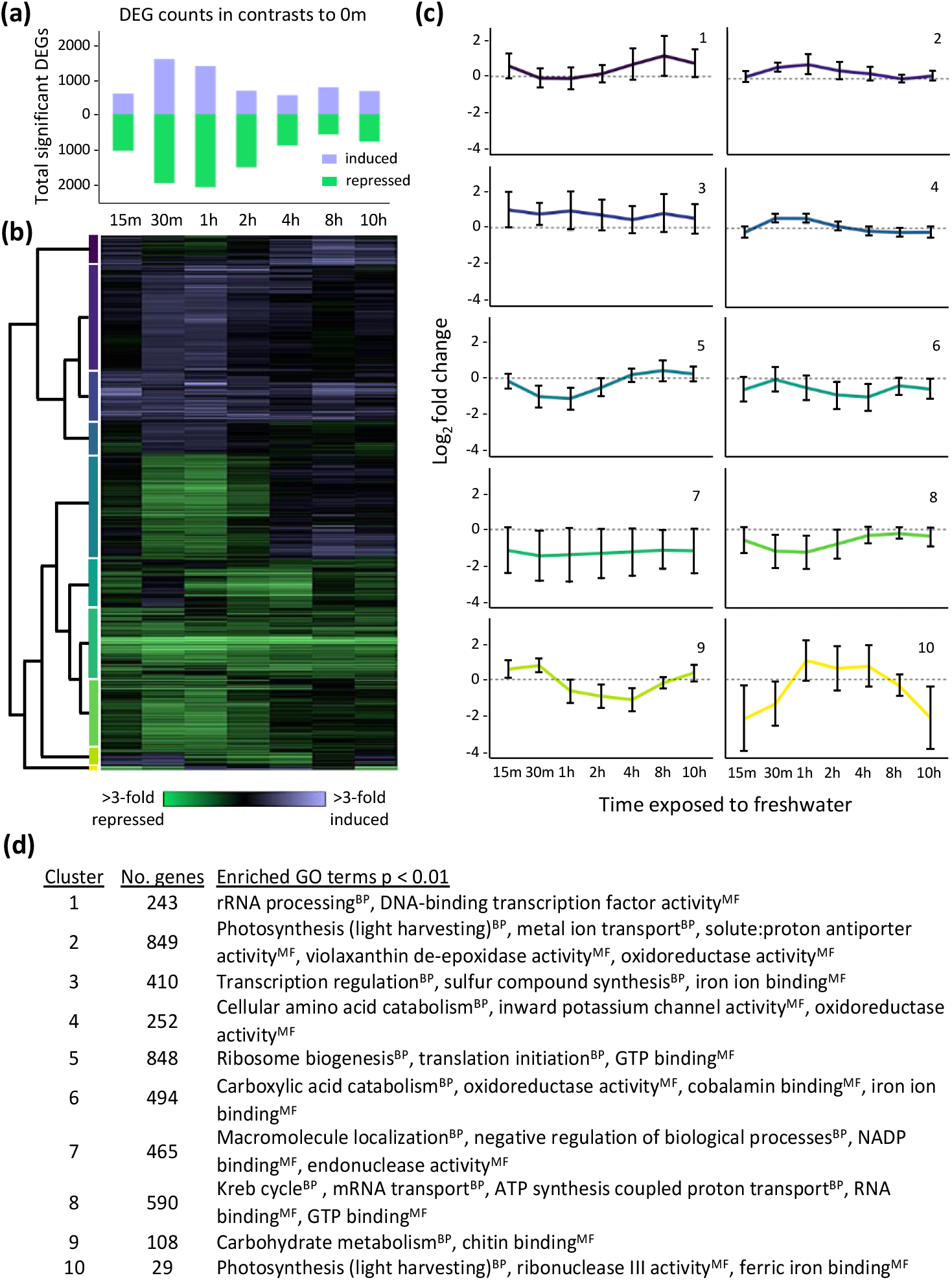
Substantial and dynamic remodeling of the *C. cryptica* transcriptome during hypoosmotic stress. (a) Total number of significant differentially expressed genes (DEGs) that respond to hypoosmotic stress at each time point relative to the control. (b) Heat map depicting hierarchical clustering of log_2_ fold changes of 4,298 genes that were differentially expressed at two or more consecutive time points. Each row represents a gene, and each column represents a time point following exposure to hypoosmotic stress. Purple indicates induced genes and green indicates repressed genes in response to hypoosmotic stress, according to the key. (c) Mean log_2_ fold changes for 10 clusters (panel b), organized from top to bottom according to the vertical color key next to the heat-map dendrogram (panel b). Error bars represent the 95% confidence interval for all genes within a cluster. (d) Enriched functional groups for each cluster (Bonferroni-corrected *P* < 0.01). Superscripts denote biological process (BP) or molecular function (MF) GO categories. Full GO annotations for each cluster can be found in Table S6.

### Key differences in the genes and processes that comprise the “peak” and “late” phases of the acute hypoosmotic stress response

Next, we sought to understand the major biological processes that were differentially regulated in response to hyposalinity stress. The 60-minute time point had the largest number of genes (1,480) with the greatest magnitude of expression change, suggesting that this time point best captured the peak of the response. These peak response genes largely returned to their pre-stress levels of expression by 8 hours (Figure 3a). Genes significantly induced at least 2-fold at 60 minutes were enriched for upregulation of photosynthetic processes, indicators of oxidative stress, and transcriptional regulation (Figure 3c, Table S7). In contrast, genes significantly repressed at least 2-fold at 60 minutes were enriched for ribosome biogenesis, transcriptional regulation, translational initiation, protein modification and localization, and ATP metabolism (Figure 3c, Table S7). Repression of genes related to protein synthesis and growth is common in eukaryotic stress responses (Gasch et al., 2000; Mayer et al., 2005), and is likely important for redirecting translational capacity towards induced stress-defense transcripts (Ho et al., 2018; M. V. Lee et al., 2011).

**FIGURE 3.**
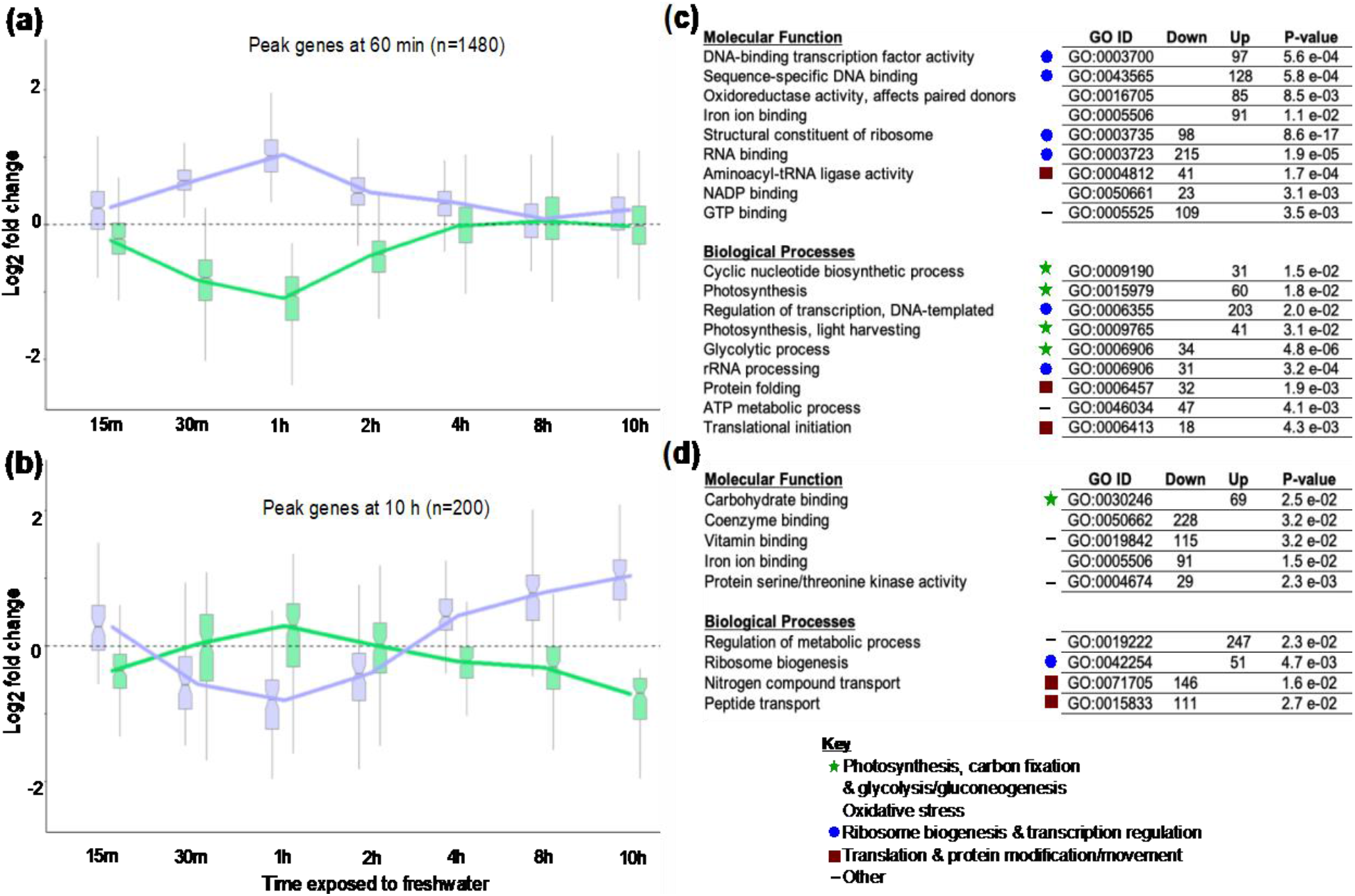
Distinct cellular processes are regulated during the peak versus late acute hypoosmotic response. (a) Box and whisker plots of significantly differentially expressed genes with the highest magnitude of change at 60 minutes (1,480 genes) versus (b) 10 hours (200 genes). Boxes represent the median and interquartile ranges, the notches bracketing the median represent the 95% confidence intervals, and the whiskers represent 1.5 times the interquartile range. Genes induced at either 60 minutes (panel a) or 10 hours (panel b) are purple, while repressed genes at the relevant time-point comparisons are green. (c and d) Functional enrichments for genes with ≥ 2-fold induction or repression at 60 minutes (c), or 10 hours (d). Symbols denote major cellular processes associated with each GO term. Full GO annotations for the 60-minute and 10-hour time points can be found in Table S7.

We were interested in understanding whether late-responding genes had different characteristics compared to peak-response genes. In contrast to the 1,480 genes with maximal expression changes at 60 minutes, only 200 genes experienced their maximal expression change at the 10 hour mark (Figure 3b). There was relatively little shared functional enrichment between genes with maximal expression at 60 minutes versus 10 hours, with only “ribosome biogenesis” and “iron ion binding” shared between the two (Figure 3d). This low level of functional overlap was driven by a majority (~80%) of the genes with maximal expression at 60 minutes returning to nearly pre-stress expression levels by 10 hours (Figure 3a). Intriguingly, genes with maximal expression changes at 10 hours had, on average, opposite expression patterns at 60 minutes (Figure 3a, b). The genes upregulated at 10 hours were functionally enriched for ribosome biogenesis, carbon fixation, and general regulation of metabolism–with ribosome biogenesis being notably repressed at 60 minutes (Figure 3d, Table S7). Likewise, the genes repressed at 10 hours were functionally enriched for peptide transport and oxidative stress–again, oxidative stress defense genes are induced at 60 minutes. One possible explanation for the discordant direction of gene expression changes between the 60-minute and 10-hour time points is that a subset of genes that respond mildly to acute stress are important for the early stages of acclimation. For example, the timing of de-repression of ribosome biogenesis genes, which are required for cell growth, just precedes growth resumption.

### Groups of functionally-related genes show distinct expression sub-dynamics during hypoosmotic stress

While the overall peak of the hypoosmotic stress response occurred approximately 60 minutes after introduction into freshwater, hierarchical clustering revealed groups of genes with distinct sub-dynamics that deviated from the behavior of peak response genes. We collapsed the expression dendrogram into 10 clusters based on their distinct expression profiles (Figure 2b, c). Notably, the expression patterns of five of these clusters (2, 4, 5, and 8) showed short-term changes across the early time points followed by acclimated gene expression patterns (i.e., a return to near baseline), that we hypothesize are related to the short-term growth arrest that occurred during the first four hours of hypoosmotic shock (Figure 1). We also found that distinct clusters with similar expression patterns often shared functional enrichments (Figure 2c, d). For example, clusters enriched for ‘ion transporter activity’ and ‘oxidoreductase activity’ (Figure 2c, d; Clusters 2, 4, and 6) were generally induced between 30–60 minutes then downregulated from 2 hours onward. Conversely, clusters enriched for genes involved in transcription, translation, and ribosome biogenesis (Figure 2c, d; Clusters 1 and 5) were repressed between 15–60 minutes, followed either by a return to baseline expression (Cluster 5) or upregulation (Cluster 1) between 2–10 hours. Cluster 8, enriched for genes involved in energy production (Krebs cycle and ATP synthesis-coupled proton transport), showed similar dynamics to the “translation-related” Clusters 1 and 5, suggesting that these processes may be coordinately regulated during hypoosmotic stress.

Although the majority of genes fell into clusters where expression changes peaked within 30–60 minutes, four clusters had unique expression patterns. Cluster 3 showed constant upregulation, and was enriched for both positive and negative regulation of transcription, and defense against oxidative stress. In contrast, Cluster 7 showed continuous downregulation at all time points and was enriched for processes related to macromolecule trafficking and negative regulation of cell division and DNA replication. Finally, clusters with the smallest gene membership (Clusters 9 and 10) showed either delayed repression (Cluster 9) or delayed induction (Cluster 10). Annotated genes in cluster 9 were enriched for genes associated with chitin metabolism, while the few known genes in cluster 10 were enriched for photosynthesis-related processes and ribonuclease III activity. Clusters 9 and 10 also had a high proportion of uncharacterized genes (25% and 50%, respectively), so the biological implications of these “late responding” clusters are not completely clear.

### Stress defense genes induced by hypoosmotic stress

To understand how *C. cryptica* maintains homeostasis during acute hypoosmotic shock, we analyzed the expression of genes known to play a role in stress defense in other species. Hypoosmotic shock initially causes an influx of water and efflux of ions and other osmolytes, so we examined how genes that encode channels and transporters responded throughout the time course. None of the genes encoding aquaporin water channels were significantly differentially expressed at two time points, though one was strongly induced at 30 min (Figure S2). A total of 15 K^+^ channels and pumps were upregulated at intermittent periods throughout the experiment, as were 17 Na^+^ exchangers and pumps. These transporters were primarily grouped into Clusters 2 and 5, which display contrasting expression patterns. This may indicate that the genes required to maintain the ion gradient changed over time. In addition to transmembrane transporters in the cell membrane, two chloroplastic K^+^ efflux pumps were upregulated. Hypoosmotic stress also triggered upregulation of a mechanosensitive anion transporter in the plasma membrane during 30 minutes to 3 hours, potentially due to cell sensing of increased osmotic pressure. Outside of the subset of genes differentially expressed at two consecutive time points, an additional 66 K^+^ and Na^+^ transporters were differentially expressed at only one time point, with no consistent pattern of when differential expression occurred and with a roughly equal distribution of up- and downregulation.

Cells can respond to increased turgor pressure from hypoosmotic stress by regulating intracellular osmolyte concentrations. Diatoms achieve osmotic balance by modulating concentrations of low-molecular weight osmolytes including dimethylsulfoniopropionate (DMSP), taurine, glycine-betaine, and proline (Boyd & Gradmann, 2002; Tevatia et al., 2015). Several essential steps in proline biosynthesis were downregulated (Figure S2), suggesting that stressed cells decreased the intracellular concentration of this key osmolyte. For taurine, two genes involved in its degradation were repressed at various time points, suggesting that cytosolic taurine levels may actually be increasing during acute hypoosmotic stress in *C. cryptica* (Figure S2). There was no transcriptional evidence that cells adjusted the concentrations of other osmolytes within the 10 hours following freshwater exposure. For DMSP and glycine-betaine biosynthesis, neither of the two methyltransferases critical for their biosynthesis were differentially expressed (Table S3).

In addition to genes directly involved in maintaining osmotic balance, other classes of stress defense genes were also differentially expressed during the time course. This includes a number of genes involved in scavenging reactive oxygen species (ROS). Out of 11 probable thioredoxins, two were upregulated at different time points (Figure 2c; clusters 1 and 2, Figure S3), while one was downregulated for the duration of the experiment. We also identified five differentially expressed chloroplastic peroxiredoxins, with four upregulated at different time points and one strongly downregulated at all time points. Several ROS scavengers were upregulated around the peak period of the hypoosmotic stress response (30–60 minutes): two superoxide dismutases, one catalase, and one L-ascorbate peroxidase (Figure S3). A total of 15 of 28 putative heat shock protein (HSP) chaperones were differentially expressed, with the majority of these (12/15) downregulated at the peak period of the stress response, and becoming slightly upregulated or returning to baseline levels by 4 hours. The remaining three HSP chaperones showed the opposite pattern, being upregulated from 30–60 minutes and returning to baseline levels or becoming slightly repressed by 4 hours (Figure S5).

### Expression dynamics of metabolic pathways during hyposaline stress

In addition to “classic” stress defense genes, hypoosmotic stress resulted in substantial transcriptional remodeling of numerous other metabolic genes in *C. cryptica*. Below, we summarize the effects on the main metabolic pathways.

#### Krebs cycle

One of the two copies of the gene that encodes citrate synthase, which catalyzes the first step of the Krebs cycle, was slightly upregulated for the duration of the time course (Figure 4, Figure S4). However, the enzymes responsible for catalyzing the next steps were either not differentially expressed or were mildly downregulated for the first 2 hours followed by recovery to approximately unstressed levels by 4–8 hours.

**FIGURE 4.**
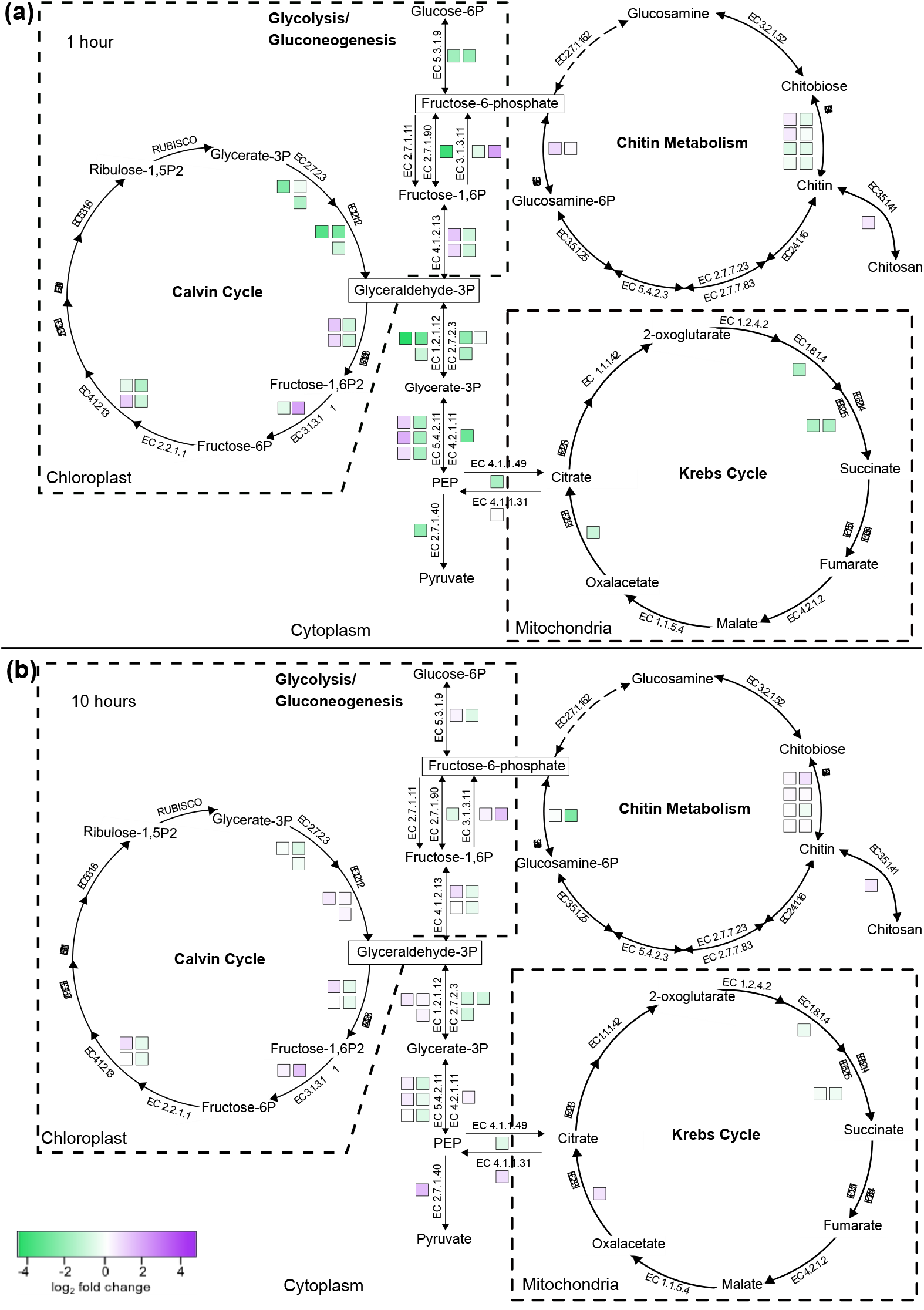
Hypoosmotic stress affects the expression of genes involved in major metabolic pathways. Each colored square denotes the expression of a single differentially expressed gene at (a) 60 minutes and (b) 10 hours, with purple indicating upregulation and green downregulation according to the key. Enzymatic steps encoded by multiple paralogs are indicated by the presence of multiple boxes (e.g., there are two paralogs for EC 5.3.1.9 responsible for the first step in glycolysis). Dotted lines circumscribe pathways with different organellar compartmentalization, and genes that participate in multiple pathways are separately color coded (e.g., EC 3.1.3.11 for both the Calvin cycle and gluconeogenesis).

**FIGURE 5.**
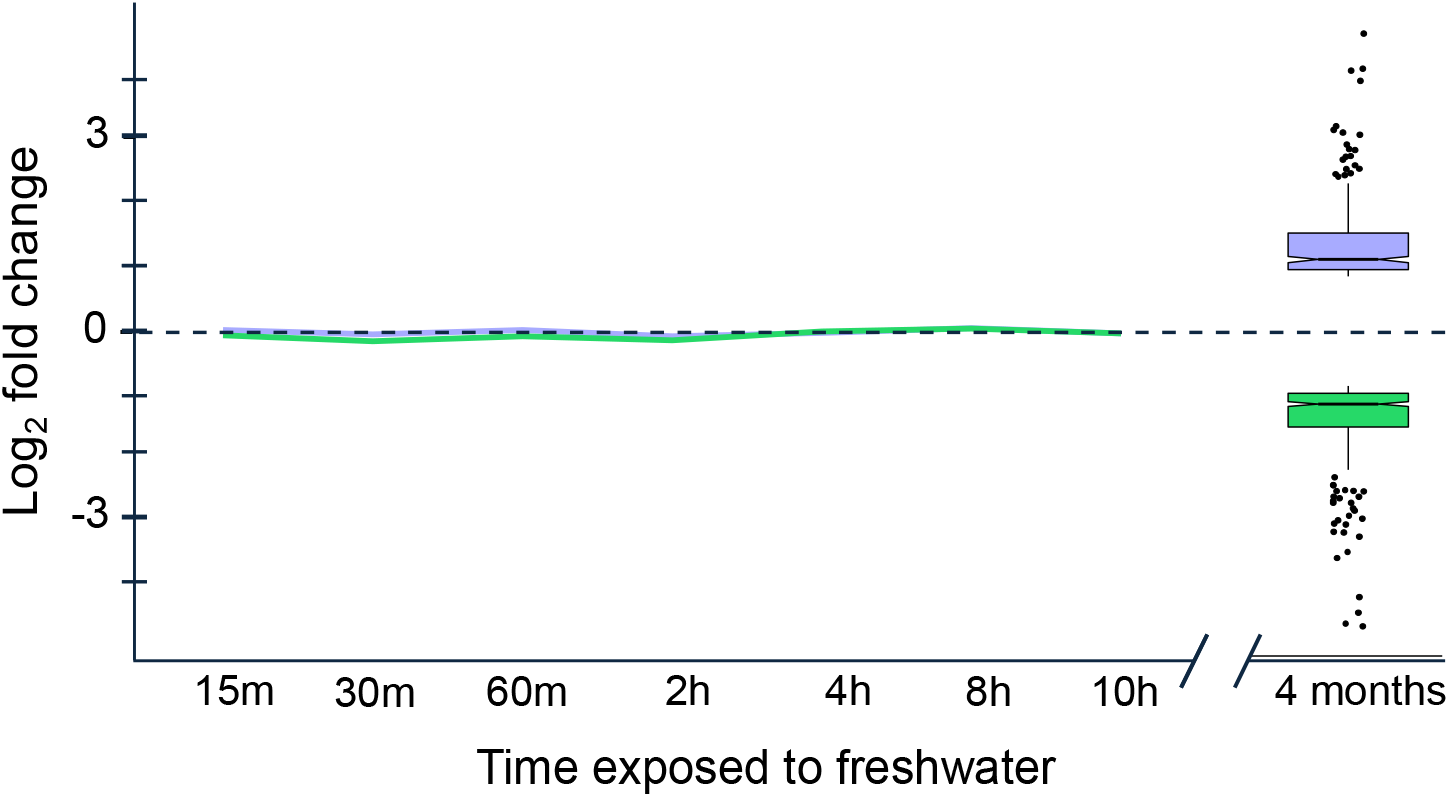
The genes differentially expressed during long-term acclimation of *C. cryptica* to freshwater are distinct from those that respond during acute hypoosmotic stress. The line graph depicts the expression pattern during our short-term transcriptional profiling for genes that we identified as differentially expressed ≥ 2-fold following months-long acclimation using data from Nakov et al. 2020. Box and whisker plots show the expression of those genes at four months, with boxes representing the median and interquartile ranges, notches bracketing the median representing the 95% confidence intervals, and the whiskers representing 1.5 times the interquartile range. Outliers are shown with dots horizontally jittered for legibility. GO annotations for differentially expressed genes at 4 months (120 days) can be found in Table S8.

#### Photosynthesis and carbon fixation

A total of 14 out of 24 genes associated with chlorophyll production, including three magnesium chelatase subunits that initiate chlorophyll biosynthesis (Papenbrock et al., 2000),, were upregulated at 30–60 minutes (Table S9). Additionally, 21 of the 26 genes that encode binding proteins for light-harvesting pigments were upregulated during this time period. Xanthophyll cycle genes coding for zeaxanthin and violaxanthin de-epoxidase were upregulated at 30–60 minutes and 30 minutes–4 hours, respectively. In contrast, genes associated with light-independent reactions of the Calvin cycle were either downregulated or not differentially expressed during the peak stress response. Most returned to baseline expression levels or became slightly upregulated at 4 hours. However, the gene necessary for the two-step conversion of glyceraldehyde-3-phosphate to fructose-6-phosphate, a key intermediate for both glycolysis and chitin biosynthesis, was upregulated during the peak response.

#### Glycolysis and gluconeogenesis

Generally, the genes repressed at 60 minutes were enriched for glycolysis and gluconeogenesis (Figure 3b). However, there were notable exceptions, including a number of paralogous genes in the pathway showing induction (Figure 4a). Thus, to better understand whether flux through glycolysis and gluconeogenesis is being regulated during hypoosmotic stress, we examined the rate-limiting and irreversible steps of each pathway. For glycolysis, the gene encoding phosphofructokinase-1 was not differentially expressed at either 60 minutes or 10 hours. The gene encoding pyruvate kinase, which is responsible for the last step in glycolysis, was repressed at 60 minutes, before recovering and ultimately becoming induced at 10 hours. For gluconeogenesis, at 60 minutes one paralog encoding the dedicated enzyme fructose 1,6-bisphosphatase was moderately repressed, while the other paralog was strongly induced. Paralogs encoding fructose-bisphosphate aldolase, the enzyme responsible for the gene directly before fructose 1,6-bisphosphatase, showed the same pattern of up- and downregulation. Differential expression of the genes encoding fructose 1,6-bisphosphatase and fructose-bisphosphate aldolase could be due to their shared role in the Calvin cycle, though no other Calvin cycle genes showed similar expression patterns. Instead, we predict that the accumulation of fructose-6-phosphate increases flux towards chitin biosynthesis (see below).

#### Chitin biosynthesis

The enrichments of (i) carbohydrate metabolic processes and carbohydrate binding at 60 minutes, (ii) carbohydrate binding at 10 hours (Figure 3b, d), and (iii) carbohydrate metabolism and chitin binding in the “delayed repression” Cluster 9 (Figure 2d) all highlight chitin biosynthesis as an important part of the response to hypoosmotic stress. The entire pathway is reversible and initiated by the breakdown of fructose-6-phosphate molecules into glucosamine by glutamine-fructose-6-phosphate transaminase (Figure 4) (Traller et al., 2016). Genes involved in chitin metabolism were largely upregulated at 60 minutes, as well as the gene responsible for its conversion into the more fibrous chitosan molecule (Figure 4a). Notably, all four paralogs for chitin synthase were strongly upregulated at 30 minutes (Figure S4). Given these expression patterns, combined with the predicted accumulation of the key intermediate fructose-6-phosphate, it is likely that chitin biosynthesis is being favored over degradation. We hypothesize that increased chitin and chitosan production might be important for cell wall remodeling during acute hypoosmotic stress (see Discussion).

### The gene expression response to acute hypoosmotic stress is distinct compared to long-term acclimation to low salinity

To understand whether the genes that were important for the acute hyposaline response were also important for long-term acclimation, we re-analyzed previously published RNA-seq data for the same *C. cryptica* CCMP332 strain that had been grown in freshwater for 120 days (Nakov et al., 2020). Of the 1,220 genes in the post-acclimation dataset that were significantly differentially expressed at least 2-fold, only 259 of them were also differentially expressed at any time point in the short-term stress experiment, representing a greater than 2.3-fold under-enrichment (*P* < 4.9 × 10^−99^, Fisher’s exact test). Although there was significant overlap between genes differentially expressed at the peak 60-minute time point and post-acclimated cells, the effect was small (1.03-fold enrichment, *P* < 1.1 × 10^−6^). Moreover, the majority of those genes showed opposing directions of differential expression in the acute versus acclimated responses. There was also little overlap between the genes responding to hypoosmotic stress at 10 hours compared to those at 120 days, with just 80 of 1,431 differential expressed genes at 10 hours overlapping with the 1,220 differential expressed genes at 120 days. Again, the majority of these had opposing expression directions at 10 hours compared to acclimated cells at 120 days. For example, many genes involved in photosynthesis transitioned from being repressed during the acute response to upregulated in fully acclimated cells. Among the few genes with similar expression patterns during the acute and acclimated responses, the majority were genes with unknown function.

Although we observed little overlap in the identity of genes that were differentially expressed during both the acute (0–10 hours) and acclimated (120 days) responses, it was still possible that the same functional groups were expressed at both timescales. Enrichment analysis showed no overlap in GO terms for genes involved in the acute versus acclimated response. In acclimated cells at 120 days, phosphorelay signal transduction was generally repressed, including several strongly repressed genes involved in histidine kinase pathways. Histidine kinases are used by both prokaryotes and eukaryotes as environmental sensors (Abriata et al., 2017), so downregulating these pathways may be a common characteristic of acclimated cells. In contrast, genes involved in DNA replication, DNA methylation, and mRNA splicing were all upregulated in post-acclimated cells, which may reflect post-acclimation differences in regulation of cell division (Table S3). Despite the caveat that enrichment analysis on the long-term acclimation data may be underpowered due to the absence of replicates, the complete lack of overlap combined with the novel enrichments for the long-term dataset are striking. Overall, this comparison suggests that the genes and processes necessary for the acute hypoosmotic stress response and long-term acclimation to freshwater are largely distinct.

## Discussion

The overall goal of this study was to understand how a euryhaline diatom responds to the stress of being immediately immersed in freshwater, a treatment that simulates a dispersal event or a rapid environmental fluctuation. The effects on the transcriptome were rapid and dramatic—more than 21% of the genes in the genome were differentially expressed within 60 minutes (Figure 2a), and over half of the genes were differentially expressed for at least one time point. The types of genes most impacted by hypoosmotic stress provide new insights into how diatoms maintain homeostasis during an abrupt salinity shift, which has implications for understanding how diatoms have repeatedly diversified into freshwaters and, more pressingly, how they will respond to changes in ocean salinity that are predicted to occur in response to climate change.

During acute hypoosmotic stress, cells face an influx of water and increased turgor pressure that risks rupturing the cell membrane, as well as an efflux of ions and other osmolytes that play important roles in cellular metabolism. In such situations, water channels (aquaporins) allow for rapid diffusion of water and sometimes other small molecules such as glycerol (Jahn et al., 2004; Matsui et al., 2018; S. D. Tyerman et al., 2002). Because aquaporins are known to play an important role in osmoadaptation (Tanghe et al., 2006), we examined genes encoding aquaporins for differential expression. Though none were significantly differentially expressed at two consecutive time points, one was strongly induced at 30 minutes (Figure S2). This was somewhat counter-intuitive, as high aquaporin expression is associated with increased water influx and potential cell lysis during hypoosmotic stress (Booth & Louis, 1999; Calamita, 2000). However, induction of an aquaporin during hypoosmotic stress was also identified recently in another diatom species (Pinseel et al., 2022), and aquaporins may have important roles beyond water transport including being involved in signal transduction via osmosensing (Stephen D. Tyerman et al., 2021; S. D. Tyerman et al., 2002).

Cells can also mitigate the effects of water influx by adjusting the concentrations of internal osmolytes to restore osmotic balance. We identified two upregulated chloroplastic K^+^ efflux pumps, which may help maintain K^+^ concentrations in the chloroplast to maintain proper turgor pressure and photosynthetic efficiency (Kunz et al., 2014; Sheng et al., 2014). Increased osmotic pressure within the cell also triggered the upregulation of a mechanosensitive anion transporter in the plasma membrane during 30 minutes to 3 hours. However, the most striking change in expression occurred for cell membrane ion transporters. We identified 17 induced Na^+^ exchangers and pumps and 15 induced K^+^ channels, suggesting the cell exports Na^+^ and H^+^ ions and imports K^+^ to maintain ion homeostasis (Table S3). An additional 66 K^+^ or Na^+^ transporter genes were differentially expressed at only a single time point, which may indicate that genes required to maintain ion homeostasis change throughout the hypoosmotic response. Maintaining optimal intracellular ion levels while acclimating to acute hypoosmotic stress is a major cellular challenge, which *C. cryptica* responds to with a finely-tuned response reflected through the complicated regulation of these myriad transporters throughout the time course. Notably, many channels involved in homeostasis during osmotic stress are mechanosensitive (Kirst, 1990; Kung et al., 2010), so the cell was likely tuning not only expression levels, but also post-translational regulation of these transporters to restore internal ion homeostasis following the initial shock.

Examination of genes involved in osmolyte homoeostasis revealed key differences between the acute response to hypoosmotic stress (this study) versus months-long acclimation (Nakov et al., 2020). Glycine-betaine, taurine, and DMSP were all identified as likely important osmolytes for long-term acclimation to freshwater in *C. cryptica*, but we found evidence for different patterns of osmolyte modulation during the short-term acute response. For long-term acclimation, levels of taurine were predicted to decrease, while our short-term data showed that taurine degradation genes are repressed, which would increase taurine levels in the cell. Additionally, unlike for long-term acclimation to freshwater, we found no evidence for differential expression of genes involved in DMSP and glycine-betaine metabolism, suggesting their levels do not change during the acute response. In contrast to gene expression levels in *C. cryptica* following acclimation, genes involved in proline metabolism were the only ones differentially regulated during acute freshwater stress. The gene encoding the rate-limiting step of proline biosynthesis (delta-1-pyrroline-5-carboxylate synthetase) was strongly repressed at 30–60 minutes. A similar pattern of repression was also seen for proline iminopeptidase, an enzyme that breaks down proline-containing peptides into the amino acid constituents (Figure S2). Based on previous carbon-labeling experiments (M. S. Liu & Hellebust, 1974; Ming Sai Liu & Hellebust, 1976), remaining “free” proline is likely being incorporated into proteins during acute hypoosmotic stress, rather than being excreted. Because proline is thought to be the main diatom osmolyte that changes in abundance during osmotic shock (Kirst, 1990; Krell et al., 2007), it was surprising at the time to find no changes in proline metabolic gene expression during long-term acclimation (Nakov et al., 2020). Our data suggests that a decrease in proline levels is only critical for the acute phase of the response, which sheds important light on this discrepancy in the literature and underscores the importance of considering different timescales for highly dynamic responses. Taken together, our analysis indicates that the osmolytes necessary during acute hypoosmotic stress are likely different from the ones that are important for long-term acclimation.

The hypoosmotic stress response of *C. cryptica* shared features of “general” environmental stress responses seen in other eukaryotes (Gasch et al., 2000; Mager & De Kruijff, 1995), including repression of genes involved in cell growth (e.g., ribosome biogenesis and RNA metabolism) and induction of genes canonically involved in stress defense such as ROS scavengers and HSP chaperones. Like other prokaryotic and eukaryotic microbes (Rojas et al., 2017; Sharfstein et al., 2007; Warner, 1999), diatoms halt cell division in the early stages of osmotic shock until ionic and osmotic equilibria are restored. Under such conditions, energy is likely redirected to cellular processes essential for acute stress management, rather than energetically expensive processes related to cell growth such as ribosome biogenesis (Albert et al., 2019). In yeast, transcriptional repression during stress appears important for redirecting translational capacity towards induced mRNAs, thereby accelerating production of induced proteins (Ho et al., 2018). Repression of ribosomal transcripts parallels transient growth arrest during stress, and the lack of concomitant ribosomal pro tein repression suggests that the steady-state level of ribosomes per cell remains similar (Ho et al., 2018; M. V. Lee et al., 2011). Ribosomal transcript repression has also been hypothesized as a mechanism for maintaining proteostasis while cells adapt to the stressful environment and express stress defense genes (Albert et al., 2019). We also observed a similar downregulation of transcripts for metabolic genes including the Krebs cycle, Calvin cycle, and glycolysis. Because transient growth arrest during stress in yeast leads to a poor correlation between mRNA repression and protein repression (M. V. Lee et al., 2011; Storey et al., 2020), caution is warranted when interpreting mRNA downregulation in the absence of proteomics.

Acute hypoosmotic stress resulted in the induction of many oxidative stress response genes, suggesting that hypoosmotic stress may increase ROS levels. In addition to increased induction of genes encoding direct ROS scavengers (i.e., superoxide dismutase, ascorbate peroxidase, and catalase), other oxidative stress response genes were also induced. Many genes that encode light-harvesting pigment-binding proteins were upregulated throughout the time course, and these proteins are known to mitigate oxidative stress in addition to their roles in photosynthesis (Latowski et al., 2011). We also observed upregulation of genes that are expected to increase the concentration of xanthophyll pigments in the cell. These pigments not only function as accessory light-harvesting pigments, but also protect the cell from photo-oxidative damage (Latowski et al., 2011). Polyamine biosynthetic genes were also induced, and polyamines are known to protect against oxidative damage of DNA (Ha et al., 1998). While there is precedent for hypoosmotic stress causing secondary ROS accumulation (Lyon et al., 2016), it is also possible that hypoosmotic stress in *C. cryptica* triggers a general stress response. In that case, induction of oxidative stress defense genes during hypoosmotic stress may reflect shared upstream signaling networks and not hypoosmotic stress causing oxidative stress per se. An open question in diatom stress biology is whether different species possess a common, generalized stress response, or whether individual stresses all lead to unique, specific responses. General stress responses are hypothesized to play a role in cross-stress protection (Berry & Gasch, 2008; McDaniel et al., 2018; Rangel, 2011). Future studies should examine the transcriptomic response of *C. cryptica* under different stress conditions, and whether hypoosmotic stress cross protects against other stressors such as elevated temperature or light.

Another striking feature of the hypoosmotic response was the regulation of many genes related to metabolism, and in particular chitin biosynthesis (Figure 4, Table S9). Chitin is a long-chain polysaccharide notable for playing an analogous role to plant cellulose as the major component of fungal cell walls. Chitin is also found in the cell membrane in a number of different eukaryotic lineages, where it supports skeletal and cell wall structures (Durkin et al., 2009). *Cyclotella* species, including *C. cryptica*, produce β-chitin, which is extruded as long crystalin threads from the specialized pores around the margin of the cell (LeDuff & Rorrer, 2019; McLachlan & Craigie, 1966). Initially, it was thought Thalassiosirales (which includes *Cyclotella*) was the only diatom lineage capable of producing chitin (McLachlan & Craigie, 1966), but chitin synthases were subsequently identified in the genome of a distantly related diatom, *Phaeodactylum tricornutum* (Kroth et al., 2008). In *Cyclotella nana*, chitin synthases are upregulated in response to different abiotic stressors that trigger elongated cell wall morphology (Davis et al., 2005; Mock et al., 2008). Increased chitin production could strengthen the cell wall in response to increased intracellular turgor pressure. Extracellular chitin threads also promote increased buoyancy in the water column (Chiriboga N & Rorrer, 2017; Morin et al., 1986). This might help diatoms optimize environmental light and temperature conditions (Boyd & Gradmann, 2002; Angela Falciatore & Bowler, 2002; A. Falciatore et al., 2000), so one hypothesis is that diatoms also regulate buoyancy in response to salinity stress. Future studies on the role of chitin in diatom salinity responses are clearly warranted.

Notably, two-thirds of 21,250 thousand predicted genes in the *C. cryptica* genome have no GO annotation, and a large fraction of those were differentially expressed in at least one time point (48%) or at least two consecutive time points (12%). These latter genes in particular likely reflect important, but not yet annotated, processes that play important roles in the response to hypoosmotic stress by diatoms. Understanding the function of uncharacterized genes is clearly a great remaining challenge for diatom biology.

The dynamics of the transcriptional response, combined with growth analysis and comparison to long-term acclimation, allows a better understanding of how transcriptional remodeling unfolds as marine- or brackish-water diatoms acclimate to freshwater. Notably, the majority of genes with peak differential expression in the early minutes and hours following freshwater exposure returned to baseline levels within 10 hours, coinciding with recovery of growth and the initial stages of freshwater acclimation. The growth rate of the newly acclimated cells was nevertheless lower than that of cells growing in the brackish control, and would likely extend for months if not in perpetuity (Nakov et al., 2020). Stress response studies in diatoms have covered a broad range of timescales (Borowitzka, 2018), with some studies solely looking at short term responses (minutes to hours) while others examine long-term responses in the range of days to weeks (Branco et al., 2010; Cheng et al., 2014; Krell et al., 2008; Krell et al., 2007; Nymark et al., 2013; Rijstenbil et al., 1989; Smith et al., 2016; Wang & Wang, 2008). Comparison of the acute expression response to hypoosmotic stress in this study to that of months-long freshwater acclimated cells from Nakov et al. 2020 revealed a striking lack of overlap, suggesting that short-term and long-term stress acclimated responses are largely distinct. This analysis also raises questions about what is happening at the gene expression level between 10 hours and 120 days, and further studies on genomic remodeling during medium- to long-term acclimation are clearly warranted. Nonetheless, our analyses strongly imply that the genes necessary for acute stress survival and tolerance during the early stages of growth acclimation are distinct from the genes necessary for long-term, stable growth. This study highlights how more finely resolved time courses improve our power to identify the genes important for an environmental response, while providing critical information about the processes important at different stages of acclimation.

## Acknowledgements

This material is based upon work supported by grants from the Simons Foundation (403249 to AJA and 725407 to EP) and the National Science Foundation (DEB-1651087 to AJA and MCB-1941824 to JAL), and multiple grants from the Arkansas Biosciences Institute (Arkansas Settlement Proceeds Act of 2000). EP also benefited from postdoctoral fellowships from Fulbright Belgium and the Belgian American Educational Foundation (BAEF). This research used resources available through the Arkansas High Performance Computing Center, which is funded through multiple NSF grants and the Arkansas Economic Development Commission.

## Data Accessibility

All RNA-seq data are available through the National Institutes of Health Gene Expression Omnibus (GEO) under accession number GSE206725. Raw and processed data plus the scripts needed to reproduce all analyses and figures are available as both “Supporting Information” and from Zenodo (https://doi.org/10.5281/zenodo.6704206).

## Author Contributions

KMD, KJJ, AJA, and JAL conceived and designed the study. KMD, KJJ conducted the experiments. KMD, KJJ, and JAL performed data analysis. EP, AJA, and JAL supervised the study. KMD wrote the original draft of the manuscript, and EP, AJA, and JAL edited the manuscript. All authors have read and approve the final manuscript.

## Supplemental Information

**FIGURE S1.**
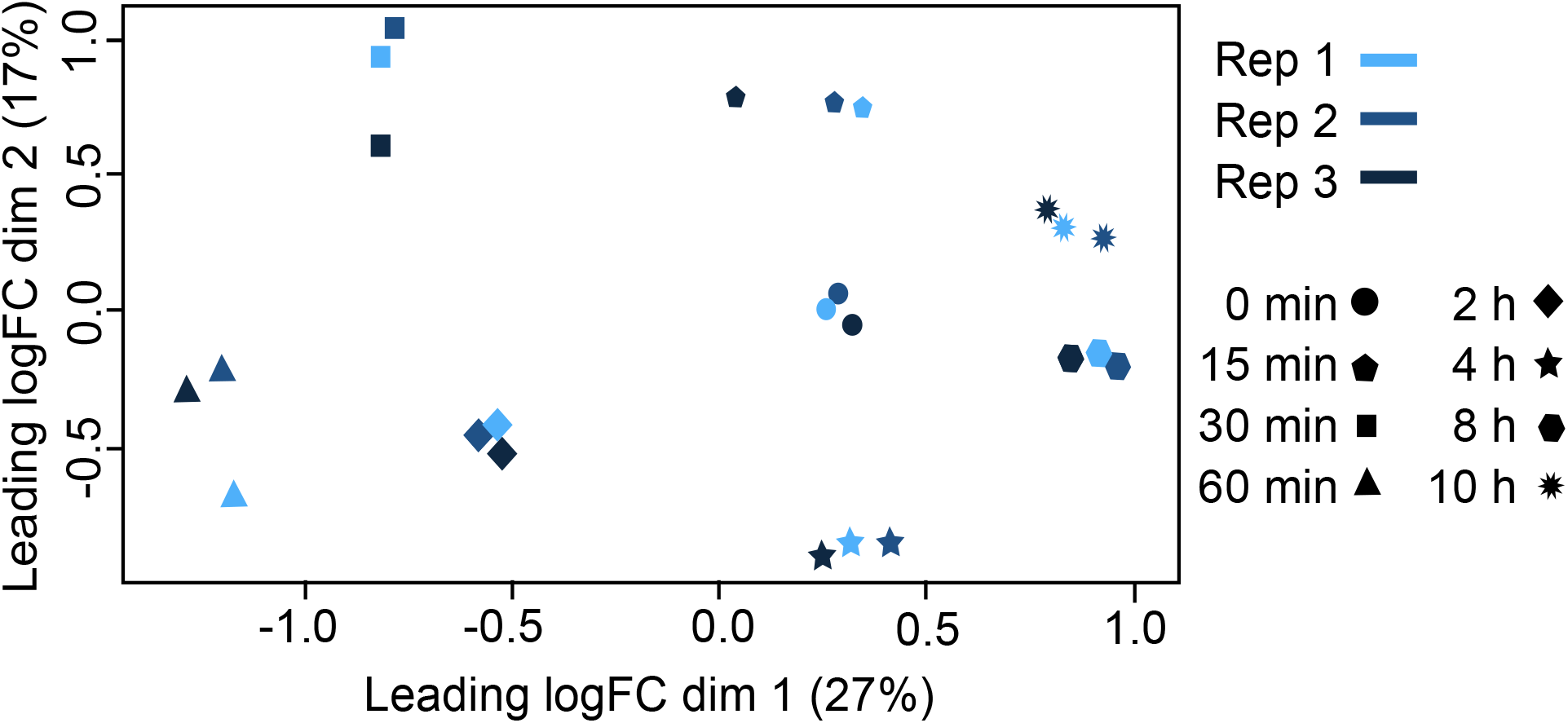
Multidimensional scaling (MDS) plot of the acute response to hypoosmotic stress. MDS analysis was performed on the logCPM for the top 500 genes with largest standard deviations across samples.

**FIGURE S2.**
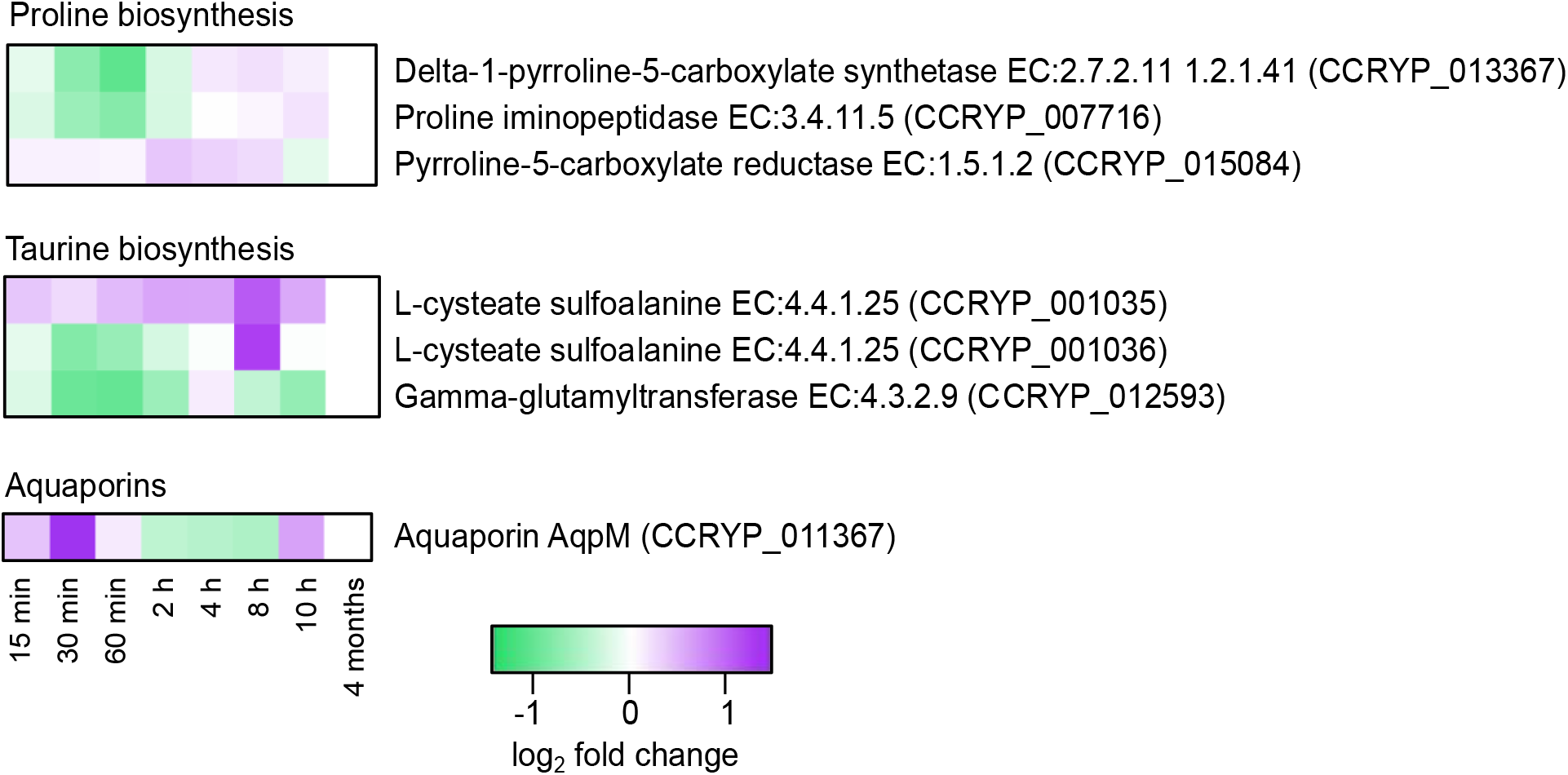
Expression dynamics of genes associated with osmotic regulation. Boxes denote log_2_ fold changes at each time point, with purple indicating induction and green indicating repression according to the key.

**FIGURE S3.**
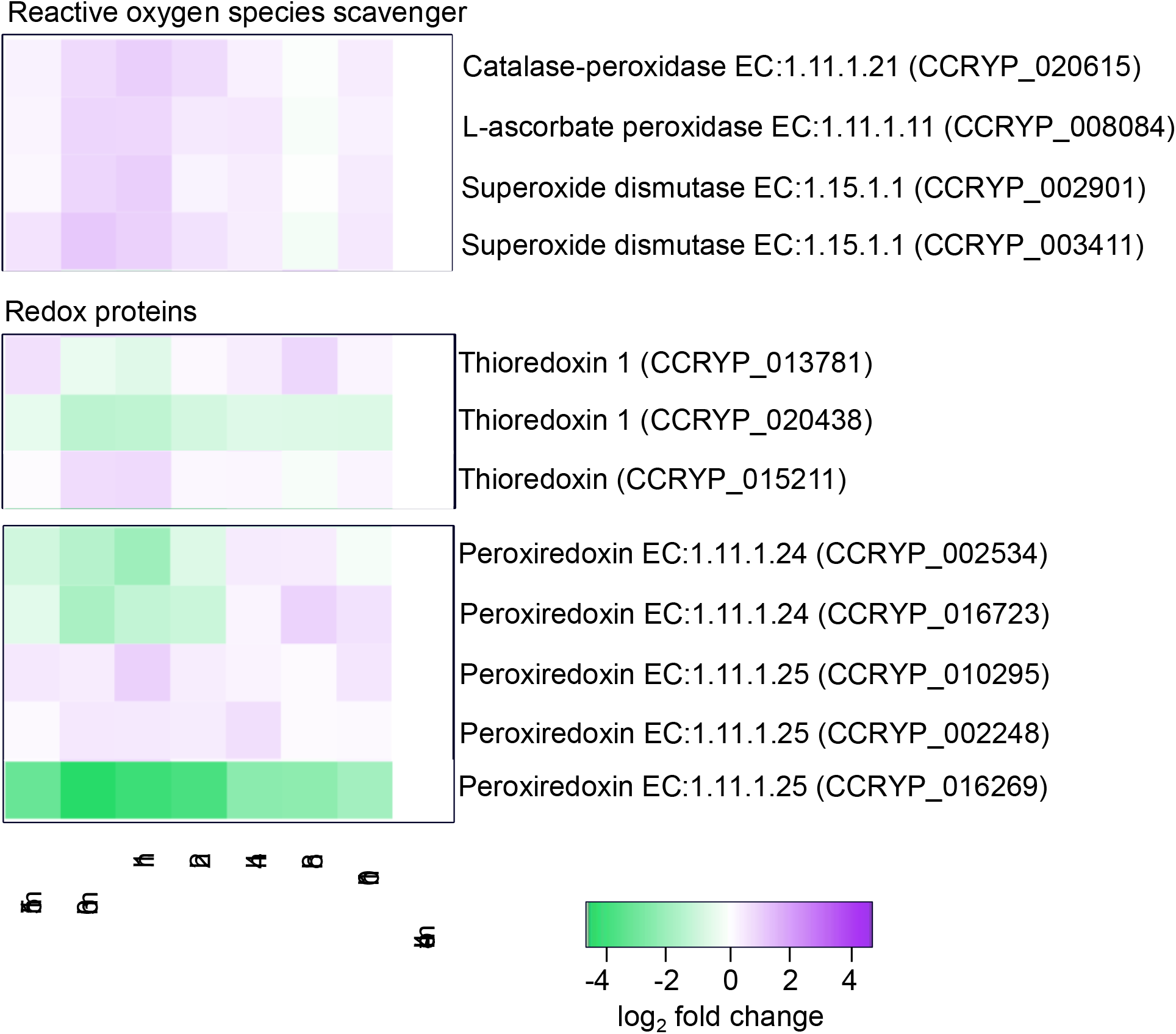
Expression dynamics of oxidative-stress defense related genes. Boxes denote log_2_ fold changes at each time point, with purple indicating induction and green indicating repression according to the key.

**FIGURE S4.**
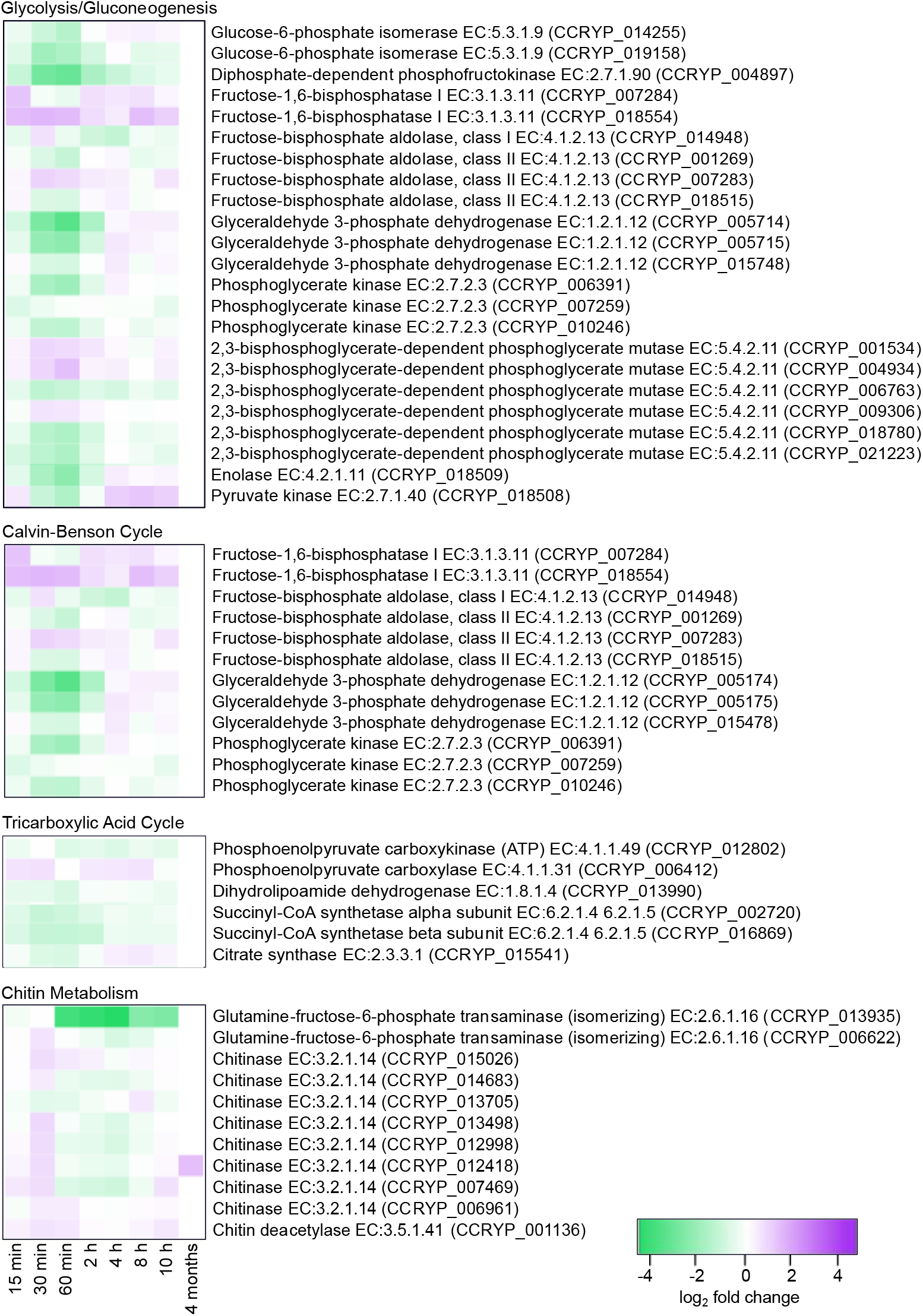
Expression dynamics of significantly differentially expressed metabolic genes noted in Figure 4. Boxes denote log_2_ fold changes at each time point, with purple indicating induction and green indicating repression according to the key.

**FIGURE S5.**
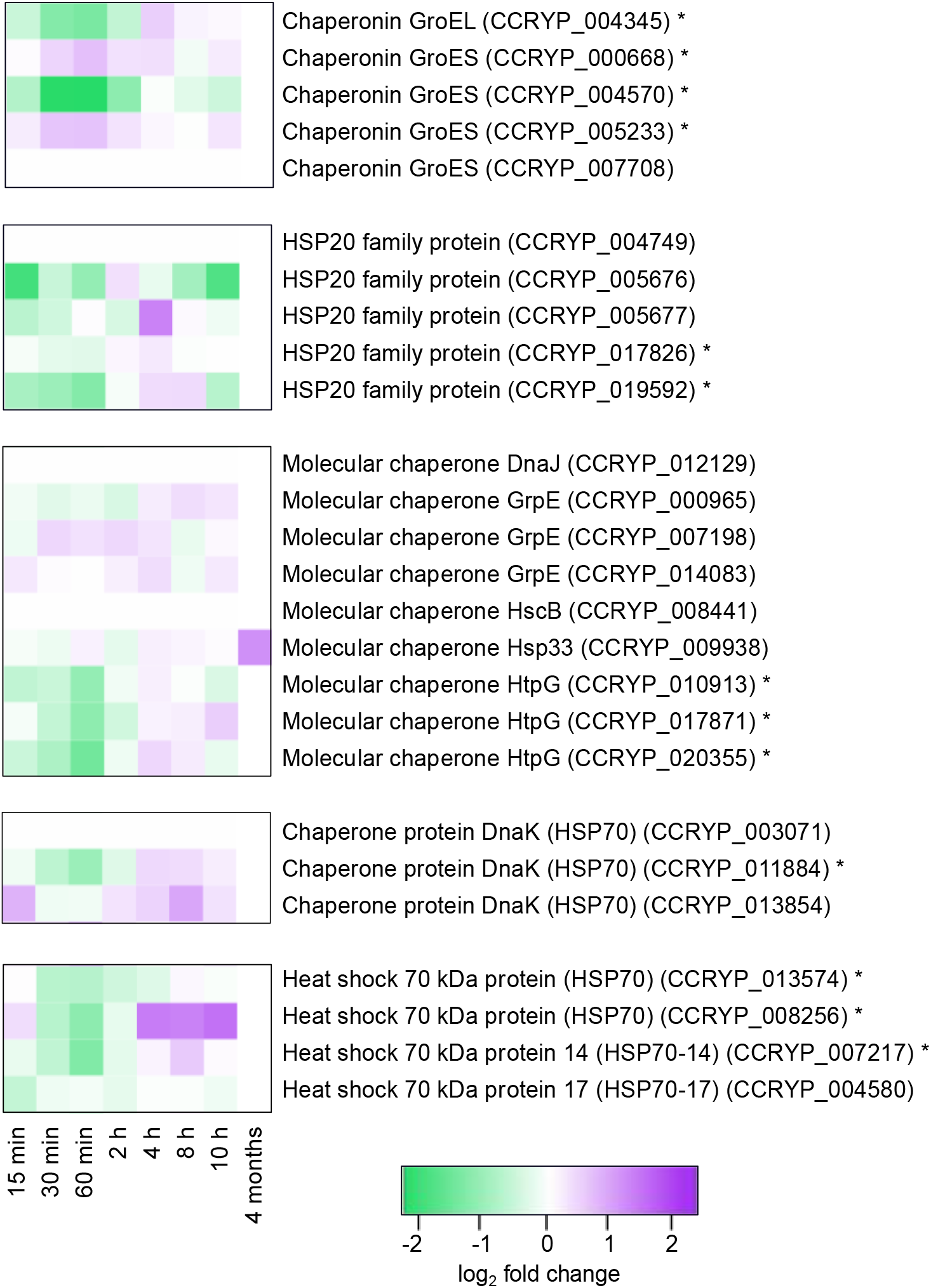
Expression dynamics of genes encoding heat shock protein (HSP) chaperones during hypoosmotic stress. Boxes denote log_2_ fold changes at each time point, with purple indicating induction and green indicating repression according to the key. Asterisks denote genes with significant differential expression for at least two time points.

## References

Abriata, L. A., Albanesi, D., Dal Peraro, M., & de Mendoza, D. (2017). Signal sensing and transduction by histidine kinases as unveiled through studies on a temperature sensor. Accounts of Chemical Research, 50(6), 1359–1366. doi:10.1021/acs.accounts.6b00593

Albert, B., Kos-Braun, I. C., Henras, A. K., Dez, C., Rueda, M. P., Zhang, X., Gadal, O., Kos, M., & Shore, D. (2019). A ribosome assembly stress response regulates transcription to maintain proteome homeostasis. Elife, 8, e45002. doi:10.7554/eLife.45002

Alexa, A., & Rahnenfuhrer, J. (2022). topGO: enrichment analysis for gene ontology. R package version 2.48.0.

Altschul, S. F., Gish, W., Miller, W., Myers, E. W., & Lipman, D. J. (1990). Basic local alignment search tool. Journal of Molecular Biology, 215(3), 403–410. doi:10.1016/S0022-2836(05)80360-2

Anders, S., Pyl, P. T., & Huber, W. (2014). HTSeq--a Python framework to work with high-throughput sequencing data. Bioinformatics, 31(2), 166–169. doi:10.1093/bioinformatics/btu638

Andrews, S. (2010). FastQC: a quality control tool for high throughput sequence data [Online]. Available online at: http://www.bioinformatics.babraham.ac.uk/projects/fastqc/.

Aramaki, T., Blanc-Mathieu, R., Endo, H., Ohkubo, K., Kanehisa, M., Goto, S., & Ogata, H. (2020). KofamKOALA: KEGG Ortholog assignment based on profile HMM and adaptive score threshold. Bioinformatics, 36(7), 2251–2252. doi:10.1093/bioinformatics/btz859

Armbrust, E. V. (2009). The life of diatoms in the world’s oceans. Nature, 459(7244), 185–192. doi:10.1038/nature08057

Balzano, S., Abs, E., & Leterme, S. C. (2015). Protist diversity along a salinity gradient in a coastal lagoon. Aquatic Microbial Ecology, 74(3), 263–277. doi:10.3354/ame01740

Berry, D. B., & Gasch, A. P. (2008). Stress-activated genomic expression changes serve a preparative role for impending stress in yeast. Molecular Biology of the Cell, 19(11), 4580–4587. doi:10.1091/mbc.e07-07-0680

Booth, I. R., & Louis, P. (1999). Managing hypoosmotic stress: Aquaporins and medianosensitive channels in *Escherichia coli*. Current Opinion in Microbiology, 2(2), 166–169. doi:10.1016/S1369-5274(99)80029-0

Borowitzka, M. A. (2018). The ‘stress’ concept in microalgal biology—homeostasis, acclimation and adaptation. Journal of Applied Phycology, 30(5), 2815–2825. doi:10.1007/s10811-018-1399-0

Boyd, C., & Gradmann, D. (2002). Impact of osmolytes on buoyancy of marine phytoplankton. Marine Biology, 141(4), 605–618. doi:10.1007/s00227-002-0872-z

Branco, D., Lima, A., Almeida, S. F. P., & Figueira, E. (2010). Sensitivity of biochemical markers to evaluate cadmium stress in the freshwater diatom *Nitzschia palea* (Kützing) W. Smith. Aquatic Toxicology, 99(2), 109–117. doi:10.1016/j.aquatox.2010.04.010

Calamita, G. (2000). The *Escherichia coli* aquaporin-Z water channel. Molecular Microbiology, 37(2), 254–262. doi:10.1046/j.1365-2958.2000.02016.x

Cheng, R.-L., Feng, J., Zhang, B.-X., Huang, Y., Cheng, J., & Zhang, C.-X. (2014). Transcriptome and gene expression analysis of an oleaginous diatom under different salinity conditions. BioEnergy Research, 7(1), 192–205. doi:10.1007/s12155-013-9360-1

Chiriboga N, O. G., & Rorrer, G. L. (2017). Control of chitin nanofiber production by the lipid-producing diatom *Cyclotella sp*. through fed-batch addition of dissolved silicon and nitrate in a bubble-column photobioreactor. Biotechnology Progress, 33(2), 407–415. doi:10.1002/btpr.2445

Conley, D. J., Kilham, S. S., & Theriot, E. (1989). Differences in silica content between marine and freshwater diatoms. Limnology and Oceanography, 34(1), 205–212. doi:10.4319/lo.1989.34.1.0205

Davis, A. K., Hildebrand, M., & Palenik, B. (2005). A stress-induced protein associated with the girdle band region of the diatom *Thalassiosira pseudonana* (bacillariophyta). Journal of Phycology, 41(3), 577–589. doi:10.1111/j.1529-8817.2005.00076.x

Dickson, D. M. J., & Kirst, G. O. (1987). Osmotic adjustment in marine eukaryotic algae: the role of inorganic ions, quaternary ammonium, tertiary sulphonium and carbohydrate solutes. New Phytologist, 106(4), 645–655. doi:10.1111/j.1469-8137.1987.tb00165.x

Dobin, A., & Gingeras, T. R. (2015). Mapping RNA-seq reads with STAR. Current Protocols in Bioinformatics, 51, 11.14.11–11.14.19. doi:10.1002/0471250953.bi1114s51

Durkin, C. A., Mock, T., & Armbrust, E. V. (2009). Chitin in diatoms and its association with the cell wall. Eukaryotic Cell, 8(7), 1038–1050. doi:10.1128/EC.00079-09

Eisen, M. B., Spellman, P. T., Brown, P. O., & Botstein, D. (1998). Cluster analysis and display of genome-wide expression patterns. Proceedings of the National Academy of Sciences, 95(25), 14863–14868. doi:10.1073/pnas.95.25.14863

Falciatore, A., & Bowler, C. (2002). Revealing the molecular secrets of marine diatoms. Annual Review of Plant Biology, 53, 109–130. doi:10.1146/annurev.arplant.53.091701.153921

Falciatore, A., d’Alcalà, M. R., Croot, P., & Bowler, C. (2000). Perception of environmental signals by a marine diatom. Science, 288(5475), 2363–2366. doi:10.1126/science.288.5475.2363

Gasch, A. P., Spellman, P. T., Kao, C. M., Carmel-Harel, O., Eisen, M. B., Storz, G., Botstein, D., & Brown, P. O. (2000). Genomic expression programs in the response of yeast cells to environmental changes. Molecular Biology of the Cell, 11(12), 4241–4257. doi:10.1091/mbc.11.12.4241

Godhe, A., Kremp, A., & Montresor, M. (2014). Genetic and microscopic evidence for sexual reproduction in the centric diatom Skeletonema marinoi. Protist, 165(4), 401–416. doi:10.1016/j.protis.2014.04.006

Ha, H. C., Yager, J. D., Woster, P. A., & Casero, R. A. (1998). Structural specificity of polyamines and polyamine analogues in the protection of DNA from strand breaks induced by reactive oxygen species. Biochemical and Biophysical Research Communications, 244(1), 298–303. doi:10.1006/bbrc.1998.8258

Heller, R., Manduchi, E., Grant, G. R., & Ewens, W. J. (2009). A flexible two-stage procedure for identifying gene sets that are differentially expressed. Bioinformatics, 25(8), 1019–1025. doi:10.1093/bioinformatics/btp076

Ho, Y.-H., Shishkova, E., Hose, J., Coon, J. J., & Gasch, A. P. (2018). Decoupling yeast cell division and stress defense implicates mRNA repression in translational reallocation during stress. Current Biology, 28(16), 2673–2680.e2674. doi:10.1016/j.cub.2018.06.044

Jahn, T. P., Møller, A. L. B., Zeuthen, T., Holm, L. M., Klaerke, D. A., Mohsin, B., Kühlbrandt, W., & Schjoerring, J. K. (2004). Aquaporin homologues in plants and mammals transport ammonia. FEBS Letters, 574(1-3), 31–36. doi:10.1016/j.febslet.2004.08.004

Kageyama, H., Tanaka, Y., & Takabe, T. (2018). Biosynthetic pathways of glycinebetaine in *Thalassiosira pseudonana;* functional characterization of enzyme catalyzing three-step methylation of glycine. Plant Physiology and Biochemistry, 127, 248–255. doi:10.1016/j.plaphy.2018.03.032

Kirst, G. O. (1990). Salinity tolerance of eukaryotic marine algae. Annual Review of Plant Physiology and Plant Molecular Biology, 41(1), 21–53. doi:10.1146/annurev.pp.41.060190.000321

Krell, A., Beszteri, B., Dieckmann, G., Glöckner, G., Valentin, K., & Mock, T. (2008). A new class of ice-binding proteins discovered in a salt-stress-induced cDNA library of the psychrophilic diatom *Fragilariopsis cylindrus* (Bacillariophyceae). European Journal of Phycology, 43(4), 423–433. doi:10.1080/09670260802348615

Krell, A., Funck, D., Plettner, I., John, U., & Dieckmann, G. (2007). Regulation of proline metabolism under salt stress in the psychrophilic diatom *Fragilariopsis cylindrus* (Bacillariophyceae). Journal of Phycology, 43(4), 753–762. doi:10.1111/j.1529-8817.2007.00366.x

Kroth, P. G., Chiovitti, A., Gruber, A., Martin-Jezequel, V., Mock, T., Parker, M. S., Stanley, M. S., Kaplan, A., Caron, L., Weber, T., Maheswari, U., Armbrust, E. V., & Bowler, C. (2008). A model for carbohydrate metabolism in the diatom *Phaeodactylum tricornutum* deduced from comparative whole genome analysis. PLoS One, 3(1), e1426. doi:10.1371/journal.pone.0001426

Kung, C., Martinac, B., & Sukharev, S. (2010). Mechanosensitive channels in microbes. Annual Review of Microbiology, 64, 313–329. doi:10.1146/annurev.micro.112408.134106

Kunz, H.-H., Gierth, M., Herdean, A., Satoh-Cruz, M., Kramer, D. M., Spetea, C., & Schroeder, J. I. (2014). Plastidial transporters KEA1, −2, and −3 are essential for chloroplast osmoregulation, integrity, and pH regulation in *Arabidopsis*. Proceedings of the National Academy of Sciences, 111(20), 7480–7485. doi:10.1073/pnas.1323899111

Latowski, D., Kuczyńska, P., & Strzałka, K. (2011). Xanthophyll cycle – a mechanism protecting plants against oxidative stress. Redox Report, 16(2), 78–90. doi:10.1179/174329211x13020951739938

LeDuff, P., & Rorrer, G. L. (2019). Formation of extracellular β-chitin nanofibers during batch cultivation of marine diatom *Cyclotella sp*. at silicon limitation. Journal of Applied Phycology, 31(6), 3479–3490. doi:10.1007/s10811-019-01879-6

Lee, C. E., Downey, K., Colby, R. S., Freire, C. A., Nichols, S., Burgess, M. N., & Judy, K. J. (2022). Recognizing salinity threats in the climate crisis. Integrative and Comparative Biology. doi:10.1093/icb/icac069

Lee, M. V., Topper, S. E., Hubler, S. L., Hose, J., Wenger, C. D., Coon, J. J., & Gasch, A. P. (2011). A dynamic model of proteome changes reveals new roles for transcript alteration in yeast. Molecular Systems Biology, 7, 514. doi:10.1038/msb.2011.48

Liu, M. S., & Hellebust, J. A. (1974). Uptake of amino acids by the marine centric diatom *Cyclotella cryptica*. Canadian Journal of Botany, 20(8), 1109–1118. doi:10.1139/m74-173

Liu, M. S., & Hellebust, J. A. (1976). Effects of salinity and osmolarity of the medium on amino acid metabolism in *Cyclotella cryptica*. Canadian Journal of Microbiology, 54(9), 938–948. doi:10.1139/b76-098

Lund, S. P., Nettleton, D., McCarthy, D. J., & Smyth, G. K. (2012). Detecting differential expression in RNA-sequence data using quasi-likelihood with shrunken dispersion estimates. Statistical Applications in Genetics and Molecular Biology, 11(5). doi:10.1515/1544-6115.1826

Lyon, B. R., Bennett-Mintz, J. M., Lee, P. A., Janech, M. G., & DiTullio, G. R. (2016). Role of dimethylsulfoniopropionate as an osmoprotectant following gradual salinity shifts in the sea-ice diatom *Fragilariopsis cylindrus*. Environmental Chemistry, 13(2), 181–194. doi:10.1071/EN14269

Lyon, B. R., Lee, P. A., Bennett, J. M., DiTullio, G. R., & Janech, M. G. (2011). Proteomic analysis of a sea-ice diatom: salinity acclimation provides new insight into the dimethylsulfoniopropionate production pathway. Plant Physiology, 157(4), 1926–1941. doi:10.1104/pp.111.185025

Mager, W. H., & De Kruijff, A. J. (1995). Stress-induced transcriptional activation. Microbiology Reviews, 59(3), 506–531. doi:10.1128/mr.59.3.506-531.1995

Matsui, H., Hopkinson, B., Nakajima, K., & Matsuda, Y. (2018). Plasma-membrane-type aquaporins from marine diatoms function as CO2/NH3 channels and provide photoprotection. Plant Physiology, 178(1). doi:10.1104/pp.18.00453

Mayer, C., Bierhoff, H., & Grummt, I. (2005). The nucleolus as a stress sensor: JNK2 inactivates the transcription factor TIF-IA and down-regulates rRNA synthesis. Genes & Development, 19(8), 933–941. doi:10.1101/gad.333205

McDaniel, E. A., Stuecker, T. N., Veluvolu, M., Gasch, A. P., & Lewis, J. A. (2018). Independent mechanisms for acquired salt tolerance versus growth resumption induced by mild ethanol pretreatment in *Saccharomyces cerevisiae*. mSphere, 3(6), e00574–00518. doi:10.1128/mSphere.00574-18

McLachlan, J., & Craigie, J. S. (1966). Chitan fibres in *Cyclotella cryptica* and growth of *C. cryptica* and *Thalassiosira fluviatilis*. Some contemporary studies in marine science, 511–517.

Mock, T., Samanta, M. P., Iverson, V., Berthiaume, C., Robison, M., Holtermann, K., Durkin, C., BonDurant, S. S., Richmond, K., Rodesch, M., Kallas, T., Huttlin, E. L., Cerrina, F., Sussman, M. R., & Armbrust, E. V. (2008). Whole-genome expression profiling of the marine diatom *Thalassiosira pseudonana* identifies genes involved in silicon bioprocesses. Proceedings of the National Academy of Sciences, 105(5), 1579–1584. doi:10.1073/pnas.0707946105

Morin, L. G., Smucker, R. A., & Herth, W. (1986). Effects of two chitin synthesis inhibitors on *Thalassiosira fluviatilis* and *Cyclotella cryptica*. FEMS Microbiology Letters, 37(3), 263–268. doi:10.1111/j.1574-6968.1986.tb01806.x

Nakov, T., Beaulieu, J. M., & Alverson, A. J. (2018). Insights into global planktonic diatom diversity: The importance of comparisons between phylogenetically equivalent units that account for time. The ISME Journal, 12(11), 2807–2810. doi:10.1038/s41396-018-0221-y

Nakov, T., Judy, K. J., Downey, K. M., Ruck, E. C., & Alverson, A. J. (2020). Transcriptional response of osmolyte synthetic pathways and membrane transporters in a euryhaline diatom during long-term acclimation to a salinity gradient. Journal of Phycology.

Nitta, M., Okamura, H., Aizawa, S., & Yamaizumi, M. (1997). Heat shock induces transient p53-dependent cell cycle arrest at G1/S. Oncogene, 15(5), 561–568. doi:10.1038/sj.onc.1201210

Nymark, M., Valle, K. C., Hancke, K., Winge, P., Andresen, K., Johnsen, G., Bones, A. M., & Brembu, T. (2013). Molecular and photosynthetic responses to prolonged darkness and subsequent acclimation to re-illumination in the diatom *Phaeodactylum tricornutum*. PLoS One, 8(3), e58722. doi:10.1371/journal.pone.0058722

Papenbrock, J., Mock, H. P., Tanaka, R., Kruse, E., & Grimm, B. (2000). Role of magnesium chelatase activity in the early steps of the tetrapyrrole biosynthetic pathway. Plant Physiology, 122(4), 1161–1169. doi:10.1104/pp.122.4.1161

Pinseel, E., Nakov, T., Van den Berge, K., Downey, K. M., Judy, K. J., Kourtchenko, O., Kremp, A., Ruck, E. C., Sjöqvist, C., Töpel, M., Godhe, A., & Alverson, A. J. (2022). Strain-specific transcriptional responses overshadow salinity effects in a marine diatom sampled along the Baltic Sea salinity cline. The ISME Journal, 16(7), 1776–1787. doi:10.1038/s41396-022-01230-x

Pörtner, H.-O., Roberts, D. C., Masson-Delmotte, V., Zhai, P., Tignor, M., Poloczanska, E., & Weyer, N. M. (2019). The ocean and cryosphere in a changing climate. IPCC Special Report on the Ocean and Cryosphere in a Changing Climate.

Rangel, D. E. N. (2011). Stress induced cross-protection against environmental challenges on prokaryotic and eukaryotic microbes. World Journal of Microbiology and Biotechnology, 27(6), 1281–1296. doi:10.1007/s11274-010-0584-3

Reimann, B. E. F., Lewin, J. M. C., & Guillard, R. R. L. (1963). *Cyclotella cryptica*, a new brackish-water diatom species. Phycologia, 3(2), 75–84. doi:10.2216/i0031-8884-3-2-75.1

Rijstenbil, J. W., Wijnholds, J. A., & Sinke, J. J. (1989). Implications of salinity fluctuation for growth and nitrogen metabolism of the marine diatom *Ditylum brightwellii* in comparison with *Skeletonema costatum*. Marine Biology, 101(1), 131–141. doi:10.1007/BF00393486

Ritchie, M. E., Phipson, B., Wu, D., Hu, Y., Law, C. W., Shi, W., & Smyth, G. K. (2015). limma powers differential expression analyses for RNA-sequencing and microarray studies. Nucleic Acids Research, 43(7), e47. doi:10.1093/nar/gkv007

Roberts, W. R., Downey, K. M., Ruck, E. C., Traller, J. C., & Alverson, A. J. (2020). Improved reference genome for *Cyclotella cryptica* CCMP332, a model for cell wall morphogenesis, salinity adaptation, and lipid production in diatoms (Bacillariophyta). G3 (Bethesda), 10(9), 2965–2974. doi:10.1534/g3.120.401408

Robinson, M. D., McCarthy, D. J., & Smyth, G. K. (2010). edgeR: a Bioconductor package for differential expression analysis of digital gene expression data. Bioinformatics, 26(1), 139–140. doi:10.1093/bioinformatics/btp616

Rojas, E. R., Huang, K. C., & Theriot, J. A. (2017). Homeostatic cell growth is accomplished mechanically through membrane tension inhibition of cell-wall synthesis. Cell Systems, 5(6), 578–590. doi:10.1016/j.cels.2017.11.005

Schultz, M. E. (1971). Salinity-related polymorphism in the brackish-water diatom *Cyclotella cryptica*. Canadian Journal of Botany, 49(8), 1285–1289. doi:10.1139/b71-182

Schultz, M. E., & Trainor, F. R. (1970). Production of male gametes and auxospores in a polymorphic clone of the centric diatom *Cyclotella*. Canadian Journal of Botany, 48(5), 947–951. doi:10.1139/b70-133

Seaton, D. D., & Krishnan, J. (2016). Model-based analysis of cell cycle responses to dynamically changing environments. PLOS Computational Biology, 12(1), e1004604. doi:10.1371/journal.pcbi.1004604

Sharfstein, S. T., Shen, D., Kiehl, T. R., & Zhou, R. (2007). Molecular response to osmotic shock. In Systems Biology (pp. 213–236).

Sheng, P., Tan, J., Jin, M., Wu, F., Zhou, K., Ma, W., Heng, Y., Wang, J., Guo, X., Zhang, X., Cheng, Z., Liu, L., Wang, C., Liu, X., & Wan, J. (2014). *Albino midrib* 1, encoding a putative potassium efflux antiporter, affects chloroplast development and drought tolerance in rice. Plant Cell Reports, 33(9), 1581–1594. doi:10.1007/s00299-014-1639-y

Shu, Q., Qiao, F., Song, Z., Zhao, J., & Li, X. (2018). Projected freshening of the arctic ocean in the 21st century. J. Geophys. Res. C: Oceans, 123(12), 9232–9244. doi:10.1029/2018jc014036

Skirycz, A., Claeys, H., De Bodt, S., Oikawa, A., Shinoda, S., Andriankaja, M., Maleux, K., Eloy, N. B., Coppens, F., Yoo, S.-D., Saito, K., & Inzé, D. (2011). Pause-and-stop: the effects of osmotic stress on cell proliferation during early leaf development in *Arabidopsis* and a role for ethylene signaling in cell cycle arrest. The Plant Cell, 23(5), 1876–1888. doi:10.1105/tpc.111.084160

Smith, S. R., Gillard, J. T. F., Kustka, A. B., McCrow, J. P., Badger, J. H., Zheng, H., Ashley, M. N., Dupont, C. L., Obata, T., Fernie, A. R., & Allen, A. E. (2016). Transcriptional orchestration of the global cellular response of a model pennate diatom to diel light cycling under iron limitation. PLOS Genetics, 12(12), e1006490. doi:10.1371/journal.pgen.1006490

Storey, A. J., Hardman, R. E., Byrum, S. D., Mackintosh, S. G., Edmondson, R. D., Wahls, W. P., Tackett, A. J., & Lewis, J. A. (2020). Accurate and sensitive quantitation of the dynamic heat shock proteome using tandem mass tags. Journal of Proteome Research, 19(3), 1183–1195. doi:10.1021/acs.jproteome.9b00704

Sun, K. (2020). Ktrim: an extra-fast and accurate adapter- and quality-trimmer for sequencing data. Bioinformatics, 36(11), 3561–3562. doi:10.1093/bioinformatics/btaa171

Supek, F., Bošnjak, M., Škunca, N., & Šmuc, T. (2011). REVIGO summarizes and visualizes long lists of gene ontology terms. PLoS One, 6(7), e21800. doi:10.1371/journal.pone.0021800

Tanghe, A., Van Dijck, P., & Thevelein, J. M. (2006). Why do microorganisms have aquaporins? Trends in Microbiology, 14(2), 78–85. doi:10.1016/j.tim.2005.12.001

Tevatia, R., Allen, J., Rudrappa, D., White, D., Clemente, T. E., Cerutti, H., Demirel, Y., & Blum, P. (2015). The taurine biosynthetic pathway of microalgae. Algal Research, 9, 21–26. doi:10.1016/j.algal.2015.02.012

Theseira, A. M., Nielsen, D. A., & Petrou, K. (2020). Uptake of dimethylsulphoniopropionate (DMSP) reduces free reactive oxygen species (ROS) during late exponential growth in the diatom *Thalassiosira weissflogii* grown under three salinities. Marine Biology, 167(9). doi:10.1007/s00227-020-03744-4

Traller, J. C., Cokus, S. J., Lopez, D. A., Gaidarenko, O., Smith, S. R., McCrow, J. P., Gallaher, S. D., Podell, S., Thompson, M., Cook, O., Morselli, M., Jaroszewicz, A., Allen, E. E., Allen, A. E., Merchant, S. S., Pellegrini, M., & Hildebrand, M. (2016). Genome and methylome of the oleaginous diatom *Cyclotella cryptica* reveal genetic flexibility toward a high lipid phenotype. Biotechnology for Biofuels and Bioproducts, 9, 258. doi:10.1186/s13068-016-0670-3

Tyerman, S. D., McGaughey, S. A., Qiu, J., Yool, A. J., & Byrt, C. S. (2021). Adaptable and multifunctional ion-conducting aquaporins. Annual Review of Plant Biology, 72, 703–736. doi:10.1146/annurev-arplant-081720-013608

Tyerman, S. D., Niemietz, C. M., & Bramley, H. (2002). Plant aquaporins: multifunctional water and solute channels with expanding roles. Plant, Cell & Environment, 25(2), 173–194. doi:10.1046/j.0016-8025.2001.00791.x

Van Bergeijk, S. A., Van der Zee, C., & Stal, L. J. (2003). Uptake and excretion of dimethylsulphoniopropionate is driven by salinity changes in the marine benthic diatom *Cylindrotheca closterium*. European Journal of Phycology, 38(4), 341–349. doi:10.1080/09670260310001612600

Van den Berge, K., Soneson, C., Robinson, M. D., & Clement, L. (2017). stageR: a general stage-wise method for controlling the gene-level false discovery rate in differential expression and differential transcript usage. Genome Biology, 18(1), 151. doi:10.1186/s13059-017-1277-0

Wang, M.-J., & Wang, W.-X. (2008). Temperature-dependent sensitivity of a marine diatom to cadmium stress explained by subcelluar distribution and thiol synthesis. Environmental Science & Technology, 42(22), 8603–8608. doi:10.1021/es801470w

Warner, J. R. (1999). The economics of ribosome biosynthesis in yeast. Trends in Biochemical Sciences, 24(11), 437–440. doi:10.1016/s0968-0004(99)01460-7

West, G., Inzé, D., & Beemster, G. T. S. (2004). Cell cycle modulation in the response of the primary root of *Arabidopsis* to salt stress. Plant Physiology, 135(2), 1050–1058. doi:10.1104/pp.104.040022

