## Appendix S1 for "The dynamic response to hypoosmotic stress reveals distinct stages of freshwater acclimation by a euryhaline diatom": Appendix 1.html

- 1. Depiction of C.
  crytpica growth data
  - 1.1 Experimental design
    and methods
  - 1.2 Packages used in
    analyses
  - 1.3 Growth data for
    short term cultures
- 2. Bioinformatic
  analysis of RNA-seq reads
  - 2.1 Quality
    control, adapter removal & trimming
- 3. Statistical
  analysis of RNA-seq data in R
  - 3.1 Model fitting in EdgeR
    - 3.1.2 Data filtering
    - 3.1.3 Data normalization
    - 3.1.4 Design
      matrix and dispersion estimation
    - 3.1.5 Model fitting
  - 3.2
    Testing for differential expression using stage-wise analysis:
    - 3.2.1 Defining contrasts to
      test
    - 3.2.2 Stage-wise testing
    - 3.2.3
      Summarize the results for downstream analyses
    - 3.2.4 Subset to genes
      DE at 2 timepoints
    - 3.2.5
      Graph of genes up and downregulated at each timepoint
    - 3.2.6 Unique and
      shared DE between timepoints
- 4. Gene enrichment analyses
  of:
  - 4.1 Full dataset
  - 4.2
    Significantly differentially expressed genes
  - 4.3 Long-term data
  - 4.4
    Comparisons between peak stress, acclimation, and post-acclimation
    - 4.4.1. Comparing 60
      min and 10 h enrichments
    - 4.4.2
      GO enrichment for different LFC cutoffs for 1h and 10 h.
  - 4.5
    Clusters identified from TreeView (centroid linkage)
- 5.
  Combine data for KO, GO, uniprot, and cluster information with the LFC
  data for perusal.
- 6. Manuscript graphs
  - 6.1 Gene
    expression behavior in short- and long-term
  - 6.2
    Graph of median expression of genes DE at 1h and 10 h
  - 6.3 Heatmaps for cycles
    of interest
  - 6.4 Cluster plots
    - 6.4.1 Summary
      of the averages of all the clusters
    - 6.4.2
      Individual graphs showing cluster averages with error bars
- 7. Paralogs simulation

This document gives an overview of the data analysis associated with
the manuscript rapid hypo-osmotic stress response of the euryhaline
diatom *Cyclotella cryptica* to freshwater.

### 1. Depiction of C. crytpica growth data

#### 1.1 Experimental design and methods

- Species: *Cyclotella cryptica*, strain CCMP332, from National
  Center for Marine Algae and Microbiota
- Culture maintained in 1 L artificial seawater (ASW) at 24 ppt in 15
  C incubator, ~20 mol photons/m2/s irradiance
- Experimental design
  - Replicates: three 2 mL aliquots taken from source to inoculate
    flasks of 500 mL ASW 24 ppt, grown until exponential growth was
    reached
  - Inoculate volumes: 3 x 10^6 cells, estimated using 500 uL of
    replicates in FlowCam
  - Treatments
    - Stress media: 38 mL freshwater (0 ppt) with ASW nutrients in a 50 mL
      falcon tube
    - Time-series exposures: 0 minutes, 15 minutes, 30 minutes, 60
      minutes, 120 minutes, 240 minutes, 480 minutes, 600 minutes
    - Mock/Control of ASW 24 ppt treated in exactly the same way to
      confirm no substantial stress occurred at 0 minutes \*Time series
      collection
  - Collected volume required for inoculation in two 50 mL falcon tubes
    - Concentrated in centrifuge 4 C for 3 minutes at 3000 rpm
    - Supernatant decanted and cells combined in 18 mL of ASW 0 ppt (2 mL
      for each time point)
    - Mock/Control suspended in 2 mL ASW 24 ppt
    - All volumes transferred to treatment vessels in 15 C incubator,
      gentle agitation on rocker
  - Collection
    - Mock/Control and 0 minute collected immediately
    - Spun down at 1500 rpm for 3 minutes at 4 C
    - Supernatant decanted and pellet flash-frozen in liquid nitrogen
- RNA extraction: RNeasy kit, quality check with Qubit and
  Tapestation
- Library development: KAPA mRNA HyperPrep kit, libraries multiplexed
  for sequencing on Illumina HiSeq 4000
  - Pooled with *Skeletonema marinoi* sequences from a similar
    short-term stress experiment conducted simultaneously

#### 1.2 Packages used in analyses

```
setwd ("/My Drive/ShortTermStress-IlluminaData/FullExp/Cryp-DE/")

#Load libraries to be used in this project into a variable.

libraries <- c("DESeq2", "edgeR", "shiny", "shinythemes", "tidyverse", "lubridate",  "viridis", "ggthemr", "ggrepel", "rlang", "reshape", "naniar", "pheatmap","rhdf5", "cgwtools", "NMF", "RColorBrewer" ,"topGO", "genefilter", "PoiClaClu", "gplots", "statmod", "stageR", "data.table", "radiant.data", "modelr", "here")

#Command to load libraries all at once.  
lapply(libraries, require, character.only = TRUE)

#You can save yourself some time and heartache by making an RData file that saves all your variables. If you ever accidentally wipe your environment, or something crashes, loading the RData will import any variables you've saved along the way. Establish an RData file with ---save("directory/file_name.RData")---. Use the `resave` function that I have in a few sections to save new variables to the file without overwriting anything. `load` will import the variables into your environment.

#load("/My Drive/ShortTermStress-IlluminaData/FullExp/Cryp-DE/data/import_rmd_vars.RData")
```

#### 1.3 Growth data for short term cultures

We monitored growth of C. cryptica in freshwater over 7 days via
chlorophyll-a fluorescence and cell counts.

```
#Import data
growth_cells <- read.delim("/My Drive/Downey_Manuscripts/Ccryptica_STS/Ccryptica_STS_files/Growth_cells.txt", header = TRUE, sep = '\t', fill = TRUE, row.names=NULL)
growth_RFU <- read.delim("/My Drive/Downey_Manuscripts/Ccryptica_STS/Ccryptica_STS_files/Growth_RFU.txt",header = TRUE, sep = '\t', fill = TRUE, row.names=NULL)

#Growth data from cell counts obtained using FlowCam
growth_cells_df <- pivot_longer(growth_cells[ ,c(1:13)], -c(Replicate,Trt), names_to = "timept")
growth_cells_df$timept <- ordered(growth_cells_df$timept, levels=c("t0m","t15m","t30m","t1h","t2h","t4h","t8h","t10h","t12h","t24h","t7d"))

growth_cells_df <- growth_cells_df %>% mutate(timept = case_when(
  growth_cells_df$timept == "t0m"  ~ 0,
  growth_cells_df$timept == "t15m" ~ 15,
  growth_cells_df$timept == "t30m" ~ 30,
  growth_cells_df$timept == "t1h" ~ 60,
  growth_cells_df$timept == "t2h" ~ 120,
  growth_cells_df$timept == "t4h" ~ 240,
  growth_cells_df$timept == "t8h" ~ 480,
  growth_cells_df$timept == "t10h" ~ 600,
  growth_cells_df$timept == "t12h" ~ 720,
  growth_cells_df$timept == "t24h" ~ 1440,
  growth_cells_df$timept == "t7d" ~ 10080
))

growth_cells_data <- growth_cells_df %>% group_by(timept,Trt) %>% summarize(sd = sd(value, na.rm = TRUE),value = mean(value), )
```

```
## `summarise()` has grouped output by 'timept'. You can override using the
## `.groups` argument.
```

```
#Visualize
ggplot(growth_cells_data,  aes(x = timept, y = value, group = Trt, color = Trt, ymin = value-sd, ymax = value+sd))  + 
    geom_errorbar(width = 0.5) +
    geom_line(size = 1) +
    theme(axis.text.x = element_text(vjust=0, face = "bold"),
          axis.ticks.x=element_blank()) +
    coord_trans(y = "log") +
    xlab("Time in minutes") + 
    ylab("Cells per mL") +
    theme_classic() + 
    scale_color_manual(values=c("#112943", "#51b0fa")) +
    theme(plot.title = element_text(hjust = 0.5, face = "bold"))
```

```
#Growth data from relative fluorescent units measured with Trilogy
growth_RFU_df <- pivot_longer(growth_RFU[ ,c(1:13)], -c(Replicate,Trt), names_to = "timept")
growth_RFU_df$timept <- ordered(growth_RFU_df$timept, levels=c("t0m","t15m","t30m","t1h","t2h","t4h","t8h","t10h","t12h","t24h","t7d"))

growth_RFU_df <- growth_RFU_df %>% mutate(timept = case_when(
  growth_RFU_df$timept == "t0m" ~ 0,
  growth_RFU_df$timept == "t15m" ~ 15,
  growth_RFU_df$timept == "t30m" ~ 30,
  growth_RFU_df$timept == "t1h" ~ 60,
  growth_RFU_df$timept == "t2h" ~ 120,
  growth_RFU_df$timept == "t4h" ~ 240,
  growth_RFU_df$timept == "t8h" ~ 480,
  growth_RFU_df$timept == "t10h" ~ 600,
  growth_RFU_df$timept == "t12h" ~ 720,
  growth_RFU_df$timept == "t24h" ~ 1440,
  growth_RFU_df$timept == "t7d" ~ 10080
))

growth_RFU_data <- growth_RFU_df %>% group_by(timept,Trt) %>% summarize(sd = sd(value, na.rm = TRUE),value = mean(value), )
```

```
## `summarise()` has grouped output by 'timept'. You can override using the
## `.groups` argument.
```

```
#Visualize
ggplot(growth_RFU_data,  aes(x = timept, y = value, group = Trt, color = Trt, ymin = value-sd, ymax = value+sd))  + 
    geom_errorbar(width = 0.5) +
    geom_line(size = 1) +
    theme(axis.text.x = element_text(vjust=0, face = "bold"),
          axis.ticks.x=element_blank()) +
    coord_trans(y = "log") +
    xlab("Time in minutes") + 
    ylab("Average growth (log)") +
    theme_classic() + 
    scale_color_manual(values=c("#112943", "#51b0fa")) +
    theme(plot.title = element_text(hjust = 0.5, face = "bold"))
```

```
#resave(growth_cells, growth_RFU, file = "/My Drive/ShortTermStress-IlluminaData/FullExp/Cryp-DE/data/import_rmd_vars.RData")
```

### 2. Bioinformatic analysis of RNA-seq reads

445,128,534 100bp paired-end reads recovered for C. cryptica

#### 2.1 Quality control, adapter removal & trimming

1. We ran FastQC on
   all fastq files:

```
# run FastQC v0.11.5
fastqc *.fastq
```

2. Adapter removal and trimming of sequences was done using Ktrim:

```
for i in $(cat original_names.txt); do ktrim \
  -1 $i_L004_R1_001.fastq \
  -2 $i_L004_R2_001.fastq \
  -o $i\_ -t 15 -p 33 -q 20 -s 36\
  -a TACACTCTTTCCCTACACGACGCTCTTCCGATCT \
  -a GTGACTGGAGTTCAGACGTGTGCTCTTCCGATCT \
  -a TACACTCTTTCCCTACACGACGCTCTTCCGATCT \
  -a AGATCGGAAGAGCGTCGTSGTAGGGAAAGAGTGTA \
  -a GTGACTGGAGTTCAGACGTGTGCTCTTCCGATCT \
  -a AGATCGGAAGAGCACACGTCTGAACTCCAGTCAC \
  -m 0.5; done
```

The output files of the Ktrim run were checked again using
FastQC:

```
# run Fastqc v0.11.5
fastqc *_L004_R1_001.fastq
fastqc *_L004_R2_001.fastq
```

##2.2 STAR/2.7.3a genome mapping

```
STAR --runThreadN 8 --runMode genomeGenerate \
  --genomeDir original_names/CCMP332_STARIndex \
  --genomeFastaFiles CCMP332_data/CCMP332_renamed.fasta \
  --sjdbGTFfile CCMP332_data/cryptica_pilon5.all.functional_ipr.functional_blast.renamed.gff \
  --sjdbGTFfeatureExon exon \
  --sjdbGTFtagExonParentTranscript Parent \
  --sjdbOverhang 99 --genomeSAindexNbases 12
#the --genomeSAindexNbases = 12 parameter was suggested after running STAR without this parameter. It is based on: min(1, log2(152305664)/2 - 1) with 152305664 being the length of the genome in bases
#--sjdbOverhang =  max read length - 1 = 100 - 1 = 99
```

We ran STAR

```
  for i in $(cat original_names.txt); do STAR  \
  --runThreadN 15 --genomeDir CCMP332_STARIndex \
  --outSAMtype BAM SortedByCoordinate --outReadsUnmapped Fastx \
  --alignIntronMin 1 --alignIntronMax 22618 \
  --readFilesIn trimmed_CCMP332_seqs/$i\.read1.fq\
   trimmed_CCMP332_seqs/$i\.read2.fq \
  --outFileNamePrefix CCMP332_STAR_output/$i\_; done
```

##2.3 Read quantification with HTSeq on gene-level

We ran HTSeq as
follows on a gff file without the fasta included at the end:

```
for i in $(cat original_names.txt);do /share/apps/python/anaconda3-python-3.7.3/lib/python3.7/site-packages/HTSeq-0.11.3-py3.7-linux-x86_64.egg-info/scripts/htseq-count \
--format=bam \
--order=pos \
--stranded=reverse \
--minaqual=10 \
--type=gene \
--idattr=ID \
--mode=union \
--nonunique=none \
--samout=$i\_HTSeq.gene-lvl.out \
CCMP332_STAR_output/$i\_Aligned.sortedByCoord.out.bam \
CCMP332_noFASTA.gff \
>> $i\_HTSeq.gene-lvl.STDOUT;done
```

### 3. Statistical analysis of RNA-seq data in R

#### 3.1 Model fitting in EdgeR

###3.1.1 Data import

Import the count data for Downey et al. 2022 and Nakov et al. 2020
(the long-term hypo-osmotic growth study for *C. cryptica*). The
Nakov et al. 2020 required re-analysis because the initial mapping used
a de-novo transcriptome, rather than the annotated genome from Wade et
al. 2021. We utilized only the CCMP332 data from the long-term
study.

```
#Data Import 
x_cnts <- read.table("/My Drive/ShortTermStress-IlluminaData/FullExp/Cryp-DE/CCMP332_HTSeq_output.txt", header = TRUE, row.names = 1)

nakov_cnts <- read.table("/My Drive/ShortTermStress-IlluminaData/FullExp/Cryp-DE/nakov_data/nakov_HTSeq_output.txt", header = FALSE, row.names = 1)

colnames(x_cnts) <- c("Cntl_R1","0m_R1","15m_R1","30m_R1","60m_R1","2hr_R1","4hr_R1","8hr_R1","10hr_R1",
                      "Cntl_R2","0m_R2","15m_R2","30m_R2","60m_R2","2hr_R2","4hr_R2","8hr_R2","10hr_R2",
                      "Cntl_R3","0m_R3","15m_R3","30m_R3","60m_R3","2hr_R3","4hr_R3","8hr_R3","10hr_R3")

colnames(nakov_cnts) <- c("R1_0","R1_24","R2_0","R2_24","R3_0","R3_24")

x_cnts = x_cnts[-c(21251:21254), ] #drop the last lines that do not contain information pertinent to gene counts

total_gene_number = nrow(x_cnts) #21250

#resave(x_cnts,nakov_cnts, file = "/My Drive/ShortTermStress-IlluminaData/FullExp/Cryp-DE/data/import_rmd_vars.RData")
```

EdgeR works with a DGEList data class object. This needed to be
created using the count data and a group object that contains
information on the different groups. We also developed multidimensional
scale plot including the mock control ASW 24 treatment to look at data
similarity.

```
#----All replicates without Control----#
# We removed the  because initial comparisons showed no significant difference
group_cntl <- c("Cntl_R1","0m","15m","30m","60m","2hr","4hr","8hr","10hr",
           "Cntl_R2","0m","15m","30m","60m","2hr","4hr","8hr","10hr",
           "Cntl_R3","0m","15m","30m","60m","2hr","4hr","8hr","10hr")

y_list_cntl <- DGEList(counts=as.matrix(x_cnts),group=group_cntl)
```

##### 3.1.2 Data filtering

Here, the genes that have very low counts across all the libraries
were removed. Filtering was done using the CPM (count per million).
Here, we retained all the genes that have least one CPM in at least
three samples:

```
keep <- rowSums(cpm(y_list_cntl)>1)>=3 #keep genes that have a least one count per million in at least three samples
y_list_cntl <- y_list_cntl[keep,]
y_list_cntl[,]
```

```
## An object of class "DGEList"
## $counts
##              Cntl_R1 0m_R1 15m_R1 30m_R1 60m_R1 2hr_R1 4hr_R1 8hr_R1 10hr_R1
## CCRYP_000001      84   146    116    358    137    137    143    122     196
## CCRYP_000002      47    44     78    305     54    144     56     69     134
## CCRYP_000003      16    45     59     98     18     52     47     35      64
## CCRYP_000005      11    10     15     35     17     11      6      9      16
## CCRYP_000006      55    86     47    143     31     52     62     55     103
##              Cntl_R2 0m_R2 15m_R2 30m_R2 60m_R2 2hr_R2 4hr_R2 8hr_R2 10hr_R2
## CCRYP_000001      77   157    107    506    164    130    169    137     114
## CCRYP_000002      40    55     79    473     81    113     60     92      68
## CCRYP_000003       9    56     54    130     45     42     52     42      55
## CCRYP_000005       4    10     14     49     17     10      5      6      18
## CCRYP_000006      48    97     43    182     58     52     68     52      68
##              Cntl_R3 0m_R3 15m_R3 30m_R3 60m_R3 2hr_R3 4hr_R3 8hr_R3 10hr_R3
## CCRYP_000001      94   149     95    209    155    123    116    106     263
## CCRYP_000002      58    47     85    136     72     82     51     45     116
## CCRYP_000003      28    48     48     66     36     32     41     28      92
## CCRYP_000005      23    10      9     20     16      8      6     11      29
## CCRYP_000006      66    89     38    103     49     51     55     58     109
## 12951 more rows ...
## 
## $samples
##           group lib.size norm.factors
## Cntl_R1 Cntl_R1  3537770            1
## 0m_R1        0m  3084445            1
## 15m_R1      15m  3460086            1
## 30m_R1      30m  5209309            1
## 60m_R1      60m  2599734            1
## 22 more rows ...
```

```
y_list_cntl$samples$lib.size <- colSums(y_list_cntl$counts)
num_kept_gene = nrow(y_list_cntl$counts)
```

##### 3.1.3 Data normalization

Next, we calculated a set of normalization factors (one for each
sample) to eliminate composition biases between libraries:

```
y_list_cntl <- calcNormFactors(y_list_cntl, method = 'TMM') #normalizes for RNA composition (highly expressed genes)
head(y_list_cntl$samples)
```

```
##           group lib.size norm.factors
## Cntl_R1 Cntl_R1  3537451    0.9430666
## 0m_R1        0m  3084098    0.9597864
## 15m_R1      15m  3459566    0.9667379
## 30m_R1      30m  5208139    1.1556713
## 60m_R1      60m  2599410    0.9086212
## 2hr_R1      2hr  3163770    0.9478741
```

```
plotMDS(y_list_cntl, top = 500)
```

Based on similarity of the mock control counts to that of the 0
minute control, we removed the “Cntl” samples and repeated the above
steps. Further analyses used this subset.

```
#MDS plot without mock control data
gene_cnts <-  subset(x_cnts, select=-c(Cntl_R1,Cntl_R2,Cntl_R3))
group <- c("0m","15m","30m","60m","2hr","4hr","8hr","10hr",
          "0m","15m","30m","60m","2hr","4hr","8hr","10hr",
          "0m","15m","30m","60m","2hr","4hr","8hr","10hr")

y_list <- DGEList(counts=as.matrix(gene_cnts),group=group)
keep <- rowSums(cpm(y_list)>1)>=3 #keep genes that have a least one count per million in at least three samples
y_list <- y_list[keep,]
y_toss <- y_list[,]
y_list$samples$lib.size <- colSums(y_list$counts)
num_kept_gene = nrow(y_list$counts)
y_list <- calcNormFactors(y_list, method = 'TMM') #normalizes for RNA composition (highly expressed genes)
head(y_list$samples)
```

```
##        group lib.size norm.factors
## 0m_R1     0m  3084082    0.9542249
## 15m_R1   15m  3459556    0.9612427
## 30m_R1   30m  5208090    1.1490092
## 60m_R1   60m  2599402    0.9040386
## 2hr_R1   2hr  3163759    0.9424528
## 4hr_R1   4hr  3123184    0.9700367
```

```
plotMDS(y_list, top = 500)
```

```
#resave(gene_cnts,y_list, file = "/My Drive/ShortTermStress-IlluminaData/FullExp/Cryp-DE/data/import_rmd_vars.RData")
```

We repeated the process with the Nakov et al. 2020 data

```
#Setting it all up with the C. cryptica long term data
nakov_332 <- subset(nakov_cnts, select=c(R2_0,R2_24))
group_nakov <- c("0", "24","0", "24","0", "24")
group_332 <- c("0", "24")
y_list_nakov <- DGEList(counts=as.matrix(nakov_332),group=group_332)
keep_nakov <- rowSums(cpm(y_list_nakov)>1)>=2 #keep genes that have a least one count per million in at least three samples
y_list_nakov <- y_list_nakov[keep_nakov,]
dim(y_list_nakov)
```

```
## [1] 12366     2
```

```
y_list_nakov$samples$lib.size <- colSums(y_list_nakov$counts)
num_kept_gene = nrow(y_list_nakov$counts)
y_list_nakov <- calcNormFactors(y_list_nakov, method = 'TMM') #normalizes for RNA composition (highly expressed genes)
head(y_list_nakov$samples)
```

```
##       group lib.size norm.factors
## R2_0      0 16433537    1.0673781
## R2_24    24 17919534    0.9368751
```

```
#resave(nakov_332,y_list_nakov, file = "/My Drive/ShortTermStress-IlluminaData/FullExp/Cryp-DE/data/import_rmd_vars.RData")
```

We calculated Poisson distances for both sets of data and visualized
with pheatmap

```
poisd <- PoissonDistance(t(y_list$counts))
poisd_nakov <- PoissonDistance(t(y_list_nakov$counts))

#define colors to be used in the plot
colors_heatmap <- colorRampPalette(rev(brewer.pal(9,"Blues")))(255)

#plot a heatmap
samplePoisDistMatrix <- as.matrix(poisd$dd)
rownames(samplePoisDistMatrix) <- paste(y_list$samples$group)
colnames(samplePoisDistMatrix) <- paste(y_list$samples$group)
pheatmap(samplePoisDistMatrix,
         clustering_distance_rows=poisd$dd,
         clustering_distance_cols=poisd$dd,
         col=colors_heatmap,cex=1)#calculate poisson distances for the normalized count data
```

```
#C. cryptica longterm data
samplePoisDistMatrix <- as.matrix(poisd_nakov$dd)
rownames(samplePoisDistMatrix) <- paste(y_list_nakov$samples$group)
colnames(samplePoisDistMatrix) <- paste(y_list_nakov$samples$group)
pheatmap(samplePoisDistMatrix,
         clustering_distance_rows=poisd_nakov$dd,
         clustering_distance_cols=poisd_nakov$dd,
         col=colors_heatmap,cex=1)#calculate poisson distances for the normalized count data
```

##### 3.1.4 Design matrix and dispersion estimation

Prior to differential expression analysis, dispersion needs to be
estimated from a data matrix. edgeR uses the negative binomial (NB)
distribution to model the read counts for each gene in each sample. The
dispersion parameter of the NB distribution accounts for variability
between biological replicates. edgeR estimates an empirical Bayes
moderated dispersion for each individual gene. It also estimates a
common dispersion, which is a global dispersion estimate averaged over
all genes, and a trended dispersion where the dispersion of a gene is
predicted from its abundance.

```
#Created a design matrix for future data contrasts

design <- model.matrix(~0+group, data = y_list$samples)
colnames(design) <- levels(y_list$samples$group)

#Dispersion estimation & model fitting
y_list <- estimateDisp(y_list, design)
plotBCV(y_list)
```

```
#Design matrix for long-term data

design_nakov <- model.matrix(~0+group_nakov, data = y_list_nakov$samples)
colnames(design_nakov) <- levels(y_list_nakov$samples$group)

#resave(design, design_nakov, file = "/My Drive/ShortTermStress-IlluminaData/FullExp/Cryp-DE/data/import_rmd_vars.RData")
```

##### 3.1.5 Model fitting

glmQLFit is a quasi-likelihood (QL) method that accounts for
gene-specific variability from both biological and technical sources.
Under the QL framework, the NB dispersion trend is used to describe the
overall biological variability across all genes, and gene-specific
variability above and below the overall level is picked up by the QL
dispersion. In the QL approach, the individual (tagwise) NB dispersions
are not used.

```
fit_group_model <- glmQLFit(y_list, design, robust = TRUE)
head(fit_group_model$coefficients)
```

```
##                      0m      10hr       15m       2hr        30m        4hr
## CCRYP_000001  -9.920891 -10.19841 -10.24761 -10.00913  -9.727104  -9.954751
## CCRYP_000002 -11.050294 -10.76839 -10.51693 -10.15107  -9.911733 -10.892128
## CCRYP_000003 -11.030161 -11.18574 -10.92747 -11.13845 -11.007729 -11.068738
## CCRYP_000005 -12.623782 -12.39362 -12.36687 -12.59870 -12.053307 -13.160445
## CCRYP_000006 -10.428704 -10.89697 -11.15657 -10.93039 -10.625916 -10.790332
## CCRYP_000007 -10.971651 -11.10870 -11.43505 -11.37961 -11.286043 -11.386840
##                     60m       8hr
## CCRYP_000001  -9.777955 -10.30644
## CCRYP_000002 -10.569688 -10.88039
## CCRYP_000003 -11.309167 -11.55029
## CCRYP_000005 -11.983033 -12.93246
## CCRYP_000006 -10.975925 -11.09640
## CCRYP_000007 -11.915511 -11.42867
```

```
plotQLDisp(fit_group_model)
```

```
#C. cryptica longterm data
DE_nakov <- exactTest(y_list_nakov,dispersion = 0.16) #dispersion assigned based on exactTest author recommendation
#For future comparison, we used a subset of the long-term data. Due to the last replication for the CCMP332 expression data, we imposed a strenuous LFC filter. 
nakov_DE_dr <- subset(DE_nakov$table, (`logFC`) <= -1) #downregulated genes
nakov_DE_ur <- subset(DE_nakov$table, (`logFC`) >= 1) #upregulated genes

nakov_DE <- rbind(nakov_DE_dr,nakov_DE_ur)

#resave(nakov_DE, fit_group_model, file = "/My Drive/ShortTermStress-IlluminaData/FullExp/Cryp-DE/data/import_rmd_vars.RData")
```

#### 3.2 Testing for differential expression using stage-wise analysis:

##### 3.2.1 Defining contrasts to test

We contrasted all timepoints against the 0 minute timepoint

```
Contrasts=matrix(0,nrow = ncol(fit_group_model$coefficients),ncol=7)
rownames(Contrasts)=colnames(fit_group_model$coefficients)
colnames(Contrasts)=c("C0m-C15m", "C0m-C30m", "C0m-C60m", "C0m-C2hr", "C0m-C4hr", "C0m-C8hr", "C0m-C10hr")

Contrasts[c("0m", "15m"), "C0m-C15m"]=c(1,-1)
Contrasts[c("0m", "30m"), "C0m-C30m"]=c(1,-1)
Contrasts[c("0m", "60m"), "C0m-C60m"]=c(1,-1)
Contrasts[c("0m", "2hr"), "C0m-C2hr"]=c(1,-1)
Contrasts[c("0m", "4hr"), "C0m-C4hr"]=c(1,-1)
Contrasts[c("0m", "8hr"), "C0m-C8hr"]=c(1,-1)
Contrasts[c("0m", "10hr"), "C0m-C10hr"]=c(1,-1)

#resave(Contrasts, file = "/My Drive/ShortTermStress-IlluminaData/FullExp/Cryp-DE/data/import_rmd_vars.RData")
```

##### 3.2.2 Stage-wise testing

We performed the stage-wise testing procedure in stageR.
StageR allows for simultaneous FDR control in all the contrasts, and
consists of two steps: the screening stage, and the confirmation
stage.

The screening stage tested whether any of the contrasts were
significant. The screening stage gives P-values as output, but these are
not yet FDR-controlled so should not be used in downstream analyses.

```
#screening stage
alpha=0.01 
screenTest <- glmQLFTest(fit_group_model, contrast=Contrasts)
pScreen <- screenTest$table$PValue
names(pScreen) <- rownames(screenTest$table)
```

The screening stage was followed by the confirmation stage. In the
confirmation stage, every contrast was assessed separately. The
confirmation stage P-values were adjusted to control the FWER across the
hypotheses within a gene and are subsequently corrected to the
BH-adjusted significance level of the screening stage. This allowed for
a direct comparison of the adjusted P-values to the provided
significance level alpha for both screening and confirmation stage
adjusted P-values. Here, we used the holm method for correction of the
P-values.

```
#confirmation stage
confirmationResults <- sapply(1:ncol(Contrasts),function(i) glmQLFTest(fit_group_model, 
                              contrast = Contrasts[,i]), simplify=FALSE) #calculates F-test for each contrast
confirmationPList <- lapply(confirmationResults, 
                            function(x) x$table$PValue) #takes the p-values from all genes for each contrast and puts them in a list
confirmationP <- as.matrix(Reduce(f=cbind,confirmationPList)) 
rownames(confirmationP) <- rownames(confirmationResults[[1]]$table)
colnames(confirmationP) <- colnames(Contrasts)
stageRObj <- stageR(pScreen=pScreen, pConfirmation=confirmationP) #constructs an object
stageRAdj <- stageWiseAdjustment(object=stageRObj, 
                                 method="holm", alpha=0.01) #adjusts the P-values using FWER correction using the holm method
res <- getResults(stageRAdj)
```

```
## The returned adjusted p-values are based on a stage-wise testing approach and are only valid for the provided target OFDR level of 1%. If a different target OFDR level is of interest,the entire adjustment should be re-run.
```

```
#Number of differentially expressed genes in every contrast
SignifGenes <- colSums(res)

#Establish adjusted p-values
adjusted_p <- getAdjustedPValues(stageRAdj, onlySignificantGenes = FALSE, order = FALSE)
```

```
## The returned adjusted p-values are based on a stage-wise testing approach and are only valid for the provided target OFDR level of 1%. If a different target OFDR level is of interest,the entire adjustment should be re-run.
```

```
padj_df <- as.data.frame(adjusted_p) #establish as a dataframe for use downstream
```

Upon finishing the stage-wise testing procedure, we checked the
number of significant genes:

```
#visualize the number of significant genes
res_df = as.data.frame(res) #res is a matrix, to be able to subset, change it to a dataframe
res_df2 = res_df
res_df2$gene = rownames(res_df2)
OnlySignGenes = res_df2[res_df2$padjScreen == 1,] #removes rows for which global test was non significant
dim(OnlySignGenes) #10569 significant genes based on stringent alpha = 0.01
```

```
## [1] 10569     9
```

```
genesSI <- rownames(adjusted_p)[adjusted_p[,"padjScreen"]<=0.01] 
length(genesSI) #10569 unique genes
```

```
## [1] 10569
```

```
genesNotFoundStageII <- genesSI[genesSI %in% rownames(res)[rowSums(res==0)==8]]
length(genesNotFoundStageII) #There are 0 genes found to be significant in screening, but not confirmation
```

```
## [1] 0
```

```
#resave(adjusted_p, OnlySignGenes, genesNotFoundStageII, file = "/My Drive/ShortTermStress-IlluminaData/FullExp/Cryp-DE/data/import_rmd_vars.RData")
```

We only included genes that were significant after the confirmation
stage

```
OnlySignGenes_ConStage <- OnlySignGenes [!rownames(OnlySignGenes) %in% genesNotFoundStageII, ]
nrow(OnlySignGenes_ConStage) #10569 rows
```

```
## [1] 10569
```

##### 3.2.3 Summarize the results for downstream analyses

```
#get information on total number of expressed genes = 12939
expressed_genes = nrow(fit_group_model$fitted.values)
expressed_genes
```

```
## [1] 12939
```

```
#relative number of expressed genes in the genome of C. cryptica = 60.89 %
rel_exp_genes = expressed_genes / total_gene_number
rel_exp_genes
```

```
## [1] 0.6088941
```

```
#get information on totaly number of DE expressed genes = 10569
DE_expressed_genes = nrow(OnlySignGenes_ConStage)
DE_expressed_genes
```

```
## [1] 10569
```

```
#relative number of DE expressed genes in the genome of C. cryptica = 81.68% %
rel_DEexp_genes = DE_expressed_genes  / expressed_genes 
rel_DEexp_genes
```

```
## [1] 0.8168328
```

Before we continued with the downstream analyses, we created a single
data object that contains some key-information of the statistical
pipeline outlined above. This included information on logFC, logCPM and
P-values for each gene for each contrast.

First, we selected the FDR adjusted P-values for each contrast using
the output of the stageR screening stage:

```
colnames(adjusted_p)[1] = "padjScreen"
colnames(adjusted_p)[2] = "C0mvsC15m_Padj"
colnames(adjusted_p)[3] = "C0mvsC30m_Padj"
colnames(adjusted_p)[4] = "C0mvsC60m_Padj"
colnames(adjusted_p)[5] = "C0mvsC2hr_Padj"
colnames(adjusted_p)[6] = "C0mvsC4hr_Padj"
colnames(adjusted_p)[7] = "C0mvsC8hr_Padj"
colnames(adjusted_p)[8] = "C0mvsC10hr_Padj"

#resave(OnlySignGenes_ConStage, adjusted_p_onlysign, file = "/My Drive/ShortTermStress-IlluminaData/FullExp/Cryp-DE/data/import_rmd_vars.RData")
```

Second, we extracted the information on logFC, logCPM, F value and
non-adjusted P-values from the *confirmationResults* object:

```
#create empty list to hold the data values
datalist = list()

#loop over the confirmationResults object to obtain the relevant information (table)
for (contrast in c(1,2,3,4,5,6,7)){
  table = confirmationResults[[contrast]]$table
  datalist[[contrast]] <- table
}

#turn list into data frame
confirmationResults_total_dataset = cbind(datalist[[1]],datalist[[2]],datalist[[3]],datalist[[4]]
                                               ,datalist[[5]],datalist[[6]],datalist[[7]])

#rename column names for tracability
#logCPM tells us about the overall presence of that gene in the entire dataset, which is why it's identical across the board. Has nothing to do with contrasts
colnames(confirmationResults_total_dataset)=c("C0mvsC15m_logFC","C0mvsC15m_logCPM","C0mvsC15m_F","C0mvsC15m_nonadj_PValue","C0mvsC30m_logFC","C0mvsC30m_logCPM","C0mvsC30m_F","C0mvsC30m_nonadj_PValue","C0mvsC60m_logFC","C0mvsC60m_logCPM","C0mvsC60m_F","C0mvsC60m_nonadj_PValue","C0mvsC2hr_logFC","C0mvsC2hr_logCPM","C0mvsC2hr_F","C0mvsC2hr_nonadj_PValue","C0mvsC4hr_logFC","C0mvsC4hr_logCPM","C0mvsC4hr_F","C0mvsC4hr_nonadj_PValue","C0mvsC8hr_logFC","C0mvsC8hr_logCPM","C0mvsC8hr_F","C0mvsC8hr_nonadj_PValue","C0mvsC10hr_logFC","C0mvsC10hr_logCPM","C0mvsC10hr_F","C0mvsC10hr_nonadj_PValue")
```

Then we combined the FDR-adjusted P-values and the table with
information on logFC etc. into a single data frame:

```
#merge the data frames
table = merge(confirmationResults_total_dataset,adjusted_p, by = 0, all = TRUE)
all_results <- table[,-1]

#use the first column (gene names) for the row names
rownames(all_results) <- table[,1]
all_results <- (na.omit(all_results))
head(all_results)
```

```
##              C0mvsC15m_logFC C0mvsC15m_logCPM C0mvsC15m_F
## CCRYP_000001       0.4713537         5.525873  24.2077232
## CCRYP_000002      -0.7694779         4.818694  10.2566718
## CCRYP_000003      -0.1481453         3.917981   0.7648136
## CCRYP_000005      -0.3706496         2.191358   2.1083980
## CCRYP_000006       1.0500842         4.348562  46.3279224
## CCRYP_000008       0.9899192         4.336509  23.7372691
##              C0mvsC15m_nonadj_PValue C0mvsC30m_logFC C0mvsC30m_logCPM
## CCRYP_000001            4.343084e-05     -0.27957525         5.525873
## CCRYP_000002            3.630795e-03     -1.64259737         4.818694
## CCRYP_000003            3.899782e-01     -0.03236368         3.917981
## CCRYP_000005            2.839490e-01     -0.82302187         2.191358
## CCRYP_000006            3.498690e-07      0.28451796         4.348562
## CCRYP_000008            4.924978e-05      1.14738413         4.336509
##              C0mvsC30m_F C0mvsC30m_nonadj_PValue C0mvsC60m_logFC
## CCRYP_000001 11.25524838            2.487663e-03      -0.2062128
## CCRYP_000002 64.23304182            1.949782e-08      -0.6933680
## CCRYP_000003  0.04260787            8.923105e-01       0.4025204
## CCRYP_000005 14.66994706            3.445773e-03      -0.9244058
## CCRYP_000006  4.94729302            3.519344e-02       0.7894733
## CCRYP_000008 38.78046335            1.486369e-06       1.7745411
##              C0mvsC60m_logCPM C0mvsC60m_F C0mvsC60m_nonadj_PValue
## CCRYP_000001         5.525873    5.161386            3.195153e-02
## CCRYP_000002         4.818694    7.885306            9.419149e-03
## CCRYP_000003         3.917981    4.710796            3.946765e-02
## CCRYP_000005         2.191358   14.793011            4.469992e-03
## CCRYP_000006         4.348562   26.840327            2.198055e-05
## CCRYP_000008         4.336509   58.380747            4.669842e-08
##              C0mvsC2hr_logFC C0mvsC2hr_logCPM C0mvsC2hr_F
## CCRYP_000001      0.12729919         5.525873  1.88218405
## CCRYP_000002     -1.29730374         4.818694 32.72827726
## CCRYP_000003      0.15622440         3.917981  0.77976939
## CCRYP_000005     -0.03618298         2.191358  0.01784375
## CCRYP_000006      0.72378363         4.348562 23.65986049
## CCRYP_000008      0.80109334         4.336509 16.11712025
##              C0mvsC2hr_nonadj_PValue C0mvsC4hr_logFC C0mvsC4hr_logCPM
## CCRYP_000001            2.034533e-01      0.04885057         5.525873
## CCRYP_000002            5.417049e-06     -0.22818579         4.818694
## CCRYP_000003            3.854502e-01      0.05565480         3.917981
## CCRYP_000005            9.210282e-01      0.77424029         2.191358
## CCRYP_000006            5.028548e-05      0.52172010         4.348562
## CCRYP_000008            4.622899e-04      0.30036179         4.336509
##              C0mvsC4hr_F C0mvsC4hr_nonadj_PValue C0mvsC8hr_logFC
## CCRYP_000001   0.2846715             0.644439092       0.5562258
## CCRYP_000002   0.7865562             0.383420949      -0.2451260
## CCRYP_000003   0.1028127             0.751089572       0.7503890
## CCRYP_000005   6.4236569             0.067416328       0.4453227
## CCRYP_000006  13.1179758             0.001268388       0.9632763
## CCRYP_000008   2.6019817             0.119022755       0.5652576
##              C0mvsC8hr_logCPM C0mvsC8hr_F C0mvsC8hr_nonadj_PValue
## CCRYP_000001         5.525873  35.2640350            3.100156e-06
## CCRYP_000002         4.818694   0.9784567            3.318617e-01
## CCRYP_000003         3.917981  16.8701397            3.634482e-04
## CCRYP_000005         2.191358   2.5839890            2.406365e-01
## CCRYP_000006         4.348562  43.1183293            6.343426e-07
## CCRYP_000008         4.336509   9.2474787            5.392498e-03
##              C0mvsC10hr_logFC C0mvsC10hr_logCPM C0mvsC10hr_F
## CCRYP_000001        0.4003748          5.525873    20.387317
## CCRYP_000002       -0.4067069          4.818694     3.048789
## CCRYP_000003        0.2244549          3.917981     1.885134
## CCRYP_000005       -0.3320500          2.191358     2.024402
## CCRYP_000006        0.6755596          4.348562    24.995884
## CCRYP_000008        0.2595381          4.336509     2.262058
##              C0mvsC10hr_nonadj_PValue   padjScreen C0mvsC15m_Padj
## CCRYP_000001             1.253717e-04 5.435395e-09   3.190188e-04
## CCRYP_000002             9.282000e-02 1.123595e-07   2.222484e-02
## CCRYP_000003             1.816904e-01 1.459588e-03   1.000000e+00
## CCRYP_000005             2.938133e-01 5.731703e-06   1.000000e+00
## CCRYP_000006             3.528087e-05 4.950012e-06   2.569943e-06
## CCRYP_000008             1.448474e-01 1.120762e-06   3.014679e-04
##              C0mvsC30m_Padj C0mvsC60m_Padj C0mvsC2hr_Padj C0mvsC4hr_Padj
## CCRYP_000001   1.218200e-02   1.173491e-01   4.981516e-01    0.788948568
## CCRYP_000002   1.432202e-07   4.612522e-02   3.979063e-05    0.812557091
## CCRYP_000003   1.000000e+00   2.899074e-01   1.000000e+00    1.000000000
## CCRYP_000005   2.531073e-02   3.283408e-02   1.000000e+00    0.412669065
## CCRYP_000006   4.308524e-02   1.345474e-04   1.846846e-04    0.003105624
## CCRYP_000008   1.091804e-05   3.430206e-07   2.263817e-03    0.291425003
##              C0mvsC8hr_Padj C0mvsC10hr_Padj
## CCRYP_000001   2.277202e-05    0.0007674256
## CCRYP_000002   8.125571e-01    0.3409020500
## CCRYP_000003   2.669688e-03    1.0000000000
## CCRYP_000005   1.000000e+00    1.0000000000
## CCRYP_000006   4.659528e-06    0.0001727691
## CCRYP_000008   1.980514e-02    0.2914250034
```

```
#resave(OnlySignGenes_ConStage, confirmationResults_total_dataset, table, file = "/My Drive/ShortTermStress-IlluminaData/FullExp/Cryp-DE/data/import_rmd_vars.RData")
```

The resulting data frame *all\_results* was used in multiple
analyses downstream to access basic stats on each gene for each
contrast.

##### 3.2.4 Subset to genes DE at 2 timepoints

In order to focus on genes most likely to play a significant role in
the short-term hypo-osmotic stress response, we limited our dataset to
those genes that were significant at two consecutive timepoints in the
same direction (i.e., both up/downregulated).

```
df <- res_df2[,-c(1,9)]
colSums(df)
```

```
##  C0m-C15m  C0m-C30m  C0m-C60m  C0m-C2hr  C0m-C4hr  C0m-C8hr C0m-C10hr 
##      2442      4900      4453      2533      1848      1963      2031
```

```
keep_2_SignDEG <- df[rowSums(df[-1] & df[-ncol(df)]) > 0,]
keep2_colSums <- colSums(keep_2_SignDEG)
DEG2sig_names = rownames(keep_2_SignDEG)

DEG2sig_results <- (na.omit(all_results[DEG2sig_names,]))

sign2LFC <- subset(DEG2sig_results, select=c("C0mvsC15m_logFC","C0mvsC15m_Padj","C0mvsC30m_logFC","C0mvsC30m_Padj",
                                                            "C0mvsC60m_logFC","C0mvsC60m_Padj","C0mvsC2hr_logFC","C0mvsC2hr_Padj",
                                                            "C0mvsC4hr_logFC","C0mvsC4hr_Padj","C0mvsC8hr_logFC","C0mvsC8hr_Padj",
                                                            "C0mvsC10hr_logFC","C0mvsC10hr_Padj"))
colnames(sign2LFC) <- c("0v15m","0v15m_Padj","0v30m","0v30m_Padj","0v1h","0v1h_Padj","0v2h","0v2h_Padj",
                             "0v4h","0v4h_Padj","0v8h","0v8h_Padj","0v10h","0v10h_Padj")
```

Plotted the heatmap of the significant data with a gene expression
data matrix

```
sign2LFC_mat <- as.matrix(sign2LFC %>% dplyr::select(`0v15m`,`0v30m`,`0v1h`,`0v2h`,`0v4h`,`0v8h`,`0v10h`))
min(sign2LFC_mat) #-9.470548
```

```
## [1] -9.470548
```

```
max(sign2LFC_mat) #7.635282
```

```
## [1] 7.635282
```

```
paletteLength <- 500
myColor <- colorRampPalette(c("blue", "snow", "red"))(paletteLength)
sign2LFC_hmp <- heatmap.2(sign2LFC_mat,dendrogram = "row",trace="none",col=myColor, labRow=TRUE, cexCol = 1, Colv = FALSE)
```

```
#resave(all_results, sign2LFC, file = "/My Drive/ShortTermStress-IlluminaData/FullExp/Cryp-DE/data/import_rmd_vars.RData")
```

We also looked at the non-significant genes as a quick
verification

```
#The genes that are not differentially expressed
all_sig_names <- rownames(all_results)
pScreen_df <- as.data.frame(pScreen)
pScreen_df$cluster <- "NotSig"
not_sig_genes <- (na.omit(pScreen_df[!row.names(pScreen_df)%in%all_sig_names,]))
not_sig_genes <- subset(not_sig_genes, select=c("cluster"))

NS_LFC_results <- (na.omit(all_results[!row.names(all_results)%in%DEG2sig_names,]))
NS_LFC_df <- subset(NS_LFC_results, select=c("C0mvsC15m_logFC","C0mvsC15m_Padj","C0mvsC30m_logFC","C0mvsC30m_Padj",
                                                "C0mvsC60m_logFC","C0mvsC60m_Padj","C0mvsC2hr_logFC","C0mvsC2hr_Padj",
                                                "C0mvsC4hr_logFC","C0mvsC4hr_Padj","C0mvsC8hr_logFC","C0mvsC8hr_Padj",
                                                "C0mvsC10hr_logFC","C0mvsC10hr_Padj"))
colnames(NS_LFC_df) <- c("0v15m","0v15m_Padj","0v30m","0v30m_Padj","0v1h","0v1h_Padj","0v2h","0v2h_Padj",
                       "0v4h","0v4h_Padj","0v8h","0v8h_Padj","0v10h","0v10h_Padj")

#Rows of significant and non-significant dataframes add up to 11528 as expected. No genes missing

NS_LFC_mat <- as.matrix(NS_LFC_df %>% dplyr::select(`0v15m`,`0v30m`,`0v1h`,`0v2h`,`0v4h`,`0v8h`,`0v10h`))

min(NS_LFC_mat) #-9.452026
```

```
## [1] -6.316635
```

```
max(NS_LFC_mat) #7.762832
```

```
## [1] 6.970724
```

```
paletteLength <- 500
myColor <- colorRampPalette(c("blue", "snow", "red"))(paletteLength)
NS_LFC_hmp <- heatmap.2(NS_LFC_mat,dendrogram = "row",trace="none",col=myColor, labRow=TRUE, cexCol = 1, Colv = FALSE)
```

##### 3.2.5 Graph of genes up and downregulated at each timepoint

```
sign_0v15m <- rownames(keep_2_SignDEG)[keep_2_SignDEG[,"C0m-C15m"]==1] 
length(sign_0v15m)
```

```
## [1] 1614
```

```
sign2LFC_0v15m <- sign2LFC [rownames(sign2LFC) %in% sign_0v15m, ]
logFCup_0v15m <- subset(sign2LFC_0v15m, (`0v15m`) > 0) 
logFCdown_0v15m <- subset(sign2LFC_0v15m, (`0v15m`) < 0)

sign_0v30m <- rownames(keep_2_SignDEG)[keep_2_SignDEG[,"C0m-C30m"]==1] 
length(sign_0v30m)
```

```
## [1] 3497
```

```
sign2LFC_0v30m <- sign2LFC [rownames(sign2LFC) %in% sign_0v30m, ]
logFCup_0v30m <- subset(sign2LFC_0v30m, (`0v30m`) > 0) 
logFCdown_0v30m <- subset(sign2LFC_0v30m, (`0v30m`) < 0)

sign_0v1h <- rownames(keep_2_SignDEG)[keep_2_SignDEG[,"C0m-C60m"]==1] 
length(sign_0v1h)
```

```
## [1] 3412
```

```
sign2LFC_0v1h <- sign2LFC [rownames(sign2LFC) %in% sign_0v1h, ]
logFCup_0v1h <- subset(sign2LFC_0v1h, (`0v1h`) > 0) 
logFCdown_0v1h <- subset(sign2LFC_0v1h, (`0v1h`) < 0)

sign_0v2h <- rownames(keep_2_SignDEG)[keep_2_SignDEG[,"C0m-C2hr"]==1] 
length(sign_0v2h)
```

```
## [1] 2162
```

```
sign2LFC_0v2h <- sign2LFC [rownames(sign2LFC) %in% sign_0v2h, ]
logFCup_0v2h <- subset(sign2LFC_0v2h, (`0v2h`) > 0) 
logFCdown_0v2h <- subset(sign2LFC_0v2h,  (`0v2h`) < 0)

sign_0v4h <- rownames(keep_2_SignDEG)[keep_2_SignDEG[,"C0m-C4hr"]==1] 
length(sign_0v4h)
```

```
## [1] 1423
```

```
sign2LFC_0v4h <- sign2LFC [rownames(sign2LFC) %in% sign_0v4h, ]
logFCup_0v4h <- subset(sign2LFC_0v4h, (`0v4h`) > 0) 
logFCdown_0v4h <- subset(sign2LFC_0v4h,  (`0v4h`) < 0)

sign_0v8h <- rownames(keep_2_SignDEG)[keep_2_SignDEG[,"C0m-C8hr"]==1] 
length(sign_0v8h)
```

```
## [1] 1325
```

```
sign2LFC_0v8h <- sign2LFC [rownames(sign2LFC) %in% sign_0v8h, ]
logFCup_0v8h <- subset(sign2LFC_0v8h, (`0v8h`) > 0) 
logFCdown_0v8h <- subset(sign2LFC_0v8h, (`0v8h`) < 0)

sign_0v10h <- rownames(keep_2_SignDEG)[keep_2_SignDEG[,"C0m-C10hr"]==1] 
length(sign_0v10h)
```

```
## [1] 1432
```

```
sign2LFC_0v10h <- sign2LFC [rownames(sign2LFC) %in% sign_0v10h, ]
logFCup_0v10h <- subset(sign2LFC_0v10h, (`0v10h`) > 0) 
logFCdown_0v10h <- subset(sign2LFC_0v10h, (`0v10h`) < 0)

up <- c(584, 1555, 1364, 664, 540, 763, 659)
down <- c(-1029, -1942, -2048, -1498, -883, -561, -772)
timept <- c("t15m","t30m","t1h","t2h","t4h","t8h","t10h","t15m","t30m","t1h","t2h","t4h","t8h","t10h")
level_order = c("t15m","t30m","t1h","t2h","t4h","t8h","t10h")
dat <- data.frame(
  group = rep(c("upregulated", "downregulated"), each=7),
  x = timept,
  y = c(up, down)
)

dat$x <- as.character(dat$x)
dat$x <- factor(dat$x, levels=(c("t15m","t30m","t1h","t2h","t4h","t8h","t10h")))
ggplot(dat, aes(x=x, y=y, fill=group)) + 
  geom_bar(stat="identity", position="identity", width=0.5) +
  scale_fill_manual(values=c("#28db69","#a9acff")) +
  scale_x_discrete(timept) +
  scale_y_continuous(breaks=seq(-2500,2500,500), limits=c(-2200,2200)) +
  theme_classic() +
  theme(axis.text.x=element_text(size=14), axis.text.y=element_text(size=14), legend.text=element_text(size=14)) +
  ylab("Total number of DE genes") +
  xlab("Time exposed to 0 ppt ASW")
```

##### 3.2.6 Unique and shared DE between timepoints

We created variables for the shared and unique sets of DE genes

```
#calculate number of DE genes for each contrast
allDE_C0mvsC15m = length(c(rownames (subset (OnlySignGenes_ConStage, OnlySignGenes_ConStage$`C0m-C15m`== 1)))) #2442
allDE_C0mvsC30m = length(c(rownames (subset (OnlySignGenes_ConStage, OnlySignGenes_ConStage$`C0m-C30m`== 1)))) #4900
allDE_C0mvsC60m = length(c(rownames (subset (OnlySignGenes_ConStage, OnlySignGenes_ConStage$`C0m-C60m`== 1)))) #4453
allDE_C0mvsC2hr = length(c(rownames (subset (OnlySignGenes_ConStage, OnlySignGenes_ConStage$`C0m-C2hr`== 1)))) #2533
allDE_C0mvsC4hr = length(c(rownames (subset (OnlySignGenes_ConStage, OnlySignGenes_ConStage$`C0m-C4hr`== 1)))) #1848
allDE_C0mvsC8hr = length(c(rownames (subset (OnlySignGenes_ConStage, OnlySignGenes_ConStage$`C0m-C8hr`== 1)))) #1963
allDE_C0mvsC10hr = length(c(rownames (subset (OnlySignGenes_ConStage, OnlySignGenes_ConStage$`C0m-C10hr`== 1)))) #2031

#calculate number of DE genes unique within in a contrast within a genotype/average response
UniqueSubDE_C0mvsC15m = length(c(rownames (subset (OnlySignGenes_ConStage, OnlySignGenes_ConStage$`C0m-C15m` == 1 
                                                   & OnlySignGenes_ConStage$`C0m-C30m` == 0 & OnlySignGenes_ConStage$`C0m-C60m` == 0 
                                                   & OnlySignGenes_ConStage$`C0m-C2hr` == 0 & OnlySignGenes_ConStage$`C0m-C4hr` == 0 
                                                   & OnlySignGenes_ConStage$`C0m-C8hr` == 0 & OnlySignGenes_ConStage$`C0m-C10hr` == 0)))) #334
UniqueSubDE_C0mvsC30m = length(c(rownames (subset (OnlySignGenes_ConStage, OnlySignGenes_ConStage$`C0m-C15m` == 0 
                                                   & OnlySignGenes_ConStage$`C0m-C30m` == 1 & OnlySignGenes_ConStage$`C0m-C60m` == 0 
                                                   & OnlySignGenes_ConStage$`C0m-C2hr` == 0 & OnlySignGenes_ConStage$`C0m-C4hr` == 0 
                                                   & OnlySignGenes_ConStage$`C0m-C8hr` == 0 & OnlySignGenes_ConStage$`C0m-C10hr` == 0)))) #1014
UniqueSubDE_C0mvsC60m = length(c(rownames (subset (OnlySignGenes_ConStage, OnlySignGenes_ConStage$`C0m-C15m` == 0 
                                                   & OnlySignGenes_ConStage$`C0m-C30m` == 0 & OnlySignGenes_ConStage$`C0m-C60m` == 1 
                                                   & OnlySignGenes_ConStage$`C0m-C2hr` == 0 & OnlySignGenes_ConStage$`C0m-C4hr` == 0 
                                                   & OnlySignGenes_ConStage$`C0m-C8hr` == 0 & OnlySignGenes_ConStage$`C0m-C10hr` == 0)))) #740
UniqueSubDE_C0mvsC2hr = length(c(rownames (subset (OnlySignGenes_ConStage, OnlySignGenes_ConStage$`C0m-C15m` == 0 
                                                   & OnlySignGenes_ConStage$`C0m-C30m` == 0 & OnlySignGenes_ConStage$`C0m-C60m` == 0 
                                                   & OnlySignGenes_ConStage$`C0m-C2hr` == 1 & OnlySignGenes_ConStage$`C0m-C4hr` == 0 
                                                   & OnlySignGenes_ConStage$`C0m-C8hr` == 0 & OnlySignGenes_ConStage$`C0m-C10hr` == 0)))) #110
UniqueSubDE_C0mvsC4hr = length(c(rownames (subset (OnlySignGenes_ConStage, OnlySignGenes_ConStage$`C0m-C15m` == 0 
                                                   & OnlySignGenes_ConStage$`C0m-C30m` == 0 & OnlySignGenes_ConStage$`C0m-C60m` == 0 
                                                   & OnlySignGenes_ConStage$`C0m-C2hr` == 0 & OnlySignGenes_ConStage$`C0m-C4hr` == 1 
                                                   & OnlySignGenes_ConStage$`C0m-C8hr` == 0 & OnlySignGenes_ConStage$`C0m-C10hr` == 0)))) #169
UniqueSubDE_C0mvsC8hr = length(c(rownames (subset (OnlySignGenes_ConStage, OnlySignGenes_ConStage$`C0m-C15m` == 0 
                                                   & OnlySignGenes_ConStage$`C0m-C30m` == 0 & OnlySignGenes_ConStage$`C0m-C60m` == 0 
                                                   & OnlySignGenes_ConStage$`C0m-C2hr` == 0 & OnlySignGenes_ConStage$`C0m-C4hr` == 0 
                                                   & OnlySignGenes_ConStage$`C0m-C8hr` == 1 & OnlySignGenes_ConStage$`C0m-C10hr` == 0)))) #303
UniqueSubDE_C0mvsC10hr = length(c(rownames (subset (OnlySignGenes_ConStage, OnlySignGenes_ConStage$`C0m-C15m` == 0 
                                                    & OnlySignGenes_ConStage$`C0m-C30m` == 0 & OnlySignGenes_ConStage$`C0m-C60m` == 0 
                                                    & OnlySignGenes_ConStage$`C0m-C2hr` == 0 & OnlySignGenes_ConStage$`C0m-C4hr` == 0 
                                                    & OnlySignGenes_ConStage$`C0m-C8hr` == 0 & OnlySignGenes_ConStage$`C0m-C10hr` == 1)))) #200


DE_info = as.vector(cbind(allDE_C0mvsC15m,UniqueSubDE_C0mvsC15m,
                               allDE_C0mvsC30m,UniqueSubDE_C0mvsC30m,
                               allDE_C0mvsC60m,UniqueSubDE_C0mvsC60m,
                               allDE_C0mvsC2hr,UniqueSubDE_C0mvsC2hr,
                               allDE_C0mvsC4hr,UniqueSubDE_C0mvsC4hr,
                               allDE_C0mvsC8hr,UniqueSubDE_C0mvsC8hr,
                               allDE_C0mvsC10hr,UniqueSubDE_C0mvsC10hr))


rownames = as.vector(cbind("allDE_C0mvsC15m","UniqueSubDE_C0mvsC15m",
                                "allDE_C0mvsC30m","UniqueSubDE_C0mvsC30m",
                                "allDE_C0mvsC60m","UniqueSubDE_C0mvsC60m",
                                "allDE_C0mvsC2hr","UniqueSubDE_C0mvsC2hr",
                                "allDE_C0mvsC4hr","UniqueSubDE_C0mvsC4hr",
                                "allDE_C0mvsC8hr","UniqueSubDE_C0mvsC8hr",
                                "allDE_C0mvs10hr","UniqueSubDE_C0mvsC10hr"))

DE_colors = rep(c("slategray1","steelblue4","gray80","gray55","sandybrown","chocolate4",
                       "slategray1","steelblue4","gray80","gray55","sandybrown","chocolate4",
                       "slategray1","steelblue4"))
table  <- as.data.frame(cbind(rownames,DE_info,DE_colors ))                        
DE_info = table[,-1]
rownames(DE_info) = table[,1]
DE_info$DE_info_rel  <- (as.numeric(as.character(DE_info$DE_info)) / DE_expressed_genes) * 100
```

Then we plotted the total and unique DE genes at each timepoint

```
par(mfrow=c(1,2))

colsums = colSums(subset(keep_2_SignDEG, select=c("C0m-C15m", "C0m-C30m", "C0m-C60m", "C0m-C2hr", "C0m-C4hr", "C0m-C8hr", "C0m-C10hr")))
par(las=2)
par(mar = c(6, 5, 3, 1), xpd = TRUE)
barplot(colsums,col = "black",
        main = "# DEG in contrasts", ylab = "Total number of DE genes", 
        border=NA, cex.axis=0.9, cex.names = 0.9, cex.lab=1, font.lab=2, ylim=c(0,4000)) #plot barplot


par(mar = c(5, 4, 4, 8), xpd = TRUE)
space<-c(0,0,1,0,1,0,1,0,1,0,1,0,1,0) #create spaces between bars

barplot(DE_info$DE_info_rel,col = as.vector(DE_info$DE_colors), space=space, xaxt='n', 
        main = "% DEG over time", ylab = "Percentage of DE genes (%)", xlab = "Exposure time points",
        border=NA, cex.axis=0.9, cex.lab=1, font.lab=1,ylim=c(0,50)) #plot barplot

legend("topleft",inset=c(1,0),
       c(rows), c( "Total DE genes [0-15m]", "Unique DE genes [0-15m]",
                   "Total DE genes [0-30m]", "Unique DE genes [0-30m]", 
                   "Total DE genes [0-60m]", "Unique DE genes [0-60m]", 
                   "Total DE genes [0-2hr]", "Unique DE genes [0-2hr]",
                   "Total DE genes [0-4hr]", "Unique DE genes [0-4hr]",
                   "Total DE genes [0-8hr]", "Unique DE genes [0-8hr]",
                   "Total DE genes [0-10hr]", "Unique DE genes [0-10hr]"),
       fill = c("slategray1","steelblue4","gray80","gray55","sandybrown","chocolate4"), 
       bty = "n", border=NA, cex=0.5) 
mtext('15 min', side=1, line=0.45, at=1, cex = 0.8)
mtext('30 min', side=1, line=0.45, at=4, cex = 0.8)
mtext('1 hour', side=1, line=0.45, at=7, cex = 0.8)
mtext('2 hour', side=1, line=0.45, at=10, cex = 0.8)
mtext('4 hour', side=1, line=0.45, at=13, cex = 0.8)
mtext('8 hour', side=1, line=0.45, at=16, cex = 0.8)
mtext('10 hour', side=1, line=0.45, at=19, cex = 0.8)
```

### 4. Gene enrichment analyses of:

#### 4.1 Full dataset

Gene enrichment analyses for the entire dataset with GO terms from
the GFF file. This does not account for up or down regulation, it’s just
a bulk overview.

```
#GO enrichment analyses: ORA (over-represetntation analysis) [topGO] for all genes
geneID2GO = readMappings(file = "/My Drive/ShortTermStress-IlluminaData/FullExp/Cryp-DE/CCMP332_GFF_GO.txt")
geneUniverse <- names(geneID2GO)
length(geneUniverse) #6309
#Example search
GOtoMatch = c("GO:0033036")
head(names(geneID2GO)[sapply(geneID2GO,function(x) all(GOtoMatch %in% x))])

number_of_sign_genes = nrow(getAdjustedPValues(stageRAdj, onlySignificantGenes = TRUE, order = FALSE)) #11,759
Padjscreen_sorted = DEG2sig_results[with(DEG2sig_results, 
                                    order(DEG2sig_results$padjScreen)),] #rank genes based on padjScreen
genesOfInterest = rownames(subset(Padjscreen_sorted, rownames(Padjscreen_sorted)%in%geneUniverse))
length(genesOfInterest)

#create gene list for input in topGO
geneList = factor(as.integer(geneUniverse %in% genesOfInterest))
names(geneList) = geneUniverse
str(geneList)

##create a topGO object (for biological process GOs)
GOdata_BP = new("topGOdata", ontology="BP", allGenes=geneList, 
                              annot = annFUN.gene2GO, gene2GO = geneID2GO)

##create a topGO object (for molecular function GOs)
GOdata_MF = new("topGOdata", ontology="MF", allGenes=geneList, 
                              annot = annFUN.gene2GO, gene2GO = geneID2GO)

#run Fisher's exact test
resultFisher_BP <- runTest(GOdata_BP, algorithm = "elim", statistic = "fisher")
resultFisher_MF <- runTest(GOdata_MF, algorithm = "elim", statistic = "fisher")

#extract the significant GO terms, trim the topNodes down to whatever you want (i.e., classic < 0.05)
allRes_BP <- GenTable(GOdata_BP, classic = resultFisher_BP, 
                                    orderBy = "weight", ranksOf = "weight", topNodes = 48)
allRes_MF <- GenTable(GOdata_MF, classic = resultFisher_MF, 
                                    orderBy = "weight", ranksOf = "weight", topNodes = 43)
```

#### 4.2 Significantly differentially expressed genes

Gene enrichment analyses of the significantly DE genes without
accounting for up/down regulation.

```
#select the set of significant DE genes with GO terms
genesOfInterest_allSign  = rownames(subset(OnlySignGenes_ConStage, rownames(OnlySignGenes_ConStage)%in%geneUniverse))
length(genesOfInterest_allSign) 

#create gene list for input in topGO
geneList_allDE = factor(as.integer(geneUniverse %in% genesOfInterest_allSign))
names(geneList_allDE) = geneUniverse
str(geneList_allDE)

##create a topGO object (for biological process GOs)
GOdata_BP_allDE = new("topGOdata", ontology="BP", allGenes=geneList_allDE, 
                           annot = annFUN.gene2GO, gene2GO = geneID2GO)
##create a topGO object (for molecular function GOs)
GOdata_MF_allDE = new("topGOdata", ontology="MF", allGenes=geneList_allDE, 
                           annot = annFUN.gene2GO, gene2GO = geneID2GO)

#run Fisher's exact test
resultFisher_BP_DEG_elim <- runTest(GOdata_BP_allDE, algorithm = "elim", statistic = "fisher")
resultFisher_MF_DEG_elim <- runTest(GOdata_MF_allDE, algorithm = "elim", statistic = "fisher")

#extract the significant GO terms, trim the topNodes down to whatever you want (i.e., classic < 0.05)
DEG_BP_elim <- GenTable(GOdata_BP_allDE, classic = resultFisher_BP_DEG_elim, 
                                        orderBy = "weight", ranksOf = "weight", topNodes = 50)
DEG_MF_elim <- GenTable(GOdata_MF_allDE, classic = resultFisher_MF_DEG_elim, 
                                        orderBy = "weight", ranksOf = "weight", topNodes = 50)


#Generate the files for REVIGO input, which only needs a GO.ID and the p-value 

#import results from REVIGO back in if you want visualize them or use them in future analyses. I don't use this set, so I won't import them again.
```

#### 4.3 Long-term data

Gene enrichment analyses for the long-term Nakov et al. 2020 subset,
separated by regulation direction

```
#C.cryptica longterm data, by FC direction (up or downregulated)

nakov_dr_GOEnrich = rownames(subset(nakov_DE_dr, rownames(nakov_DE_dr)%in%geneUniverse))
nakov_ur_GOEnrich = rownames(subset(nakov_DE_ur, rownames(nakov_DE_ur)%in%geneUniverse))

geneList_nakov_dr = factor(as.integer(geneUniverse %in% nakov_dr_GOEnrich))
names(geneList_nakov_dr) = geneUniverse
str(geneList_nakov_dr)
geneList_nakov_ur = factor(as.integer(geneUniverse %in% nakov_ur_GOEnrich))
names(geneList_nakov_ur) = geneUniverse
str(geneList_nakov_ur)

##create a topGO object (for up GOs)
GOdata_BP_nakov_dr = new("topGOdata", ontology="BP", allGenes=geneList_nakov_dr, 
                        annot = annFUN.gene2GO, gene2GO = geneID2GO)
GOdata_MF_nakov_dr = new("topGOdata", ontology="MF", allGenes=geneList_nakov_dr, 
                        annot = annFUN.gene2GO, gene2GO = geneID2GO)
GOdata_CC_nakov_dr = new("topGOdata", ontology="CC", allGenes=geneList_nakov_dr, 
                        annot = annFUN.gene2GO, gene2GO = geneID2GO)
##create a topGO object (for down GOs)
GOdata_BP_nakov_ur  = new("topGOdata", ontology="BP", allGenes=geneList_nakov_ur , 
                           annot = annFUN.gene2GO, gene2GO = geneID2GO)
GOdata_MF_nakov_ur  = new("topGOdata", ontology="MF", allGenes=geneList_nakov_ur , 
                           annot = annFUN.gene2GO, gene2GO = geneID2GO)
GOdata_CC_nakov_ur  = new("topGOdata", ontology="CC", allGenes=geneList_nakov_ur , 
                           annot = annFUN.gene2GO, gene2GO = geneID2GO)

#run Fisher's exact test
resultFisher_BP_nakov_dr_elim <- runTest(GOdata_BP_nakov_dr, algorithm = "elim", statistic = "fisher")
resultFisher_MF_nakov_dr_elim <- runTest(GOdata_MF_nakov_dr, algorithm = "elim", statistic = "fisher")
resultFisher_CC_nakov_dr_elim <- runTest(GOdata_CC_nakov_dr, algorithm = "elim", statistic = "fisher")
resultFisher_BP_nakov_ur_elim <- runTest(GOdata_BP_nakov_ur, algorithm = "elim", statistic = "fisher")
resultFisher_MF_nakov_ur_elim <- runTest(GOdata_MF_nakov_ur, algorithm = "elim", statistic = "fisher")
resultFisher_CC_nakov_ur_elim <- runTest(GOdata_CC_nakov_ur, algorithm = "elim", statistic = "fisher")

#extract the significant GO terms
nakov_dr_BP_elim <- GenTable(GOdata_BP_nakov_dr, classic = resultFisher_BP_nakov_dr_elim, 
                            orderBy = "weight", ranksOf = "weight", topNodes = 50)
nakov_dr_MF_elim <- GenTable(GOdata_MF_nakov_dr, classic = resultFisher_MF_nakov_dr_elim, 
                            orderBy = "weight", ranksOf = "weight", topNodes = 50)
nakov_dr_CC_elim <- GenTable(GOdata_CC_nakov_dr, classic = resultFisher_CC_nakov_dr_elim, 
                            orderBy = "weight", ranksOf = "weight", topNodes = 50)
nakov_ur_BP_elim <- GenTable(GOdata_BP_nakov_ur, classic = resultFisher_BP_nakov_ur_elim, 
                              orderBy = "weight", ranksOf = "weight", topNodes = 50)
nakov_ur_MF_elim <- GenTable(GOdata_MF_nakov_ur, classic = resultFisher_MF_nakov_ur_elim, 
                              orderBy = "weight", ranksOf = "weight", topNodes = 50)
nakov_ur_CC_elim <- GenTable(GOdata_CC_nakov_ur, classic = resultFisher_CC_nakov_ur_elim, 
                              orderBy = "weight", ranksOf = "weight", topNodes = 50)

#Generate the files for REVIGO input, which only needs a GO.ID and the p-value 
#Write the tables to you desired location to run through revigo using code below as a template.
#write.table(nakov_ur_BP_elim,
#            "/My Drive/ShortTermStress-IlluminaData/FullExp/Cryp-DE/data/ReviGo_files/BP_upLT_elim.txt")
#write.table(nakov_ur_MF_elim,
#            "/My Drive/ShortTermStress-IlluminaData/FullExp/Cryp-DE/data/ReviGo_files/MF_upLT_elim.txt")
#write.table(nakov_ur_CC_elim,
#            "/My Drive/ShortTermStress-IlluminaData/FullExp/Cryp-DE/data/ReviGo_files/CC_upLT_elim.txt")
#write.table(nakov_dr_BP_elim,
#            "/My Drive/ShortTermStress-IlluminaData/FullExp/Cryp-DE/data/ReviGo_files/BP_dnLT_elim.txt")
#write.table(nakov_dr_MF_elim,
#            "/My Drive/ShortTermStress-IlluminaData/FullExp/Cryp-DE/data/ReviGo_files/MF_dnLT_elim.txt")
#write.table(nakov_dr_CC_elim,
#            "/My Drive/ShortTermStress-IlluminaData/FullExp/Cryp-DE/data/ReviGo_files/CC_dnLT_elim.txt")

#import results from REVIGO back in if you want visualize them or use them in future analyses. The results were so small, no reduction was necessary
```

#### 4.4 Comparisons between peak stress, acclimation, and post-acclimation

Gene enrichment analyses for only the significant genes at 60 min and
10 hour timepoints, which we identified as the “peak” stress response
and the beginning of acclimation, separated by direction of regulation.
We then compile tables and figures to compare against long-term data

15 minute timepoint

```
#Select only the genes significant at a timepoint from the reduced subset of 2 consecutive gene. This can be done for all time points, but is limited here to the ones discussed extensively in the manuscript
sign_0v15m <- rownames(keep_2_SignDEG)[keep_2_SignDEG[,"C0m-C60m"]==1] 

length(sign_0v15m) #3497 and this matches the output from SignifGenes
```

```
## [1] 3412
```

```
sign2LFC_0v15m <- sign2LFC [rownames(sign2LFC) %in% sign_0v15m, ] #3019 rows, which means one row had NAs and got omitted. 
logFCup_0v15m <- subset(sign2LFC_0v15m, (`0v15m_Padj`) <= 0.01 & (`0v15m`) > 0)
logFCdown_0v15m <- subset(sign2LFC_0v15m, (`0v15m_Padj`) <= 0.01 & (`0v15m`) < 0)

logFCup_0v15m_GOEnrich  = rownames(subset(logFCup_0v15m, rownames(logFCup_0v15m)%in%genesOfInterest_allSign))
logFCdown_0v15m_GOEnrich  = rownames(subset(logFCdown_0v15m, rownames(logFCdown_0v15m)%in%genesOfInterest_allSign))


#create gene list for input in topGO for 30 minutes
geneList_up0v15m = factor(as.integer(geneUniverse %in% logFCup_0v15m_GOEnrich))
names(geneList_up0v15m) = geneUniverse
str(geneList_up0v15m)
```

```
##  Factor w/ 2 levels "0","1": 1 1 1 1 1 1 1 1 1 1 ...
##  - attr(*, "names")= chr [1:6309] "gene_idGO.ID" "CCRYP_013698" "CCRYP_013706" "CCRYP_013704" ...
```

```
geneList_down0v15m = factor(as.integer(geneUniverse %in% logFCdown_0v15m_GOEnrich))
names(geneList_down0v15m) = geneUniverse
str(geneList_down0v15m)
```

```
##  Factor w/ 2 levels "0","1": 1 1 1 1 1 1 1 2 1 2 ...
##  - attr(*, "names")= chr [1:6309] "gene_idGO.ID" "CCRYP_013698" "CCRYP_013706" "CCRYP_013704" ...
```

```
##create a topGO object (for up GOs)
GOdata_BP_up0v15m = new("topGOdata", ontology="BP", allGenes=geneList_up0v15m, 
                        annot = annFUN.gene2GO, gene2GO = geneID2GO)
```

```
## 
## Building most specific GOs .....
```

```
##  ( 624 GO terms found. )
```

```
## 
## Build GO DAG topology ..........
```

```
##  ( 1729 GO terms and 3515 relations. )
```

```
## 
## Annotating nodes ...............
```

```
##  ( 3345 genes annotated to the GO terms. )
```

```
GOdata_MF_up0v15m = new("topGOdata", ontology="MF", allGenes=geneList_up0v15m, 
                        annot = annFUN.gene2GO, gene2GO = geneID2GO)
```

```
## 
## Building most specific GOs .....
```

```
##  ( 736 GO terms found. )
```

```
## 
## Build GO DAG topology ..........
```

```
##  ( 1096 GO terms and 1406 relations. )
```

```
## 
## Annotating nodes ...............
```

```
##  ( 5189 genes annotated to the GO terms. )
```

```
GOdata_CC_up0v15m = new("topGOdata", ontology="CC", allGenes=geneList_up0v15m, 
                        annot = annFUN.gene2GO, gene2GO = geneID2GO)
```

```
## 
## Building most specific GOs .....
```

```
##  ( 235 GO terms found. )
```

```
## 
## Build GO DAG topology ..........
```

```
##  ( 452 GO terms and 812 relations. )
```

```
## 
## Annotating nodes ...............
```

```
##  ( 1987 genes annotated to the GO terms. )
```

```
##create a topGO object (for down GOs)
GOdata_BP_down0v15m  = new("topGOdata", ontology="BP", allGenes=geneList_down0v15m , 
                           annot = annFUN.gene2GO, gene2GO = geneID2GO)
```

```
## 
## Building most specific GOs .....
```

```
##  ( 624 GO terms found. )
```

```
## 
## Build GO DAG topology ..........
```

```
##  ( 1729 GO terms and 3515 relations. )
```

```
## 
## Annotating nodes ...............
```

```
##  ( 3345 genes annotated to the GO terms. )
```

```
GOdata_MF_down0v15m  = new("topGOdata", ontology="MF", allGenes=geneList_down0v15m , 
                           annot = annFUN.gene2GO, gene2GO = geneID2GO)
```

```
## 
## Building most specific GOs .....
```

```
##  ( 736 GO terms found. )
```

```
## 
## Build GO DAG topology ..........
```

```
##  ( 1096 GO terms and 1406 relations. )
```

```
## 
## Annotating nodes ...............
```

```
##  ( 5189 genes annotated to the GO terms. )
```

```
GOdata_CC_down0v15m  = new("topGOdata", ontology="CC", allGenes=geneList_down0v15m , 
                           annot = annFUN.gene2GO, gene2GO = geneID2GO)
```

```
## 
## Building most specific GOs .....
```

```
##  ( 235 GO terms found. )
```

```
## 
## Build GO DAG topology ..........
```

```
##  ( 452 GO terms and 812 relations. )
```

```
## 
## Annotating nodes ...............
```

```
##  ( 1987 genes annotated to the GO terms. )
```

```
#run Fisher's exact test
resultFisher_BP_up0v15m_elim <- runTest(GOdata_BP_up0v15m, algorithm = "elim", statistic = "fisher")
```

```
## 
##           -- Elim Algorithm -- 
## 
##       the algorithm is scoring 353 nontrivial nodes
##       parameters: 
##           test statistic: fisher
##           cutOff: 0.01
```

```
## 
##   Level 13:  3 nodes to be scored    (0 eliminated genes)
```

```
## 
##   Level 12:  5 nodes to be scored    (0 eliminated genes)
```

```
## 
##   Level 11:  12 nodes to be scored   (0 eliminated genes)
```

```
## 
##   Level 10:  27 nodes to be scored   (0 eliminated genes)
```

```
## 
##   Level 9:   39 nodes to be scored   (0 eliminated genes)
```

```
## 
##   Level 8:   38 nodes to be scored   (3 eliminated genes)
```

```
## 
##   Level 7:   41 nodes to be scored   (3 eliminated genes)
```

```
## 
##   Level 6:   54 nodes to be scored   (6 eliminated genes)
```

```
## 
##   Level 5:   63 nodes to be scored   (51 eliminated genes)
```

```
## 
##   Level 4:   38 nodes to be scored   (51 eliminated genes)
```

```
## 
##   Level 3:   25 nodes to be scored   (51 eliminated genes)
```

```
## 
##   Level 2:   7 nodes to be scored    (51 eliminated genes)
```

```
## 
##   Level 1:   1 nodes to be scored    (51 eliminated genes)
```

```
resultFisher_MF_up0v15m_elim <- runTest(GOdata_MF_up0v15m, algorithm = "elim", statistic = "fisher")
```

```
## 
##           -- Elim Algorithm -- 
## 
##       the algorithm is scoring 199 nontrivial nodes
##       parameters: 
##           test statistic: fisher
##           cutOff: 0.01
```

```
## 
##   Level 9:   4 nodes to be scored    (0 eliminated genes)
```

```
## 
##   Level 8:   11 nodes to be scored   (3 eliminated genes)
```

```
## 
##   Level 7:   23 nodes to be scored   (3 eliminated genes)
```

```
## 
##   Level 6:   39 nodes to be scored   (6 eliminated genes)
```

```
## 
##   Level 5:   47 nodes to be scored   (6 eliminated genes)
```

```
## 
##   Level 4:   46 nodes to be scored   (55 eliminated genes)
```

```
## 
##   Level 3:   21 nodes to be scored   (55 eliminated genes)
```

```
## 
##   Level 2:   7 nodes to be scored    (112 eliminated genes)
```

```
## 
##   Level 1:   1 nodes to be scored    (112 eliminated genes)
```

```
resultFisher_CC_up0v15m_elim <- runTest(GOdata_CC_up0v15m, algorithm = "elim", statistic = "fisher")
```

```
## 
##           -- Elim Algorithm -- 
## 
##       the algorithm is scoring 73 nontrivial nodes
##       parameters: 
##           test statistic: fisher
##           cutOff: 0.01
```

```
## 
##   Level 10:  1 nodes to be scored    (0 eliminated genes)
```

```
## 
##   Level 9:   3 nodes to be scored    (0 eliminated genes)
```

```
## 
##   Level 8:   7 nodes to be scored    (0 eliminated genes)
```

```
## 
##   Level 7:   8 nodes to be scored    (0 eliminated genes)
```

```
## 
##   Level 6:   13 nodes to be scored   (0 eliminated genes)
```

```
## 
##   Level 5:   12 nodes to be scored   (0 eliminated genes)
```

```
## 
##   Level 4:   14 nodes to be scored   (0 eliminated genes)
```

```
## 
##   Level 3:   12 nodes to be scored   (0 eliminated genes)
```

```
## 
##   Level 2:   2 nodes to be scored    (41 eliminated genes)
```

```
## 
##   Level 1:   1 nodes to be scored    (41 eliminated genes)
```

```
resultFisher_BP_down0v15m_elim <- runTest(GOdata_BP_down0v15m, algorithm = "elim", statistic = "fisher")
```

```
## 
##           -- Elim Algorithm -- 
## 
##       the algorithm is scoring 526 nontrivial nodes
##       parameters: 
##           test statistic: fisher
##           cutOff: 0.01
```

```
## 
##   Level 14:  2 nodes to be scored    (0 eliminated genes)
```

```
## 
##   Level 13:  5 nodes to be scored    (0 eliminated genes)
```

```
## 
##   Level 12:  10 nodes to be scored   (0 eliminated genes)
```

```
## 
##   Level 11:  18 nodes to be scored   (0 eliminated genes)
```

```
## 
##   Level 10:  40 nodes to be scored   (0 eliminated genes)
```

```
## 
##   Level 9:   59 nodes to be scored   (7 eliminated genes)
```

```
## 
##   Level 8:   66 nodes to be scored   (7 eliminated genes)
```

```
## 
##   Level 7:   70 nodes to be scored   (46 eliminated genes)
```

```
## 
##   Level 6:   83 nodes to be scored   (46 eliminated genes)
```

```
## 
##   Level 5:   84 nodes to be scored   (338 eliminated genes)
```

```
## 
##   Level 4:   49 nodes to be scored   (338 eliminated genes)
```

```
## 
##   Level 3:   32 nodes to be scored   (338 eliminated genes)
```

```
## 
##   Level 2:   7 nodes to be scored    (368 eliminated genes)
```

```
## 
##   Level 1:   1 nodes to be scored    (368 eliminated genes)
```

```
resultFisher_MF_down0v15m_elim <- runTest(GOdata_MF_down0v15m, algorithm = "elim", statistic = "fisher")
```

```
## 
##           -- Elim Algorithm -- 
## 
##       the algorithm is scoring 330 nontrivial nodes
##       parameters: 
##           test statistic: fisher
##           cutOff: 0.01
```

```
## 
##   Level 11:  1 nodes to be scored    (0 eliminated genes)
```

```
## 
##   Level 10:  1 nodes to be scored    (7 eliminated genes)
```

```
## 
##   Level 9:   6 nodes to be scored    (7 eliminated genes)
```

```
## 
##   Level 8:   17 nodes to be scored   (7 eliminated genes)
```

```
## 
##   Level 7:   42 nodes to be scored   (7 eliminated genes)
```

```
## 
##   Level 6:   79 nodes to be scored   (98 eliminated genes)
```

```
## 
##   Level 5:   78 nodes to be scored   (100 eliminated genes)
```

```
## 
##   Level 4:   64 nodes to be scored   (141 eliminated genes)
```

```
## 
##   Level 3:   30 nodes to be scored   (231 eliminated genes)
```

```
## 
##   Level 2:   11 nodes to be scored   (1310 eliminated genes)
```

```
## 
##   Level 1:   1 nodes to be scored    (1310 eliminated genes)
```

```
resultFisher_CC_down0v15m_elim <- runTest(GOdata_CC_down0v15m, algorithm = "elim", statistic = "fisher")
```

```
## 
##           -- Elim Algorithm -- 
## 
##       the algorithm is scoring 141 nontrivial nodes
##       parameters: 
##           test statistic: fisher
##           cutOff: 0.01
```

```
## 
##   Level 12:  1 nodes to be scored    (0 eliminated genes)
```

```
## 
##   Level 11:  4 nodes to be scored    (0 eliminated genes)
```

```
## 
##   Level 10:  8 nodes to be scored    (0 eliminated genes)
```

```
## 
##   Level 9:   12 nodes to be scored   (0 eliminated genes)
```

```
## 
##   Level 8:   20 nodes to be scored   (3 eliminated genes)
```

```
## 
##   Level 7:   19 nodes to be scored   (6 eliminated genes)
```

```
## 
##   Level 6:   21 nodes to be scored   (6 eliminated genes)
```

```
## 
##   Level 5:   18 nodes to be scored   (31 eliminated genes)
```

```
## 
##   Level 4:   18 nodes to be scored   (31 eliminated genes)
```

```
## 
##   Level 3:   17 nodes to be scored   (338 eliminated genes)
```

```
## 
##   Level 2:   2 nodes to be scored    (338 eliminated genes)
```

```
## 
##   Level 1:   1 nodes to be scored    (338 eliminated genes)
```

```
#extract the significant GO terms
up0v15m_BP_elim <- GenTable(GOdata_BP_up0v15m, classic = resultFisher_BP_up0v15m_elim, 
                            orderBy = "weight", ranksOf = "weight", topNodes = 50)
up0v15m_MF_elim <- GenTable(GOdata_MF_up0v15m, classic = resultFisher_MF_up0v15m_elim, 
                            orderBy = "weight", ranksOf = "weight", topNodes = 50)
up0v15m_CC_elim <- GenTable(GOdata_CC_up0v15m, classic = resultFisher_CC_up0v15m_elim, 
                            orderBy = "weight", ranksOf = "weight", topNodes = 50)
down0v15m_BP_elim <- GenTable(GOdata_BP_down0v15m, classic = resultFisher_BP_down0v15m_elim, 
                              orderBy = "weight", ranksOf = "weight", topNodes = 50)
down0v15m_MF_elim <- GenTable(GOdata_MF_down0v15m, classic = resultFisher_MF_down0v15m_elim, 
                              orderBy = "weight", ranksOf = "weight", topNodes = 50)
down0v15m_CC_elim <- GenTable(GOdata_CC_down0v15m, classic = resultFisher_CC_down0v15m_elim, 
                              orderBy = "weight", ranksOf = "weight", topNodes = 50)

#Write the tables to you desired location to run through revigo using code below as a template.
#write.table(up0v15m_BP_elim,
#            "/My Drive/ShortTermStress-IlluminaData/FullExp/Cryp-DE/data/ReviGo_files/BP_up15m_elim.txt")
#write.table(up0v15m_MF_elim,
#            "/My Drive/ShortTermStress-IlluminaData/FullExp/Cryp-DE/data/ReviGo_files/MF_up15m_elim.txt")
#write.table(up0v15m_CC_elim,
#            "/My Drive/ShortTermStress-IlluminaData/FullExp/Cryp-DE/data/ReviGo_files/CC_up15m_elim.txt")
#write.table(down0v15m_BP_elim,
#            "/My Drive/ShortTermStress-IlluminaData/FullExp/Cryp-DE/data/ReviGo_files/BP_dn15m_elim.txt")
#write.table(down0v15m_MF_elim,
#            "/My Drive/ShortTermStress-IlluminaData/FullExp/Cryp-DE/data/ReviGo_files/MF_dn15m_elim.txt")
#write.table(down0v15m_CC_elim,
#            "/My Drive/ShortTermStress-IlluminaData/FullExp/Cryp-DE/data/ReviGo_files/CC_dn15m_elim.txt")
```

30 minute timepoint

```
#Select only the genes significant at a timepoint from the reduced subset of 2 consecutive gene. This can be done for all time points, but is limited here to the ones discussed extensively in the manuscript
sign_0v30m <- rownames(keep_2_SignDEG)[keep_2_SignDEG[,"C0m-C60m"]==1] 

length(sign_0v30m) #3497 and this matches the output from SignifGenes
```

```
## [1] 3412
```

```
sign2LFC_0v30m <- sign2LFC [rownames(sign2LFC) %in% sign_0v30m, ] #3019 rows, which means one row had NAs and got omitted. 
logFCup_0v30m <- subset(sign2LFC_0v30m, (`0v30m_Padj`) <= 0.01 & (`0v30m`) > 0)
logFCdown_0v30m <- subset(sign2LFC_0v30m, (`0v30m_Padj`) <= 0.01 & (`0v30m`) < 0)

logFCup_0v30m_GOEnrich  = rownames(subset(logFCup_0v30m, rownames(logFCup_0v30m)%in%genesOfInterest_allSign))
logFCdown_0v30m_GOEnrich  = rownames(subset(logFCdown_0v30m, rownames(logFCdown_0v30m)%in%genesOfInterest_allSign))


#create gene list for input in topGO for 30 minutes
geneList_up0v30m = factor(as.integer(geneUniverse %in% logFCup_0v30m_GOEnrich))
names(geneList_up0v30m) = geneUniverse
str(geneList_up0v30m)
```

```
##  Factor w/ 2 levels "0","1": 1 1 1 1 1 1 1 1 2 1 ...
##  - attr(*, "names")= chr [1:6309] "gene_idGO.ID" "CCRYP_013698" "CCRYP_013706" "CCRYP_013704" ...
```

```
geneList_down0v30m = factor(as.integer(geneUniverse %in% logFCdown_0v30m_GOEnrich))
names(geneList_down0v30m) = geneUniverse
str(geneList_down0v30m)
```

```
##  Factor w/ 2 levels "0","1": 1 1 1 1 2 1 1 2 1 2 ...
##  - attr(*, "names")= chr [1:6309] "gene_idGO.ID" "CCRYP_013698" "CCRYP_013706" "CCRYP_013704" ...
```

```
##create a topGO object (for up GOs)
GOdata_BP_up0v30m = new("topGOdata", ontology="BP", allGenes=geneList_up0v30m, 
                        annot = annFUN.gene2GO, gene2GO = geneID2GO)
```

```
## 
## Building most specific GOs .....
```

```
##  ( 624 GO terms found. )
```

```
## 
## Build GO DAG topology ..........
```

```
##  ( 1729 GO terms and 3515 relations. )
```

```
## 
## Annotating nodes ...............
```

```
##  ( 3345 genes annotated to the GO terms. )
```

```
GOdata_MF_up0v30m = new("topGOdata", ontology="MF", allGenes=geneList_up0v30m, 
                        annot = annFUN.gene2GO, gene2GO = geneID2GO)
```

```
## 
## Building most specific GOs .....
```

```
##  ( 736 GO terms found. )
```

```
## 
## Build GO DAG topology ..........
```

```
##  ( 1096 GO terms and 1406 relations. )
```

```
## 
## Annotating nodes ...............
```

```
##  ( 5189 genes annotated to the GO terms. )
```

```
GOdata_CC_up0v30m = new("topGOdata", ontology="CC", allGenes=geneList_up0v30m, 
                        annot = annFUN.gene2GO, gene2GO = geneID2GO)
```

```
## 
## Building most specific GOs .....
```

```
##  ( 235 GO terms found. )
```

```
## 
## Build GO DAG topology ..........
```

```
##  ( 452 GO terms and 812 relations. )
```

```
## 
## Annotating nodes ...............
```

```
##  ( 1987 genes annotated to the GO terms. )
```

```
##create a topGO object (for down GOs)
GOdata_BP_down0v30m  = new("topGOdata", ontology="BP", allGenes=geneList_down0v30m , 
                           annot = annFUN.gene2GO, gene2GO = geneID2GO)
```

```
## 
## Building most specific GOs .....
```

```
##  ( 624 GO terms found. )
```

```
## 
## Build GO DAG topology ..........
```

```
##  ( 1729 GO terms and 3515 relations. )
```

```
## 
## Annotating nodes ...............
```

```
##  ( 3345 genes annotated to the GO terms. )
```

```
GOdata_MF_down0v30m  = new("topGOdata", ontology="MF", allGenes=geneList_down0v30m , 
                           annot = annFUN.gene2GO, gene2GO = geneID2GO)
```

```
## 
## Building most specific GOs .....
```

```
##  ( 736 GO terms found. )
```

```
## 
## Build GO DAG topology ..........
```

```
##  ( 1096 GO terms and 1406 relations. )
```

```
## 
## Annotating nodes ...............
```

```
##  ( 5189 genes annotated to the GO terms. )
```

```
GOdata_CC_down0v30m  = new("topGOdata", ontology="CC", allGenes=geneList_down0v30m , 
                           annot = annFUN.gene2GO, gene2GO = geneID2GO)
```

```
## 
## Building most specific GOs .....
```

```
##  ( 235 GO terms found. )
```

```
## 
## Build GO DAG topology ..........
```

```
##  ( 452 GO terms and 812 relations. )
```

```
## 
## Annotating nodes ...............
```

```
##  ( 1987 genes annotated to the GO terms. )
```

```
#run Fisher's exact test
resultFisher_BP_up0v30m_elim <- runTest(GOdata_BP_up0v30m, algorithm = "elim", statistic = "fisher")
```

```
## 
##           -- Elim Algorithm -- 
## 
##       the algorithm is scoring 554 nontrivial nodes
##       parameters: 
##           test statistic: fisher
##           cutOff: 0.01
```

```
## 
##   Level 14:  1 nodes to be scored    (0 eliminated genes)
```

```
## 
##   Level 13:  3 nodes to be scored    (0 eliminated genes)
```

```
## 
##   Level 12:  12 nodes to be scored   (0 eliminated genes)
```

```
## 
##   Level 11:  20 nodes to be scored   (0 eliminated genes)
```

```
## 
##   Level 10:  39 nodes to be scored   (0 eliminated genes)
```

```
## 
##   Level 9:   57 nodes to be scored   (31 eliminated genes)
```

```
## 
##   Level 8:   70 nodes to be scored   (40 eliminated genes)
```

```
## 
##   Level 7:   80 nodes to be scored   (52 eliminated genes)
```

```
## 
##   Level 6:   99 nodes to be scored   (63 eliminated genes)
```

```
## 
##   Level 5:   86 nodes to be scored   (201 eliminated genes)
```

```
## 
##   Level 4:   50 nodes to be scored   (202 eliminated genes)
```

```
## 
##   Level 3:   29 nodes to be scored   (202 eliminated genes)
```

```
## 
##   Level 2:   7 nodes to be scored    (202 eliminated genes)
```

```
## 
##   Level 1:   1 nodes to be scored    (202 eliminated genes)
```

```
resultFisher_MF_up0v30m_elim <- runTest(GOdata_MF_up0v30m, algorithm = "elim", statistic = "fisher")
```

```
## 
##           -- Elim Algorithm -- 
## 
##       the algorithm is scoring 406 nontrivial nodes
##       parameters: 
##           test statistic: fisher
##           cutOff: 0.01
```

```
## 
##   Level 11:  1 nodes to be scored    (0 eliminated genes)
```

```
## 
##   Level 10:  3 nodes to be scored    (0 eliminated genes)
```

```
## 
##   Level 9:   17 nodes to be scored   (0 eliminated genes)
```

```
## 
##   Level 8:   29 nodes to be scored   (2 eliminated genes)
```

```
## 
##   Level 7:   52 nodes to be scored   (29 eliminated genes)
```

```
## 
##   Level 6:   92 nodes to be scored   (41 eliminated genes)
```

```
## 
##   Level 5:   92 nodes to be scored   (55 eliminated genes)
```

```
## 
##   Level 4:   78 nodes to be scored   (104 eliminated genes)
```

```
## 
##   Level 3:   30 nodes to be scored   (274 eliminated genes)
```

```
## 
##   Level 2:   11 nodes to be scored   (1375 eliminated genes)
```

```
## 
##   Level 1:   1 nodes to be scored    (1375 eliminated genes)
```

```
resultFisher_CC_up0v30m_elim <- runTest(GOdata_CC_up0v30m, algorithm = "elim", statistic = "fisher")
```

```
## 
##           -- Elim Algorithm -- 
## 
##       the algorithm is scoring 141 nontrivial nodes
##       parameters: 
##           test statistic: fisher
##           cutOff: 0.01
```

```
## 
##   Level 11:  2 nodes to be scored    (0 eliminated genes)
```

```
## 
##   Level 10:  6 nodes to be scored    (0 eliminated genes)
```

```
## 
##   Level 9:   11 nodes to be scored   (0 eliminated genes)
```

```
## 
##   Level 8:   17 nodes to be scored   (0 eliminated genes)
```

```
## 
##   Level 7:   27 nodes to be scored   (0 eliminated genes)
```

```
## 
##   Level 6:   28 nodes to be scored   (7 eliminated genes)
```

```
## 
##   Level 5:   16 nodes to be scored   (7 eliminated genes)
```

```
## 
##   Level 4:   16 nodes to be scored   (478 eliminated genes)
```

```
## 
##   Level 3:   15 nodes to be scored   (478 eliminated genes)
```

```
## 
##   Level 2:   2 nodes to be scored    (844 eliminated genes)
```

```
## 
##   Level 1:   1 nodes to be scored    (844 eliminated genes)
```

```
resultFisher_BP_down0v30m_elim <- runTest(GOdata_BP_down0v30m, algorithm = "elim", statistic = "fisher")
```

```
## 
##           -- Elim Algorithm -- 
## 
##       the algorithm is scoring 901 nontrivial nodes
##       parameters: 
##           test statistic: fisher
##           cutOff: 0.01
```

```
## 
##   Level 14:  4 nodes to be scored    (0 eliminated genes)
```

```
## 
##   Level 13:  11 nodes to be scored   (0 eliminated genes)
```

```
## 
##   Level 12:  36 nodes to be scored   (41 eliminated genes)
```

```
## 
##   Level 11:  52 nodes to be scored   (41 eliminated genes)
```

```
## 
##   Level 10:  90 nodes to be scored   (80 eliminated genes)
```

```
## 
##   Level 9:   117 nodes to be scored  (120 eliminated genes)
```

```
## 
##   Level 8:   125 nodes to be scored  (138 eliminated genes)
```

```
## 
##   Level 7:   129 nodes to be scored  (308 eliminated genes)
```

```
## 
##   Level 6:   122 nodes to be scored  (401 eliminated genes)
```

```
## 
##   Level 5:   110 nodes to be scored  (427 eliminated genes)
```

```
## 
##   Level 4:   59 nodes to be scored   (435 eliminated genes)
```

```
## 
##   Level 3:   38 nodes to be scored   (435 eliminated genes)
```

```
## 
##   Level 2:   7 nodes to be scored    (502 eliminated genes)
```

```
## 
##   Level 1:   1 nodes to be scored    (2785 eliminated genes)
```

```
resultFisher_MF_down0v30m_elim <- runTest(GOdata_MF_down0v30m, algorithm = "elim", statistic = "fisher")
```

```
## 
##           -- Elim Algorithm -- 
## 
##       the algorithm is scoring 495 nontrivial nodes
##       parameters: 
##           test statistic: fisher
##           cutOff: 0.01
```

```
## 
##   Level 10:  4 nodes to be scored    (0 eliminated genes)
```

```
## 
##   Level 9:   15 nodes to be scored   (0 eliminated genes)
```

```
## 
##   Level 8:   29 nodes to be scored   (758 eliminated genes)
```

```
## 
##   Level 7:   77 nodes to be scored   (789 eliminated genes)
```

```
## 
##   Level 6:   133 nodes to be scored  (815 eliminated genes)
```

```
## 
##   Level 5:   111 nodes to be scored  (822 eliminated genes)
```

```
## 
##   Level 4:   81 nodes to be scored   (1013 eliminated genes)
```

```
## 
##   Level 3:   33 nodes to be scored   (1030 eliminated genes)
```

```
## 
##   Level 2:   11 nodes to be scored   (1118 eliminated genes)
```

```
## 
##   Level 1:   1 nodes to be scored    (1138 eliminated genes)
```

```
resultFisher_CC_down0v30m_elim <- runTest(GOdata_CC_down0v30m, algorithm = "elim", statistic = "fisher")
```

```
## 
##           -- Elim Algorithm -- 
## 
##       the algorithm is scoring 247 nontrivial nodes
##       parameters: 
##           test statistic: fisher
##           cutOff: 0.01
```

```
## 
##   Level 12:  2 nodes to be scored    (0 eliminated genes)
```

```
## 
##   Level 11:  10 nodes to be scored   (0 eliminated genes)
```

```
## 
##   Level 10:  14 nodes to be scored   (0 eliminated genes)
```

```
## 
##   Level 9:   27 nodes to be scored   (0 eliminated genes)
```

```
## 
##   Level 8:   37 nodes to be scored   (3 eliminated genes)
```

```
## 
##   Level 7:   37 nodes to be scored   (25 eliminated genes)
```

```
## 
##   Level 6:   37 nodes to be scored   (54 eliminated genes)
```

```
## 
##   Level 5:   28 nodes to be scored   (146 eliminated genes)
```

```
## 
##   Level 4:   29 nodes to be scored   (165 eliminated genes)
```

```
## 
##   Level 3:   23 nodes to be scored   (423 eliminated genes)
```

```
## 
##   Level 2:   2 nodes to be scored    (1159 eliminated genes)
```

```
## 
##   Level 1:   1 nodes to be scored    (1159 eliminated genes)
```

```
#extract the significant GO terms
up0v30m_BP_elim <- GenTable(GOdata_BP_up0v30m, classic = resultFisher_BP_up0v30m_elim, 
                            orderBy = "weight", ranksOf = "weight", topNodes = 50)
up0v30m_MF_elim <- GenTable(GOdata_MF_up0v30m, classic = resultFisher_MF_up0v30m_elim, 
                            orderBy = "weight", ranksOf = "weight", topNodes = 50)
up0v30m_CC_elim <- GenTable(GOdata_CC_up0v30m, classic = resultFisher_CC_up0v30m_elim, 
                            orderBy = "weight", ranksOf = "weight", topNodes = 50)
down0v30m_BP_elim <- GenTable(GOdata_BP_down0v30m, classic = resultFisher_BP_down0v30m_elim, 
                              orderBy = "weight", ranksOf = "weight", topNodes = 50)
down0v30m_MF_elim <- GenTable(GOdata_MF_down0v30m, classic = resultFisher_MF_down0v30m_elim, 
                              orderBy = "weight", ranksOf = "weight", topNodes = 50)
down0v30m_CC_elim <- GenTable(GOdata_CC_down0v30m, classic = resultFisher_CC_down0v30m_elim, 
                              orderBy = "weight", ranksOf = "weight", topNodes = 50)

#Write the tables to you desired location to run through revigo using code below as a template.
#write.table(up0v30m_BP_elim,
#            "/My Drive/ShortTermStress-IlluminaData/FullExp/Cryp-DE/data/ReviGo_files/BP_up30m_elim.txt")
#write.table(up0v30m_MF_elim,
#            "/My Drive/ShortTermStress-IlluminaData/FullExp/Cryp-DE/data/ReviGo_files/MF_up30m_elim.txt")
#write.table(up0v30m_CC_elim,
#            "/My Drive/ShortTermStress-IlluminaData/FullExp/Cryp-DE/data/ReviGo_files/CC_up30m_elim.txt")
#write.table(down0v30m_BP_elim,
#            "/My Drive/ShortTermStress-IlluminaData/FullExp/Cryp-DE/data/ReviGo_files/BP_dn30m_elim.txt")
#write.table(down0v30m_MF_elim,
#            "/My Drive/ShortTermStress-IlluminaData/FullExp/Cryp-DE/data/ReviGo_files/MF_dn30m_elim.txt")
#write.table(down0v30m_CC_elim,
#            "/My Drive/ShortTermStress-IlluminaData/FullExp/Cryp-DE/data/ReviGo_files/CC_dn30m_elim.txt")
```

1 hour timepoint; peak period of low salinity stress

```
#Select only the genes significant at a timepoint from the reduced subset of 2 consecutive gene. This can be done for all time points, but is limited here to the ones discussed extensively in the manuscript
sign_0v1h <- rownames(keep_2_SignDEG)[keep_2_SignDEG[,"C0m-C60m"]==1] 

length(sign_0v1h) #3497 and this matches the output from SignifGenes
```

```
## [1] 3412
```

```
sign2LFC_0v1h <- sign2LFC [rownames(sign2LFC) %in% sign_0v1h, ] #3019 rows, which means one row had NAs and got omitted. 
logFCup_0v1h <- subset(sign2LFC_0v1h, (`0v1h_Padj`) <= 0.01 & (`0v1h`) > 0)
logFCdown_0v1h <- subset(sign2LFC_0v1h, (`0v1h_Padj`) <= 0.01 & (`0v1h`) < 0)

logFCup_0v1h_GOEnrich  = rownames(subset(logFCup_0v1h, rownames(logFCup_0v1h)%in%genesOfInterest_allSign))
logFCdown_0v1h_GOEnrich  = rownames(subset(logFCdown_0v1h, rownames(logFCdown_0v1h)%in%genesOfInterest_allSign))


#create gene list for input in topGO for 30 minutes
geneList_up0v1h = factor(as.integer(geneUniverse %in% logFCup_0v1h_GOEnrich))
names(geneList_up0v1h) = geneUniverse
str(geneList_up0v1h)
```

```
##  Factor w/ 2 levels "0","1": 1 1 1 1 1 1 1 1 2 1 ...
##  - attr(*, "names")= chr [1:6309] "gene_idGO.ID" "CCRYP_013698" "CCRYP_013706" "CCRYP_013704" ...
```

```
geneList_down0v1h = factor(as.integer(geneUniverse %in% logFCdown_0v1h_GOEnrich))
names(geneList_down0v1h) = geneUniverse
str(geneList_down0v1h)
```

```
##  Factor w/ 2 levels "0","1": 1 1 1 1 2 1 1 2 1 2 ...
##  - attr(*, "names")= chr [1:6309] "gene_idGO.ID" "CCRYP_013698" "CCRYP_013706" "CCRYP_013704" ...
```

```
##create a topGO object (for up GOs)
GOdata_BP_up0v1h = new("topGOdata", ontology="BP", allGenes=geneList_up0v1h, 
                        annot = annFUN.gene2GO, gene2GO = geneID2GO)
```

```
## 
## Building most specific GOs .....
```

```
##  ( 624 GO terms found. )
```

```
## 
## Build GO DAG topology ..........
```

```
##  ( 1729 GO terms and 3515 relations. )
```

```
## 
## Annotating nodes ...............
```

```
##  ( 3345 genes annotated to the GO terms. )
```

```
GOdata_MF_up0v1h = new("topGOdata", ontology="MF", allGenes=geneList_up0v1h, 
                        annot = annFUN.gene2GO, gene2GO = geneID2GO)
```

```
## 
## Building most specific GOs .....
```

```
##  ( 736 GO terms found. )
```

```
## 
## Build GO DAG topology ..........
```

```
##  ( 1096 GO terms and 1406 relations. )
```

```
## 
## Annotating nodes ...............
```

```
##  ( 5189 genes annotated to the GO terms. )
```

```
GOdata_CC_up0v1h = new("topGOdata", ontology="CC", allGenes=geneList_up0v1h, 
                        annot = annFUN.gene2GO, gene2GO = geneID2GO)
```

```
## 
## Building most specific GOs .....
```

```
##  ( 235 GO terms found. )
```

```
## 
## Build GO DAG topology ..........
```

```
##  ( 452 GO terms and 812 relations. )
```

```
## 
## Annotating nodes ...............
```

```
##  ( 1987 genes annotated to the GO terms. )
```

```
##create a topGO object (for down GOs)
GOdata_BP_down0v1h  = new("topGOdata", ontology="BP", allGenes=geneList_down0v1h , 
                           annot = annFUN.gene2GO, gene2GO = geneID2GO)
```

```
## 
## Building most specific GOs .....
```

```
##  ( 624 GO terms found. )
```

```
## 
## Build GO DAG topology ..........
```

```
##  ( 1729 GO terms and 3515 relations. )
```

```
## 
## Annotating nodes ...............
```

```
##  ( 3345 genes annotated to the GO terms. )
```

```
GOdata_MF_down0v1h  = new("topGOdata", ontology="MF", allGenes=geneList_down0v1h , 
                           annot = annFUN.gene2GO, gene2GO = geneID2GO)
```

```
## 
## Building most specific GOs .....
```

```
##  ( 736 GO terms found. )
```

```
## 
## Build GO DAG topology ..........
```

```
##  ( 1096 GO terms and 1406 relations. )
```

```
## 
## Annotating nodes ...............
```

```
##  ( 5189 genes annotated to the GO terms. )
```

```
GOdata_CC_down0v1h  = new("topGOdata", ontology="CC", allGenes=geneList_down0v1h , 
                           annot = annFUN.gene2GO, gene2GO = geneID2GO)
```

```
## 
## Building most specific GOs .....
```

```
##  ( 235 GO terms found. )
```

```
## 
## Build GO DAG topology ..........
```

```
##  ( 452 GO terms and 812 relations. )
```

```
## 
## Annotating nodes ...............
```

```
##  ( 1987 genes annotated to the GO terms. )
```

```
#run Fisher's exact test
resultFisher_BP_up0v1h_elim <- runTest(GOdata_BP_up0v1h, algorithm = "elim", statistic = "fisher")
```

```
## 
##           -- Elim Algorithm -- 
## 
##       the algorithm is scoring 563 nontrivial nodes
##       parameters: 
##           test statistic: fisher
##           cutOff: 0.01
```

```
## 
##   Level 14:  1 nodes to be scored    (0 eliminated genes)
```

```
## 
##   Level 13:  5 nodes to be scored    (0 eliminated genes)
```

```
## 
##   Level 12:  14 nodes to be scored   (0 eliminated genes)
```

```
## 
##   Level 11:  23 nodes to be scored   (0 eliminated genes)
```

```
## 
##   Level 10:  42 nodes to be scored   (0 eliminated genes)
```

```
## 
##   Level 9:   61 nodes to be scored   (31 eliminated genes)
```

```
## 
##   Level 8:   68 nodes to be scored   (31 eliminated genes)
```

```
## 
##   Level 7:   78 nodes to be scored   (72 eliminated genes)
```

```
## 
##   Level 6:   99 nodes to be scored   (83 eliminated genes)
```

```
## 
##   Level 5:   85 nodes to be scored   (124 eliminated genes)
```

```
## 
##   Level 4:   50 nodes to be scored   (417 eliminated genes)
```

```
## 
##   Level 3:   29 nodes to be scored   (417 eliminated genes)
```

```
## 
##   Level 2:   7 nodes to be scored    (417 eliminated genes)
```

```
## 
##   Level 1:   1 nodes to be scored    (417 eliminated genes)
```

```
resultFisher_MF_up0v1h_elim <- runTest(GOdata_MF_up0v1h, algorithm = "elim", statistic = "fisher")
```

```
## 
##           -- Elim Algorithm -- 
## 
##       the algorithm is scoring 417 nontrivial nodes
##       parameters: 
##           test statistic: fisher
##           cutOff: 0.01
```

```
## 
##   Level 11:  1 nodes to be scored    (0 eliminated genes)
```

```
## 
##   Level 10:  4 nodes to be scored    (7 eliminated genes)
```

```
## 
##   Level 9:   17 nodes to be scored   (7 eliminated genes)
```

```
## 
##   Level 8:   33 nodes to be scored   (9 eliminated genes)
```

```
## 
##   Level 7:   54 nodes to be scored   (27 eliminated genes)
```

```
## 
##   Level 6:   95 nodes to be scored   (39 eliminated genes)
```

```
## 
##   Level 5:   91 nodes to be scored   (53 eliminated genes)
```

```
## 
##   Level 4:   78 nodes to be scored   (53 eliminated genes)
```

```
## 
##   Level 3:   31 nodes to be scored   (94 eliminated genes)
```

```
## 
##   Level 2:   12 nodes to be scored   (1276 eliminated genes)
```

```
## 
##   Level 1:   1 nodes to be scored    (1276 eliminated genes)
```

```
resultFisher_CC_up0v1h_elim <- runTest(GOdata_CC_up0v1h, algorithm = "elim", statistic = "fisher")
```

```
## 
##           -- Elim Algorithm -- 
## 
##       the algorithm is scoring 153 nontrivial nodes
##       parameters: 
##           test statistic: fisher
##           cutOff: 0.01
```

```
## 
##   Level 11:  2 nodes to be scored    (0 eliminated genes)
```

```
## 
##   Level 10:  7 nodes to be scored    (0 eliminated genes)
```

```
## 
##   Level 9:   12 nodes to be scored   (0 eliminated genes)
```

```
## 
##   Level 8:   18 nodes to be scored   (0 eliminated genes)
```

```
## 
##   Level 7:   29 nodes to be scored   (0 eliminated genes)
```

```
## 
##   Level 6:   30 nodes to be scored   (7 eliminated genes)
```

```
## 
##   Level 5:   19 nodes to be scored   (7 eliminated genes)
```

```
## 
##   Level 4:   18 nodes to be scored   (478 eliminated genes)
```

```
## 
##   Level 3:   15 nodes to be scored   (478 eliminated genes)
```

```
## 
##   Level 2:   2 nodes to be scored    (844 eliminated genes)
```

```
## 
##   Level 1:   1 nodes to be scored    (844 eliminated genes)
```

```
resultFisher_BP_down0v1h_elim <- runTest(GOdata_BP_down0v1h, algorithm = "elim", statistic = "fisher")
```

```
## 
##           -- Elim Algorithm -- 
## 
##       the algorithm is scoring 965 nontrivial nodes
##       parameters: 
##           test statistic: fisher
##           cutOff: 0.01
```

```
## 
##   Level 14:  4 nodes to be scored    (0 eliminated genes)
```

```
## 
##   Level 13:  11 nodes to be scored   (0 eliminated genes)
```

```
## 
##   Level 12:  37 nodes to be scored   (41 eliminated genes)
```

```
## 
##   Level 11:  54 nodes to be scored   (41 eliminated genes)
```

```
## 
##   Level 10:  95 nodes to be scored   (80 eliminated genes)
```

```
## 
##   Level 9:   125 nodes to be scored  (111 eliminated genes)
```

```
## 
##   Level 8:   138 nodes to be scored  (129 eliminated genes)
```

```
## 
##   Level 7:   140 nodes to be scored  (303 eliminated genes)
```

```
## 
##   Level 6:   134 nodes to be scored  (396 eliminated genes)
```

```
## 
##   Level 5:   118 nodes to be scored  (413 eliminated genes)
```

```
## 
##   Level 4:   61 nodes to be scored   (421 eliminated genes)
```

```
## 
##   Level 3:   40 nodes to be scored   (421 eliminated genes)
```

```
## 
##   Level 2:   7 nodes to be scored    (453 eliminated genes)
```

```
## 
##   Level 1:   1 nodes to be scored    (453 eliminated genes)
```

```
resultFisher_MF_down0v1h_elim <- runTest(GOdata_MF_down0v1h, algorithm = "elim", statistic = "fisher")
```

```
## 
##           -- Elim Algorithm -- 
## 
##       the algorithm is scoring 544 nontrivial nodes
##       parameters: 
##           test statistic: fisher
##           cutOff: 0.01
```

```
## 
##   Level 10:  4 nodes to be scored    (0 eliminated genes)
```

```
## 
##   Level 9:   17 nodes to be scored   (0 eliminated genes)
```

```
## 
##   Level 8:   32 nodes to be scored   (758 eliminated genes)
```

```
## 
##   Level 7:   82 nodes to be scored   (789 eliminated genes)
```

```
## 
##   Level 6:   151 nodes to be scored  (815 eliminated genes)
```

```
## 
##   Level 5:   124 nodes to be scored  (822 eliminated genes)
```

```
## 
##   Level 4:   87 nodes to be scored   (1016 eliminated genes)
```

```
## 
##   Level 3:   35 nodes to be scored   (1024 eliminated genes)
```

```
## 
##   Level 2:   11 nodes to be scored   (1112 eliminated genes)
```

```
## 
##   Level 1:   1 nodes to be scored    (1132 eliminated genes)
```

```
resultFisher_CC_down0v1h_elim <- runTest(GOdata_CC_down0v1h, algorithm = "elim", statistic = "fisher")
```

```
## 
##           -- Elim Algorithm -- 
## 
##       the algorithm is scoring 247 nontrivial nodes
##       parameters: 
##           test statistic: fisher
##           cutOff: 0.01
```

```
## 
##   Level 12:  2 nodes to be scored    (0 eliminated genes)
```

```
## 
##   Level 11:  10 nodes to be scored   (0 eliminated genes)
```

```
## 
##   Level 10:  13 nodes to be scored   (0 eliminated genes)
```

```
## 
##   Level 9:   26 nodes to be scored   (0 eliminated genes)
```

```
## 
##   Level 8:   38 nodes to be scored   (3 eliminated genes)
```

```
## 
##   Level 7:   38 nodes to be scored   (25 eliminated genes)
```

```
## 
##   Level 6:   37 nodes to be scored   (54 eliminated genes)
```

```
## 
##   Level 5:   28 nodes to be scored   (146 eliminated genes)
```

```
## 
##   Level 4:   29 nodes to be scored   (156 eliminated genes)
```

```
## 
##   Level 3:   23 nodes to be scored   (423 eliminated genes)
```

```
## 
##   Level 2:   2 nodes to be scored    (423 eliminated genes)
```

```
## 
##   Level 1:   1 nodes to be scored    (423 eliminated genes)
```

```
#extract the significant GO terms
up0v1h_BP_elim <- GenTable(GOdata_BP_up0v1h, classic = resultFisher_BP_up0v1h_elim, 
                            orderBy = "weight", ranksOf = "weight", topNodes = 50)
up0v1h_MF_elim <- GenTable(GOdata_MF_up0v1h, classic = resultFisher_MF_up0v1h_elim, 
                            orderBy = "weight", ranksOf = "weight", topNodes = 50)
up0v1h_CC_elim <- GenTable(GOdata_CC_up0v1h, classic = resultFisher_CC_up0v1h_elim, 
                            orderBy = "weight", ranksOf = "weight", topNodes = 50)
down0v1h_BP_elim <- GenTable(GOdata_BP_down0v1h, classic = resultFisher_BP_down0v1h_elim, 
                              orderBy = "weight", ranksOf = "weight", topNodes = 50)
down0v1h_MF_elim <- GenTable(GOdata_MF_down0v1h, classic = resultFisher_MF_down0v1h_elim, 
                              orderBy = "weight", ranksOf = "weight", topNodes = 50)
down0v1h_CC_elim <- GenTable(GOdata_CC_down0v1h, classic = resultFisher_CC_down0v1h_elim, 
                              orderBy = "weight", ranksOf = "weight", topNodes = 50)

#Write the tables to you desired location to run through revigo using code below as a template.
#write.table(up0v1h_BP_elim,
#            "/My Drive/ShortTermStress-IlluminaData/FullExp/Cryp-DE/data/ReviGo_files/BP_up1h_elim.txt")
#write.table(up0v1h_MF_elim,
#            "/My Drive/ShortTermStress-IlluminaData/FullExp/Cryp-DE/data/ReviGo_files/MF_up1h_elim.txt")
#write.table(up0v1h_CC_elim,
#            "/My Drive/ShortTermStress-IlluminaData/FullExp/Cryp-DE/data/ReviGo_files/CC_up1h_elim.txt")
#write.table(down0v1h_BP_elim,
#            "/My Drive/ShortTermStress-IlluminaData/FullExp/Cryp-DE/data/ReviGo_files/BP_dn1h_elim.txt")
#write.table(down0v1h_MF_elim,
#            "/My Drive/ShortTermStress-IlluminaData/FullExp/Cryp-DE/data/ReviGo_files/MF_dn1h_elim.txt")
#write.table(down0v1h_CC_elim,
#            "/My Drive/ShortTermStress-IlluminaData/FullExp/Cryp-DE/data/ReviGo_files/CC_dn1h_elim.txt")
```

2 hour timepoint

```
#Select only the genes significant at a timepoint from the reduced subset of 2 consecutive gene. This can be done for all time points, but is limited here to the ones discussed extensively in the manuscript
sign_0v2h <- rownames(keep_2_SignDEG)[keep_2_SignDEG[,"C0m-C60m"]==1] 

length(sign_0v2h) #3497 and this matches the output from SignifGenes
```

```
## [1] 3412
```

```
sign2LFC_0v2h <- sign2LFC [rownames(sign2LFC) %in% sign_0v2h, ] #3019 rows, which means one row had NAs and got omitted. 
logFCup_0v2h <- subset(sign2LFC_0v2h, (`0v2h_Padj`) <= 0.01 & (`0v2h`) > 0)
logFCdown_0v2h <- subset(sign2LFC_0v2h, (`0v2h_Padj`) <= 0.01 & (`0v2h`) < 0)

logFCup_0v2h_GOEnrich  = rownames(subset(logFCup_0v2h, rownames(logFCup_0v2h)%in%genesOfInterest_allSign))
logFCdown_0v2h_GOEnrich  = rownames(subset(logFCdown_0v2h, rownames(logFCdown_0v2h)%in%genesOfInterest_allSign))


#create gene list for input in topGO for 30 minutes
geneList_up0v2h = factor(as.integer(geneUniverse %in% logFCup_0v2h_GOEnrich))
names(geneList_up0v2h) = geneUniverse
str(geneList_up0v2h)
```

```
##  Factor w/ 2 levels "0","1": 1 1 1 1 1 1 1 1 1 1 ...
##  - attr(*, "names")= chr [1:6309] "gene_idGO.ID" "CCRYP_013698" "CCRYP_013706" "CCRYP_013704" ...
```

```
geneList_down0v2h = factor(as.integer(geneUniverse %in% logFCdown_0v2h_GOEnrich))
names(geneList_down0v2h) = geneUniverse
str(geneList_down0v2h)
```

```
##  Factor w/ 2 levels "0","1": 1 1 1 1 1 1 1 1 1 2 ...
##  - attr(*, "names")= chr [1:6309] "gene_idGO.ID" "CCRYP_013698" "CCRYP_013706" "CCRYP_013704" ...
```

```
##create a topGO object (for up GOs)
GOdata_BP_up0v2h = new("topGOdata", ontology="BP", allGenes=geneList_up0v2h, 
                        annot = annFUN.gene2GO, gene2GO = geneID2GO)
```

```
## 
## Building most specific GOs .....
```

```
##  ( 624 GO terms found. )
```

```
## 
## Build GO DAG topology ..........
```

```
##  ( 1729 GO terms and 3515 relations. )
```

```
## 
## Annotating nodes ...............
```

```
##  ( 3345 genes annotated to the GO terms. )
```

```
GOdata_MF_up0v2h = new("topGOdata", ontology="MF", allGenes=geneList_up0v2h, 
                        annot = annFUN.gene2GO, gene2GO = geneID2GO)
```

```
## 
## Building most specific GOs .....
```

```
##  ( 736 GO terms found. )
```

```
## 
## Build GO DAG topology ..........
```

```
##  ( 1096 GO terms and 1406 relations. )
```

```
## 
## Annotating nodes ...............
```

```
##  ( 5189 genes annotated to the GO terms. )
```

```
GOdata_CC_up0v2h = new("topGOdata", ontology="CC", allGenes=geneList_up0v2h, 
                        annot = annFUN.gene2GO, gene2GO = geneID2GO)
```

```
## 
## Building most specific GOs .....
```

```
##  ( 235 GO terms found. )
```

```
## 
## Build GO DAG topology ..........
```

```
##  ( 452 GO terms and 812 relations. )
```

```
## 
## Annotating nodes ...............
```

```
##  ( 1987 genes annotated to the GO terms. )
```

```
##create a topGO object (for down GOs)
GOdata_BP_down0v2h  = new("topGOdata", ontology="BP", allGenes=geneList_down0v2h , 
                           annot = annFUN.gene2GO, gene2GO = geneID2GO)
```

```
## 
## Building most specific GOs .....
```

```
##  ( 624 GO terms found. )
```

```
## 
## Build GO DAG topology ..........
```

```
##  ( 1729 GO terms and 3515 relations. )
```

```
## 
## Annotating nodes ...............
```

```
##  ( 3345 genes annotated to the GO terms. )
```

```
GOdata_MF_down0v2h  = new("topGOdata", ontology="MF", allGenes=geneList_down0v2h , 
                           annot = annFUN.gene2GO, gene2GO = geneID2GO)
```

```
## 
## Building most specific GOs .....
```

```
##  ( 736 GO terms found. )
```

```
## 
## Build GO DAG topology ..........
```

```
##  ( 1096 GO terms and 1406 relations. )
```

```
## 
## Annotating nodes ...............
```

```
##  ( 5189 genes annotated to the GO terms. )
```

```
GOdata_CC_down0v2h  = new("topGOdata", ontology="CC", allGenes=geneList_down0v2h , 
                           annot = annFUN.gene2GO, gene2GO = geneID2GO)
```

```
## 
## Building most specific GOs .....
```

```
##  ( 235 GO terms found. )
```

```
## 
## Build GO DAG topology ..........
```

```
##  ( 452 GO terms and 812 relations. )
```

```
## 
## Annotating nodes ...............
```

```
##  ( 1987 genes annotated to the GO terms. )
```

```
#run Fisher's exact test
resultFisher_BP_up0v2h_elim <- runTest(GOdata_BP_up0v2h, algorithm = "elim", statistic = "fisher")
```

```
## 
##           -- Elim Algorithm -- 
## 
##       the algorithm is scoring 371 nontrivial nodes
##       parameters: 
##           test statistic: fisher
##           cutOff: 0.01
```

```
## 
##   Level 13:  3 nodes to be scored    (0 eliminated genes)
```

```
## 
##   Level 12:  5 nodes to be scored    (0 eliminated genes)
```

```
## 
##   Level 11:  14 nodes to be scored   (0 eliminated genes)
```

```
## 
##   Level 10:  24 nodes to be scored   (0 eliminated genes)
```

```
## 
##   Level 9:   36 nodes to be scored   (0 eliminated genes)
```

```
## 
##   Level 8:   42 nodes to be scored   (3 eliminated genes)
```

```
## 
##   Level 7:   49 nodes to be scored   (3 eliminated genes)
```

```
## 
##   Level 6:   66 nodes to be scored   (3 eliminated genes)
```

```
## 
##   Level 5:   63 nodes to be scored   (44 eliminated genes)
```

```
## 
##   Level 4:   37 nodes to be scored   (55 eliminated genes)
```

```
## 
##   Level 3:   25 nodes to be scored   (55 eliminated genes)
```

```
## 
##   Level 2:   6 nodes to be scored    (55 eliminated genes)
```

```
## 
##   Level 1:   1 nodes to be scored    (55 eliminated genes)
```

```
resultFisher_MF_up0v2h_elim <- runTest(GOdata_MF_up0v2h, algorithm = "elim", statistic = "fisher")
```

```
## 
##           -- Elim Algorithm -- 
## 
##       the algorithm is scoring 297 nontrivial nodes
##       parameters: 
##           test statistic: fisher
##           cutOff: 0.01
```

```
## 
##   Level 11:  1 nodes to be scored    (0 eliminated genes)
```

```
## 
##   Level 10:  3 nodes to be scored    (0 eliminated genes)
```

```
## 
##   Level 9:   12 nodes to be scored   (0 eliminated genes)
```

```
## 
##   Level 8:   24 nodes to be scored   (3 eliminated genes)
```

```
## 
##   Level 7:   36 nodes to be scored   (12 eliminated genes)
```

```
## 
##   Level 6:   61 nodes to be scored   (15 eliminated genes)
```

```
## 
##   Level 5:   61 nodes to be scored   (22 eliminated genes)
```

```
## 
##   Level 4:   59 nodes to be scored   (22 eliminated genes)
```

```
## 
##   Level 3:   29 nodes to be scored   (24 eliminated genes)
```

```
## 
##   Level 2:   10 nodes to be scored   (24 eliminated genes)
```

```
## 
##   Level 1:   1 nodes to be scored    (55 eliminated genes)
```

```
resultFisher_CC_up0v2h_elim <- runTest(GOdata_CC_up0v2h, algorithm = "elim", statistic = "fisher")
```

```
## 
##           -- Elim Algorithm -- 
## 
##       the algorithm is scoring 100 nontrivial nodes
##       parameters: 
##           test statistic: fisher
##           cutOff: 0.01
```

```
## 
##   Level 11:  1 nodes to be scored    (0 eliminated genes)
```

```
## 
##   Level 10:  4 nodes to be scored    (0 eliminated genes)
```

```
## 
##   Level 9:   8 nodes to be scored    (0 eliminated genes)
```

```
## 
##   Level 8:   11 nodes to be scored   (0 eliminated genes)
```

```
## 
##   Level 7:   16 nodes to be scored   (0 eliminated genes)
```

```
## 
##   Level 6:   17 nodes to be scored   (7 eliminated genes)
```

```
## 
##   Level 5:   13 nodes to be scored   (7 eliminated genes)
```

```
## 
##   Level 4:   14 nodes to be scored   (7 eliminated genes)
```

```
## 
##   Level 3:   13 nodes to be scored   (7 eliminated genes)
```

```
## 
##   Level 2:   2 nodes to be scored    (844 eliminated genes)
```

```
## 
##   Level 1:   1 nodes to be scored    (844 eliminated genes)
```

```
resultFisher_BP_down0v2h_elim <- runTest(GOdata_BP_down0v2h, algorithm = "elim", statistic = "fisher")
```

```
## 
##           -- Elim Algorithm -- 
## 
##       the algorithm is scoring 719 nontrivial nodes
##       parameters: 
##           test statistic: fisher
##           cutOff: 0.01
```

```
## 
##   Level 14:  3 nodes to be scored    (0 eliminated genes)
```

```
## 
##   Level 13:  6 nodes to be scored    (0 eliminated genes)
```

```
## 
##   Level 12:  21 nodes to be scored   (0 eliminated genes)
```

```
## 
##   Level 11:  39 nodes to be scored   (0 eliminated genes)
```

```
## 
##   Level 10:  73 nodes to be scored   (0 eliminated genes)
```

```
## 
##   Level 9:   89 nodes to be scored   (65 eliminated genes)
```

```
## 
##   Level 8:   94 nodes to be scored   (65 eliminated genes)
```

```
## 
##   Level 7:   97 nodes to be scored   (243 eliminated genes)
```

```
## 
##   Level 6:   105 nodes to be scored  (243 eliminated genes)
```

```
## 
##   Level 5:   100 nodes to be scored  (275 eliminated genes)
```

```
## 
##   Level 4:   52 nodes to be scored   (275 eliminated genes)
```

```
## 
##   Level 3:   33 nodes to be scored   (275 eliminated genes)
```

```
## 
##   Level 2:   6 nodes to be scored    (315 eliminated genes)
```

```
## 
##   Level 1:   1 nodes to be scored    (315 eliminated genes)
```

```
resultFisher_MF_down0v2h_elim <- runTest(GOdata_MF_down0v2h, algorithm = "elim", statistic = "fisher")
```

```
## 
##           -- Elim Algorithm -- 
## 
##       the algorithm is scoring 437 nontrivial nodes
##       parameters: 
##           test statistic: fisher
##           cutOff: 0.01
```

```
## 
##   Level 10:  1 nodes to be scored    (0 eliminated genes)
```

```
## 
##   Level 9:   8 nodes to be scored    (0 eliminated genes)
```

```
## 
##   Level 8:   21 nodes to be scored   (0 eliminated genes)
```

```
## 
##   Level 7:   61 nodes to be scored   (0 eliminated genes)
```

```
## 
##   Level 6:   114 nodes to be scored  (789 eliminated genes)
```

```
## 
##   Level 5:   109 nodes to be scored  (796 eliminated genes)
```

```
## 
##   Level 4:   79 nodes to be scored   (986 eliminated genes)
```

```
## 
##   Level 3:   32 nodes to be scored   (1129 eliminated genes)
```

```
## 
##   Level 2:   11 nodes to be scored   (1231 eliminated genes)
```

```
## 
##   Level 1:   1 nodes to be scored    (1231 eliminated genes)
```

```
resultFisher_CC_down0v2h_elim <- runTest(GOdata_CC_down0v2h, algorithm = "elim", statistic = "fisher")
```

```
## 
##           -- Elim Algorithm -- 
## 
##       the algorithm is scoring 192 nontrivial nodes
##       parameters: 
##           test statistic: fisher
##           cutOff: 0.01
```

```
## 
##   Level 12:  2 nodes to be scored    (0 eliminated genes)
```

```
## 
##   Level 11:  7 nodes to be scored    (0 eliminated genes)
```

```
## 
##   Level 10:  10 nodes to be scored   (0 eliminated genes)
```

```
## 
##   Level 9:   20 nodes to be scored   (0 eliminated genes)
```

```
## 
##   Level 8:   29 nodes to be scored   (0 eliminated genes)
```

```
## 
##   Level 7:   25 nodes to be scored   (14 eliminated genes)
```

```
## 
##   Level 6:   28 nodes to be scored   (47 eliminated genes)
```

```
## 
##   Level 5:   26 nodes to be scored   (140 eliminated genes)
```

```
## 
##   Level 4:   22 nodes to be scored   (152 eliminated genes)
```

```
## 
##   Level 3:   20 nodes to be scored   (152 eliminated genes)
```

```
## 
##   Level 2:   2 nodes to be scored    (193 eliminated genes)
```

```
## 
##   Level 1:   1 nodes to be scored    (193 eliminated genes)
```

```
#extract the significant GO terms
up0v2h_BP_elim <- GenTable(GOdata_BP_up0v2h, classic = resultFisher_BP_up0v2h_elim, 
                            orderBy = "weight", ranksOf = "weight", topNodes = 50)
up0v2h_MF_elim <- GenTable(GOdata_MF_up0v2h, classic = resultFisher_MF_up0v2h_elim, 
                            orderBy = "weight", ranksOf = "weight", topNodes = 50)
up0v2h_CC_elim <- GenTable(GOdata_CC_up0v2h, classic = resultFisher_CC_up0v2h_elim, 
                            orderBy = "weight", ranksOf = "weight", topNodes = 50)
down0v2h_BP_elim <- GenTable(GOdata_BP_down0v2h, classic = resultFisher_BP_down0v2h_elim, 
                              orderBy = "weight", ranksOf = "weight", topNodes = 50)
down0v2h_MF_elim <- GenTable(GOdata_MF_down0v2h, classic = resultFisher_MF_down0v2h_elim, 
                              orderBy = "weight", ranksOf = "weight", topNodes = 50)
down0v2h_CC_elim <- GenTable(GOdata_CC_down0v2h, classic = resultFisher_CC_down0v2h_elim, 
                              orderBy = "weight", ranksOf = "weight", topNodes = 50)

#Write the tables to you desired location to run through revigo using code below as a template.
#write.table(up0v2h_BP_elim,
#            "/My Drive/ShortTermStress-IlluminaData/FullExp/Cryp-DE/data/ReviGo_files/BP_up2h_elim.txt")
#write.table(up0v2h_MF_elim,
#            "/My Drive/ShortTermStress-IlluminaData/FullExp/Cryp-DE/data/ReviGo_files/MF_up2h_elim.txt")
#write.table(up0v2h_CC_elim,
#            "/My Drive/ShortTermStress-IlluminaData/FullExp/Cryp-DE/data/ReviGo_files/CC_up2h_elim.txt")
#write.table(down0v2h_BP_elim,
#            "/My Drive/ShortTermStress-IlluminaData/FullExp/Cryp-DE/data/ReviGo_files/BP_dn2h_elim.txt")
#write.table(down0v2h_MF_elim,
#            "/My Drive/ShortTermStress-IlluminaData/FullExp/Cryp-DE/data/ReviGo_files/MF_dn2h_elim.txt")
#write.table(down0v2h_CC_elim,
#            "/My Drive/ShortTermStress-IlluminaData/FullExp/Cryp-DE/data/ReviGo_files/CC_dn2h_elim.txt")
```

4 hour timepoint

```
#Select only the genes significant at a timepoint from the reduced subset of 2 consecutive gene. This can be done for all time points, but is limited here to the ones discussed extensively in the manuscript
sign_0v4h <- rownames(keep_2_SignDEG)[keep_2_SignDEG[,"C0m-C60m"]==1] 

length(sign_0v4h) #3497 and this matches the output from SignifGenes
```

```
## [1] 3412
```

```
sign2LFC_0v4h <- sign2LFC [rownames(sign2LFC) %in% sign_0v4h, ] #3019 rows, which means one row had NAs and got omitted. 
logFCup_0v4h <- subset(sign2LFC_0v4h, (`0v4h_Padj`) <= 0.01 & (`0v4h`) > 0)
logFCdown_0v4h <- subset(sign2LFC_0v4h, (`0v4h_Padj`) <= 0.01 & (`0v4h`) < 0)

logFCup_0v4h_GOEnrich  = rownames(subset(logFCup_0v4h, rownames(logFCup_0v4h)%in%genesOfInterest_allSign))
logFCdown_0v4h_GOEnrich  = rownames(subset(logFCdown_0v4h, rownames(logFCdown_0v4h)%in%genesOfInterest_allSign))


#create gene list for input in topGO for 30 minutes
geneList_up0v4h = factor(as.integer(geneUniverse %in% logFCup_0v4h_GOEnrich))
names(geneList_up0v4h) = geneUniverse
str(geneList_up0v4h)
```

```
##  Factor w/ 2 levels "0","1": 1 1 1 1 1 1 1 1 1 1 ...
##  - attr(*, "names")= chr [1:6309] "gene_idGO.ID" "CCRYP_013698" "CCRYP_013706" "CCRYP_013704" ...
```

```
geneList_down0v4h = factor(as.integer(geneUniverse %in% logFCdown_0v4h_GOEnrich))
names(geneList_down0v4h) = geneUniverse
str(geneList_down0v4h)
```

```
##  Factor w/ 2 levels "0","1": 1 1 1 1 1 1 1 1 1 1 ...
##  - attr(*, "names")= chr [1:6309] "gene_idGO.ID" "CCRYP_013698" "CCRYP_013706" "CCRYP_013704" ...
```

```
##create a topGO object (for up GOs)
GOdata_BP_up0v4h = new("topGOdata", ontology="BP", allGenes=geneList_up0v4h, 
                        annot = annFUN.gene2GO, gene2GO = geneID2GO)
```

```
## 
## Building most specific GOs .....
```

```
##  ( 624 GO terms found. )
```

```
## 
## Build GO DAG topology ..........
```

```
##  ( 1729 GO terms and 3515 relations. )
```

```
## 
## Annotating nodes ...............
```

```
##  ( 3345 genes annotated to the GO terms. )
```

```
GOdata_MF_up0v4h = new("topGOdata", ontology="MF", allGenes=geneList_up0v4h, 
                        annot = annFUN.gene2GO, gene2GO = geneID2GO)
```

```
## 
## Building most specific GOs .....
```

```
##  ( 736 GO terms found. )
```

```
## 
## Build GO DAG topology ..........
```

```
##  ( 1096 GO terms and 1406 relations. )
```

```
## 
## Annotating nodes ...............
```

```
##  ( 5189 genes annotated to the GO terms. )
```

```
GOdata_CC_up0v4h = new("topGOdata", ontology="CC", allGenes=geneList_up0v4h, 
                        annot = annFUN.gene2GO, gene2GO = geneID2GO)
```

```
## 
## Building most specific GOs .....
```

```
##  ( 235 GO terms found. )
```

```
## 
## Build GO DAG topology ..........
```

```
##  ( 452 GO terms and 812 relations. )
```

```
## 
## Annotating nodes ...............
```

```
##  ( 1987 genes annotated to the GO terms. )
```

```
##create a topGO object (for down GOs)
GOdata_BP_down0v4h  = new("topGOdata", ontology="BP", allGenes=geneList_down0v4h , 
                           annot = annFUN.gene2GO, gene2GO = geneID2GO)
```

```
## 
## Building most specific GOs .....
```

```
##  ( 624 GO terms found. )
```

```
## 
## Build GO DAG topology ..........
```

```
##  ( 1729 GO terms and 3515 relations. )
```

```
## 
## Annotating nodes ...............
```

```
##  ( 3345 genes annotated to the GO terms. )
```

```
GOdata_MF_down0v4h  = new("topGOdata", ontology="MF", allGenes=geneList_down0v4h , 
                           annot = annFUN.gene2GO, gene2GO = geneID2GO)
```

```
## 
## Building most specific GOs .....
```

```
##  ( 736 GO terms found. )
```

```
## 
## Build GO DAG topology ..........
```

```
##  ( 1096 GO terms and 1406 relations. )
```

```
## 
## Annotating nodes ...............
```

```
##  ( 5189 genes annotated to the GO terms. )
```

```
GOdata_CC_down0v4h  = new("topGOdata", ontology="CC", allGenes=geneList_down0v4h , 
                           annot = annFUN.gene2GO, gene2GO = geneID2GO)
```

```
## 
## Building most specific GOs .....
```

```
##  ( 235 GO terms found. )
```

```
## 
## Build GO DAG topology ..........
```

```
##  ( 452 GO terms and 812 relations. )
```

```
## 
## Annotating nodes ...............
```

```
##  ( 1987 genes annotated to the GO terms. )
```

```
#run Fisher's exact test
resultFisher_BP_up0v4h_elim <- runTest(GOdata_BP_up0v4h, algorithm = "elim", statistic = "fisher")
```

```
## 
##           -- Elim Algorithm -- 
## 
##       the algorithm is scoring 404 nontrivial nodes
##       parameters: 
##           test statistic: fisher
##           cutOff: 0.01
```

```
## 
##   Level 13:  3 nodes to be scored    (0 eliminated genes)
```

```
## 
##   Level 12:  4 nodes to be scored    (0 eliminated genes)
```

```
## 
##   Level 11:  11 nodes to be scored   (0 eliminated genes)
```

```
## 
##   Level 10:  27 nodes to be scored   (0 eliminated genes)
```

```
## 
##   Level 9:   42 nodes to be scored   (0 eliminated genes)
```

```
## 
##   Level 8:   44 nodes to be scored   (0 eliminated genes)
```

```
## 
##   Level 7:   57 nodes to be scored   (0 eliminated genes)
```

```
## 
##   Level 6:   67 nodes to be scored   (3 eliminated genes)
```

```
## 
##   Level 5:   73 nodes to be scored   (92 eliminated genes)
```

```
## 
##   Level 4:   42 nodes to be scored   (92 eliminated genes)
```

```
## 
##   Level 3:   27 nodes to be scored   (92 eliminated genes)
```

```
## 
##   Level 2:   6 nodes to be scored    (92 eliminated genes)
```

```
## 
##   Level 1:   1 nodes to be scored    (92 eliminated genes)
```

```
resultFisher_MF_up0v4h_elim <- runTest(GOdata_MF_up0v4h, algorithm = "elim", statistic = "fisher")
```

```
## 
##           -- Elim Algorithm -- 
## 
##       the algorithm is scoring 252 nontrivial nodes
##       parameters: 
##           test statistic: fisher
##           cutOff: 0.01
```

```
## 
##   Level 11:  1 nodes to be scored    (0 eliminated genes)
```

```
## 
##   Level 10:  1 nodes to be scored    (0 eliminated genes)
```

```
## 
##   Level 9:   7 nodes to be scored    (0 eliminated genes)
```

```
## 
##   Level 8:   14 nodes to be scored   (644 eliminated genes)
```

```
## 
##   Level 7:   28 nodes to be scored   (644 eliminated genes)
```

```
## 
##   Level 6:   52 nodes to be scored   (648 eliminated genes)
```

```
## 
##   Level 5:   58 nodes to be scored   (650 eliminated genes)
```

```
## 
##   Level 4:   55 nodes to be scored   (650 eliminated genes)
```

```
## 
##   Level 3:   25 nodes to be scored   (942 eliminated genes)
```

```
## 
##   Level 2:   10 nodes to be scored   (942 eliminated genes)
```

```
## 
##   Level 1:   1 nodes to be scored    (942 eliminated genes)
```

```
resultFisher_CC_up0v4h_elim <- runTest(GOdata_CC_up0v4h, algorithm = "elim", statistic = "fisher")
```

```
## 
##           -- Elim Algorithm -- 
## 
##       the algorithm is scoring 68 nontrivial nodes
##       parameters: 
##           test statistic: fisher
##           cutOff: 0.01
```

```
## 
##   Level 9:   3 nodes to be scored    (0 eliminated genes)
```

```
## 
##   Level 8:   4 nodes to be scored    (0 eliminated genes)
```

```
## 
##   Level 7:   8 nodes to be scored    (14 eliminated genes)
```

```
## 
##   Level 6:   12 nodes to be scored   (14 eliminated genes)
```

```
## 
##   Level 5:   12 nodes to be scored   (14 eliminated genes)
```

```
## 
##   Level 4:   14 nodes to be scored   (14 eliminated genes)
```

```
## 
##   Level 3:   12 nodes to be scored   (14 eliminated genes)
```

```
## 
##   Level 2:   2 nodes to be scored    (852 eliminated genes)
```

```
## 
##   Level 1:   1 nodes to be scored    (852 eliminated genes)
```

```
resultFisher_BP_down0v4h_elim <- runTest(GOdata_BP_down0v4h, algorithm = "elim", statistic = "fisher")
```

```
## 
##           -- Elim Algorithm -- 
## 
##       the algorithm is scoring 421 nontrivial nodes
##       parameters: 
##           test statistic: fisher
##           cutOff: 0.01
```

```
## 
##   Level 14:  1 nodes to be scored    (0 eliminated genes)
```

```
## 
##   Level 13:  2 nodes to be scored    (0 eliminated genes)
```

```
## 
##   Level 12:  4 nodes to be scored    (0 eliminated genes)
```

```
## 
##   Level 11:  10 nodes to be scored   (0 eliminated genes)
```

```
## 
##   Level 10:  25 nodes to be scored   (0 eliminated genes)
```

```
## 
##   Level 9:   35 nodes to be scored   (0 eliminated genes)
```

```
## 
##   Level 8:   50 nodes to be scored   (0 eliminated genes)
```

```
## 
##   Level 7:   60 nodes to be scored   (0 eliminated genes)
```

```
## 
##   Level 6:   77 nodes to be scored   (32 eliminated genes)
```

```
## 
##   Level 5:   78 nodes to be scored   (32 eliminated genes)
```

```
## 
##   Level 4:   44 nodes to be scored   (32 eliminated genes)
```

```
## 
##   Level 3:   28 nodes to be scored   (32 eliminated genes)
```

```
## 
##   Level 2:   6 nodes to be scored    (32 eliminated genes)
```

```
## 
##   Level 1:   1 nodes to be scored    (32 eliminated genes)
```

```
resultFisher_MF_down0v4h_elim <- runTest(GOdata_MF_down0v4h, algorithm = "elim", statistic = "fisher")
```

```
## 
##           -- Elim Algorithm -- 
## 
##       the algorithm is scoring 247 nontrivial nodes
##       parameters: 
##           test statistic: fisher
##           cutOff: 0.01
```

```
## 
##   Level 11:  1 nodes to be scored    (0 eliminated genes)
```

```
## 
##   Level 10:  2 nodes to be scored    (0 eliminated genes)
```

```
## 
##   Level 9:   8 nodes to be scored    (0 eliminated genes)
```

```
## 
##   Level 8:   15 nodes to be scored   (0 eliminated genes)
```

```
## 
##   Level 7:   28 nodes to be scored   (0 eliminated genes)
```

```
## 
##   Level 6:   51 nodes to be scored   (91 eliminated genes)
```

```
## 
##   Level 5:   55 nodes to be scored   (91 eliminated genes)
```

```
## 
##   Level 4:   52 nodes to be scored   (242 eliminated genes)
```

```
## 
##   Level 3:   24 nodes to be scored   (358 eliminated genes)
```

```
## 
##   Level 2:   10 nodes to be scored   (362 eliminated genes)
```

```
## 
##   Level 1:   1 nodes to be scored    (362 eliminated genes)
```

```
resultFisher_CC_down0v4h_elim <- runTest(GOdata_CC_down0v4h, algorithm = "elim", statistic = "fisher")
```

```
## 
##           -- Elim Algorithm -- 
## 
##       the algorithm is scoring 95 nontrivial nodes
##       parameters: 
##           test statistic: fisher
##           cutOff: 0.01
```

```
## 
##   Level 12:  1 nodes to be scored    (0 eliminated genes)
```

```
## 
##   Level 11:  2 nodes to be scored    (0 eliminated genes)
```

```
## 
##   Level 10:  5 nodes to be scored    (0 eliminated genes)
```

```
## 
##   Level 9:   8 nodes to be scored    (0 eliminated genes)
```

```
## 
##   Level 8:   10 nodes to be scored   (0 eliminated genes)
```

```
## 
##   Level 7:   10 nodes to be scored   (0 eliminated genes)
```

```
## 
##   Level 6:   15 nodes to be scored   (0 eliminated genes)
```

```
## 
##   Level 5:   15 nodes to be scored   (0 eliminated genes)
```

```
## 
##   Level 4:   15 nodes to be scored   (0 eliminated genes)
```

```
## 
##   Level 3:   11 nodes to be scored   (0 eliminated genes)
```

```
## 
##   Level 2:   2 nodes to be scored    (41 eliminated genes)
```

```
## 
##   Level 1:   1 nodes to be scored    (41 eliminated genes)
```

```
#extract the significant GO terms
up0v4h_BP_elim <- GenTable(GOdata_BP_up0v4h, classic = resultFisher_BP_up0v4h_elim, 
                            orderBy = "weight", ranksOf = "weight", topNodes = 50)
up0v4h_MF_elim <- GenTable(GOdata_MF_up0v4h, classic = resultFisher_MF_up0v4h_elim, 
                            orderBy = "weight", ranksOf = "weight", topNodes = 50)
up0v4h_CC_elim <- GenTable(GOdata_CC_up0v4h, classic = resultFisher_CC_up0v4h_elim, 
                            orderBy = "weight", ranksOf = "weight", topNodes = 50)
down0v4h_BP_elim <- GenTable(GOdata_BP_down0v4h, classic = resultFisher_BP_down0v4h_elim, 
                              orderBy = "weight", ranksOf = "weight", topNodes = 50)
down0v4h_MF_elim <- GenTable(GOdata_MF_down0v4h, classic = resultFisher_MF_down0v4h_elim, 
                              orderBy = "weight", ranksOf = "weight", topNodes = 50)
down0v4h_CC_elim <- GenTable(GOdata_CC_down0v4h, classic = resultFisher_CC_down0v4h_elim, 
                              orderBy = "weight", ranksOf = "weight", topNodes = 50)

#Write the tables to you desired location to run through revigo using code below as a template.
#write.table(up0v4h_BP_elim,
#            "/My Drive/ShortTermStress-IlluminaData/FullExp/Cryp-DE/data/ReviGo_files/BP_up4h_elim.txt")
#write.table(up0v4h_MF_elim,
#            "/My Drive/ShortTermStress-IlluminaData/FullExp/Cryp-DE/data/ReviGo_files/MF_up4h_elim.txt")
#write.table(up0v4h_CC_elim,
#            "/My Drive/ShortTermStress-IlluminaData/FullExp/Cryp-DE/data/ReviGo_files/CC_up4h_elim.txt")
#write.table(down0v4h_BP_elim,
#            "/My Drive/ShortTermStress-IlluminaData/FullExp/Cryp-DE/data/ReviGo_files/BP_dn4h_elim.txt")
#write.table(down0v4h_MF_elim,
#            "/My Drive/ShortTermStress-IlluminaData/FullExp/Cryp-DE/data/ReviGo_files/MF_dn4h_elim.txt")
#write.table(down0v4h_CC_elim,
#            "/My Drive/ShortTermStress-IlluminaData/FullExp/Cryp-DE/data/ReviGo_files/CC_dn4h_elim.txt")
```

8 hour timepoint

```
#Select only the genes significant at a timepoint from the reduced subset of 2 consecutive gene. This can be done for all time points, but is limited here to the ones discussed extensively in the manuscript
sign_0v8h <- rownames(keep_2_SignDEG)[keep_2_SignDEG[,"C0m-C60m"]==1] 

length(sign_0v8h) #3497 and this matches the output from SignifGenes
```

```
## [1] 3412
```

```
sign2LFC_0v8h <- sign2LFC [rownames(sign2LFC) %in% sign_0v8h, ] #3019 rows, which means one row had NAs and got omitted. 
logFCup_0v8h <- subset(sign2LFC_0v8h, (`0v8h_Padj`) <= 0.01 & (`0v8h`) > 0)
logFCdown_0v8h <- subset(sign2LFC_0v8h, (`0v8h_Padj`) <= 0.01 & (`0v8h`) < 0)

logFCup_0v8h_GOEnrich  = rownames(subset(logFCup_0v8h, rownames(logFCup_0v8h)%in%genesOfInterest_allSign))
logFCdown_0v8h_GOEnrich  = rownames(subset(logFCdown_0v8h, rownames(logFCdown_0v8h)%in%genesOfInterest_allSign))


#create gene list for input in topGO for 30 minutes
geneList_up0v8h = factor(as.integer(geneUniverse %in% logFCup_0v8h_GOEnrich))
names(geneList_up0v8h) = geneUniverse
str(geneList_up0v8h)
```

```
##  Factor w/ 2 levels "0","1": 1 1 1 1 2 1 1 1 1 1 ...
##  - attr(*, "names")= chr [1:6309] "gene_idGO.ID" "CCRYP_013698" "CCRYP_013706" "CCRYP_013704" ...
```

```
geneList_down0v8h = factor(as.integer(geneUniverse %in% logFCdown_0v8h_GOEnrich))
names(geneList_down0v8h) = geneUniverse
str(geneList_down0v8h)
```

```
##  Factor w/ 2 levels "0","1": 1 1 1 1 1 1 1 2 1 1 ...
##  - attr(*, "names")= chr [1:6309] "gene_idGO.ID" "CCRYP_013698" "CCRYP_013706" "CCRYP_013704" ...
```

```
##create a topGO object (for up GOs)
GOdata_BP_up0v8h = new("topGOdata", ontology="BP", allGenes=geneList_up0v8h, 
                        annot = annFUN.gene2GO, gene2GO = geneID2GO)
```

```
## 
## Building most specific GOs .....
```

```
##  ( 624 GO terms found. )
```

```
## 
## Build GO DAG topology ..........
```

```
##  ( 1729 GO terms and 3515 relations. )
```

```
## 
## Annotating nodes ...............
```

```
##  ( 3345 genes annotated to the GO terms. )
```

```
GOdata_MF_up0v8h = new("topGOdata", ontology="MF", allGenes=geneList_up0v8h, 
                        annot = annFUN.gene2GO, gene2GO = geneID2GO)
```

```
## 
## Building most specific GOs .....
```

```
##  ( 736 GO terms found. )
```

```
## 
## Build GO DAG topology ..........
```

```
##  ( 1096 GO terms and 1406 relations. )
```

```
## 
## Annotating nodes ...............
```

```
##  ( 5189 genes annotated to the GO terms. )
```

```
GOdata_CC_up0v8h = new("topGOdata", ontology="CC", allGenes=geneList_up0v8h, 
                        annot = annFUN.gene2GO, gene2GO = geneID2GO)
```

```
## 
## Building most specific GOs .....
```

```
##  ( 235 GO terms found. )
```

```
## 
## Build GO DAG topology ..........
```

```
##  ( 452 GO terms and 812 relations. )
```

```
## 
## Annotating nodes ...............
```

```
##  ( 1987 genes annotated to the GO terms. )
```

```
##create a topGO object (for down GOs)
GOdata_BP_down0v8h  = new("topGOdata", ontology="BP", allGenes=geneList_down0v8h , 
                           annot = annFUN.gene2GO, gene2GO = geneID2GO)
```

```
## 
## Building most specific GOs .....
```

```
##  ( 624 GO terms found. )
```

```
## 
## Build GO DAG topology ..........
```

```
##  ( 1729 GO terms and 3515 relations. )
```

```
## 
## Annotating nodes ...............
```

```
##  ( 3345 genes annotated to the GO terms. )
```

```
GOdata_MF_down0v8h  = new("topGOdata", ontology="MF", allGenes=geneList_down0v8h , 
                           annot = annFUN.gene2GO, gene2GO = geneID2GO)
```

```
## 
## Building most specific GOs .....
```

```
##  ( 736 GO terms found. )
```

```
## 
## Build GO DAG topology ..........
```

```
##  ( 1096 GO terms and 1406 relations. )
```

```
## 
## Annotating nodes ...............
```

```
##  ( 5189 genes annotated to the GO terms. )
```

```
GOdata_CC_down0v8h  = new("topGOdata", ontology="CC", allGenes=geneList_down0v8h , 
                           annot = annFUN.gene2GO, gene2GO = geneID2GO)
```

```
## 
## Building most specific GOs .....
```

```
##  ( 235 GO terms found. )
```

```
## 
## Build GO DAG topology ..........
```

```
##  ( 452 GO terms and 812 relations. )
```

```
## 
## Annotating nodes ...............
```

```
##  ( 1987 genes annotated to the GO terms. )
```

```
#run Fisher's exact test
resultFisher_BP_up0v8h_elim <- runTest(GOdata_BP_up0v8h, algorithm = "elim", statistic = "fisher")
```

```
## 
##           -- Elim Algorithm -- 
## 
##       the algorithm is scoring 436 nontrivial nodes
##       parameters: 
##           test statistic: fisher
##           cutOff: 0.01
```

```
## 
##   Level 13:  3 nodes to be scored    (0 eliminated genes)
```

```
## 
##   Level 12:  7 nodes to be scored    (0 eliminated genes)
```

```
## 
##   Level 11:  12 nodes to be scored   (0 eliminated genes)
```

```
## 
##   Level 10:  31 nodes to be scored   (0 eliminated genes)
```

```
## 
##   Level 9:   54 nodes to be scored   (31 eliminated genes)
```

```
## 
##   Level 8:   53 nodes to be scored   (31 eliminated genes)
```

```
## 
##   Level 7:   58 nodes to be scored   (210 eliminated genes)
```

```
## 
##   Level 6:   65 nodes to be scored   (224 eliminated genes)
```

```
## 
##   Level 5:   77 nodes to be scored   (240 eliminated genes)
```

```
## 
##   Level 4:   43 nodes to be scored   (240 eliminated genes)
```

```
## 
##   Level 3:   26 nodes to be scored   (240 eliminated genes)
```

```
## 
##   Level 2:   6 nodes to be scored    (240 eliminated genes)
```

```
## 
##   Level 1:   1 nodes to be scored    (240 eliminated genes)
```

```
resultFisher_MF_up0v8h_elim <- runTest(GOdata_MF_up0v8h, algorithm = "elim", statistic = "fisher")
```

```
## 
##           -- Elim Algorithm -- 
## 
##       the algorithm is scoring 251 nontrivial nodes
##       parameters: 
##           test statistic: fisher
##           cutOff: 0.01
```

```
## 
##   Level 9:   4 nodes to be scored    (0 eliminated genes)
```

```
## 
##   Level 8:   14 nodes to be scored   (647 eliminated genes)
```

```
## 
##   Level 7:   35 nodes to be scored   (647 eliminated genes)
```

```
## 
##   Level 6:   55 nodes to be scored   (656 eliminated genes)
```

```
## 
##   Level 5:   57 nodes to be scored   (663 eliminated genes)
```

```
## 
##   Level 4:   52 nodes to be scored   (663 eliminated genes)
```

```
## 
##   Level 3:   24 nodes to be scored   (670 eliminated genes)
```

```
## 
##   Level 2:   9 nodes to be scored    (764 eliminated genes)
```

```
## 
##   Level 1:   1 nodes to be scored    (3076 eliminated genes)
```

```
resultFisher_CC_up0v8h_elim <- runTest(GOdata_CC_up0v8h, algorithm = "elim", statistic = "fisher")
```

```
## 
##           -- Elim Algorithm -- 
## 
##       the algorithm is scoring 115 nontrivial nodes
##       parameters: 
##           test statistic: fisher
##           cutOff: 0.01
```

```
## 
##   Level 12:  1 nodes to be scored    (0 eliminated genes)
```

```
## 
##   Level 11:  2 nodes to be scored    (0 eliminated genes)
```

```
## 
##   Level 10:  5 nodes to be scored    (0 eliminated genes)
```

```
## 
##   Level 9:   10 nodes to be scored   (0 eliminated genes)
```

```
## 
##   Level 8:   16 nodes to be scored   (0 eliminated genes)
```

```
## 
##   Level 7:   12 nodes to be scored   (17 eliminated genes)
```

```
## 
##   Level 6:   17 nodes to be scored   (17 eliminated genes)
```

```
## 
##   Level 5:   17 nodes to be scored   (110 eliminated genes)
```

```
## 
##   Level 4:   16 nodes to be scored   (110 eliminated genes)
```

```
## 
##   Level 3:   16 nodes to be scored   (110 eliminated genes)
```

```
## 
##   Level 2:   2 nodes to be scored    (110 eliminated genes)
```

```
## 
##   Level 1:   1 nodes to be scored    (110 eliminated genes)
```

```
resultFisher_BP_down0v8h_elim <- runTest(GOdata_BP_down0v8h, algorithm = "elim", statistic = "fisher")
```

```
## 
##           -- Elim Algorithm -- 
## 
##       the algorithm is scoring 368 nontrivial nodes
##       parameters: 
##           test statistic: fisher
##           cutOff: 0.01
```

```
## 
##   Level 13:  2 nodes to be scored    (0 eliminated genes)
```

```
## 
##   Level 12:  2 nodes to be scored    (0 eliminated genes)
```

```
## 
##   Level 11:  8 nodes to be scored    (0 eliminated genes)
```

```
## 
##   Level 10:  23 nodes to be scored   (0 eliminated genes)
```

```
## 
##   Level 9:   32 nodes to be scored   (0 eliminated genes)
```

```
## 
##   Level 8:   45 nodes to be scored   (7 eliminated genes)
```

```
## 
##   Level 7:   47 nodes to be scored   (7 eliminated genes)
```

```
## 
##   Level 6:   61 nodes to be scored   (12 eliminated genes)
```

```
## 
##   Level 5:   67 nodes to be scored   (12 eliminated genes)
```

```
## 
##   Level 4:   43 nodes to be scored   (12 eliminated genes)
```

```
## 
##   Level 3:   30 nodes to be scored   (12 eliminated genes)
```

```
## 
##   Level 2:   7 nodes to be scored    (126 eliminated genes)
```

```
## 
##   Level 1:   1 nodes to be scored    (126 eliminated genes)
```

```
resultFisher_MF_down0v8h_elim <- runTest(GOdata_MF_down0v8h, algorithm = "elim", statistic = "fisher")
```

```
## 
##           -- Elim Algorithm -- 
## 
##       the algorithm is scoring 228 nontrivial nodes
##       parameters: 
##           test statistic: fisher
##           cutOff: 0.01
```

```
## 
##   Level 11:  1 nodes to be scored    (0 eliminated genes)
```

```
## 
##   Level 10:  1 nodes to be scored    (0 eliminated genes)
```

```
## 
##   Level 9:   7 nodes to be scored    (0 eliminated genes)
```

```
## 
##   Level 8:   10 nodes to be scored   (0 eliminated genes)
```

```
## 
##   Level 7:   25 nodes to be scored   (0 eliminated genes)
```

```
## 
##   Level 6:   50 nodes to be scored   (0 eliminated genes)
```

```
## 
##   Level 5:   54 nodes to be scored   (40 eliminated genes)
```

```
## 
##   Level 4:   46 nodes to be scored   (75 eliminated genes)
```

```
## 
##   Level 3:   23 nodes to be scored   (75 eliminated genes)
```

```
## 
##   Level 2:   10 nodes to be scored   (75 eliminated genes)
```

```
## 
##   Level 1:   1 nodes to be scored    (75 eliminated genes)
```

```
resultFisher_CC_down0v8h_elim <- runTest(GOdata_CC_down0v8h, algorithm = "elim", statistic = "fisher")
```

```
## 
##           -- Elim Algorithm -- 
## 
##       the algorithm is scoring 117 nontrivial nodes
##       parameters: 
##           test statistic: fisher
##           cutOff: 0.01
```

```
## 
##   Level 12:  2 nodes to be scored    (0 eliminated genes)
```

```
## 
##   Level 11:  4 nodes to be scored    (0 eliminated genes)
```

```
## 
##   Level 10:  5 nodes to be scored    (0 eliminated genes)
```

```
## 
##   Level 9:   9 nodes to be scored    (0 eliminated genes)
```

```
## 
##   Level 8:   12 nodes to be scored   (0 eliminated genes)
```

```
## 
##   Level 7:   15 nodes to be scored   (0 eliminated genes)
```

```
## 
##   Level 6:   20 nodes to be scored   (29 eliminated genes)
```

```
## 
##   Level 5:   17 nodes to be scored   (29 eliminated genes)
```

```
## 
##   Level 4:   18 nodes to be scored   (29 eliminated genes)
```

```
## 
##   Level 3:   12 nodes to be scored   (312 eliminated genes)
```

```
## 
##   Level 2:   2 nodes to be scored    (312 eliminated genes)
```

```
## 
##   Level 1:   1 nodes to be scored    (312 eliminated genes)
```

```
#extract the significant GO terms
up0v8h_BP_elim <- GenTable(GOdata_BP_up0v8h, classic = resultFisher_BP_up0v8h_elim, 
                            orderBy = "weight", ranksOf = "weight", topNodes = 50)
up0v8h_MF_elim <- GenTable(GOdata_MF_up0v8h, classic = resultFisher_MF_up0v8h_elim, 
                            orderBy = "weight", ranksOf = "weight", topNodes = 50)
up0v8h_CC_elim <- GenTable(GOdata_CC_up0v8h, classic = resultFisher_CC_up0v8h_elim, 
                            orderBy = "weight", ranksOf = "weight", topNodes = 50)
down0v8h_BP_elim <- GenTable(GOdata_BP_down0v8h, classic = resultFisher_BP_down0v8h_elim, 
                              orderBy = "weight", ranksOf = "weight", topNodes = 50)
down0v8h_MF_elim <- GenTable(GOdata_MF_down0v8h, classic = resultFisher_MF_down0v8h_elim, 
                              orderBy = "weight", ranksOf = "weight", topNodes = 50)
down0v8h_CC_elim <- GenTable(GOdata_CC_down0v8h, classic = resultFisher_CC_down0v8h_elim, 
                              orderBy = "weight", ranksOf = "weight", topNodes = 50)

#Write the tables to you desired location to run through revigo using code below as a template.
#write.table(up0v8h_BP_elim,
#            "/My Drive/ShortTermStress-IlluminaData/FullExp/Cryp-DE/data/ReviGo_files/BP_up8h_elim.txt")
#write.table(up0v8h_MF_elim,
#            "/My Drive/ShortTermStress-IlluminaData/FullExp/Cryp-DE/data/ReviGo_files/MF_up8h_elim.txt")
#write.table(up0v8h_CC_elim,
#            "/My Drive/ShortTermStress-IlluminaData/FullExp/Cryp-DE/data/ReviGo_files/CC_up8h_elim.txt")
#write.table(down0v8h_BP_elim,
#            "/My Drive/ShortTermStress-IlluminaData/FullExp/Cryp-DE/data/ReviGo_files/BP_dn8h_elim.txt")
#write.table(down0v8h_MF_elim,
#            "/My Drive/ShortTermStress-IlluminaData/FullExp/Cryp-DE/data/ReviGo_files/MF_dn8h_elim.txt")
#write.table(down0v8h_CC_elim,
#            "/My Drive/ShortTermStress-IlluminaData/FullExp/Cryp-DE/data/ReviGo_files/CC_dn8h_elim.txt")
```

10 hour timepoint; early acclimation

```
#Time 0 vs 10 hour
sign_0v10h <- rownames(keep_2_SignDEG)[keep_2_SignDEG[,"C0m-C10hr"]==1] 

length(sign_0v10h)
```

```
## [1] 1432
```

```
sign2LFC_0v10h <- sign2LFC [rownames(sign2LFC) %in% sign_0v10h, ]  
logFCup_0v10h <- subset(sign2LFC_0v10h, (`0v10h_Padj`) <= 0.01 & (`0v10h`) > 0)
logFCdown_0v10h <- subset(sign2LFC_0v10h, (`0v10h_Padj`) <= 0.01 & (`0v10h`) < 0)

logFCup_0v10h_GOEnrich  = rownames(subset(logFCup_0v10h, rownames(logFCup_0v10h)%in%genesOfInterest_allSign))
logFCdown_0v10h_GOEnrich  = rownames(subset(logFCdown_0v10h, rownames(logFCdown_0v10h)%in%genesOfInterest_allSign))


#create gene list for input in topGO for 10 hours
geneList_up0v10h = factor(as.integer(geneUniverse %in% logFCup_0v10h_GOEnrich))
names(geneList_up0v10h) = geneUniverse
str(geneList_up0v10h)
```

```
##  Factor w/ 2 levels "0","1": 1 1 1 1 1 1 1 1 1 1 ...
##  - attr(*, "names")= chr [1:6309] "gene_idGO.ID" "CCRYP_013698" "CCRYP_013706" "CCRYP_013704" ...
```

```
geneList_down0v10h = factor(as.integer(geneUniverse %in% logFCdown_0v10h_GOEnrich))
names(geneList_down0v10h) = geneUniverse
str(geneList_down0v10h)
```

```
##  Factor w/ 2 levels "0","1": 1 1 1 1 1 1 1 2 1 2 ...
##  - attr(*, "names")= chr [1:6309] "gene_idGO.ID" "CCRYP_013698" "CCRYP_013706" "CCRYP_013704" ...
```

```
##create a topGO object (for up GOs)
GOdata_BP_up0v10h = new("topGOdata", ontology="BP", allGenes=geneList_up0v10h, 
                        annot = annFUN.gene2GO, gene2GO = geneID2GO)
```

```
## 
## Building most specific GOs .....
```

```
##  ( 624 GO terms found. )
```

```
## 
## Build GO DAG topology ..........
```

```
##  ( 1729 GO terms and 3515 relations. )
```

```
## 
## Annotating nodes ...............
```

```
##  ( 3345 genes annotated to the GO terms. )
```

```
GOdata_MF_up0v10h = new("topGOdata", ontology="MF", allGenes=geneList_up0v10h, 
                        annot = annFUN.gene2GO, gene2GO = geneID2GO)
```

```
## 
## Building most specific GOs .....
```

```
##  ( 736 GO terms found. )
```

```
## 
## Build GO DAG topology ..........
```

```
##  ( 1096 GO terms and 1406 relations. )
```

```
## 
## Annotating nodes ...............
```

```
##  ( 5189 genes annotated to the GO terms. )
```

```
GOdata_CC_up0v10h = new("topGOdata", ontology="CC", allGenes=geneList_up0v10h, 
                        annot = annFUN.gene2GO, gene2GO = geneID2GO)
```

```
## 
## Building most specific GOs .....
```

```
##  ( 235 GO terms found. )
```

```
## 
## Build GO DAG topology ..........
```

```
##  ( 452 GO terms and 812 relations. )
```

```
## 
## Annotating nodes ...............
```

```
##  ( 1987 genes annotated to the GO terms. )
```

```
##create a topGO object (for down GOs)
GOdata_BP_down0v10h  = new("topGOdata", ontology="BP", allGenes=geneList_down0v10h , 
                           annot = annFUN.gene2GO, gene2GO = geneID2GO)
```

```
## 
## Building most specific GOs .....
```

```
##  ( 624 GO terms found. )
```

```
## 
## Build GO DAG topology ..........
```

```
##  ( 1729 GO terms and 3515 relations. )
```

```
## 
## Annotating nodes ...............
```

```
##  ( 3345 genes annotated to the GO terms. )
```

```
GOdata_MF_down0v10h  = new("topGOdata", ontology="MF", allGenes=geneList_down0v10h , 
                           annot = annFUN.gene2GO, gene2GO = geneID2GO)
```

```
## 
## Building most specific GOs .....
```

```
##  ( 736 GO terms found. )
```

```
## 
## Build GO DAG topology ..........
```

```
##  ( 1096 GO terms and 1406 relations. )
```

```
## 
## Annotating nodes ...............
```

```
##  ( 5189 genes annotated to the GO terms. )
```

```
GOdata_CC_down0v10h  = new("topGOdata", ontology="CC", allGenes=geneList_down0v10h , 
                           annot = annFUN.gene2GO, gene2GO = geneID2GO)
```

```
## 
## Building most specific GOs .....
```

```
##  ( 235 GO terms found. )
```

```
## 
## Build GO DAG topology ..........
```

```
##  ( 452 GO terms and 812 relations. )
```

```
## 
## Annotating nodes ...............
```

```
##  ( 1987 genes annotated to the GO terms. )
```

```
#run Fisher's exact test
resultFisher_BP_up0v10h_elim <- runTest(GOdata_BP_up0v10h, algorithm = "elim", statistic = "fisher")
```

```
## 
##           -- Elim Algorithm -- 
## 
##       the algorithm is scoring 498 nontrivial nodes
##       parameters: 
##           test statistic: fisher
##           cutOff: 0.01
```

```
## 
##   Level 13:  4 nodes to be scored    (0 eliminated genes)
```

```
## 
##   Level 12:  11 nodes to be scored   (6 eliminated genes)
```

```
## 
##   Level 11:  25 nodes to be scored   (6 eliminated genes)
```

```
## 
##   Level 10:  36 nodes to be scored   (30 eliminated genes)
```

```
## 
##   Level 9:   57 nodes to be scored   (61 eliminated genes)
```

```
## 
##   Level 8:   58 nodes to be scored   (64 eliminated genes)
```

```
## 
##   Level 7:   70 nodes to be scored   (64 eliminated genes)
```

```
## 
##   Level 6:   82 nodes to be scored   (67 eliminated genes)
```

```
## 
##   Level 5:   76 nodes to be scored   (84 eliminated genes)
```

```
## 
##   Level 4:   44 nodes to be scored   (84 eliminated genes)
```

```
## 
##   Level 3:   28 nodes to be scored   (84 eliminated genes)
```

```
## 
##   Level 2:   6 nodes to be scored    (84 eliminated genes)
```

```
## 
##   Level 1:   1 nodes to be scored    (84 eliminated genes)
```

```
resultFisher_MF_up0v10h_elim <- runTest(GOdata_MF_up0v10h, algorithm = "elim", statistic = "fisher")
```

```
## 
##           -- Elim Algorithm -- 
## 
##       the algorithm is scoring 313 nontrivial nodes
##       parameters: 
##           test statistic: fisher
##           cutOff: 0.01
```

```
## 
##   Level 9:   6 nodes to be scored    (0 eliminated genes)
```

```
## 
##   Level 8:   14 nodes to be scored   (0 eliminated genes)
```

```
## 
##   Level 7:   39 nodes to be scored   (0 eliminated genes)
```

```
## 
##   Level 6:   74 nodes to be scored   (3 eliminated genes)
```

```
## 
##   Level 5:   73 nodes to be scored   (3 eliminated genes)
```

```
## 
##   Level 4:   66 nodes to be scored   (3 eliminated genes)
```

```
## 
##   Level 3:   29 nodes to be scored   (5 eliminated genes)
```

```
## 
##   Level 2:   11 nodes to be scored   (5 eliminated genes)
```

```
## 
##   Level 1:   1 nodes to be scored    (5 eliminated genes)
```

```
resultFisher_CC_up0v10h_elim <- runTest(GOdata_CC_up0v10h, algorithm = "elim", statistic = "fisher")
```

```
## 
##           -- Elim Algorithm -- 
## 
##       the algorithm is scoring 126 nontrivial nodes
##       parameters: 
##           test statistic: fisher
##           cutOff: 0.01
```

```
## 
##   Level 13:  1 nodes to be scored    (0 eliminated genes)
```

```
## 
##   Level 12:  1 nodes to be scored    (0 eliminated genes)
```

```
## 
##   Level 11:  2 nodes to be scored    (0 eliminated genes)
```

```
## 
##   Level 10:  5 nodes to be scored    (0 eliminated genes)
```

```
## 
##   Level 9:   8 nodes to be scored    (0 eliminated genes)
```

```
## 
##   Level 8:   15 nodes to be scored   (0 eliminated genes)
```

```
## 
##   Level 7:   18 nodes to be scored   (14 eliminated genes)
```

```
## 
##   Level 6:   18 nodes to be scored   (14 eliminated genes)
```

```
## 
##   Level 5:   22 nodes to be scored   (14 eliminated genes)
```

```
## 
##   Level 4:   17 nodes to be scored   (14 eliminated genes)
```

```
## 
##   Level 3:   16 nodes to be scored   (33 eliminated genes)
```

```
## 
##   Level 2:   2 nodes to be scored    (74 eliminated genes)
```

```
## 
##   Level 1:   1 nodes to be scored    (74 eliminated genes)
```

```
resultFisher_BP_down0v10h_elim <- runTest(GOdata_BP_down0v10h, algorithm = "elim", statistic = "fisher")
```

```
## 
##           -- Elim Algorithm -- 
## 
##       the algorithm is scoring 485 nontrivial nodes
##       parameters: 
##           test statistic: fisher
##           cutOff: 0.01
```

```
## 
##   Level 13:  2 nodes to be scored    (0 eliminated genes)
```

```
## 
##   Level 12:  5 nodes to be scored    (0 eliminated genes)
```

```
## 
##   Level 11:  16 nodes to be scored   (0 eliminated genes)
```

```
## 
##   Level 10:  38 nodes to be scored   (0 eliminated genes)
```

```
## 
##   Level 9:   52 nodes to be scored   (0 eliminated genes)
```

```
## 
##   Level 8:   67 nodes to be scored   (0 eliminated genes)
```

```
## 
##   Level 7:   69 nodes to be scored   (53 eliminated genes)
```

```
## 
##   Level 6:   79 nodes to be scored   (65 eliminated genes)
```

```
## 
##   Level 5:   77 nodes to be scored   (65 eliminated genes)
```

```
## 
##   Level 4:   46 nodes to be scored   (65 eliminated genes)
```

```
## 
##   Level 3:   27 nodes to be scored   (65 eliminated genes)
```

```
## 
##   Level 2:   6 nodes to be scored    (65 eliminated genes)
```

```
## 
##   Level 1:   1 nodes to be scored    (65 eliminated genes)
```

```
resultFisher_MF_down0v10h_elim <- runTest(GOdata_MF_down0v10h, algorithm = "elim", statistic = "fisher")
```

```
## 
##           -- Elim Algorithm -- 
## 
##       the algorithm is scoring 302 nontrivial nodes
##       parameters: 
##           test statistic: fisher
##           cutOff: 0.01
```

```
## 
##   Level 11:  1 nodes to be scored    (0 eliminated genes)
```

```
## 
##   Level 10:  2 nodes to be scored    (0 eliminated genes)
```

```
## 
##   Level 9:   9 nodes to be scored    (0 eliminated genes)
```

```
## 
##   Level 8:   14 nodes to be scored   (0 eliminated genes)
```

```
## 
##   Level 7:   35 nodes to be scored   (0 eliminated genes)
```

```
## 
##   Level 6:   75 nodes to be scored   (91 eliminated genes)
```

```
## 
##   Level 5:   68 nodes to be scored   (204 eliminated genes)
```

```
## 
##   Level 4:   62 nodes to be scored   (215 eliminated genes)
```

```
## 
##   Level 3:   26 nodes to be scored   (579 eliminated genes)
```

```
## 
##   Level 2:   9 nodes to be scored    (579 eliminated genes)
```

```
## 
##   Level 1:   1 nodes to be scored    (579 eliminated genes)
```

```
resultFisher_CC_down0v10h_elim <- runTest(GOdata_CC_down0v10h, algorithm = "elim", statistic = "fisher")
```

```
## 
##           -- Elim Algorithm -- 
## 
##       the algorithm is scoring 115 nontrivial nodes
##       parameters: 
##           test statistic: fisher
##           cutOff: 0.01
```

```
## 
##   Level 12:  2 nodes to be scored    (0 eliminated genes)
```

```
## 
##   Level 11:  3 nodes to be scored    (0 eliminated genes)
```

```
## 
##   Level 10:  5 nodes to be scored    (0 eliminated genes)
```

```
## 
##   Level 9:   8 nodes to be scored    (0 eliminated genes)
```

```
## 
##   Level 8:   11 nodes to be scored   (0 eliminated genes)
```

```
## 
##   Level 7:   16 nodes to be scored   (0 eliminated genes)
```

```
## 
##   Level 6:   18 nodes to be scored   (0 eliminated genes)
```

```
## 
##   Level 5:   17 nodes to be scored   (0 eliminated genes)
```

```
## 
##   Level 4:   16 nodes to be scored   (471 eliminated genes)
```

```
## 
##   Level 3:   16 nodes to be scored   (474 eliminated genes)
```

```
## 
##   Level 2:   2 nodes to be scored    (474 eliminated genes)
```

```
## 
##   Level 1:   1 nodes to be scored    (474 eliminated genes)
```

```
#extract the significant GO terms
up0v10h_BP_elim <- GenTable(GOdata_BP_up0v10h, classic = resultFisher_BP_up0v10h_elim, 
                            orderBy = "weight", ranksOf = "weight", topNodes = 50)
up0v10h_MF_elim <- GenTable(GOdata_MF_up0v10h, classic = resultFisher_MF_up0v10h_elim, 
                            orderBy = "weight", ranksOf = "weight", topNodes = 50)
up0v10h_CC_elim <- GenTable(GOdata_CC_up0v10h, classic = resultFisher_CC_up0v10h_elim, 
                            orderBy = "weight", ranksOf = "weight", topNodes = 50)
down0v10h_BP_elim <- GenTable(GOdata_BP_down0v10h, classic = resultFisher_BP_down0v10h_elim, 
                              orderBy = "weight", ranksOf = "weight", topNodes = 50)
down0v10h_MF_elim <- GenTable(GOdata_MF_down0v10h, classic = resultFisher_MF_down0v10h_elim, 
                              orderBy = "weight", ranksOf = "weight", topNodes = 50)
down0v10h_CC_elim <- GenTable(GOdata_CC_down0v10h, classic = resultFisher_CC_down0v10h_elim, 
                              orderBy = "weight", ranksOf = "weight", topNodes = 50)

#Write the tables to you desired location to run through revigo using code below as a template.
#write.table(up0v10h_BP_elim,
#            "/My Drive/ShortTermStress-IlluminaData/FullExp/Cryp-DE/data/ReviGo_files/BP_up10h_elim.txt")
#write.table(up0v10h_MF_elim,
#            "/My Drive/ShortTermStress-IlluminaData/FullExp/Cryp-DE/data/ReviGo_files/MF_up10h_elim.txt")
#write.table(up0v10h_CC_elim,
#            "/My Drive/ShortTermStress-IlluminaData/FullExp/Cryp-DE/data/ReviGo_files/CC_up10h_elim.txt")
#write.table(down0v10h_BP_elim,
#            "/My Drive/ShortTermStress-IlluminaData/FullExp/Cryp-DE/data/ReviGo_files/BP_dn10h_elim.txt")
#write.table(down0v10h_MF_elim,
#            "/My Drive/ShortTermStress-IlluminaData/FullExp/Cryp-DE/data/ReviGo_files/MF_dn10h_elim.txt")
#write.table(down0v10h_CC_elim,
#            "/My Drive/ShortTermStress-IlluminaData/FullExp/Cryp-DE/data/ReviGo_files/CC_dn10h_elim.txt")
```

Import the necessary Revigo output columns back in. These files are
not available because you’ll want to run ReviGo yourself to pick your
similarity values.

```
dr_1h_bp = read.table("/My Drive/ShortTermStress-IlluminaData/FullExp/Cryp-DE/data/ReviGo_files/BP_dn1h_Revigo.tsv",
                       colClasses = c(NA,"NULL","NULL","NULL","NULL","NULL","NULL","NULL","NULL","NULL","NULL","NULL","NULL",NA),
                       header=TRUE) #biological process results
ur_1h_bp = read.table("/My Drive/ShortTermStress-IlluminaData/FullExp/Cryp-DE/data/ReviGo_files/BP_up1h_Revigo.tsv",
                     colClasses = c(NA,"NULL","NULL","NULL","NULL","NULL","NULL","NULL","NULL","NULL","NULL","NULL","NULL",NA),
                     header=TRUE) #biological process results
dr_1h_cc = read.table("/My Drive/ShortTermStress-IlluminaData/FullExp/Cryp-DE/data/ReviGo_files/CC_dn1h_Revigo.tsv",
                     colClasses = c(NA,"NULL","NULL","NULL","NULL","NULL","NULL","NULL","NULL","NULL","NULL","NULL","NULL",NA),
                     header=TRUE) #cellular component results
ur_1h_cc = read.table("/My Drive/ShortTermStress-IlluminaData/FullExp/Cryp-DE/data/ReviGo_files/CC_up1h_Revigo.tsv",
                     colClasses = c(NA,"NULL","NULL","NULL","NULL","NULL","NULL","NULL","NULL","NULL","NULL","NULL","NULL",NA),
                     header=TRUE) #cellular component results
dr_1h_mf = read.table("/My Drive/ShortTermStress-IlluminaData/FullExp/Cryp-DE/data/ReviGo_files/MF_dn1h_Revigo.tsv",
                     colClasses = c(NA,"NULL","NULL","NULL","NULL","NULL","NULL","NULL","NULL","NULL","NULL","NULL","NULL",NA),
                     header=TRUE) #molecular function results
ur_1h_mf = read.table("/My Drive/ShortTermStress-IlluminaData/FullExp/Cryp-DE/data/ReviGo_files/MF_up1h_Revigo.tsv",
                     colClasses = c(NA,"NULL","NULL","NULL","NULL","NULL","NULL","NULL","NULL","NULL","NULL","NULL","NULL",NA),
                     header=TRUE) #molecular function results
#
dr_10h_bp = read.table("/My Drive/ShortTermStress-IlluminaData/FullExp/Cryp-DE/data/ReviGo_files/BP_dn10h_Revigo.tsv",
                     colClasses = c(NA,"NULL","NULL","NULL","NULL","NULL","NULL","NULL","NULL","NULL","NULL","NULL","NULL",NA),
                     header=TRUE) #biological process results
ur_10h_bp = read.table("/My Drive/ShortTermStress-IlluminaData/FullExp/Cryp-DE/data/ReviGo_files/BP_up10h_Revigo.tsv",
                     colClasses = c(NA,"NULL","NULL","NULL","NULL","NULL","NULL","NULL","NULL","NULL","NULL","NULL","NULL",NA),
                     header=TRUE) #biological process results
dr_10h_cc = read.table("/My Drive/ShortTermStress-IlluminaData/FullExp/Cryp-DE/data/ReviGo_files/CC_dn10h_Revigo.tsv",
                     colClasses = c(NA,"NULL","NULL","NULL","NULL","NULL","NULL","NULL","NULL","NULL","NULL","NULL","NULL",NA),
                     header=TRUE) #cellular component results
ur_10h_cc = read.table("/My Drive/ShortTermStress-IlluminaData/FullExp/Cryp-DE/data/ReviGo_files/CC_up10h_Revigo.tsv",
                     colClasses = c(NA,"NULL","NULL","NULL","NULL","NULL","NULL","NULL","NULL","NULL","NULL","NULL","NULL",NA),
                     header=TRUE) #cellular component results
dr_10h_mf = read.table("/My Drive/ShortTermStress-IlluminaData/FullExp/Cryp-DE/data/ReviGo_files/MF_dn10h_Revigo.tsv",
                     colClasses = c(NA,"NULL","NULL","NULL","NULL","NULL","NULL","NULL","NULL","NULL","NULL","NULL","NULL",NA),
                     header=TRUE) #molecular function results
ur_10h_mf = read.table("/My Drive/ShortTermStress-IlluminaData/FullExp/Cryp-DE/data/ReviGo_files/MF_up10h_Revigo.tsv",
                     colClasses = c(NA,"NULL","NULL","NULL","NULL","NULL","NULL","NULL","NULL","NULL","NULL","NULL","NULL",NA),
                     header=TRUE) #molecular function results

names(dr_1h_bp) <- c('GO.ID','Eliminated')
names(dr_1h_cc) <- c('GO.ID','Eliminated')
names(dr_1h_mf) <- c('GO.ID','Eliminated')
names(ur_1h_bp) <- c('GO.ID','Eliminated')
names(ur_1h_cc) <- c('GO.ID','Eliminated')
names(ur_1h_mf) <- c('GO.ID','Eliminated')

names(dr_10h_bp) <- c('GO.ID','Eliminated')
names(dr_10h_cc) <- c('GO.ID','Eliminated')
names(dr_10h_mf) <- c('GO.ID','Eliminated')
names(ur_10h_bp) <- c('GO.ID','Eliminated')
names(ur_10h_cc) <- c('GO.ID','Eliminated')
names(ur_10h_mf) <- c('GO.ID','Eliminated')

#resave(dr_1h_bp,dr_1h_cc,dr_1h_mf,ur_1h_bp,ur_1h_cc,ur_1h_mf,
#       dr_10h_bp,dr_10h_cc,dr_10h_mf,ur_10h_bp,ur_10h_cc,ur_10h_mf,
#       file = "/My Drive/ShortTermStress-IlluminaData/FullExp/Cryp-DE/data/import_rmd_vars.RData")
```

##### 4.4.1. Comparing 60 min and 10 h enrichments

###### 4.4.1.1 For a quick glance, we compiled it all into a single table.

```
ur_10h_bp <- up0v10h_BP_elim %>% 
  full_join(., ur_10h_bp, by = "GO.ID", copy = FALSE, suffix = c(".x",".y"))
ur_10h_bp$timept <- '10h'
ur_10h_bp$category <- 'biological process'
ur_10h_bp$degree <- 'upregulated'
ur_10h_cc <- up0v10h_CC_elim %>% 
  full_join(., ur_10h_cc, by = "GO.ID", copy = FALSE, suffix = c(".x",".y"))
ur_10h_cc$timept <- '10h'
ur_10h_cc$category <- 'cellular component'
ur_10h_cc$degree <- 'upregulated'
ur_10h_mf <- up0v10h_MF_elim %>% 
  full_join(., ur_10h_mf, by = "GO.ID", copy = FALSE, suffix = c(".x",".y"))
ur_10h_mf$timept <- '10h'
ur_10h_mf$category <- 'molecular function'
ur_10h_mf$degree <- 'upregulated'
dr_10h_bp <- down0v10h_BP_elim %>% 
  full_join(., dr_10h_bp, by = "GO.ID", copy = FALSE, suffix = c(".x",".y"))
dr_10h_bp$timept <- '10h'
dr_10h_bp$category <- 'biological process'
dr_10h_bp$degree <- 'downregulated'
dr_10h_cc <- down0v10h_CC_elim %>% 
  full_join(., dr_10h_cc, by = "GO.ID", copy = FALSE, suffix = c(".x",".y"))
dr_10h_cc$timept <- '10h'
dr_10h_cc$category <- 'cellular component'
dr_10h_cc$degree <- 'downregulated'
dr_10h_mf <- down0v10h_MF_elim %>% 
  full_join(., dr_10h_mf, by = "GO.ID", copy = FALSE, suffix = c(".x",".y"))
dr_10h_mf$timept <- '10h'
dr_10h_mf$category <- 'molecular function'
dr_10h_mf$degree <- 'downregulated'

ur_1h_bp <- up0v1h_BP_elim %>% 
  full_join(., ur_1h_bp, by = "GO.ID", copy = FALSE, suffix = c(".x",".y"))
ur_1h_bp$timept <- '1h'
ur_1h_bp$category <- 'biological process'
ur_1h_bp$degree <- 'upregulated'
ur_1h_cc <- up0v1h_CC_elim %>% 
  full_join(., ur_1h_cc, by = "GO.ID", copy = FALSE, suffix = c(".x",".y"))
ur_1h_cc$timept <- '1h'
ur_1h_cc$category <- 'cellular component'
ur_1h_cc$degree <- 'upregulated'
ur_1h_mf <- up0v1h_MF_elim %>% 
  full_join(., ur_1h_mf, by = "GO.ID", copy = FALSE, suffix = c(".x",".y"))
ur_1h_mf$timept <- '1h'
ur_1h_mf$category <- 'molecular function'
ur_1h_mf$degree <- 'upregulated'
dr_1h_bp <- down0v1h_BP_elim %>% 
  full_join(., dr_1h_bp, by = "GO.ID", copy = FALSE, suffix = c(".x",".y"))
dr_1h_bp$timept <- '1h'
dr_1h_bp$category <- 'biological process'
dr_1h_bp$degree <- 'downregulated'
dr_1h_cc <- down0v1h_CC_elim %>% 
  full_join(., dr_1h_cc, by = "GO.ID", copy = FALSE, suffix = c(".x",".y"))
dr_1h_cc$timept <- '1h'
dr_1h_cc$category <- 'cellular component'
dr_1h_cc$degree <- 'downregulated'
dr_1h_mf <- down0v1h_MF_elim %>% 
  full_join(., dr_1h_mf, by = "GO.ID", copy = FALSE, suffix = c(".x",".y"))
dr_1h_mf$timept <- '1h'
dr_1h_mf$category <- 'molecular function'
dr_1h_mf$degree <- 'downregulated'

#After reviewing the cellular component data, we opted not to include it in the comparison between timepoints since it didn't provide much additional depth.
go_data <- rbind(ur_10h_bp,ur_10h_mf,dr_10h_bp,dr_10h_mf,ur_1h_bp,ur_1h_mf,dr_1h_bp,dr_1h_mf)

names(go_data) <- c("GO.ID","Term","Annotated","Sign","Exp","Pvalue","Eliminated","timept","category","degree")
go_data$GO.Term <- paste(go_data$Term, go_data$GO.ID, sep=" ")
go_data_reduced <- filter(go_data, Eliminated == "False") 
go_data_reduced <- subset(go_data_reduced,select=c("GO.Term","Annotated","Pvalue","timept","category","degree"))
go_data_reduced <- transform(go_data_reduced, Percentage = (Annotated / 3193) * 100)
go_data_reduced <- subset(go_data_reduced, (`Percentage`) > 1) 
go_data_reduced <- subset(go_data_reduced, (`Pvalue`) < 0.05) 
go_data_reduced <- go_data_reduced %>%
  mutate(Percentage = ifelse(degree == "upregulated",
                             Percentage,
                             -1*Percentage))
go_data_reduced$Pvalue<-gsub("<","",as.character(go_data_reduced$Pvalue))
```

###### 4.4.1.2 Create the variables for the graphs and plot.

```
#Data for graph

data_1h <- filter(go_data_reduced, timept == "1h")
data_1h <- data_1h[
    order( data_1h[,5], data_1h[,6], data_1h[,7]),
]
data_1h$GO.Term <- factor(data_1h$GO.Term, levels = data_1h$GO.Term)
data_1h$Pvalue <- as.numeric(data_1h$Pvalue)

data_10h <- filter(go_data_reduced, timept == "10h")
data_10h <- data_10h[
    order( data_10h[,5], data_10h[,6], data_10h[,7]),
]
data_10h$GO.Term <- factor(data_10h$GO.Term, levels = data_10h$GO.Term)
data_10h$Pvalue <- as.numeric(data_10h$Pvalue)
y_breaks <- pretty(go_data_reduced$Percentage)

GO_plot_1h <- ggplot(data_1h,
                      aes(x=GO.Term,
                         y=Percentage,
                         fill=Pvalue)) +
  geom_bar(stat = "identity") +
  coord_flip()+
  ylim(-25,25) +
  xlab("Enriched Gene Ontology Terms") + 
  ylab("Percentage of Significant DEG") +
  ggtitle("0m vs 1h") +
  theme_classic() +
  theme(plot.title = element_text(hjust = 0.4))

GO_plot_10h <- ggplot(data_10h,
                      aes(x=GO.Term,
                         y=Percentage,
                         fill=Pvalue)) +
  geom_bar(stat = "identity") +
  coord_flip() +
  ylim(-25,25) +
  xlab("Enriched Gene Ontology Terms") + 
  ylab("Percentage of Significant DEG") +
  ggtitle("0m vs 10h") +
  theme_classic() +
  theme(plot.title = element_text(hjust = 0.4))

#Figure plotting
(GO_plot_1h / GO_plot_10h)
```

```
## Warning: Removed 1 rows containing missing values (position_stack).
```

###### 4.4.1.3 Hypergeometric tests

We also looked at the significance of shared DE genes between 60 min,
10 h, and the long-term datasets.

```
#Hypergeometric assessment of overlap of DE genes between basically everything we can think of.
genes_1h_up <- rownames(logFCup_0v1h)
genes_1h_down <- rownames(logFCdown_0v1h)
genes_10h_up <- rownames(logFCup_0v10h)
genes_10h_down <- rownames(logFCdown_0v10h)
genes_LT_up <- rownames(nakov_DE_ur)
genes_LT_down <- rownames(nakov_DE_dr)
genes_1h <- rownames(sign2LFC_0v1h)
genes_10h <- rownames(sign2LFC_0v10h)
genes_LT <- rownames(nakov_DE)
genes_all_signDE <- rownames(Padjscreen_sorted)
genes_all <- rownames(OnlySignGenes_ConStage)
genes_2 <- rownames(DEG2sig_results)

#ID the overlapping genes

int_up <- intersect (genes_1h_up, genes_10h_up)
int_down <- intersect (genes_1h_down, genes_10h_down)
int_updown <- intersect (genes_1h_up, genes_10h_down)
int_downup <- intersect (genes_1h_down, genes_10h_up)
int_1v10 <- intersect (genes_1h, genes_10h)
int_LTv1_up <- intersect (genes_1h_up, genes_LT_up)
int_LTv10_up <- intersect (genes_10h_up, genes_LT_up)
int_LTv1_dn <- intersect (genes_1h_down, genes_LT_down)
int_LTv10_dn <- intersect (genes_10h_down, genes_LT_down)
int_LTv1_updown <- intersect (genes_1h_up, genes_LT_down)
int_LTv1_downup <- intersect (genes_1h_down, genes_LT_up)
int_LTv10_updown <- intersect (genes_LT_up, genes_10h_down)
int_LTv10_downup <- intersect (genes_LT_down, genes_10h_up)
int_LTv10 <- intersect (genes_10h, genes_LT)
int_LTv1h <- intersect (genes_1h, genes_LT)
int_LTvAll <- intersect (genes_all_signDE, genes_LT)
int_All <- intersect (genes_all, genes_LT)
int_LTv2sig <- intersect (genes_2, genes_LT)

#Execute the hypergeometric test to determine if that number of overlapping genes is significant.

phyper(416,417,(2048-417),772,lower.tail = FALSE) # 1h vs 10h repressed (8.349016e-186)
```

```
## [1] 8.656944e-219
```

```
phyper(243,244,(1364-244),660,lower.tail = FALSE) #1h vs 10h induced (1.496532e-113)
```

```
## [1] 4.476689e-90
```

```
phyper(56,57,(1364-57),772,lower.tail = FALSE) #10h repressed vs 1h induced (8.057105e-42)
```

```
## [1] 3.195299e-15
```

```
phyper(92,93,(2048-93),660,lower.tail = FALSE) #1h repressed vs 10h induced (7.854042e-106)
```

```
## [1] 1.692235e-48
```

```
phyper(969,970,(3497-970),1431,lower.tail = FALSE) #1h signDE vs 10h signDE (0?????)
```

```
## [1] 0
```

```
phyper(258,259,(4298-259),1220,lower.tail = FALSE) #All short-term signDE vs long-term signDE (9.145136e-152)
```

```
## [1] 9.145136e-152
```

```
phyper(79,80,(1431-80),1220,lower.tail = FALSE) #All 10h signDE vs LT signDE (1.926452e-06)
```

```
## [1] 1.926452e-06
```

```
phyper(205,206,(3497-206),1220,lower.tail = FALSE) #All 1h signDE vs LT signDE (2.592496e-103)
```

```
## [1] 3.061351e-100
```

```
phyper(50,51,(1942-51),695,lower.tail = FALSE)#1h vs LT repressed (5.145085e-24)
```

```
## [1] 5.145085e-24
```

```
phyper(22,23,(772-23),695,lower.tail = FALSE)#10h vs LT repressed 0.0859712)
```

```
## [1] 0.0859712
```

```
phyper(42,43,(1555-43),525,lower.tail = FALSE)#1h vs LT induced (1.617967e-21)
```

```
## [1] 1.617967e-21
```

```
phyper(17,18,(659-18),525,lower.tail = FALSE)#10h vs LT induced (0.01572783)
```

```
## [1] 0.01572783
```

```
phyper(53,54,(1942-54),525,lower.tail = FALSE)#1h repressed vs LT induced (2.630185e-32)
```

```
## [1] 2.630185e-32
```

```
phyper(16,17,(772-17),525,lower.tail = FALSE)#10h repressed vs LT induced (0.001307894
```

```
## [1] 0.001307894
```

```
phyper(63,64,(1555-64),695,lower.tail = FALSE)#1h induced vs LT repressed (7.711969e-24)
```

```
## [1] 7.711969e-24
```

```
phyper(21,22,(695-22),659,lower.tail = FALSE)#10h induced vs LT repressed (0.3046189)
```

```
## [1] 0.3046189
```

##### 4.4.2 GO enrichment for different LFC cutoffs for 1h and 10 h.

We started with the genes at 1h that had an LFC less than 0.585 to
look at the primary processes enriched in lowly expressed genes during
peak hyposalinity stress.

```
##This is for looking at LFC 0.585 in 1 h 
LFCupless0.585_0v1h <- subset(sign2LFC_0v1h, (`0v1h_Padj`) <= 0.01 & (`0v1h`) < 0.585 & (`0v1h`) > 0 )
LFCdnless0.585_0v1h <- subset(sign2LFC_0v1h, (`0v1h_Padj`) <= 0.01 & (`0v1h`) > -0.585 & (`0v1h`) < 0)

LFCupless0.585_0v1h_GOEnrich  = rownames(subset(LFCupless0.585_0v1h, rownames(LFCupless0.585_0v1h)%in%genesOfInterest_allSign))
LFCdnless0.585_0v1h_GOEnrich  = rownames(subset(LFCdnless0.585_0v1h, rownames(LFCdnless0.585_0v1h)%in%genesOfInterest_allSign))


#create gene list for input in topGO
geneList_up1hLFCless0.585 = factor(as.integer(geneUniverse %in% LFCupless0.585_0v1h_GOEnrich))
names(geneList_up1hLFCless0.585) = geneUniverse
str(geneList_up1hLFCless0.585)
```

```
##  Factor w/ 2 levels "0","1": 1 1 1 1 1 1 1 1 1 1 ...
##  - attr(*, "names")= chr [1:6309] "gene_idGO.ID" "CCRYP_013698" "CCRYP_013706" "CCRYP_013704" ...
```

```
geneList_dn1hLFCless0.585 = factor(as.integer(geneUniverse %in% LFCdnless0.585_0v1h_GOEnrich))
names(geneList_dn1hLFCless0.585) = geneUniverse
str(geneList_dn1hLFCless0.585)
```

```
##  Factor w/ 2 levels "0","1": 1 1 1 1 2 1 1 2 1 1 ...
##  - attr(*, "names")= chr [1:6309] "gene_idGO.ID" "CCRYP_013698" "CCRYP_013706" "CCRYP_013704" ...
```

```
##create a topGO object (for up GOs)
GOdata_BP_up1hLFCless0.585 = new("topGOdata", ontology="BP", allGenes=geneList_up1hLFCless0.585, 
                        annot = annFUN.gene2GO, gene2GO = geneID2GO)
```

```
## 
## Building most specific GOs .....
```

```
##  ( 624 GO terms found. )
```

```
## 
## Build GO DAG topology ..........
```

```
##  ( 1729 GO terms and 3515 relations. )
```

```
## 
## Annotating nodes ...............
```

```
##  ( 3345 genes annotated to the GO terms. )
```

```
GOdata_MF_up1hLFCless0.585 = new("topGOdata", ontology="MF", allGenes=geneList_up1hLFCless0.585, 
                        annot = annFUN.gene2GO, gene2GO = geneID2GO)
```

```
## 
## Building most specific GOs .....
```

```
##  ( 736 GO terms found. )
```

```
## 
## Build GO DAG topology ..........
```

```
##  ( 1096 GO terms and 1406 relations. )
```

```
## 
## Annotating nodes ...............
```

```
##  ( 5189 genes annotated to the GO terms. )
```

```
GOdata_CC_up1hLFCless0.585 = new("topGOdata", ontology="CC", allGenes=geneList_up1hLFCless0.585, 
                        annot = annFUN.gene2GO, gene2GO = geneID2GO)
```

```
## 
## Building most specific GOs .....
```

```
##  ( 235 GO terms found. )
```

```
## 
## Build GO DAG topology ..........
```

```
##  ( 452 GO terms and 812 relations. )
```

```
## 
## Annotating nodes ...............
```

```
##  ( 1987 genes annotated to the GO terms. )
```

```
##create a topGO object (for down GOs)
GOdata_BP_dn1hLFCless0.585  = new("topGOdata", ontology="BP", allGenes=geneList_dn1hLFCless0.585, 
                           annot = annFUN.gene2GO, gene2GO = geneID2GO)
```

```
## 
## Building most specific GOs .....
```

```
##  ( 624 GO terms found. )
```

```
## 
## Build GO DAG topology ..........
```

```
##  ( 1729 GO terms and 3515 relations. )
```

```
## 
## Annotating nodes ...............
```

```
##  ( 3345 genes annotated to the GO terms. )
```

```
GOdata_MF_dn1hLFCless0.585  = new("topGOdata", ontology="MF", allGenes=geneList_dn1hLFCless0.585, 
                           annot = annFUN.gene2GO, gene2GO = geneID2GO)
```

```
## 
## Building most specific GOs .....
```

```
##  ( 736 GO terms found. )
```

```
## 
## Build GO DAG topology ..........
```

```
##  ( 1096 GO terms and 1406 relations. )
```

```
## 
## Annotating nodes ...............
```

```
##  ( 5189 genes annotated to the GO terms. )
```

```
GOdata_CC_dn1hLFCless0.585  = new("topGOdata", ontology="CC", allGenes=geneList_dn1hLFCless0.585, 
                           annot = annFUN.gene2GO, gene2GO = geneID2GO)
```

```
## 
## Building most specific GOs .....
```

```
##  ( 235 GO terms found. )
```

```
## 
## Build GO DAG topology ..........
```

```
##  ( 452 GO terms and 812 relations. )
```

```
## 
## Annotating nodes ...............
```

```
##  ( 1987 genes annotated to the GO terms. )
```

```
#run Fisher's exact test
resultFisher_BP_up1hLFCless0.585_elim <- runTest(GOdata_BP_up1hLFCless0.585, algorithm = "elim", statistic = "fisher")
```

```
## 
##           -- Elim Algorithm -- 
## 
##       the algorithm is scoring 333 nontrivial nodes
##       parameters: 
##           test statistic: fisher
##           cutOff: 0.01
```

```
## 
##   Level 13:  1 nodes to be scored    (0 eliminated genes)
```

```
## 
##   Level 12:  5 nodes to be scored    (0 eliminated genes)
```

```
## 
##   Level 11:  8 nodes to be scored    (0 eliminated genes)
```

```
## 
##   Level 10:  19 nodes to be scored   (0 eliminated genes)
```

```
## 
##   Level 9:   32 nodes to be scored   (0 eliminated genes)
```

```
## 
##   Level 8:   42 nodes to be scored   (0 eliminated genes)
```

```
## 
##   Level 7:   39 nodes to be scored   (30 eliminated genes)
```

```
## 
##   Level 6:   59 nodes to be scored   (41 eliminated genes)
```

```
## 
##   Level 5:   63 nodes to be scored   (41 eliminated genes)
```

```
## 
##   Level 4:   37 nodes to be scored   (41 eliminated genes)
```

```
## 
##   Level 3:   21 nodes to be scored   (41 eliminated genes)
```

```
## 
##   Level 2:   6 nodes to be scored    (41 eliminated genes)
```

```
## 
##   Level 1:   1 nodes to be scored    (41 eliminated genes)
```

```
resultFisher_MF_up1hLFCless0.585_elim <- runTest(GOdata_MF_up1hLFCless0.585, algorithm = "elim", statistic = "fisher")
```

```
## 
##           -- Elim Algorithm -- 
## 
##       the algorithm is scoring 295 nontrivial nodes
##       parameters: 
##           test statistic: fisher
##           cutOff: 0.01
```

```
## 
##   Level 11:  1 nodes to be scored    (0 eliminated genes)
```

```
## 
##   Level 10:  2 nodes to be scored    (7 eliminated genes)
```

```
## 
##   Level 9:   10 nodes to be scored   (7 eliminated genes)
```

```
## 
##   Level 8:   22 nodes to be scored   (7 eliminated genes)
```

```
## 
##   Level 7:   35 nodes to be scored   (28 eliminated genes)
```

```
## 
##   Level 6:   63 nodes to be scored   (28 eliminated genes)
```

```
## 
##   Level 5:   68 nodes to be scored   (28 eliminated genes)
```

```
## 
##   Level 4:   60 nodes to be scored   (39 eliminated genes)
```

```
## 
##   Level 3:   24 nodes to be scored   (342 eliminated genes)
```

```
## 
##   Level 2:   9 nodes to be scored    (342 eliminated genes)
```

```
## 
##   Level 1:   1 nodes to be scored    (342 eliminated genes)
```

```
resultFisher_CC_up1hLFCless0.585_elim <- runTest(GOdata_CC_up1hLFCless0.585, algorithm = "elim", statistic = "fisher")
```

```
## 
##           -- Elim Algorithm -- 
## 
##       the algorithm is scoring 70 nontrivial nodes
##       parameters: 
##           test statistic: fisher
##           cutOff: 0.01
```

```
## 
##   Level 10:  1 nodes to be scored    (0 eliminated genes)
```

```
## 
##   Level 9:   2 nodes to be scored    (0 eliminated genes)
```

```
## 
##   Level 8:   4 nodes to be scored    (0 eliminated genes)
```

```
## 
##   Level 7:   11 nodes to be scored   (0 eliminated genes)
```

```
## 
##   Level 6:   15 nodes to be scored   (0 eliminated genes)
```

```
## 
##   Level 5:   12 nodes to be scored   (0 eliminated genes)
```

```
## 
##   Level 4:   12 nodes to be scored   (471 eliminated genes)
```

```
## 
##   Level 3:   10 nodes to be scored   (471 eliminated genes)
```

```
## 
##   Level 2:   2 nodes to be scored    (838 eliminated genes)
```

```
## 
##   Level 1:   1 nodes to be scored    (838 eliminated genes)
```

```
resultFisher_BP_dn1hLFCless0.585_elim <- runTest(GOdata_BP_dn1hLFCless0.585, algorithm = "elim", statistic = "fisher")
```

```
## 
##           -- Elim Algorithm -- 
## 
##       the algorithm is scoring 405 nontrivial nodes
##       parameters: 
##           test statistic: fisher
##           cutOff: 0.01
```

```
## 
##   Level 14:  1 nodes to be scored    (0 eliminated genes)
```

```
## 
##   Level 13:  3 nodes to be scored    (0 eliminated genes)
```

```
## 
##   Level 12:  5 nodes to be scored    (0 eliminated genes)
```

```
## 
##   Level 11:  9 nodes to be scored    (0 eliminated genes)
```

```
## 
##   Level 10:  21 nodes to be scored   (0 eliminated genes)
```

```
## 
##   Level 9:   36 nodes to be scored   (0 eliminated genes)
```

```
## 
##   Level 8:   51 nodes to be scored   (5 eliminated genes)
```

```
## 
##   Level 7:   57 nodes to be scored   (183 eliminated genes)
```

```
## 
##   Level 6:   70 nodes to be scored   (183 eliminated genes)
```

```
## 
##   Level 5:   73 nodes to be scored   (197 eliminated genes)
```

```
## 
##   Level 4:   44 nodes to be scored   (345 eliminated genes)
```

```
## 
##   Level 3:   28 nodes to be scored   (345 eliminated genes)
```

```
## 
##   Level 2:   6 nodes to be scored    (350 eliminated genes)
```

```
## 
##   Level 1:   1 nodes to be scored    (350 eliminated genes)
```

```
resultFisher_MF_dn1hLFCless0.585_elim <- runTest(GOdata_MF_dn1hLFCless0.585, algorithm = "elim", statistic = "fisher")
```

```
## 
##           -- Elim Algorithm -- 
## 
##       the algorithm is scoring 216 nontrivial nodes
##       parameters: 
##           test statistic: fisher
##           cutOff: 0.01
```

```
## 
##   Level 10:  1 nodes to be scored    (0 eliminated genes)
```

```
## 
##   Level 9:   6 nodes to be scored    (0 eliminated genes)
```

```
## 
##   Level 8:   11 nodes to be scored   (0 eliminated genes)
```

```
## 
##   Level 7:   21 nodes to be scored   (0 eliminated genes)
```

```
## 
##   Level 6:   41 nodes to be scored   (4 eliminated genes)
```

```
## 
##   Level 5:   49 nodes to be scored   (4 eliminated genes)
```

```
## 
##   Level 4:   51 nodes to be scored   (397 eliminated genes)
```

```
## 
##   Level 3:   25 nodes to be scored   (397 eliminated genes)
```

```
## 
##   Level 2:   10 nodes to be scored   (497 eliminated genes)
```

```
## 
##   Level 1:   1 nodes to be scored    (506 eliminated genes)
```

```
resultFisher_CC_dn1hLFCless0.585_elim <- runTest(GOdata_CC_dn1hLFCless0.585, algorithm = "elim", statistic = "fisher")
```

```
## 
##           -- Elim Algorithm -- 
## 
##       the algorithm is scoring 104 nontrivial nodes
##       parameters: 
##           test statistic: fisher
##           cutOff: 0.01
```

```
## 
##   Level 11:  1 nodes to be scored    (0 eliminated genes)
```

```
## 
##   Level 10:  3 nodes to be scored    (3 eliminated genes)
```

```
## 
##   Level 9:   6 nodes to be scored    (3 eliminated genes)
```

```
## 
##   Level 8:   12 nodes to be scored   (3 eliminated genes)
```

```
## 
##   Level 7:   14 nodes to be scored   (8 eliminated genes)
```

```
## 
##   Level 6:   16 nodes to be scored   (8 eliminated genes)
```

```
## 
##   Level 5:   16 nodes to be scored   (101 eliminated genes)
```

```
## 
##   Level 4:   17 nodes to be scored   (101 eliminated genes)
```

```
## 
##   Level 3:   16 nodes to be scored   (101 eliminated genes)
```

```
## 
##   Level 2:   2 nodes to be scored    (101 eliminated genes)
```

```
## 
##   Level 1:   1 nodes to be scored    (101 eliminated genes)
```

```
#extract the significant GO terms
up1hLFCless0.585_BP_elim <- GenTable(GOdata_BP_up1hLFCless0.585, classic = resultFisher_BP_up1hLFCless0.585_elim, 
                            orderBy = "weight", ranksOf = "weight", topNodes = 50)
up1hLFCless0.585_MF_elim <- GenTable(GOdata_MF_up1hLFCless0.585, classic = resultFisher_MF_up1hLFCless0.585_elim, 
                            orderBy = "weight", ranksOf = "weight", topNodes = 50)
up1hLFCless0.585_CC_elim <- GenTable(GOdata_CC_up1hLFCless0.585, classic = resultFisher_CC_up1hLFCless0.585_elim, 
                            orderBy = "weight", ranksOf = "weight", topNodes = 50)
dn1hLFCless0.585_BP_elim <- GenTable(GOdata_BP_dn1hLFCless0.585, classic = resultFisher_BP_dn1hLFCless0.585_elim, 
                              orderBy = "weight", ranksOf = "weight", topNodes = 50)
dn1hLFCless0.585_MF_elim <- GenTable(GOdata_MF_dn1hLFCless0.585, classic = resultFisher_MF_dn1hLFCless0.585_elim, 
                              orderBy = "weight", ranksOf = "weight", topNodes = 50)
dn1hLFCless0.585_CC_elim <- GenTable(GOdata_CC_dn1hLFCless0.585, classic = resultFisher_CC_dn1hLFCless0.585_elim, 
                              orderBy = "weight", ranksOf = "weight", topNodes = 50)

#write.table(up1hLFCless0.585_BP_elim,
#            "/My Drive/ShortTermStress-IlluminaData/FullExp/Cryp-DE/data/ReviGo_files/BP_up0.585_elim.txt")
#write.table(up1hLFCless0.585_MF_elim,
#            "/My Drive/ShortTermStress-IlluminaData/FullExp/Cryp-DE/data/ReviGo_files/MF_up0.585_elim.txt")
#write.table(up1hLFCless0.585_CC_elim,
#            "/My Drive/ShortTermStress-IlluminaData/FullExp/Cryp-DE/data/ReviGo_files/CC_up0.585_elim.txt")
#write.table(dn1hLFCless0.585_BP_elim,
#            "/My Drive/ShortTermStress-IlluminaData/FullExp/Cryp-DE/data/ReviGo_files/BP_dn0.585_elim.txt")
#write.table(dn1hLFCless0.585_MF_elim,
#            "/My Drive/ShortTermStress-IlluminaData/FullExp/Cryp-DE/data/ReviGo_files/MF_dn0.585_elim.txt")
#write.table(dn1hLFCless0.585_CC_elim,
#            "/My Drive/ShortTermStress-IlluminaData/FullExp/Cryp-DE/data/ReviGo_files/CC_dn0.585_elim.txt")

#Not used in further anayses, so we didn't import it back into the code after revigo.
```

Enrichment analyses on timepoints 1 h and 10 h for genes with an LFC
< -1 and > 1

```
##This is for looking at LFC 1 in 1 h and 10 h time points
LFCup1_0v1h <- subset(sign2LFC_0v1h, (`0v1h_Padj`) <= 0.01 & (`0v1h`) > 1)
LFCdn1_0v1h <- subset(sign2LFC_0v1h, (`0v1h_Padj`) <= 0.01 & (`0v1h`) < -1)

LFCup1_0v1h_GOEnrich  = rownames(subset(LFCup1_0v1h, rownames(LFCup1_0v1h)%in%genesOfInterest_allSign))
LFCdn1_0v1h_GOEnrich  = rownames(subset(LFCdn1_0v1h, rownames(LFCdn1_0v1h)%in%genesOfInterest_allSign))


#create gene list for input in topGO for cluster 1
geneList_up1hLFC1 = factor(as.integer(geneUniverse %in% LFCup1_0v1h_GOEnrich))
names(geneList_up1hLFC1) = geneUniverse
str(geneList_up1hLFC1)
```

```
##  Factor w/ 2 levels "0","1": 1 1 1 1 1 1 1 1 1 1 ...
##  - attr(*, "names")= chr [1:6309] "gene_idGO.ID" "CCRYP_013698" "CCRYP_013706" "CCRYP_013704" ...
```

```
geneList_dn1hLFC1 = factor(as.integer(geneUniverse %in% LFCdn1_0v1h_GOEnrich))
names(geneList_dn1hLFC1) = geneUniverse
str(geneList_dn1hLFC1)
```

```
##  Factor w/ 2 levels "0","1": 1 1 1 1 1 1 1 1 1 2 ...
##  - attr(*, "names")= chr [1:6309] "gene_idGO.ID" "CCRYP_013698" "CCRYP_013706" "CCRYP_013704" ...
```

```
##create a topGO object (for up GOs)
GOdata_BP_up1hLFC1 = new("topGOdata", ontology="BP", allGenes=geneList_up1hLFC1, 
                        annot = annFUN.gene2GO, gene2GO = geneID2GO)
```

```
## 
## Building most specific GOs .....
```

```
##  ( 624 GO terms found. )
```

```
## 
## Build GO DAG topology ..........
```

```
##  ( 1729 GO terms and 3515 relations. )
```

```
## 
## Annotating nodes ...............
```

```
##  ( 3345 genes annotated to the GO terms. )
```

```
GOdata_MF_up1hLFC1 = new("topGOdata", ontology="MF", allGenes=geneList_up1hLFC1, 
                        annot = annFUN.gene2GO, gene2GO = geneID2GO)
```

```
## 
## Building most specific GOs .....
```

```
##  ( 736 GO terms found. )
```

```
## 
## Build GO DAG topology ..........
```

```
##  ( 1096 GO terms and 1406 relations. )
```

```
## 
## Annotating nodes ...............
```

```
##  ( 5189 genes annotated to the GO terms. )
```

```
GOdata_CC_up1hLFC1 = new("topGOdata", ontology="CC", allGenes=geneList_up1hLFC1, 
                        annot = annFUN.gene2GO, gene2GO = geneID2GO)
```

```
## 
## Building most specific GOs .....
```

```
##  ( 235 GO terms found. )
```

```
## 
## Build GO DAG topology ..........
```

```
##  ( 452 GO terms and 812 relations. )
```

```
## 
## Annotating nodes ...............
```

```
##  ( 1987 genes annotated to the GO terms. )
```

```
##create a topGO object (for down GOs)
GOdata_BP_dn1hLFC1  = new("topGOdata", ontology="BP", allGenes=geneList_dn1hLFC1 , 
                           annot = annFUN.gene2GO, gene2GO = geneID2GO)
```

```
## 
## Building most specific GOs .....
```

```
##  ( 624 GO terms found. )
```

```
## 
## Build GO DAG topology ..........
```

```
##  ( 1729 GO terms and 3515 relations. )
```

```
## 
## Annotating nodes ...............
```

```
##  ( 3345 genes annotated to the GO terms. )
```

```
GOdata_MF_dn1hLFC1  = new("topGOdata", ontology="MF", allGenes=geneList_dn1hLFC1 , 
                           annot = annFUN.gene2GO, gene2GO = geneID2GO)
```

```
## 
## Building most specific GOs .....
```

```
##  ( 736 GO terms found. )
```

```
## 
## Build GO DAG topology ..........
```

```
##  ( 1096 GO terms and 1406 relations. )
```

```
## 
## Annotating nodes ...............
```

```
##  ( 5189 genes annotated to the GO terms. )
```

```
GOdata_CC_dn1hLFC1  = new("topGOdata", ontology="CC", allGenes=geneList_dn1hLFC1 , 
                           annot = annFUN.gene2GO, gene2GO = geneID2GO)
```

```
## 
## Building most specific GOs .....
```

```
##  ( 235 GO terms found. )
```

```
## 
## Build GO DAG topology ..........
```

```
##  ( 452 GO terms and 812 relations. )
```

```
## 
## Annotating nodes ...............
```

```
##  ( 1987 genes annotated to the GO terms. )
```

```
#run Fisher's exact test
resultFisher_BP_up1hLFC1_elim <- runTest(GOdata_BP_up1hLFC1, algorithm = "elim", statistic = "fisher")
```

```
## 
##           -- Elim Algorithm -- 
## 
##       the algorithm is scoring 196 nontrivial nodes
##       parameters: 
##           test statistic: fisher
##           cutOff: 0.01
```

```
## 
##   Level 13:  1 nodes to be scored    (0 eliminated genes)
```

```
## 
##   Level 12:  1 nodes to be scored    (0 eliminated genes)
```

```
## 
##   Level 11:  3 nodes to be scored    (0 eliminated genes)
```

```
## 
##   Level 10:  7 nodes to be scored    (0 eliminated genes)
```

```
## 
##   Level 9:   16 nodes to be scored   (0 eliminated genes)
```

```
## 
##   Level 8:   22 nodes to be scored   (3 eliminated genes)
```

```
## 
##   Level 7:   20 nodes to be scored   (3 eliminated genes)
```

```
## 
##   Level 6:   34 nodes to be scored   (3 eliminated genes)
```

```
## 
##   Level 5:   40 nodes to be scored   (329 eliminated genes)
```

```
## 
##   Level 4:   25 nodes to be scored   (329 eliminated genes)
```

```
## 
##   Level 3:   19 nodes to be scored   (329 eliminated genes)
```

```
## 
##   Level 2:   7 nodes to be scored    (329 eliminated genes)
```

```
## 
##   Level 1:   1 nodes to be scored    (329 eliminated genes)
```

```
resultFisher_MF_up1hLFC1_elim <- runTest(GOdata_MF_up1hLFC1, algorithm = "elim", statistic = "fisher")
```

```
## 
##           -- Elim Algorithm -- 
## 
##       the algorithm is scoring 134 nontrivial nodes
##       parameters: 
##           test statistic: fisher
##           cutOff: 0.01
```

```
## 
##   Level 9:   2 nodes to be scored    (0 eliminated genes)
```

```
## 
##   Level 8:   7 nodes to be scored    (0 eliminated genes)
```

```
## 
##   Level 7:   13 nodes to be scored   (0 eliminated genes)
```

```
## 
##   Level 6:   25 nodes to be scored   (3 eliminated genes)
```

```
## 
##   Level 5:   28 nodes to be scored   (128 eliminated genes)
```

```
## 
##   Level 4:   34 nodes to be scored   (128 eliminated genes)
```

```
## 
##   Level 3:   17 nodes to be scored   (212 eliminated genes)
```

```
## 
##   Level 2:   7 nodes to be scored    (248 eliminated genes)
```

```
## 
##   Level 1:   1 nodes to be scored    (248 eliminated genes)
```

```
resultFisher_CC_up1hLFC1_elim <- runTest(GOdata_CC_up1hLFC1, algorithm = "elim", statistic = "fisher")
```

```
## 
##           -- Elim Algorithm -- 
## 
##       the algorithm is scoring 56 nontrivial nodes
##       parameters: 
##           test statistic: fisher
##           cutOff: 0.01
```

```
## 
##   Level 10:  1 nodes to be scored    (0 eliminated genes)
```

```
## 
##   Level 9:   1 nodes to be scored    (0 eliminated genes)
```

```
## 
##   Level 8:   5 nodes to be scored    (0 eliminated genes)
```

```
## 
##   Level 7:   6 nodes to be scored    (0 eliminated genes)
```

```
## 
##   Level 6:   7 nodes to be scored    (0 eliminated genes)
```

```
## 
##   Level 5:   8 nodes to be scored    (0 eliminated genes)
```

```
## 
##   Level 4:   14 nodes to be scored   (0 eliminated genes)
```

```
## 
##   Level 3:   11 nodes to be scored   (0 eliminated genes)
```

```
## 
##   Level 2:   2 nodes to be scored    (0 eliminated genes)
```

```
## 
##   Level 1:   1 nodes to be scored    (0 eliminated genes)
```

```
resultFisher_BP_dn1hLFC1_elim <- runTest(GOdata_BP_dn1hLFC1, algorithm = "elim", statistic = "fisher")
```

```
## 
##           -- Elim Algorithm -- 
## 
##       the algorithm is scoring 714 nontrivial nodes
##       parameters: 
##           test statistic: fisher
##           cutOff: 0.01
```

```
## 
##   Level 14:  3 nodes to be scored    (0 eliminated genes)
```

```
## 
##   Level 13:  9 nodes to be scored    (0 eliminated genes)
```

```
## 
##   Level 12:  29 nodes to be scored   (34 eliminated genes)
```

```
## 
##   Level 11:  38 nodes to be scored   (34 eliminated genes)
```

```
## 
##   Level 10:  70 nodes to be scored   (73 eliminated genes)
```

```
## 
##   Level 9:   93 nodes to be scored   (138 eliminated genes)
```

```
## 
##   Level 8:   94 nodes to be scored   (156 eliminated genes)
```

```
## 
##   Level 7:   98 nodes to be scored   (278 eliminated genes)
```

```
## 
##   Level 6:   103 nodes to be scored  (278 eliminated genes)
```

```
## 
##   Level 5:   94 nodes to be scored   (409 eliminated genes)
```

```
## 
##   Level 4:   46 nodes to be scored   (409 eliminated genes)
```

```
## 
##   Level 3:   29 nodes to be scored   (409 eliminated genes)
```

```
## 
##   Level 2:   7 nodes to be scored    (447 eliminated genes)
```

```
## 
##   Level 1:   1 nodes to be scored    (447 eliminated genes)
```

```
resultFisher_MF_dn1hLFC1_elim <- runTest(GOdata_MF_dn1hLFC1, algorithm = "elim", statistic = "fisher")
```

```
## 
##           -- Elim Algorithm -- 
## 
##       the algorithm is scoring 376 nontrivial nodes
##       parameters: 
##           test statistic: fisher
##           cutOff: 0.01
```

```
## 
##   Level 10:  2 nodes to be scored    (0 eliminated genes)
```

```
## 
##   Level 9:   7 nodes to be scored    (0 eliminated genes)
```

```
## 
##   Level 8:   17 nodes to be scored   (109 eliminated genes)
```

```
## 
##   Level 7:   60 nodes to be scored   (109 eliminated genes)
```

```
## 
##   Level 6:   99 nodes to be scored   (141 eliminated genes)
```

```
## 
##   Level 5:   89 nodes to be scored   (222 eliminated genes)
```

```
## 
##   Level 4:   63 nodes to be scored   (993 eliminated genes)
```

```
## 
##   Level 3:   28 nodes to be scored   (1001 eliminated genes)
```

```
## 
##   Level 2:   10 nodes to be scored   (1361 eliminated genes)
```

```
## 
##   Level 1:   1 nodes to be scored    (1361 eliminated genes)
```

```
resultFisher_CC_dn1hLFC1_elim <- runTest(GOdata_CC_dn1hLFC1, algorithm = "elim", statistic = "fisher")
```

```
## 
##           -- Elim Algorithm -- 
## 
##       the algorithm is scoring 187 nontrivial nodes
##       parameters: 
##           test statistic: fisher
##           cutOff: 0.01
```

```
## 
##   Level 12:  2 nodes to be scored    (0 eliminated genes)
```

```
## 
##   Level 11:  6 nodes to be scored    (6 eliminated genes)
```

```
## 
##   Level 10:  8 nodes to be scored    (6 eliminated genes)
```

```
## 
##   Level 9:   19 nodes to be scored   (6 eliminated genes)
```

```
## 
##   Level 8:   27 nodes to be scored   (10 eliminated genes)
```

```
## 
##   Level 7:   29 nodes to be scored   (10 eliminated genes)
```

```
## 
##   Level 6:   29 nodes to be scored   (10 eliminated genes)
```

```
## 
##   Level 5:   22 nodes to be scored   (150 eliminated genes)
```

```
## 
##   Level 4:   22 nodes to be scored   (160 eliminated genes)
```

```
## 
##   Level 3:   20 nodes to be scored   (160 eliminated genes)
```

```
## 
##   Level 2:   2 nodes to be scored    (1157 eliminated genes)
```

```
## 
##   Level 1:   1 nodes to be scored    (1157 eliminated genes)
```

```
#extract the significant GO terms
up1hLFC1_BP_elim <- GenTable(GOdata_BP_up1hLFC1, classic = resultFisher_BP_up1hLFC1_elim, 
                            orderBy = "weight", ranksOf = "weight", topNodes = 50)
up1hLFC1_MF_elim <- GenTable(GOdata_MF_up1hLFC1, classic = resultFisher_MF_up1hLFC1_elim, 
                            orderBy = "weight", ranksOf = "weight", topNodes = 50)
up1hLFC1_CC_elim <- GenTable(GOdata_CC_up1hLFC1, classic = resultFisher_CC_up1hLFC1_elim, 
                            orderBy = "weight", ranksOf = "weight", topNodes = 50)
dn1hLFC1_BP_elim <- GenTable(GOdata_BP_dn1hLFC1, classic = resultFisher_BP_dn1hLFC1_elim, 
                              orderBy = "weight", ranksOf = "weight", topNodes = 50)
dn1hLFC1_MF_elim <- GenTable(GOdata_MF_dn1hLFC1, classic = resultFisher_MF_dn1hLFC1_elim, 
                              orderBy = "weight", ranksOf = "weight", topNodes = 50)
dn1hLFC1_CC_elim <- GenTable(GOdata_CC_dn1hLFC1, classic = resultFisher_CC_dn1hLFC1_elim, 
                              orderBy = "weight", ranksOf = "weight", topNodes = 50)

##Time 0 vs 10 hour
LFCup1_0v10h <- subset(sign2LFC_0v10h, (`0v10h_Padj`) <= 0.01 & (`0v10h`) > 1)
LFCdn1_0v10h <- subset(sign2LFC_0v10h, (`0v10h_Padj`) <= 0.01 & (`0v10h`) < -1)

LFCup1_0v10h_GOEnrich  = rownames(subset(LFCup1_0v10h, rownames(LFCup1_0v10h)%in%genesOfInterest_allSign))
LFCdn1_0v10h_GOEnrich  = rownames(subset(LFCdn1_0v10h, rownames(LFCdn1_0v10h)%in%genesOfInterest_allSign))


#create gene list for input in topGO for cluster 1
geneList_up10hLFC1 = factor(as.integer(geneUniverse %in% LFCup1_0v10h_GOEnrich))
names(geneList_up10hLFC1) = geneUniverse
str(geneList_up10hLFC1)
```

```
##  Factor w/ 2 levels "0","1": 1 1 1 1 1 1 1 1 1 1 ...
##  - attr(*, "names")= chr [1:6309] "gene_idGO.ID" "CCRYP_013698" "CCRYP_013706" "CCRYP_013704" ...
```

```
geneList_dn10hLFC1 = factor(as.integer(geneUniverse %in% LFCdn1_0v10h_GOEnrich))
names(geneList_dn10hLFC1) = geneUniverse
str(geneList_dn10hLFC1)
```

```
##  Factor w/ 2 levels "0","1": 1 1 1 1 1 1 1 1 1 2 ...
##  - attr(*, "names")= chr [1:6309] "gene_idGO.ID" "CCRYP_013698" "CCRYP_013706" "CCRYP_013704" ...
```

```
##create a topGO object (for up GOs)
GOdata_BP_up10hLFC1 = new("topGOdata", ontology="BP", allGenes=geneList_up10hLFC1, 
                        annot = annFUN.gene2GO, gene2GO = geneID2GO)
```

```
## 
## Building most specific GOs .....
```

```
##  ( 624 GO terms found. )
```

```
## 
## Build GO DAG topology ..........
```

```
##  ( 1729 GO terms and 3515 relations. )
```

```
## 
## Annotating nodes ...............
```

```
##  ( 3345 genes annotated to the GO terms. )
```

```
GOdata_MF_up10hLFC1 = new("topGOdata", ontology="MF", allGenes=geneList_up10hLFC1, 
                        annot = annFUN.gene2GO, gene2GO = geneID2GO)
```

```
## 
## Building most specific GOs .....
```

```
##  ( 736 GO terms found. )
```

```
## 
## Build GO DAG topology ..........
```

```
##  ( 1096 GO terms and 1406 relations. )
```

```
## 
## Annotating nodes ...............
```

```
##  ( 5189 genes annotated to the GO terms. )
```

```
GOdata_CC_up10hLFC1 = new("topGOdata", ontology="CC", allGenes=geneList_up10hLFC1, 
                        annot = annFUN.gene2GO, gene2GO = geneID2GO)
```

```
## 
## Building most specific GOs .....
```

```
##  ( 235 GO terms found. )
```

```
## 
## Build GO DAG topology ..........
```

```
##  ( 452 GO terms and 812 relations. )
```

```
## 
## Annotating nodes ...............
```

```
##  ( 1987 genes annotated to the GO terms. )
```

```
##create a topGO object (for down GOs)
GOdata_BP_dn10hLFC1  = new("topGOdata", ontology="BP", allGenes=geneList_dn10hLFC1 , 
                           annot = annFUN.gene2GO, gene2GO = geneID2GO)
```

```
## 
## Building most specific GOs .....
```

```
##  ( 624 GO terms found. )
```

```
## 
## Build GO DAG topology ..........
```

```
##  ( 1729 GO terms and 3515 relations. )
```

```
## 
## Annotating nodes ...............
```

```
##  ( 3345 genes annotated to the GO terms. )
```

```
GOdata_MF_dn10hLFC1  = new("topGOdata", ontology="MF", allGenes=geneList_dn10hLFC1 , 
                           annot = annFUN.gene2GO, gene2GO = geneID2GO)
```

```
## 
## Building most specific GOs .....
```

```
##  ( 736 GO terms found. )
```

```
## 
## Build GO DAG topology ..........
```

```
##  ( 1096 GO terms and 1406 relations. )
```

```
## 
## Annotating nodes ...............
```

```
##  ( 5189 genes annotated to the GO terms. )
```

```
GOdata_CC_dn10hLFC1  = new("topGOdata", ontology="CC", allGenes=geneList_dn10hLFC1 , 
                           annot = annFUN.gene2GO, gene2GO = geneID2GO)
```

```
## 
## Building most specific GOs .....
```

```
##  ( 235 GO terms found. )
```

```
## 
## Build GO DAG topology ..........
```

```
##  ( 452 GO terms and 812 relations. )
```

```
## 
## Annotating nodes ...............
```

```
##  ( 1987 genes annotated to the GO terms. )
```

```
#run Fisher's exact test
resultFisher_BP_up10hLFC1_elim <- runTest(GOdata_BP_up10hLFC1, algorithm = "elim", statistic = "fisher")
```

```
## 
##           -- Elim Algorithm -- 
## 
##       the algorithm is scoring 154 nontrivial nodes
##       parameters: 
##           test statistic: fisher
##           cutOff: 0.01
```

```
## 
##   Level 11:  4 nodes to be scored    (0 eliminated genes)
```

```
## 
##   Level 10:  6 nodes to be scored    (1 eliminated genes)
```

```
## 
##   Level 9:   12 nodes to be scored   (1 eliminated genes)
```

```
## 
##   Level 8:   16 nodes to be scored   (1 eliminated genes)
```

```
## 
##   Level 7:   19 nodes to be scored   (1 eliminated genes)
```

```
## 
##   Level 6:   26 nodes to be scored   (1 eliminated genes)
```

```
## 
##   Level 5:   31 nodes to be scored   (52 eliminated genes)
```

```
## 
##   Level 4:   21 nodes to be scored   (52 eliminated genes)
```

```
## 
##   Level 3:   13 nodes to be scored   (52 eliminated genes)
```

```
## 
##   Level 2:   5 nodes to be scored    (52 eliminated genes)
```

```
## 
##   Level 1:   1 nodes to be scored    (52 eliminated genes)
```

```
resultFisher_MF_up10hLFC1_elim <- runTest(GOdata_MF_up10hLFC1, algorithm = "elim", statistic = "fisher")
```

```
## 
##           -- Elim Algorithm -- 
## 
##       the algorithm is scoring 118 nontrivial nodes
##       parameters: 
##           test statistic: fisher
##           cutOff: 0.01
```

```
## 
##   Level 9:   3 nodes to be scored    (0 eliminated genes)
```

```
## 
##   Level 8:   4 nodes to be scored    (0 eliminated genes)
```

```
## 
##   Level 7:   11 nodes to be scored   (0 eliminated genes)
```

```
## 
##   Level 6:   18 nodes to be scored   (768 eliminated genes)
```

```
## 
##   Level 5:   29 nodes to be scored   (768 eliminated genes)
```

```
## 
##   Level 4:   28 nodes to be scored   (768 eliminated genes)
```

```
## 
##   Level 3:   18 nodes to be scored   (775 eliminated genes)
```

```
## 
##   Level 2:   6 nodes to be scored    (775 eliminated genes)
```

```
## 
##   Level 1:   1 nodes to be scored    (775 eliminated genes)
```

```
resultFisher_CC_up10hLFC1_elim <- runTest(GOdata_CC_up10hLFC1, algorithm = "elim", statistic = "fisher")
```

```
## 
##           -- Elim Algorithm -- 
## 
##       the algorithm is scoring 29 nontrivial nodes
##       parameters: 
##           test statistic: fisher
##           cutOff: 0.01
```

```
## 
##   Level 8:   2 nodes to be scored    (0 eliminated genes)
```

```
## 
##   Level 7:   2 nodes to be scored    (14 eliminated genes)
```

```
## 
##   Level 6:   2 nodes to be scored    (14 eliminated genes)
```

```
## 
##   Level 5:   6 nodes to be scored    (14 eliminated genes)
```

```
## 
##   Level 4:   7 nodes to be scored    (14 eliminated genes)
```

```
## 
##   Level 3:   7 nodes to be scored    (14 eliminated genes)
```

```
## 
##   Level 2:   2 nodes to be scored    (55 eliminated genes)
```

```
## 
##   Level 1:   1 nodes to be scored    (55 eliminated genes)
```

```
resultFisher_BP_dn10hLFC1_elim <- runTest(GOdata_BP_dn10hLFC1, algorithm = "elim", statistic = "fisher")
```

```
## 
##           -- Elim Algorithm -- 
## 
##       the algorithm is scoring 137 nontrivial nodes
##       parameters: 
##           test statistic: fisher
##           cutOff: 0.01
```

```
## 
##   Level 11:  1 nodes to be scored    (0 eliminated genes)
```

```
## 
##   Level 10:  3 nodes to be scored    (0 eliminated genes)
```

```
## 
##   Level 9:   5 nodes to be scored    (0 eliminated genes)
```

```
## 
##   Level 8:   9 nodes to be scored    (0 eliminated genes)
```

```
## 
##   Level 7:   11 nodes to be scored   (1 eliminated genes)
```

```
## 
##   Level 6:   20 nodes to be scored   (1 eliminated genes)
```

```
## 
##   Level 5:   37 nodes to be scored   (1 eliminated genes)
```

```
## 
##   Level 4:   28 nodes to be scored   (1 eliminated genes)
```

```
## 
##   Level 3:   17 nodes to be scored   (1 eliminated genes)
```

```
## 
##   Level 2:   5 nodes to be scored    (1 eliminated genes)
```

```
## 
##   Level 1:   1 nodes to be scored    (1 eliminated genes)
```

```
resultFisher_MF_dn10hLFC1_elim <- runTest(GOdata_MF_dn10hLFC1, algorithm = "elim", statistic = "fisher")
```

```
## 
##           -- Elim Algorithm -- 
## 
##       the algorithm is scoring 113 nontrivial nodes
##       parameters: 
##           test statistic: fisher
##           cutOff: 0.01
```

```
## 
##   Level 9:   1 nodes to be scored    (0 eliminated genes)
```

```
## 
##   Level 8:   3 nodes to be scored    (0 eliminated genes)
```

```
## 
##   Level 7:   9 nodes to be scored    (0 eliminated genes)
```

```
## 
##   Level 6:   21 nodes to be scored   (0 eliminated genes)
```

```
## 
##   Level 5:   25 nodes to be scored   (0 eliminated genes)
```

```
## 
##   Level 4:   29 nodes to be scored   (0 eliminated genes)
```

```
## 
##   Level 3:   18 nodes to be scored   (0 eliminated genes)
```

```
## 
##   Level 2:   6 nodes to be scored    (0 eliminated genes)
```

```
## 
##   Level 1:   1 nodes to be scored    (0 eliminated genes)
```

```
resultFisher_CC_dn10hLFC1_elim <- runTest(GOdata_CC_dn10hLFC1, algorithm = "elim", statistic = "fisher")
```

```
## 
##           -- Elim Algorithm -- 
## 
##       the algorithm is scoring 59 nontrivial nodes
##       parameters: 
##           test statistic: fisher
##           cutOff: 0.01
```

```
## 
##   Level 12:  1 nodes to be scored    (0 eliminated genes)
```

```
## 
##   Level 11:  1 nodes to be scored    (0 eliminated genes)
```

```
## 
##   Level 10:  2 nodes to be scored    (0 eliminated genes)
```

```
## 
##   Level 9:   3 nodes to be scored    (1 eliminated genes)
```

```
## 
##   Level 8:   6 nodes to be scored    (1 eliminated genes)
```

```
## 
##   Level 7:   6 nodes to be scored    (1 eliminated genes)
```

```
## 
##   Level 6:   7 nodes to be scored    (1 eliminated genes)
```

```
## 
##   Level 5:   9 nodes to be scored    (1 eliminated genes)
```

```
## 
##   Level 4:   11 nodes to be scored   (1 eliminated genes)
```

```
## 
##   Level 3:   10 nodes to be scored   (1 eliminated genes)
```

```
## 
##   Level 2:   2 nodes to be scored    (1 eliminated genes)
```

```
## 
##   Level 1:   1 nodes to be scored    (1 eliminated genes)
```

```
#extract the significant GO terms
up10hLFC1_BP_elim <- GenTable(GOdata_BP_up10hLFC1, classic = resultFisher_BP_up10hLFC1_elim, 
                            orderBy = "weight", ranksOf = "weight", topNodes = 50)
up10hLFC1_MF_elim <- GenTable(GOdata_MF_up10hLFC1, classic = resultFisher_MF_up10hLFC1_elim, 
                            orderBy = "weight", ranksOf = "weight", topNodes = 50)
up10hLFC1_CC_elim <- GenTable(GOdata_CC_up10hLFC1, classic = resultFisher_CC_up10hLFC1_elim, 
                            orderBy = "weight", ranksOf = "weight", topNodes = 50)
dn10hLFC1_BP_elim <- GenTable(GOdata_BP_dn10hLFC1, classic = resultFisher_BP_dn10hLFC1_elim, 
                              orderBy = "weight", ranksOf = "weight", topNodes = 50)
dn10hLFC1_MF_elim <- GenTable(GOdata_MF_dn10hLFC1, classic = resultFisher_MF_dn10hLFC1_elim, 
                              orderBy = "weight", ranksOf = "weight", topNodes = 50)
dn10hLFC1_CC_elim <- GenTable(GOdata_CC_dn10hLFC1, classic = resultFisher_CC_dn10hLFC1_elim, 
                              orderBy = "weight", ranksOf = "weight", topNodes = 50)

#write.table(up1hLFC1_BP_elim,
#            "/My Drive/ShortTermStress-IlluminaData/FullExp/Cryp-DE/data/ReviGo_files/BP_up1hLFC1_elim.txt")
#write.table(up1hLFC1_MF_elim,
#            "/My Drive/ShortTermStress-IlluminaData/FullExp/Cryp-DE/data/ReviGo_files/MF_up1hLFC1_elim.txt")
#write.table(up1hLFC1_CC_elim,
#            "/My Drive/ShortTermStress-IlluminaData/FullExp/Cryp-DE/data/ReviGo_files/CC_up1hLFC1_elim.txt")
#write.table(dn1hLFC1_BP_elim,
#            "/My Drive/ShortTermStress-IlluminaData/FullExp/Cryp-DE/data/ReviGo_files/BP_dn1hLFC1_elim.txt")
#write.table(dn1hLFC1_MF_elim,
#            "/My Drive/ShortTermStress-IlluminaData/FullExp/Cryp-DE/data/ReviGo_files/MF_dn1hLFC1_elim.txt")
#write.table(dn1hLFC1_CC_elim,
#            "/My Drive/ShortTermStress-IlluminaData/FullExp/Cryp-DE/data/ReviGo_files/CC_dn1hLFC1_elim.txt")

#write.table(up10hLFC1_BP_elim,
#            "/My Drive/ShortTermStress-IlluminaData/FullExp/Cryp-DE/data/ReviGo_files/BP_up10hLFC1_elim.txt")
#write.table(up10hLFC1_MF_elim,
#            "/My Drive/ShortTermStress-IlluminaData/FullExp/Cryp-DE/data/ReviGo_files/MF_up10hLFC1_elim.txt")
#write.table(up10hLFC1_CC_elim,
#            "/My Drive/ShortTermStress-IlluminaData/FullExp/Cryp-DE/data/ReviGo_files/CC_up10hLFC1_elim.txt")
#write.table(dn10hLFC1_BP_elim,
#            "/My Drive/ShortTermStress-IlluminaData/FullExp/Cryp-DE/data/ReviGo_files/BP_dn10hLFC1_elim.txt")
#write.table(dn10hLFC1_MF_elim,
#            "/My Drive/ShortTermStress-IlluminaData/FullExp/Cryp-DE/data/ReviGo_files/MF_dn10hLFC1_elim.txt")
#write.table(dn10hLFC1_CC_elim,
#            "/My Drive/ShortTermStress-IlluminaData/FullExp/Cryp-DE/data/ReviGo_files/CC_dn10hLFC1_elim.txt")
```

Import the data back in from ReviGO

```
dr_1hLFC1_bp = read.table("/My Drive/ShortTermStress-IlluminaData/FullExp/Cryp-DE/data/ReviGo_files/BP_dn1hLFC1.tsv",
                     colClasses = c(NA,"NULL","NULL","NULL","NULL","NULL","NULL","NULL","NULL","NULL","NULL","NULL","NULL",NA),
                     header=TRUE) #biological process results
ur_1hLFC1_bp = read.table("/My Drive/ShortTermStress-IlluminaData/FullExp/Cryp-DE/data/ReviGo_files/BP_up1hLFC1.tsv",
                     colClasses = c(NA,"NULL","NULL","NULL","NULL","NULL","NULL","NULL","NULL","NULL","NULL","NULL","NULL",NA),
                     header=TRUE) #biological process results
dr_1hLFC1_mf = read.table("/My Drive/ShortTermStress-IlluminaData/FullExp/Cryp-DE/data/ReviGo_files/MF_dn1hLFC1.tsv",
                     colClasses = c(NA,"NULL","NULL","NULL","NULL","NULL","NULL","NULL","NULL","NULL","NULL","NULL","NULL",NA),
                     header=TRUE) #molecular function results
ur_1hLFC1_mf = read.table("/My Drive/ShortTermStress-IlluminaData/FullExp/Cryp-DE/data/ReviGo_files/MF_up1hLFC1.tsv",
                     colClasses = c(NA,"NULL","NULL","NULL","NULL","NULL","NULL","NULL","NULL","NULL","NULL","NULL","NULL",NA),
                     header=TRUE) #molecular function results

dr_10hLFC1_bp = read.table("/My Drive/ShortTermStress-IlluminaData/FullExp/Cryp-DE/data/ReviGo_files/BP_dn10hLFC1.tsv",
                     colClasses = c(NA,"NULL","NULL","NULL","NULL","NULL","NULL","NULL","NULL","NULL","NULL","NULL","NULL",NA),
                     header=TRUE) #biological process results
ur_10hLFC1_bp = read.table("/My Drive/ShortTermStress-IlluminaData/FullExp/Cryp-DE/data/ReviGo_files/BP_up10hLFC1.tsv",
                     colClasses = c(NA,"NULL","NULL","NULL","NULL","NULL","NULL","NULL","NULL","NULL","NULL","NULL","NULL",NA),
                     header=TRUE) #biological process results
dr_10hLFC1_mf = read.table("/My Drive/ShortTermStress-IlluminaData/FullExp/Cryp-DE/data/ReviGo_files/MF_dn10hLFC1.tsv",
                     colClasses = c(NA,"NULL","NULL","NULL","NULL","NULL","NULL","NULL","NULL","NULL","NULL","NULL","NULL",NA),
                     header=TRUE) #molecular function results
ur_10hLFC1_mf = read.table("/My Drive/ShortTermStress-IlluminaData/FullExp/Cryp-DE/data/ReviGo_files/MF_up10hLFC1.tsv",
                     colClasses = c(NA,"NULL","NULL","NULL","NULL","NULL","NULL","NULL","NULL","NULL","NULL","NULL","NULL",NA),
                     header=TRUE) #molecular function results

#
names(dr_1hLFC1_bp) <- c('GO.ID','Eliminated')
names(dr_1hLFC1_mf) <- c('GO.ID','Eliminated')
names(ur_1hLFC1_bp) <- c('GO.ID','Eliminated')
names(ur_1hLFC1_mf) <- c('GO.ID','Eliminated')

names(dr_10hLFC1_bp) <- c('GO.ID','Eliminated')
names(dr_10hLFC1_mf) <- c('GO.ID','Eliminated')
names(ur_10hLFC1_bp) <- c('GO.ID','Eliminated')
names(ur_10hLFC1_mf) <- c('GO.ID','Eliminated')
#

#resave(dr_1hLFC1_bp,dr_1hLFC1_mf,ur_1hLFC1_bp,ur_1hLFC1_mf,
#       dr_10hLFC1_bp,dr_10hLFC1_mf,ur_10hLFC1_bp,ur_10hLFC1_mf,
#       file = "/My Drive/ShortTermStress-IlluminaData/FullExp/Cryp-DE/data/import_rmd_vars.RData")
```

Make a table for quick comparison of the genes that meet the LFC
cutoff for 1 h and 10 h

```
ur_10hLFC1_bp <- up10hLFC1_BP_elim %>% 
  full_join(., ur_10hLFC1_bp, by = "GO.ID", copy = FALSE, suffix = c(".x",".y"))
ur_10hLFC1_bp$timept <- '10h'
ur_10hLFC1_bp$category <- 'biological process'
ur_10hLFC1_bp$degree <- 'upregulated'
ur_10hLFC1_mf <- up10hLFC1_MF_elim %>% 
  full_join(., ur_10hLFC1_mf, by = "GO.ID", copy = FALSE, suffix = c(".x",".y"))
ur_10hLFC1_mf$timept <- '10h'
ur_10hLFC1_mf$category <- 'molecular function'
ur_10hLFC1_mf$degree <- 'upregulated'
dr_10hLFC1_bp <- dn10hLFC1_BP_elim %>% 
  full_join(., dr_10hLFC1_bp, by = "GO.ID", copy = FALSE, suffix = c(".x",".y"))
dr_10hLFC1_bp$timept <- '10h'
dr_10hLFC1_bp$category <- 'biological process'
dr_10hLFC1_bp$degree <- 'downregulated'
dr_10hLFC1_mf <- dn10hLFC1_MF_elim %>% 
  full_join(., dr_10hLFC1_mf, by = "GO.ID", copy = FALSE, suffix = c(".x",".y"))
dr_10hLFC1_mf$timept <- '10h'
dr_10hLFC1_mf$category <- 'molecular function'
dr_10hLFC1_mf$degree <- 'downregulated'

ur_1hLFC1_bp <- up1hLFC1_BP_elim %>% 
  full_join(., ur_1hLFC1_bp, by = "GO.ID", copy = FALSE, suffix = c(".x",".y"))
ur_1hLFC1_bp$timept <- '1h'
ur_1hLFC1_bp$category <- 'biological process'
ur_1hLFC1_bp$degree <- 'upregulated'
ur_1hLFC1_mf <- up1hLFC1_MF_elim %>% 
  full_join(., ur_1hLFC1_mf, by = "GO.ID", copy = FALSE, suffix = c(".x",".y"))
ur_1hLFC1_mf$timept <- '1h'
ur_1hLFC1_mf$category <- 'molecular function'
ur_1hLFC1_mf$degree <- 'upregulated'
dr_1hLFC1_bp <- dn1hLFC1_BP_elim %>% 
  full_join(., dr_1hLFC1_bp, by = "GO.ID", copy = FALSE, suffix = c(".x",".y"))
dr_1hLFC1_bp$timept <- '1h'
dr_1hLFC1_bp$category <- 'biological process'
dr_1hLFC1_bp$degree <- 'downregulated'
dr_1hLFC1_mf <- dn1hLFC1_MF_elim %>% 
  full_join(., dr_1hLFC1_mf, by = "GO.ID", copy = FALSE, suffix = c(".x",".y"))
dr_1hLFC1_mf$timept <- '1h'
dr_1hLFC1_mf$category <- 'molecular function'
dr_1hLFC1_mf$degree <- 'downregulated'

go_data_LFC1 <- rbind(ur_10hLFC1_bp,ur_10hLFC1_mf,dr_10hLFC1_bp,dr_10hLFC1_mf)

names(go_data_LFC1) <- c("GO.ID","Term","Annotated","Sign","Exp","Pvalue","Eliminated","timept","category","degree")
go_data_LFC1$GO.Term <- paste(go_data_LFC1$Term, go_data_LFC1$GO.ID, sep=" ")
go_data_LFC1_reduced <- filter(go_data_LFC1, Eliminated == " False") 
go_data_LFC1_reduced <- subset(go_data_LFC1_reduced,select=c("GO.Term","Annotated","Pvalue","timept","category","degree"))
go_data_LFC1_reduced <- transform(go_data_LFC1_reduced, Percentage = (Annotated / 3193) * 100)
go_data_LFC1_reduced <- subset(go_data_LFC1_reduced, (`Percentage`) > 0.5 & (`Percentage`) < 23) 
go_data_LFC1_reduced <- subset(go_data_LFC1_reduced, (`Pvalue`) < 0.05 |(`Pvalue`) > 0.06) #Necessary to remove > 0.05 but keep e- data
go_data_LFC1_reduced <- go_data_LFC1_reduced %>%
  mutate(Percentage = ifelse(degree == "upregulated",
                             Percentage,
                             -1*Percentage))
```

#### 4.5 Clusters identified from TreeView (centroid linkage)

Cluster 1

```
#Cluster 1 
clust1_genes = read.delim("/My Drive/ShortTermStress-IlluminaData/FullExp/Cryp-DE/data/Clusters_11-19/sign2LFC_Holm_C1.txt",header = TRUE,row.names = 1)
clust1_genes = clust1_genes[-1, ]

clust1_GOEnrich = rownames(clust1_genes)

#create gene list for input in topGO for cluster 1
geneList_clust1 = factor(as.integer(geneUniverse %in% clust1_GOEnrich))
names(geneList_clust1) = geneUniverse
str(geneList_clust1)
```

```
##  Factor w/ 2 levels "0","1": 1 1 1 1 1 1 1 1 1 1 ...
##  - attr(*, "names")= chr [1:6309] "gene_idGO.ID" "CCRYP_013698" "CCRYP_013706" "CCRYP_013704" ...
```

```
##create a topGO object cluster1
GOdata_BP_clust1 = new("topGOdata", ontology="BP", allGenes=geneList_clust1, 
                           annot = annFUN.gene2GO, gene2GO = geneID2GO)
```

```
## 
## Building most specific GOs .....
```

```
##  ( 624 GO terms found. )
```

```
## 
## Build GO DAG topology ..........
```

```
##  ( 1729 GO terms and 3515 relations. )
```

```
## 
## Annotating nodes ...............
```

```
##  ( 3345 genes annotated to the GO terms. )
```

```
GOdata_MF_clust1 = new("topGOdata", ontology="MF", allGenes=geneList_clust1, 
                           annot = annFUN.gene2GO, gene2GO = geneID2GO)
```

```
## 
## Building most specific GOs .....
```

```
##  ( 736 GO terms found. )
```

```
## 
## Build GO DAG topology ..........
```

```
##  ( 1096 GO terms and 1406 relations. )
```

```
## 
## Annotating nodes ...............
```

```
##  ( 5189 genes annotated to the GO terms. )
```

```
GOdata_CC_clust1 = new("topGOdata", ontology="CC", allGenes=geneList_clust1, 
                           annot = annFUN.gene2GO, gene2GO = geneID2GO)
```

```
## 
## Building most specific GOs .....
```

```
##  ( 235 GO terms found. )
```

```
## 
## Build GO DAG topology ..........
```

```
##  ( 452 GO terms and 812 relations. )
```

```
## 
## Annotating nodes ...............
```

```
##  ( 1987 genes annotated to the GO terms. )
```

```
#run Fisher's exact test
resultFisher_BP_clust1_elim <- runTest(GOdata_BP_clust1, algorithm = "elim", statistic = "fisher")
```

```
## 
##           -- Elim Algorithm -- 
## 
##       the algorithm is scoring 247 nontrivial nodes
##       parameters: 
##           test statistic: fisher
##           cutOff: 0.01
```

```
## 
##   Level 13:  1 nodes to be scored    (0 eliminated genes)
```

```
## 
##   Level 12:  3 nodes to be scored    (0 eliminated genes)
```

```
## 
##   Level 11:  11 nodes to be scored   (0 eliminated genes)
```

```
## 
##   Level 10:  14 nodes to be scored   (201 eliminated genes)
```

```
## 
##   Level 9:   25 nodes to be scored   (232 eliminated genes)
```

```
## 
##   Level 8:   29 nodes to be scored   (239 eliminated genes)
```

```
## 
##   Level 7:   25 nodes to be scored   (239 eliminated genes)
```

```
## 
##   Level 6:   36 nodes to be scored   (239 eliminated genes)
```

```
## 
##   Level 5:   49 nodes to be scored   (239 eliminated genes)
```

```
## 
##   Level 4:   31 nodes to be scored   (239 eliminated genes)
```

```
## 
##   Level 3:   18 nodes to be scored   (239 eliminated genes)
```

```
## 
##   Level 2:   4 nodes to be scored    (239 eliminated genes)
```

```
## 
##   Level 1:   1 nodes to be scored    (239 eliminated genes)
```

```
resultFisher_MF_clust1_elim <- runTest(GOdata_MF_clust1, algorithm = "elim", statistic = "fisher")
```

```
## 
##           -- Elim Algorithm -- 
## 
##       the algorithm is scoring 188 nontrivial nodes
##       parameters: 
##           test statistic: fisher
##           cutOff: 0.01
```

```
## 
##   Level 9:   4 nodes to be scored    (0 eliminated genes)
```

```
## 
##   Level 8:   7 nodes to be scored    (0 eliminated genes)
```

```
## 
##   Level 7:   21 nodes to be scored   (0 eliminated genes)
```

```
## 
##   Level 6:   41 nodes to be scored   (0 eliminated genes)
```

```
## 
##   Level 5:   40 nodes to be scored   (5 eliminated genes)
```

```
## 
##   Level 4:   39 nodes to be scored   (5 eliminated genes)
```

```
## 
##   Level 3:   24 nodes to be scored   (5 eliminated genes)
```

```
## 
##   Level 2:   11 nodes to be scored   (133 eliminated genes)
```

```
## 
##   Level 1:   1 nodes to be scored    (133 eliminated genes)
```

```
resultFisher_CC_clust1_elim <- runTest(GOdata_CC_clust1, algorithm = "elim", statistic = "fisher")
```

```
## 
##           -- Elim Algorithm -- 
## 
##       the algorithm is scoring 95 nontrivial nodes
##       parameters: 
##           test statistic: fisher
##           cutOff: 0.01
```

```
## 
##   Level 13:  1 nodes to be scored    (0 eliminated genes)
```

```
## 
##   Level 12:  1 nodes to be scored    (0 eliminated genes)
```

```
## 
##   Level 11:  2 nodes to be scored    (0 eliminated genes)
```

```
## 
##   Level 10:  4 nodes to be scored    (0 eliminated genes)
```

```
## 
##   Level 9:   6 nodes to be scored    (0 eliminated genes)
```

```
## 
##   Level 8:   9 nodes to be scored    (0 eliminated genes)
```

```
## 
##   Level 7:   11 nodes to be scored   (0 eliminated genes)
```

```
## 
##   Level 6:   14 nodes to be scored   (0 eliminated genes)
```

```
## 
##   Level 5:   15 nodes to be scored   (351 eliminated genes)
```

```
## 
##   Level 4:   15 nodes to be scored   (351 eliminated genes)
```

```
## 
##   Level 3:   14 nodes to be scored   (351 eliminated genes)
```

```
## 
##   Level 2:   2 nodes to be scored    (351 eliminated genes)
```

```
## 
##   Level 1:   1 nodes to be scored    (351 eliminated genes)
```

```
#extract the significant GO terms
clust1_BP_elim <- GenTable(GOdata_BP_clust1, classic = resultFisher_BP_clust1_elim, 
                                        orderBy = "weight", ranksOf = "weight", topNodes = 50)
clust1_MF_elim <- GenTable(GOdata_MF_clust1, classic = resultFisher_MF_clust1_elim, 
                                        orderBy = "weight", ranksOf = "weight", topNodes = 50)
clust1_CC_elim <- GenTable(GOdata_CC_clust1, classic = resultFisher_CC_clust1_elim, 
                                        orderBy = "weight", ranksOf = "weight", topNodes = 50)

#files for REVIGO input
#write.table(clust1_BP_elim,
#            "/My Drive/ShortTermStress-IlluminaData/FullExp/Cryp-DE/data/ReviGo_files/Clusters/clust1_BP_elim.txt")
#write.table(clust1_MF_elim,
#            "/My Drive/ShortTermStress-IlluminaData/FullExp/Cryp-DE/data/ReviGo_files/Clusters/clust1_MF_elim.txt")
#write.table(clust1_CC_elim,
#            "/My Drive/ShortTermStress-IlluminaData/FullExp/Cryp-DE/data/ReviGo_files/Clusters/clust1_CC_elim.txt")
```

Cluster 2

```
clust2_genes = read.delim("/My Drive/ShortTermStress-IlluminaData/FullExp/Cryp-DE/data/Clusters_11-19/sign2LFC_Holm_C2.txt",header = TRUE,row.names = 1)
clust2_genes = clust2_genes[-1, ]

clust2_GOEnrich = rownames(clust2_genes)

#create gene list for input in topGO for cluster 1
geneList_clust2 = factor(as.integer(geneUniverse %in% clust2_GOEnrich))
names(geneList_clust2) = geneUniverse


##create a topGO object cluster1
GOdata_BP_clust2 = new("topGOdata", ontology="BP", allGenes=geneList_clust2, 
                           annot = annFUN.gene2GO, gene2GO = geneID2GO)
```

```
## 
## Building most specific GOs .....
```

```
##  ( 624 GO terms found. )
```

```
## 
## Build GO DAG topology ..........
```

```
##  ( 1729 GO terms and 3515 relations. )
```

```
## 
## Annotating nodes ...............
```

```
##  ( 3345 genes annotated to the GO terms. )
```

```
GOdata_MF_clust2 = new("topGOdata", ontology="MF", allGenes=geneList_clust2, 
                           annot = annFUN.gene2GO, gene2GO = geneID2GO)
```

```
## 
## Building most specific GOs .....
```

```
##  ( 736 GO terms found. )
```

```
## 
## Build GO DAG topology ..........
```

```
##  ( 1096 GO terms and 1406 relations. )
```

```
## 
## Annotating nodes ...............
```

```
##  ( 5189 genes annotated to the GO terms. )
```

```
GOdata_CC_clust2 = new("topGOdata", ontology="CC", allGenes=geneList_clust2, 
                           annot = annFUN.gene2GO, gene2GO = geneID2GO)
```

```
## 
## Building most specific GOs .....
```

```
##  ( 235 GO terms found. )
```

```
## 
## Build GO DAG topology ..........
```

```
##  ( 452 GO terms and 812 relations. )
```

```
## 
## Annotating nodes ...............
```

```
##  ( 1987 genes annotated to the GO terms. )
```

```
#run Fisher's exact test
resultFisher_BP_clust2_elim <- runTest(GOdata_BP_clust2, algorithm = "elim", statistic = "fisher")
```

```
## 
##           -- Elim Algorithm -- 
## 
##       the algorithm is scoring 454 nontrivial nodes
##       parameters: 
##           test statistic: fisher
##           cutOff: 0.01
```

```
## 
##   Level 14:  1 nodes to be scored    (0 eliminated genes)
```

```
## 
##   Level 13:  3 nodes to be scored    (0 eliminated genes)
```

```
## 
##   Level 12:  10 nodes to be scored   (0 eliminated genes)
```

```
## 
##   Level 11:  18 nodes to be scored   (0 eliminated genes)
```

```
## 
##   Level 10:  28 nodes to be scored   (0 eliminated genes)
```

```
## 
##   Level 9:   46 nodes to be scored   (31 eliminated genes)
```

```
## 
##   Level 8:   57 nodes to be scored   (31 eliminated genes)
```

```
## 
##   Level 7:   62 nodes to be scored   (43 eliminated genes)
```

```
## 
##   Level 6:   81 nodes to be scored   (113 eliminated genes)
```

```
## 
##   Level 5:   73 nodes to be scored   (154 eliminated genes)
```

```
## 
##   Level 4:   42 nodes to be scored   (154 eliminated genes)
```

```
## 
##   Level 3:   26 nodes to be scored   (154 eliminated genes)
```

```
## 
##   Level 2:   6 nodes to be scored    (154 eliminated genes)
```

```
## 
##   Level 1:   1 nodes to be scored    (154 eliminated genes)
```

```
resultFisher_MF_clust2_elim <- runTest(GOdata_MF_clust2, algorithm = "elim", statistic = "fisher")
```

```
## 
##           -- Elim Algorithm -- 
## 
##       the algorithm is scoring 355 nontrivial nodes
##       parameters: 
##           test statistic: fisher
##           cutOff: 0.01
```

```
## 
##   Level 11:  1 nodes to be scored    (0 eliminated genes)
```

```
## 
##   Level 10:  4 nodes to be scored    (0 eliminated genes)
```

```
## 
##   Level 9:   14 nodes to be scored   (0 eliminated genes)
```

```
## 
##   Level 8:   27 nodes to be scored   (0 eliminated genes)
```

```
## 
##   Level 7:   43 nodes to be scored   (12 eliminated genes)
```

```
## 
##   Level 6:   77 nodes to be scored   (24 eliminated genes)
```

```
## 
##   Level 5:   73 nodes to be scored   (45 eliminated genes)
```

```
## 
##   Level 4:   73 nodes to be scored   (135 eliminated genes)
```

```
## 
##   Level 3:   30 nodes to be scored   (176 eliminated genes)
```

```
## 
##   Level 2:   12 nodes to be scored   (1269 eliminated genes)
```

```
## 
##   Level 1:   1 nodes to be scored    (1269 eliminated genes)
```

```
resultFisher_CC_clust2_elim <- runTest(GOdata_CC_clust2, algorithm = "elim", statistic = "fisher")
```

```
## 
##           -- Elim Algorithm -- 
## 
##       the algorithm is scoring 103 nontrivial nodes
##       parameters: 
##           test statistic: fisher
##           cutOff: 0.01
```

```
## 
##   Level 11:  2 nodes to be scored    (0 eliminated genes)
```

```
## 
##   Level 10:  4 nodes to be scored    (0 eliminated genes)
```

```
## 
##   Level 9:   6 nodes to be scored    (0 eliminated genes)
```

```
## 
##   Level 8:   10 nodes to be scored   (0 eliminated genes)
```

```
## 
##   Level 7:   18 nodes to be scored   (0 eliminated genes)
```

```
## 
##   Level 6:   19 nodes to be scored   (7 eliminated genes)
```

```
## 
##   Level 5:   14 nodes to be scored   (7 eliminated genes)
```

```
## 
##   Level 4:   15 nodes to be scored   (478 eliminated genes)
```

```
## 
##   Level 3:   12 nodes to be scored   (478 eliminated genes)
```

```
## 
##   Level 2:   2 nodes to be scored    (844 eliminated genes)
```

```
## 
##   Level 1:   1 nodes to be scored    (844 eliminated genes)
```

```
#extract the significant GO terms
clust2_BP_elim <- GenTable(GOdata_BP_clust2, classic = resultFisher_BP_clust2_elim, 
                                        orderBy = "weight", ranksOf = "weight", topNodes = 50)
clust2_MF_elim <- GenTable(GOdata_MF_clust2, classic = resultFisher_MF_clust2_elim, 
                                        orderBy = "weight", ranksOf = "weight", topNodes = 50)
clust2_CC_elim <- GenTable(GOdata_CC_clust2, classic = resultFisher_CC_clust2_elim, 
                                        orderBy = "weight", ranksOf = "weight", topNodes = 50)


#files for REVIGO input2.
#write.table(clust2_BP_elim,
#            "/My Drive/ShortTermStress-IlluminaData/FullExp/Cryp-DE/data/ReviGo_files/Clusters/clust2_BP_elim.txt")
#write.table(clust2_MF_elim,
#            "/My Drive/ShortTermStress-IlluminaData/FullExp/Cryp-DE/data/ReviGo_files/Clusters/clust2_MF_elim.txt")
#write.table(clust2_CC_elim,
#            "/My Drive/ShortTermStress-IlluminaData/FullExp/Cryp-DE/data/ReviGo_files/Clusters/clust2_CC_elim.txt")
```

Cluster 3

```
clust3_genes = read.delim("/My Drive/ShortTermStress-IlluminaData/FullExp/Cryp-DE/data/Clusters_11-19/sign2LFC_Holm_C3.txt",header = TRUE,row.names = 1)
clust3_genes = clust3_genes[-1, ]

clust3_GOEnrich = rownames(clust3_genes)

#create gene list for input in topGO for cluster 1
geneList_clust3 = factor(as.integer(geneUniverse %in% clust3_GOEnrich))
names(geneList_clust3) = geneUniverse

##create a topGO object cluster1
GOdata_BP_clust3 = new("topGOdata", ontology="BP", allGenes=geneList_clust3, 
                           annot = annFUN.gene2GO, gene2GO = geneID2GO)
```

```
## 
## Building most specific GOs .....
```

```
##  ( 624 GO terms found. )
```

```
## 
## Build GO DAG topology ..........
```

```
##  ( 1729 GO terms and 3515 relations. )
```

```
## 
## Annotating nodes ...............
```

```
##  ( 3345 genes annotated to the GO terms. )
```

```
GOdata_MF_clust3 = new("topGOdata", ontology="MF", allGenes=geneList_clust3, 
                           annot = annFUN.gene2GO, gene2GO = geneID2GO)
```

```
## 
## Building most specific GOs .....
```

```
##  ( 736 GO terms found. )
```

```
## 
## Build GO DAG topology ..........
```

```
##  ( 1096 GO terms and 1406 relations. )
```

```
## 
## Annotating nodes ...............
```

```
##  ( 5189 genes annotated to the GO terms. )
```

```
GOdata_CC_clust3 = new("topGOdata", ontology="CC", allGenes=geneList_clust3, 
                           annot = annFUN.gene2GO, gene2GO = geneID2GO)
```

```
## 
## Building most specific GOs .....
```

```
##  ( 235 GO terms found. )
```

```
## 
## Build GO DAG topology ..........
```

```
##  ( 452 GO terms and 812 relations. )
```

```
## 
## Annotating nodes ...............
```

```
##  ( 1987 genes annotated to the GO terms. )
```

```
#run Fisher's exact test
resultFisher_BP_clust3_elim <- runTest(GOdata_BP_clust3, algorithm = "elim", statistic = "fisher")
```

```
## 
##           -- Elim Algorithm -- 
## 
##       the algorithm is scoring 327 nontrivial nodes
##       parameters: 
##           test statistic: fisher
##           cutOff: 0.01
```

```
## 
##   Level 14:  1 nodes to be scored    (0 eliminated genes)
```

```
## 
##   Level 13:  2 nodes to be scored    (0 eliminated genes)
```

```
## 
##   Level 12:  4 nodes to be scored    (0 eliminated genes)
```

```
## 
##   Level 11:  8 nodes to be scored    (0 eliminated genes)
```

```
## 
##   Level 10:  20 nodes to be scored   (201 eliminated genes)
```

```
## 
##   Level 9:   32 nodes to be scored   (201 eliminated genes)
```

```
## 
##   Level 8:   35 nodes to be scored   (201 eliminated genes)
```

```
## 
##   Level 7:   36 nodes to be scored   (201 eliminated genes)
```

```
## 
##   Level 6:   56 nodes to be scored   (201 eliminated genes)
```

```
## 
##   Level 5:   63 nodes to be scored   (205 eliminated genes)
```

```
## 
##   Level 4:   39 nodes to be scored   (220 eliminated genes)
```

```
## 
##   Level 3:   23 nodes to be scored   (220 eliminated genes)
```

```
## 
##   Level 2:   7 nodes to be scored    (220 eliminated genes)
```

```
## 
##   Level 1:   1 nodes to be scored    (220 eliminated genes)
```

```
resultFisher_MF_clust3_elim <- runTest(GOdata_MF_clust3, algorithm = "elim", statistic = "fisher")
```

```
## 
##           -- Elim Algorithm -- 
## 
##       the algorithm is scoring 204 nontrivial nodes
##       parameters: 
##           test statistic: fisher
##           cutOff: 0.01
```

```
## 
##   Level 10:  1 nodes to be scored    (0 eliminated genes)
```

```
## 
##   Level 9:   4 nodes to be scored    (0 eliminated genes)
```

```
## 
##   Level 8:   14 nodes to be scored   (0 eliminated genes)
```

```
## 
##   Level 7:   27 nodes to be scored   (0 eliminated genes)
```

```
## 
##   Level 6:   36 nodes to be scored   (91 eliminated genes)
```

```
## 
##   Level 5:   47 nodes to be scored   (188 eliminated genes)
```

```
## 
##   Level 4:   45 nodes to be scored   (188 eliminated genes)
```

```
## 
##   Level 3:   21 nodes to be scored   (188 eliminated genes)
```

```
## 
##   Level 2:   8 nodes to be scored    (224 eliminated genes)
```

```
## 
##   Level 1:   1 nodes to be scored    (224 eliminated genes)
```

```
resultFisher_CC_clust3_elim <- runTest(GOdata_CC_clust3, algorithm = "elim", statistic = "fisher")
```

```
## 
##           -- Elim Algorithm -- 
## 
##       the algorithm is scoring 71 nontrivial nodes
##       parameters: 
##           test statistic: fisher
##           cutOff: 0.01
```

```
## 
##   Level 10:  1 nodes to be scored    (0 eliminated genes)
```

```
## 
##   Level 9:   3 nodes to be scored    (0 eliminated genes)
```

```
## 
##   Level 8:   6 nodes to be scored    (0 eliminated genes)
```

```
## 
##   Level 7:   8 nodes to be scored    (0 eliminated genes)
```

```
## 
##   Level 6:   16 nodes to be scored   (0 eliminated genes)
```

```
## 
##   Level 5:   12 nodes to be scored   (351 eliminated genes)
```

```
## 
##   Level 4:   10 nodes to be scored   (351 eliminated genes)
```

```
## 
##   Level 3:   12 nodes to be scored   (351 eliminated genes)
```

```
## 
##   Level 2:   2 nodes to be scored    (351 eliminated genes)
```

```
## 
##   Level 1:   1 nodes to be scored    (351 eliminated genes)
```

```
#extract the significant GO terms
clust3_BP_elim <- GenTable(GOdata_BP_clust3, classic = resultFisher_BP_clust3_elim, 
                                        orderBy = "weight", ranksOf = "weight", topNodes = 50)
clust3_MF_elim <- GenTable(GOdata_MF_clust3, classic = resultFisher_MF_clust3_elim, 
                                        orderBy = "weight", ranksOf = "weight", topNodes = 50)
clust3_CC_elim <- GenTable(GOdata_CC_clust3, classic = resultFisher_CC_clust3_elim, 
                                        orderBy = "weight", ranksOf = "weight", topNodes = 50)

#files for REVIGO input2.
#write.table(clust3_BP_elim,
#            "/My Drive/ShortTermStress-IlluminaData/FullExp/Cryp-DE/data/ReviGo_files/Clusters/clust3_BP_elim.txt")
#write.table(clust3_MF_elim,
#            "/My Drive/ShortTermStress-IlluminaData/FullExp/Cryp-DE/data/ReviGo_files/Clusters/clust3_MF_elim.txt")
#write.table(clust3_CC_elim,
#            "/My Drive/ShortTermStress-IlluminaData/FullExp/Cryp-DE/data/ReviGo_files/Clusters/clust3_CC_elim.txt")
```

Cluster 4

```
clust4_genes = read.delim("/My Drive/ShortTermStress-IlluminaData/FullExp/Cryp-DE/data/Clusters_11-19/sign2LFC_Holm_C4.txt",header = TRUE,row.names = 1)
clust4_genes = clust4_genes[-1, ]

clust4_GOEnrich = rownames(clust4_genes)

#create gene list for input in topGO for cluster 1
geneList_clust4 = factor(as.integer(geneUniverse %in% clust4_GOEnrich))
names(geneList_clust4) = geneUniverse

##create a topGO object cluster1
GOdata_BP_clust4 = new("topGOdata", ontology="BP", allGenes=geneList_clust4, 
                           annot = annFUN.gene2GO, gene2GO = geneID2GO)
```

```
## 
## Building most specific GOs .....
```

```
##  ( 624 GO terms found. )
```

```
## 
## Build GO DAG topology ..........
```

```
##  ( 1729 GO terms and 3515 relations. )
```

```
## 
## Annotating nodes ...............
```

```
##  ( 3345 genes annotated to the GO terms. )
```

```
GOdata_MF_clust4 = new("topGOdata", ontology="MF", allGenes=geneList_clust4, 
                           annot = annFUN.gene2GO, gene2GO = geneID2GO)
```

```
## 
## Building most specific GOs .....
```

```
##  ( 736 GO terms found. )
```

```
## 
## Build GO DAG topology ..........
```

```
##  ( 1096 GO terms and 1406 relations. )
```

```
## 
## Annotating nodes ...............
```

```
##  ( 5189 genes annotated to the GO terms. )
```

```
GOdata_CC_clust4 = new("topGOdata", ontology="CC", allGenes=geneList_clust4, 
                           annot = annFUN.gene2GO, gene2GO = geneID2GO)
```

```
## 
## Building most specific GOs .....
```

```
##  ( 235 GO terms found. )
```

```
## 
## Build GO DAG topology ..........
```

```
##  ( 452 GO terms and 812 relations. )
```

```
## 
## Annotating nodes ...............
```

```
##  ( 1987 genes annotated to the GO terms. )
```

```
#run Fisher's exact test
resultFisher_BP_clust4_elim <- runTest(GOdata_BP_clust4, algorithm = "elim", statistic = "fisher")
```

```
## 
##           -- Elim Algorithm -- 
## 
##       the algorithm is scoring 234 nontrivial nodes
##       parameters: 
##           test statistic: fisher
##           cutOff: 0.01
```

```
## 
##   Level 12:  1 nodes to be scored    (0 eliminated genes)
```

```
## 
##   Level 11:  5 nodes to be scored    (0 eliminated genes)
```

```
## 
##   Level 10:  15 nodes to be scored   (0 eliminated genes)
```

```
## 
##   Level 9:   22 nodes to be scored   (0 eliminated genes)
```

```
## 
##   Level 8:   27 nodes to be scored   (0 eliminated genes)
```

```
## 
##   Level 7:   23 nodes to be scored   (6 eliminated genes)
```

```
## 
##   Level 6:   40 nodes to be scored   (6 eliminated genes)
```

```
## 
##   Level 5:   45 nodes to be scored   (6 eliminated genes)
```

```
## 
##   Level 4:   30 nodes to be scored   (6 eliminated genes)
```

```
## 
##   Level 3:   19 nodes to be scored   (6 eliminated genes)
```

```
## 
##   Level 2:   6 nodes to be scored    (6 eliminated genes)
```

```
## 
##   Level 1:   1 nodes to be scored    (6 eliminated genes)
```

```
resultFisher_MF_clust4_elim <- runTest(GOdata_MF_clust4, algorithm = "elim", statistic = "fisher")
```

```
## 
##           -- Elim Algorithm -- 
## 
##       the algorithm is scoring 194 nontrivial nodes
##       parameters: 
##           test statistic: fisher
##           cutOff: 0.01
```

```
## 
##   Level 11:  1 nodes to be scored    (0 eliminated genes)
```

```
## 
##   Level 10:  1 nodes to be scored    (7 eliminated genes)
```

```
## 
##   Level 9:   7 nodes to be scored    (7 eliminated genes)
```

```
## 
##   Level 8:   11 nodes to be scored   (7 eliminated genes)
```

```
## 
##   Level 7:   20 nodes to be scored   (7 eliminated genes)
```

```
## 
##   Level 6:   40 nodes to be scored   (7 eliminated genes)
```

```
## 
##   Level 5:   45 nodes to be scored   (7 eliminated genes)
```

```
## 
##   Level 4:   44 nodes to be scored   (7 eliminated genes)
```

```
## 
##   Level 3:   18 nodes to be scored   (7 eliminated genes)
```

```
## 
##   Level 2:   6 nodes to be scored    (522 eliminated genes)
```

```
## 
##   Level 1:   1 nodes to be scored    (522 eliminated genes)
```

```
resultFisher_CC_clust4_elim <- runTest(GOdata_CC_clust4, algorithm = "elim", statistic = "fisher")
```

```
## 
##           -- Elim Algorithm -- 
## 
##       the algorithm is scoring 41 nontrivial nodes
##       parameters: 
##           test statistic: fisher
##           cutOff: 0.01
```

```
## 
##   Level 10:  1 nodes to be scored    (0 eliminated genes)
```

```
## 
##   Level 9:   1 nodes to be scored    (0 eliminated genes)
```

```
## 
##   Level 8:   2 nodes to be scored    (0 eliminated genes)
```

```
## 
##   Level 7:   5 nodes to be scored    (0 eliminated genes)
```

```
## 
##   Level 6:   9 nodes to be scored    (0 eliminated genes)
```

```
## 
##   Level 5:   6 nodes to be scored    (0 eliminated genes)
```

```
## 
##   Level 4:   7 nodes to be scored    (471 eliminated genes)
```

```
## 
##   Level 3:   7 nodes to be scored    (471 eliminated genes)
```

```
## 
##   Level 2:   2 nodes to be scored    (471 eliminated genes)
```

```
## 
##   Level 1:   1 nodes to be scored    (471 eliminated genes)
```

```
#extract the significant GO terms
clust4_BP_elim <- GenTable(GOdata_BP_clust4, classic = resultFisher_BP_clust4_elim, 
                                        orderBy = "weight", ranksOf = "weight", topNodes = 50)
clust4_MF_elim <- GenTable(GOdata_MF_clust4, classic = resultFisher_MF_clust4_elim, 
                                        orderBy = "weight", ranksOf = "weight", topNodes = 50)
clust4_CC_elim <- GenTable(GOdata_CC_clust4, classic = resultFisher_CC_clust4_elim, 
                                        orderBy = "weight", ranksOf = "weight", topNodes = 50)

#files for REVIGO input2.
#write.table(clust4_BP_elim,
#            "/My Drive/ShortTermStress-IlluminaData/FullExp/Cryp-DE/data/ReviGo_files/Clusters/clust4_BP_elim.txt")
#write.table(clust4_MF_elim,
#            "/My Drive/ShortTermStress-IlluminaData/FullExp/Cryp-DE/data/ReviGo_files/Clusters/clust4_MF_elim.txt")
#write.table(clust4_CC_elim,
#            "/My Drive/ShortTermStress-IlluminaData/FullExp/Cryp-DE/data/ReviGo_files/Clusters/clust4_CC_elim.txt")
```

Cluster 5

```
clust5_genes = read.delim("/My Drive/ShortTermStress-IlluminaData/FullExp/Cryp-DE/data/Clusters_11-19/sign2LFC_Holm_C5.txt",header = TRUE,row.names = 1)
clust5_genes = clust5_genes[-1, ]

clust5_GOEnrich = rownames(clust5_genes)

#create gene list for input in topGO for cluster 1
geneList_clust5 = factor(as.integer(geneUniverse %in% clust5_GOEnrich))
names(geneList_clust5) = geneUniverse

##create a topGO object cluster1
GOdata_BP_clust5 = new("topGOdata", ontology="BP", allGenes=geneList_clust5, 
                           annot = annFUN.gene2GO, gene2GO = geneID2GO)
```

```
## 
## Building most specific GOs .....
```

```
##  ( 624 GO terms found. )
```

```
## 
## Build GO DAG topology ..........
```

```
##  ( 1729 GO terms and 3515 relations. )
```

```
## 
## Annotating nodes ...............
```

```
##  ( 3345 genes annotated to the GO terms. )
```

```
GOdata_MF_clust5 = new("topGOdata", ontology="MF", allGenes=geneList_clust5, 
                           annot = annFUN.gene2GO, gene2GO = geneID2GO)
```

```
## 
## Building most specific GOs .....
```

```
##  ( 736 GO terms found. )
```

```
## 
## Build GO DAG topology ..........
```

```
##  ( 1096 GO terms and 1406 relations. )
```

```
## 
## Annotating nodes ...............
```

```
##  ( 5189 genes annotated to the GO terms. )
```

```
GOdata_CC_clust5 = new("topGOdata", ontology="CC", allGenes=geneList_clust5, 
                           annot = annFUN.gene2GO, gene2GO = geneID2GO)
```

```
## 
## Building most specific GOs .....
```

```
##  ( 235 GO terms found. )
```

```
## 
## Build GO DAG topology ..........
```

```
##  ( 452 GO terms and 812 relations. )
```

```
## 
## Annotating nodes ...............
```

```
##  ( 1987 genes annotated to the GO terms. )
```

```
#run Fisher's exact test
resultFisher_BP_clust5_elim <- runTest(GOdata_BP_clust5, algorithm = "elim", statistic = "fisher")
```

```
## 
##           -- Elim Algorithm -- 
## 
##       the algorithm is scoring 665 nontrivial nodes
##       parameters: 
##           test statistic: fisher
##           cutOff: 0.01
```

```
## 
##   Level 14:  1 nodes to be scored    (0 eliminated genes)
```

```
## 
##   Level 13:  5 nodes to be scored    (0 eliminated genes)
```

```
## 
##   Level 12:  20 nodes to be scored   (34 eliminated genes)
```

```
## 
##   Level 11:  37 nodes to be scored   (34 eliminated genes)
```

```
## 
##   Level 10:  67 nodes to be scored   (34 eliminated genes)
```

```
## 
##   Level 9:   90 nodes to be scored   (101 eliminated genes)
```

```
## 
##   Level 8:   90 nodes to be scored   (212 eliminated genes)
```

```
## 
##   Level 7:   88 nodes to be scored   (334 eliminated genes)
```

```
## 
##   Level 6:   89 nodes to be scored   (336 eliminated genes)
```

```
## 
##   Level 5:   91 nodes to be scored   (362 eliminated genes)
```

```
## 
##   Level 4:   52 nodes to be scored   (362 eliminated genes)
```

```
## 
##   Level 3:   28 nodes to be scored   (362 eliminated genes)
```

```
## 
##   Level 2:   6 nodes to be scored    (394 eliminated genes)
```

```
## 
##   Level 1:   1 nodes to be scored    (394 eliminated genes)
```

```
resultFisher_MF_clust5_elim <- runTest(GOdata_MF_clust5, algorithm = "elim", statistic = "fisher")
```

```
## 
##           -- Elim Algorithm -- 
## 
##       the algorithm is scoring 377 nontrivial nodes
##       parameters: 
##           test statistic: fisher
##           cutOff: 0.01
```

```
## 
##   Level 10:  3 nodes to be scored    (0 eliminated genes)
```

```
## 
##   Level 9:   13 nodes to be scored   (0 eliminated genes)
```

```
## 
##   Level 8:   20 nodes to be scored   (109 eliminated genes)
```

```
## 
##   Level 7:   47 nodes to be scored   (118 eliminated genes)
```

```
## 
##   Level 6:   93 nodes to be scored   (794 eliminated genes)
```

```
## 
##   Level 5:   91 nodes to be scored   (801 eliminated genes)
```

```
## 
##   Level 4:   69 nodes to be scored   (986 eliminated genes)
```

```
## 
##   Level 3:   29 nodes to be scored   (999 eliminated genes)
```

```
## 
##   Level 2:   11 nodes to be scored   (1132 eliminated genes)
```

```
## 
##   Level 1:   1 nodes to be scored    (1132 eliminated genes)
```

```
resultFisher_CC_clust5_elim <- runTest(GOdata_CC_clust5, algorithm = "elim", statistic = "fisher")
```

```
## 
##           -- Elim Algorithm -- 
## 
##       the algorithm is scoring 174 nontrivial nodes
##       parameters: 
##           test statistic: fisher
##           cutOff: 0.01
```

```
## 
##   Level 12:  1 nodes to be scored    (0 eliminated genes)
```

```
## 
##   Level 11:  6 nodes to be scored    (0 eliminated genes)
```

```
## 
##   Level 10:  9 nodes to be scored    (0 eliminated genes)
```

```
## 
##   Level 9:   14 nodes to be scored   (2 eliminated genes)
```

```
## 
##   Level 8:   26 nodes to be scored   (2 eliminated genes)
```

```
## 
##   Level 7:   23 nodes to be scored   (24 eliminated genes)
```

```
## 
##   Level 6:   28 nodes to be scored   (24 eliminated genes)
```

```
## 
##   Level 5:   23 nodes to be scored   (109 eliminated genes)
```

```
## 
##   Level 4:   20 nodes to be scored   (121 eliminated genes)
```

```
## 
##   Level 3:   21 nodes to be scored   (121 eliminated genes)
```

```
## 
##   Level 2:   2 nodes to be scored    (121 eliminated genes)
```

```
## 
##   Level 1:   1 nodes to be scored    (121 eliminated genes)
```

```
#extract the significant GO terms
clust5_BP_elim <- GenTable(GOdata_BP_clust5, classic = resultFisher_BP_clust5_elim, 
                                        orderBy = "weight", ranksOf = "weight", topNodes = 50)
clust5_MF_elim <- GenTable(GOdata_MF_clust5, classic = resultFisher_MF_clust5_elim, 
                                        orderBy = "weight", ranksOf = "weight", topNodes = 50)
clust5_CC_elim <- GenTable(GOdata_CC_clust5, classic = resultFisher_CC_clust5_elim, 
                                        orderBy = "weight", ranksOf = "weight", topNodes = 50)

#files for REVIGO input.
#write.table(clust5_BP_elim,
#            "/My Drive/ShortTermStress-IlluminaData/FullExp/Cryp-DE/data/ReviGo_files/Clusters/clust5_BP_elim.txt")
#write.table(clust5_MF_elim,
#            "/My Drive/ShortTermStress-IlluminaData/FullExp/Cryp-DE/data/ReviGo_files/Clusters/clust5_MF_elim.txt")
#write.table(clust5_CC_elim,
#            "/My Drive/ShortTermStress-IlluminaData/FullExp/Cryp-DE/data/ReviGo_files/Clusters/clust5_CC_elim.txt")
```

Cluster 6

```
clust6_genes = read.delim("/My Drive/ShortTermStress-IlluminaData/FullExp/Cryp-DE/data/Clusters_11-19/sign2LFC_Holm_C6.txt",header = TRUE,row.names = 1)
clust6_genes = clust6_genes[-1, ]

clust6_GOEnrich = rownames(clust6_genes)

#create gene list for input in topGO for cluster 1
geneList_clust6 = factor(as.integer(geneUniverse %in% clust6_GOEnrich))
names(geneList_clust6) = geneUniverse

##create a topGO object cluster1
GOdata_BP_clust6 = new("topGOdata", ontology="BP", allGenes=geneList_clust6, 
                           annot = annFUN.gene2GO, gene2GO = geneID2GO)
```

```
## 
## Building most specific GOs .....
```

```
##  ( 624 GO terms found. )
```

```
## 
## Build GO DAG topology ..........
```

```
##  ( 1729 GO terms and 3515 relations. )
```

```
## 
## Annotating nodes ...............
```

```
##  ( 3345 genes annotated to the GO terms. )
```

```
GOdata_MF_clust6 = new("topGOdata", ontology="MF", allGenes=geneList_clust6, 
                           annot = annFUN.gene2GO, gene2GO = geneID2GO)
```

```
## 
## Building most specific GOs .....
```

```
##  ( 736 GO terms found. )
```

```
## 
## Build GO DAG topology ..........
```

```
##  ( 1096 GO terms and 1406 relations. )
```

```
## 
## Annotating nodes ...............
```

```
##  ( 5189 genes annotated to the GO terms. )
```

```
GOdata_CC_clust6 = new("topGOdata", ontology="CC", allGenes=geneList_clust6, 
                           annot = annFUN.gene2GO, gene2GO = geneID2GO)
```

```
## 
## Building most specific GOs .....
```

```
##  ( 235 GO terms found. )
```

```
## 
## Build GO DAG topology ..........
```

```
##  ( 452 GO terms and 812 relations. )
```

```
## 
## Annotating nodes ...............
```

```
##  ( 1987 genes annotated to the GO terms. )
```

```
#run Fisher's exact test
resultFisher_BP_clust6_elim <- runTest(GOdata_BP_clust6, algorithm = "elim", statistic = "fisher")
```

```
## 
##           -- Elim Algorithm -- 
## 
##       the algorithm is scoring 359 nontrivial nodes
##       parameters: 
##           test statistic: fisher
##           cutOff: 0.01
```

```
## 
##   Level 14:  1 nodes to be scored    (0 eliminated genes)
```

```
## 
##   Level 13:  2 nodes to be scored    (0 eliminated genes)
```

```
## 
##   Level 12:  9 nodes to be scored    (0 eliminated genes)
```

```
## 
##   Level 11:  14 nodes to be scored   (0 eliminated genes)
```

```
## 
##   Level 10:  27 nodes to be scored   (0 eliminated genes)
```

```
## 
##   Level 9:   35 nodes to be scored   (0 eliminated genes)
```

```
## 
##   Level 8:   48 nodes to be scored   (16 eliminated genes)
```

```
## 
##   Level 7:   41 nodes to be scored   (16 eliminated genes)
```

```
## 
##   Level 6:   55 nodes to be scored   (19 eliminated genes)
```

```
## 
##   Level 5:   60 nodes to be scored   (19 eliminated genes)
```

```
## 
##   Level 4:   37 nodes to be scored   (19 eliminated genes)
```

```
## 
##   Level 3:   23 nodes to be scored   (205 eliminated genes)
```

```
## 
##   Level 2:   6 nodes to be scored    (205 eliminated genes)
```

```
## 
##   Level 1:   1 nodes to be scored    (205 eliminated genes)
```

```
resultFisher_MF_clust6_elim <- runTest(GOdata_MF_clust6, algorithm = "elim", statistic = "fisher")
```

```
## 
##           -- Elim Algorithm -- 
## 
##       the algorithm is scoring 220 nontrivial nodes
##       parameters: 
##           test statistic: fisher
##           cutOff: 0.01
```

```
## 
##   Level 10:  3 nodes to be scored    (0 eliminated genes)
```

```
## 
##   Level 9:   8 nodes to be scored    (0 eliminated genes)
```

```
## 
##   Level 8:   10 nodes to be scored   (0 eliminated genes)
```

```
## 
##   Level 7:   20 nodes to be scored   (0 eliminated genes)
```

```
## 
##   Level 6:   48 nodes to be scored   (109 eliminated genes)
```

```
## 
##   Level 5:   52 nodes to be scored   (197 eliminated genes)
```

```
## 
##   Level 4:   49 nodes to be scored   (199 eliminated genes)
```

```
## 
##   Level 3:   22 nodes to be scored   (213 eliminated genes)
```

```
## 
##   Level 2:   7 nodes to be scored    (621 eliminated genes)
```

```
## 
##   Level 1:   1 nodes to be scored    (621 eliminated genes)
```

```
resultFisher_CC_clust6_elim <- runTest(GOdata_CC_clust6, algorithm = "elim", statistic = "fisher")
```

```
## 
##           -- Elim Algorithm -- 
## 
##       the algorithm is scoring 90 nontrivial nodes
##       parameters: 
##           test statistic: fisher
##           cutOff: 0.01
```

```
## 
##   Level 11:  2 nodes to be scored    (0 eliminated genes)
```

```
## 
##   Level 10:  5 nodes to be scored    (0 eliminated genes)
```

```
## 
##   Level 9:   6 nodes to be scored    (0 eliminated genes)
```

```
## 
##   Level 8:   8 nodes to be scored    (0 eliminated genes)
```

```
## 
##   Level 7:   10 nodes to be scored   (0 eliminated genes)
```

```
## 
##   Level 6:   12 nodes to be scored   (0 eliminated genes)
```

```
## 
##   Level 5:   13 nodes to be scored   (0 eliminated genes)
```

```
## 
##   Level 4:   16 nodes to be scored   (0 eliminated genes)
```

```
## 
##   Level 3:   15 nodes to be scored   (3 eliminated genes)
```

```
## 
##   Level 2:   2 nodes to be scored    (841 eliminated genes)
```

```
## 
##   Level 1:   1 nodes to be scored    (841 eliminated genes)
```

```
#extract the significant GO terms
clust6_BP_elim <- GenTable(GOdata_BP_clust6, classic = resultFisher_BP_clust6_elim, 
                                        orderBy = "weight", ranksOf = "weight", topNodes = 50)
clust6_MF_elim <- GenTable(GOdata_MF_clust6, classic = resultFisher_MF_clust6_elim, 
                                        orderBy = "weight", ranksOf = "weight", topNodes = 50)
clust6_CC_elim <- GenTable(GOdata_CC_clust6, classic = resultFisher_CC_clust6_elim, 
                                        orderBy = "weight", ranksOf = "weight", topNodes = 50)

#files for REVIGO input2.
#write.table(clust6_BP_elim,
#            "/My Drive/ShortTermStress-IlluminaData/FullExp/Cryp-DE/data/ReviGo_files/Clusters/clust6_BP_elim.txt")
#write.table(clust6_MF_elim,
#            "/My Drive/ShortTermStress-IlluminaData/FullExp/Cryp-DE/data/ReviGo_files/Clusters/clust6_MF_elim.txt")
#write.table(clust6_CC_elim,
#            "/My Drive/ShortTermStress-IlluminaData/FullExp/Cryp-DE/data/ReviGo_files/Clusters/clust6_CC_elim.txt")
```

Cluster 7

```
clust7_genes = read.delim("/My Drive/ShortTermStress-IlluminaData/FullExp/Cryp-DE/data/Clusters_11-19/sign2LFC_Holm_C7.txt",header = TRUE,row.names = 1)
clust7_genes = clust7_genes[-1, ]

clust7_GOEnrich = rownames(clust7_genes)

#create gene list for input in topGO for cluster 1
geneList_clust7 = factor(as.integer(geneUniverse %in% clust7_GOEnrich))
names(geneList_clust7) = geneUniverse

##create a topGO object cluster1
GOdata_BP_clust7 = new("topGOdata", ontology="BP", allGenes=geneList_clust7, 
                           annot = annFUN.gene2GO, gene2GO = geneID2GO)
```

```
## 
## Building most specific GOs .....
```

```
##  ( 624 GO terms found. )
```

```
## 
## Build GO DAG topology ..........
```

```
##  ( 1729 GO terms and 3515 relations. )
```

```
## 
## Annotating nodes ...............
```

```
##  ( 3345 genes annotated to the GO terms. )
```

```
GOdata_MF_clust7 = new("topGOdata", ontology="MF", allGenes=geneList_clust7, 
                           annot = annFUN.gene2GO, gene2GO = geneID2GO)
```

```
## 
## Building most specific GOs .....
```

```
##  ( 736 GO terms found. )
```

```
## 
## Build GO DAG topology ..........
```

```
##  ( 1096 GO terms and 1406 relations. )
```

```
## 
## Annotating nodes ...............
```

```
##  ( 5189 genes annotated to the GO terms. )
```

```
GOdata_CC_clust7 = new("topGOdata", ontology="CC", allGenes=geneList_clust7, 
                           annot = annFUN.gene2GO, gene2GO = geneID2GO)
```

```
## 
## Building most specific GOs .....
```

```
##  ( 235 GO terms found. )
```

```
## 
## Build GO DAG topology ..........
```

```
##  ( 452 GO terms and 812 relations. )
```

```
## 
## Annotating nodes ...............
```

```
##  ( 1987 genes annotated to the GO terms. )
```

```
#run Fisher's exact test
resultFisher_BP_clust7_elim <- runTest(GOdata_BP_clust7, algorithm = "elim", statistic = "fisher")
```

```
## 
##           -- Elim Algorithm -- 
## 
##       the algorithm is scoring 487 nontrivial nodes
##       parameters: 
##           test statistic: fisher
##           cutOff: 0.01
```

```
## 
##   Level 15:  1 nodes to be scored    (0 eliminated genes)
```

```
## 
##   Level 14:  3 nodes to be scored    (0 eliminated genes)
```

```
## 
##   Level 13:  4 nodes to be scored    (0 eliminated genes)
```

```
## 
##   Level 12:  6 nodes to be scored    (0 eliminated genes)
```

```
## 
##   Level 11:  16 nodes to be scored   (0 eliminated genes)
```

```
## 
##   Level 10:  30 nodes to be scored   (0 eliminated genes)
```

```
## 
##   Level 9:   43 nodes to be scored   (0 eliminated genes)
```

```
## 
##   Level 8:   55 nodes to be scored   (0 eliminated genes)
```

```
## 
##   Level 7:   77 nodes to be scored   (5 eliminated genes)
```

```
## 
##   Level 6:   80 nodes to be scored   (20 eliminated genes)
```

```
## 
##   Level 5:   87 nodes to be scored   (20 eliminated genes)
```

```
## 
##   Level 4:   47 nodes to be scored   (190 eliminated genes)
```

```
## 
##   Level 3:   30 nodes to be scored   (190 eliminated genes)
```

```
## 
##   Level 2:   7 nodes to be scored    (195 eliminated genes)
```

```
## 
##   Level 1:   1 nodes to be scored    (195 eliminated genes)
```

```
resultFisher_MF_clust7_elim <- runTest(GOdata_MF_clust7, algorithm = "elim", statistic = "fisher")
```

```
## 
##           -- Elim Algorithm -- 
## 
##       the algorithm is scoring 221 nontrivial nodes
##       parameters: 
##           test statistic: fisher
##           cutOff: 0.01
```

```
## 
##   Level 9:   3 nodes to be scored    (0 eliminated genes)
```

```
## 
##   Level 8:   12 nodes to be scored   (0 eliminated genes)
```

```
## 
##   Level 7:   26 nodes to be scored   (0 eliminated genes)
```

```
## 
##   Level 6:   50 nodes to be scored   (0 eliminated genes)
```

```
## 
##   Level 5:   51 nodes to be scored   (53 eliminated genes)
```

```
## 
##   Level 4:   46 nodes to be scored   (53 eliminated genes)
```

```
## 
##   Level 3:   22 nodes to be scored   (53 eliminated genes)
```

```
## 
##   Level 2:   10 nodes to be scored   (53 eliminated genes)
```

```
## 
##   Level 1:   1 nodes to be scored    (53 eliminated genes)
```

```
resultFisher_CC_clust7_elim <- runTest(GOdata_CC_clust7, algorithm = "elim", statistic = "fisher")
```

```
## 
##           -- Elim Algorithm -- 
## 
##       the algorithm is scoring 121 nontrivial nodes
##       parameters: 
##           test statistic: fisher
##           cutOff: 0.01
```

```
## 
##   Level 12:  1 nodes to be scored    (0 eliminated genes)
```

```
## 
##   Level 11:  3 nodes to be scored    (0 eliminated genes)
```

```
## 
##   Level 10:  6 nodes to be scored    (0 eliminated genes)
```

```
## 
##   Level 9:   10 nodes to be scored   (0 eliminated genes)
```

```
## 
##   Level 8:   16 nodes to be scored   (0 eliminated genes)
```

```
## 
##   Level 7:   16 nodes to be scored   (0 eliminated genes)
```

```
## 
##   Level 6:   18 nodes to be scored   (0 eliminated genes)
```

```
## 
##   Level 5:   17 nodes to be scored   (0 eliminated genes)
```

```
## 
##   Level 4:   17 nodes to be scored   (0 eliminated genes)
```

```
## 
##   Level 3:   14 nodes to be scored   (309 eliminated genes)
```

```
## 
##   Level 2:   2 nodes to be scored    (309 eliminated genes)
```

```
## 
##   Level 1:   1 nodes to be scored    (309 eliminated genes)
```

```
#extract the significant GO terms
clust7_BP_elim <- GenTable(GOdata_BP_clust7, classic = resultFisher_BP_clust7_elim, 
                                        orderBy = "weight", ranksOf = "weight", topNodes = 50)
clust7_MF_elim <- GenTable(GOdata_MF_clust7, classic = resultFisher_MF_clust7_elim, 
                                        orderBy = "weight", ranksOf = "weight", topNodes = 50)
clust7_CC_elim <- GenTable(GOdata_CC_clust7, classic = resultFisher_CC_clust7_elim, 
                                        orderBy = "weight", ranksOf = "weight", topNodes = 50)

#files for REVIGO input.
#write.table(clust7_BP_elim,
#            "/My Drive/ShortTermStress-IlluminaData/FullExp/Cryp-DE/data/ReviGo_files/Clusters/clust7_BP_elim.txt")
#write.table(clust7_MF_elim,
#            "/My Drive/ShortTermStress-IlluminaData/FullExp/Cryp-DE/data/ReviGo_files/Clusters/clust7_MF_elim.txt")
#write.table(clust7_CC_elim,
#            "/My Drive/ShortTermStress-IlluminaData/FullExp/Cryp-DE/data/ReviGo_files/Clusters/clust7_CC_elim.txt")
```

Cluster 8

```
clust8_genes = read.delim("/My Drive/ShortTermStress-IlluminaData/FullExp/Cryp-DE/data/Clusters_11-19/sign2LFC_Holm_C8.txt",header = TRUE,row.names = 1)
clust8_genes = clust8_genes[-1, ]

clust8_GOEnrich = rownames(clust8_genes)

#create gene list for input in topGO for cluster 1
geneList_clust8 = factor(as.integer(geneUniverse %in% clust8_GOEnrich))
names(geneList_clust8) = geneUniverse

##create a topGO object cluster1
GOdata_BP_clust8 = new("topGOdata", ontology="BP", allGenes=geneList_clust8, 
                           annot = annFUN.gene2GO, gene2GO = geneID2GO)
```

```
## 
## Building most specific GOs .....
```

```
##  ( 624 GO terms found. )
```

```
## 
## Build GO DAG topology ..........
```

```
##  ( 1729 GO terms and 3515 relations. )
```

```
## 
## Annotating nodes ...............
```

```
##  ( 3345 genes annotated to the GO terms. )
```

```
GOdata_MF_clust8 = new("topGOdata", ontology="MF", allGenes=geneList_clust8, 
                           annot = annFUN.gene2GO, gene2GO = geneID2GO)
```

```
## 
## Building most specific GOs .....
```

```
##  ( 736 GO terms found. )
```

```
## 
## Build GO DAG topology ..........
```

```
##  ( 1096 GO terms and 1406 relations. )
```

```
## 
## Annotating nodes ...............
```

```
##  ( 5189 genes annotated to the GO terms. )
```

```
GOdata_CC_clust8 = new("topGOdata", ontology="CC", allGenes=geneList_clust8, 
                           annot = annFUN.gene2GO, gene2GO = geneID2GO)
```

```
## 
## Building most specific GOs .....
```

```
##  ( 235 GO terms found. )
```

```
## 
## Build GO DAG topology ..........
```

```
##  ( 452 GO terms and 812 relations. )
```

```
## 
## Annotating nodes ...............
```

```
##  ( 1987 genes annotated to the GO terms. )
```

```
#run Fisher's exact test
resultFisher_BP_clust8_elim <- runTest(GOdata_BP_clust8, algorithm = "elim", statistic = "fisher")
```

```
## 
##           -- Elim Algorithm -- 
## 
##       the algorithm is scoring 590 nontrivial nodes
##       parameters: 
##           test statistic: fisher
##           cutOff: 0.01
```

```
## 
##   Level 14:  2 nodes to be scored    (0 eliminated genes)
```

```
## 
##   Level 13:  8 nodes to be scored    (0 eliminated genes)
```

```
## 
##   Level 12:  24 nodes to be scored   (41 eliminated genes)
```

```
## 
##   Level 11:  28 nodes to be scored   (41 eliminated genes)
```

```
## 
##   Level 10:  53 nodes to be scored   (41 eliminated genes)
```

```
## 
##   Level 9:   72 nodes to be scored   (86 eliminated genes)
```

```
## 
##   Level 8:   73 nodes to be scored   (93 eliminated genes)
```

```
## 
##   Level 7:   77 nodes to be scored   (232 eliminated genes)
```

```
## 
##   Level 6:   91 nodes to be scored   (232 eliminated genes)
```

```
## 
##   Level 5:   83 nodes to be scored   (232 eliminated genes)
```

```
## 
##   Level 4:   42 nodes to be scored   (1100 eliminated genes)
```

```
## 
##   Level 3:   30 nodes to be scored   (1100 eliminated genes)
```

```
## 
##   Level 2:   6 nodes to be scored    (1100 eliminated genes)
```

```
## 
##   Level 1:   1 nodes to be scored    (2769 eliminated genes)
```

```
resultFisher_MF_clust8_elim <- runTest(GOdata_MF_clust8, algorithm = "elim", statistic = "fisher")
```

```
## 
##           -- Elim Algorithm -- 
## 
##       the algorithm is scoring 287 nontrivial nodes
##       parameters: 
##           test statistic: fisher
##           cutOff: 0.01
```

```
## 
##   Level 10:  1 nodes to be scored    (0 eliminated genes)
```

```
## 
##   Level 9:   6 nodes to be scored    (0 eliminated genes)
```

```
## 
##   Level 8:   13 nodes to be scored   (109 eliminated genes)
```

```
## 
##   Level 7:   37 nodes to be scored   (140 eliminated genes)
```

```
## 
##   Level 6:   68 nodes to be scored   (155 eliminated genes)
```

```
## 
##   Level 5:   70 nodes to be scored   (207 eliminated genes)
```

```
## 
##   Level 4:   56 nodes to be scored   (383 eliminated genes)
```

```
## 
##   Level 3:   24 nodes to be scored   (398 eliminated genes)
```

```
## 
##   Level 2:   11 nodes to be scored   (398 eliminated genes)
```

```
## 
##   Level 1:   1 nodes to be scored    (506 eliminated genes)
```

```
resultFisher_CC_clust8_elim <- runTest(GOdata_CC_clust8, algorithm = "elim", statistic = "fisher")
```

```
## 
##           -- Elim Algorithm -- 
## 
##       the algorithm is scoring 173 nontrivial nodes
##       parameters: 
##           test statistic: fisher
##           cutOff: 0.01
```

```
## 
##   Level 12:  1 nodes to be scored    (0 eliminated genes)
```

```
## 
##   Level 11:  5 nodes to be scored    (0 eliminated genes)
```

```
## 
##   Level 10:  11 nodes to be scored   (0 eliminated genes)
```

```
## 
##   Level 9:   19 nodes to be scored   (0 eliminated genes)
```

```
## 
##   Level 8:   27 nodes to be scored   (2 eliminated genes)
```

```
## 
##   Level 7:   26 nodes to be scored   (11 eliminated genes)
```

```
## 
##   Level 6:   27 nodes to be scored   (39 eliminated genes)
```

```
## 
##   Level 5:   19 nodes to be scored   (111 eliminated genes)
```

```
## 
##   Level 4:   18 nodes to be scored   (171 eliminated genes)
```

```
## 
##   Level 3:   17 nodes to be scored   (351 eliminated genes)
```

```
## 
##   Level 2:   2 nodes to be scored    (1160 eliminated genes)
```

```
## 
##   Level 1:   1 nodes to be scored    (1160 eliminated genes)
```

```
#extract the significant GO terms
clust8_BP_elim <- GenTable(GOdata_BP_clust8, classic = resultFisher_BP_clust8_elim, 
                                        orderBy = "weight", ranksOf = "weight", topNodes = 50)
clust8_MF_elim <- GenTable(GOdata_MF_clust8, classic = resultFisher_MF_clust8_elim, 
                                        orderBy = "weight", ranksOf = "weight", topNodes = 50)
clust8_CC_elim <- GenTable(GOdata_CC_clust8, classic = resultFisher_CC_clust8_elim, 
                                        orderBy = "weight", ranksOf = "weight", topNodes = 50)

#files for REVIGO input.
#write.table(clust8_BP_elim,
#            "/My Drive/ShortTermStress-IlluminaData/FullExp/Cryp-DE/data/ReviGo_files/Clusters/clust8_BP_elim.txt")
#write.table(clust8_MF_elim,
#            "/My Drive/ShortTermStress-IlluminaData/FullExp/Cryp-DE/data/ReviGo_files/Clusters/clust8_MF_elim.txt")
#write.table(clust8_CC_elim,
#            "/My Drive/ShortTermStress-IlluminaData/FullExp/Cryp-DE/data/ReviGo_files/Clusters/clust8_CC_elim.txt")
```

Cluster 9

```
clust9_genes = read.delim("/My Drive/ShortTermStress-IlluminaData/FullExp/Cryp-DE/data/Clusters_11-19/sign2LFC_Holm_C9.txt",header = TRUE,row.names = 1)
clust9_genes = clust9_genes[-1, ]

clust9_GOEnrich = rownames(clust9_genes)

#create gene list for input in topGO for cluster 1
geneList_clust9 = factor(as.integer(geneUniverse %in% clust9_GOEnrich))
names(geneList_clust9) = geneUniverse

##create a topGO object cluster1
GOdata_BP_clust9 = new("topGOdata", ontology="BP", allGenes=geneList_clust9, 
                           annot = annFUN.gene2GO, gene2GO = geneID2GO)
```

```
## 
## Building most specific GOs .....
```

```
##  ( 624 GO terms found. )
```

```
## 
## Build GO DAG topology ..........
```

```
##  ( 1729 GO terms and 3515 relations. )
```

```
## 
## Annotating nodes ...............
```

```
##  ( 3345 genes annotated to the GO terms. )
```

```
GOdata_MF_clust9 = new("topGOdata", ontology="MF", allGenes=geneList_clust9, 
                           annot = annFUN.gene2GO, gene2GO = geneID2GO)
```

```
## 
## Building most specific GOs .....
```

```
##  ( 736 GO terms found. )
```

```
## 
## Build GO DAG topology ..........
```

```
##  ( 1096 GO terms and 1406 relations. )
```

```
## 
## Annotating nodes ...............
```

```
##  ( 5189 genes annotated to the GO terms. )
```

```
GOdata_CC_clust9 = new("topGOdata", ontology="CC", allGenes=geneList_clust9, 
                           annot = annFUN.gene2GO, gene2GO = geneID2GO)
```

```
## 
## Building most specific GOs .....
```

```
##  ( 235 GO terms found. )
```

```
## 
## Build GO DAG topology ..........
```

```
##  ( 452 GO terms and 812 relations. )
```

```
## 
## Annotating nodes ...............
```

```
##  ( 1987 genes annotated to the GO terms. )
```

```
#run Fisher's exact test
resultFisher_BP_clust9_elim <- runTest(GOdata_BP_clust9, algorithm = "elim", statistic = "fisher")
```

```
## 
##           -- Elim Algorithm -- 
## 
##       the algorithm is scoring 98 nontrivial nodes
##       parameters: 
##           test statistic: fisher
##           cutOff: 0.01
```

```
## 
##   Level 11:  1 nodes to be scored    (0 eliminated genes)
```

```
## 
##   Level 10:  3 nodes to be scored    (0 eliminated genes)
```

```
## 
##   Level 9:   3 nodes to be scored    (0 eliminated genes)
```

```
## 
##   Level 8:   6 nodes to be scored    (0 eliminated genes)
```

```
## 
##   Level 7:   8 nodes to be scored    (8 eliminated genes)
```

```
## 
##   Level 6:   18 nodes to be scored   (32 eliminated genes)
```

```
## 
##   Level 5:   25 nodes to be scored   (32 eliminated genes)
```

```
## 
##   Level 4:   15 nodes to be scored   (32 eliminated genes)
```

```
## 
##   Level 3:   12 nodes to be scored   (185 eliminated genes)
```

```
## 
##   Level 2:   6 nodes to be scored    (185 eliminated genes)
```

```
## 
##   Level 1:   1 nodes to be scored    (185 eliminated genes)
```

```
resultFisher_MF_clust9_elim <- runTest(GOdata_MF_clust9, algorithm = "elim", statistic = "fisher")
```

```
## 
##           -- Elim Algorithm -- 
## 
##       the algorithm is scoring 60 nontrivial nodes
##       parameters: 
##           test statistic: fisher
##           cutOff: 0.01
```

```
## 
##   Level 8:   1 nodes to be scored    (0 eliminated genes)
```

```
## 
##   Level 7:   3 nodes to be scored    (0 eliminated genes)
```

```
## 
##   Level 6:   8 nodes to be scored    (0 eliminated genes)
```

```
## 
##   Level 5:   14 nodes to be scored   (8 eliminated genes)
```

```
## 
##   Level 4:   15 nodes to be scored   (49 eliminated genes)
```

```
## 
##   Level 3:   13 nodes to be scored   (94 eliminated genes)
```

```
## 
##   Level 2:   5 nodes to be scored    (151 eliminated genes)
```

```
## 
##   Level 1:   1 nodes to be scored    (151 eliminated genes)
```

```
resultFisher_CC_clust9_elim <- runTest(GOdata_CC_clust9, algorithm = "elim", statistic = "fisher")
```

```
## 
##           -- Elim Algorithm -- 
## 
##       the algorithm is scoring 18 nontrivial nodes
##       parameters: 
##           test statistic: fisher
##           cutOff: 0.01
```

```
## 
##   Level 7:   1 nodes to be scored    (0 eliminated genes)
```

```
## 
##   Level 6:   3 nodes to be scored    (0 eliminated genes)
```

```
## 
##   Level 5:   3 nodes to be scored    (0 eliminated genes)
```

```
## 
##   Level 4:   4 nodes to be scored    (0 eliminated genes)
```

```
## 
##   Level 3:   5 nodes to be scored    (0 eliminated genes)
```

```
## 
##   Level 2:   1 nodes to be scored    (41 eliminated genes)
```

```
## 
##   Level 1:   1 nodes to be scored    (41 eliminated genes)
```

```
#extract the significant GO terms
clust9_BP_elim <- GenTable(GOdata_BP_clust9, classic = resultFisher_BP_clust9_elim, 
                                        orderBy = "weight", ranksOf = "weight", topNodes = 50)
clust9_MF_elim <- GenTable(GOdata_MF_clust9, classic = resultFisher_MF_clust9_elim, 
                                        orderBy = "weight", ranksOf = "weight", topNodes = 50)
clust9_CC_elim <- GenTable(GOdata_CC_clust9, classic = resultFisher_CC_clust9_elim, 
                                        orderBy = "weight", ranksOf = "weight", topNodes = 50)

#files for REVIGO input2.
#write.table(clust9_BP_elim,
#            "/My Drive/ShortTermStress-IlluminaData/FullExp/Cryp-DE/data/ReviGo_files/Clusters/clust9_BP_elim.txt")
#write.table(clust9_MF_elim,
#            "/My Drive/ShortTermStress-IlluminaData/FullExp/Cryp-DE/data/ReviGo_files/Clusters/clust9_MF_elim.txt")
#write.table(clust9_CC_elim,
#            "/My Drive/ShortTermStress-IlluminaData/FullExp/Cryp-DE/data/ReviGo_files/Clusters/clust9_CC_elim.txt")
```

Cluster 10

```
clust10_genes = read.delim("/My Drive/ShortTermStress-IlluminaData/FullExp/Cryp-DE/data/Clusters_11-19/sign2LFC_Holm_C10.txt",header = TRUE,row.names = 1)
clust10_genes = clust10_genes[-1, ]

clust10_GOEnrich = rownames(clust10_genes)

#create gene list for input in topGO for cluster 1
geneList_clust10 = factor(as.integer(geneUniverse %in% clust10_GOEnrich))
names(geneList_clust10) = geneUniverse

##create a topGO object cluster1
GOdata_BP_clust10 = new("topGOdata", ontology="BP", allGenes=geneList_clust10, 
                           annot = annFUN.gene2GO, gene2GO = geneID2GO)
```

```
## 
## Building most specific GOs .....
```

```
##  ( 624 GO terms found. )
```

```
## 
## Build GO DAG topology ..........
```

```
##  ( 1729 GO terms and 3515 relations. )
```

```
## 
## Annotating nodes ...............
```

```
##  ( 3345 genes annotated to the GO terms. )
```

```
GOdata_MF_clust10 = new("topGOdata", ontology="MF", allGenes=geneList_clust10, 
                           annot = annFUN.gene2GO, gene2GO = geneID2GO)
```

```
## 
## Building most specific GOs .....
```

```
##  ( 736 GO terms found. )
```

```
## 
## Build GO DAG topology ..........
```

```
##  ( 1096 GO terms and 1406 relations. )
```

```
## 
## Annotating nodes ...............
```

```
##  ( 5189 genes annotated to the GO terms. )
```

```
GOdata_CC_clust10 = new("topGOdata", ontology="CC", allGenes=geneList_clust10, 
                           annot = annFUN.gene2GO, gene2GO = geneID2GO)
```

```
## 
## Building most specific GOs .....
```

```
##  ( 235 GO terms found. )
```

```
## 
## Build GO DAG topology ..........
```

```
##  ( 452 GO terms and 812 relations. )
```

```
## 
## Annotating nodes ...............
```

```
##  ( 1987 genes annotated to the GO terms. )
```

```
#run Fisher's exact test
resultFisher_BP_clust10_elim <- runTest(GOdata_BP_clust10, algorithm = "elim", statistic = "fisher")
```

```
## 
##           -- Elim Algorithm -- 
## 
##       the algorithm is scoring 64 nontrivial nodes
##       parameters: 
##           test statistic: fisher
##           cutOff: 0.01
```

```
## 
##   Level 13:  1 nodes to be scored    (0 eliminated genes)
```

```
## 
##   Level 12:  1 nodes to be scored    (0 eliminated genes)
```

```
## 
##   Level 11:  1 nodes to be scored    (0 eliminated genes)
```

```
## 
##   Level 10:  2 nodes to be scored    (0 eliminated genes)
```

```
## 
##   Level 9:   5 nodes to be scored    (0 eliminated genes)
```

```
## 
##   Level 8:   5 nodes to be scored    (0 eliminated genes)
```

```
## 
##   Level 7:   5 nodes to be scored    (0 eliminated genes)
```

```
## 
##   Level 6:   7 nodes to be scored    (0 eliminated genes)
```

```
## 
##   Level 5:   11 nodes to be scored   (41 eliminated genes)
```

```
## 
##   Level 4:   14 nodes to be scored   (41 eliminated genes)
```

```
## 
##   Level 3:   8 nodes to be scored    (41 eliminated genes)
```

```
## 
##   Level 2:   3 nodes to be scored    (41 eliminated genes)
```

```
## 
##   Level 1:   1 nodes to be scored    (41 eliminated genes)
```

```
resultFisher_MF_clust10_elim <- runTest(GOdata_MF_clust10, algorithm = "elim", statistic = "fisher")
```

```
## 
##           -- Elim Algorithm -- 
## 
##       the algorithm is scoring 50 nontrivial nodes
##       parameters: 
##           test statistic: fisher
##           cutOff: 0.01
```

```
## 
##   Level 9:   2 nodes to be scored    (0 eliminated genes)
```

```
## 
##   Level 8:   3 nodes to be scored    (2 eliminated genes)
```

```
## 
##   Level 7:   6 nodes to be scored    (4 eliminated genes)
```

```
## 
##   Level 6:   7 nodes to be scored    (4 eliminated genes)
```

```
## 
##   Level 5:   9 nodes to be scored    (8 eliminated genes)
```

```
## 
##   Level 4:   10 nodes to be scored   (8 eliminated genes)
```

```
## 
##   Level 3:   10 nodes to be scored   (8 eliminated genes)
```

```
## 
##   Level 2:   2 nodes to be scored    (8 eliminated genes)
```

```
## 
##   Level 1:   1 nodes to be scored    (8 eliminated genes)
```

```
resultFisher_CC_clust10_elim <- runTest(GOdata_CC_clust10, algorithm = "elim", statistic = "fisher")
```

```
## 
##           -- Elim Algorithm -- 
## 
##       the algorithm is scoring 29 nontrivial nodes
##       parameters: 
##           test statistic: fisher
##           cutOff: 0.01
```

```
## 
##   Level 10:  1 nodes to be scored    (0 eliminated genes)
```

```
## 
##   Level 9:   1 nodes to be scored    (0 eliminated genes)
```

```
## 
##   Level 8:   2 nodes to be scored    (0 eliminated genes)
```

```
## 
##   Level 7:   2 nodes to be scored    (0 eliminated genes)
```

```
## 
##   Level 6:   3 nodes to be scored    (0 eliminated genes)
```

```
## 
##   Level 5:   5 nodes to be scored    (0 eliminated genes)
```

```
## 
##   Level 4:   6 nodes to be scored    (0 eliminated genes)
```

```
## 
##   Level 3:   6 nodes to be scored    (0 eliminated genes)
```

```
## 
##   Level 2:   2 nodes to be scored    (0 eliminated genes)
```

```
## 
##   Level 1:   1 nodes to be scored    (0 eliminated genes)
```

```
#extract the significant GO terms
clust10_BP_elim <- GenTable(GOdata_BP_clust10, classic = resultFisher_BP_clust10_elim, 
                                        orderBy = "weight", ranksOf = "weight", topNodes = 50)
clust10_MF_elim <- GenTable(GOdata_MF_clust10, classic = resultFisher_MF_clust10_elim, 
                                        orderBy = "weight", ranksOf = "weight", topNodes = 50)
clust10_CC_elim <- GenTable(GOdata_CC_clust10, classic = resultFisher_CC_clust10_elim, 
                                        orderBy = "weight", ranksOf = "weight", topNodes = 50)

#files for REVIGO input
#write.table(clust10_BP_elim,
#            "/My Drive/ShortTermStress-IlluminaData/FullExp/Cryp-DE/data/ReviGo_files/Clusters/clust10_BP_elim.txt")
#write.table(clust10_MF_elim,
#            "/My Drive/ShortTermStress-IlluminaData/FullExp/Cryp-DE/data/ReviGo_files/Clusters/clust10_MF_elim.txt")
#write.table(clust10_CC_elim,
#            "/My Drive/ShortTermStress-IlluminaData/FullExp/Cryp-DE/data/ReviGo_files/Clusters/clust10_CC_elim.txt")
```

### 5. Combine data for KO, GO, uniprot, and cluster information with the LFC data for perusal.

Here we create a couple metadata tables to reference for a wholistic
view of the data

```
kofamKO <- read.delim("/My Drive/ShortTermStress-IlluminaData/FullExp/Cryp-DE/CCMP332_KOtoMerge.txt", header = TRUE, sep = '\t', fill = TRUE, row.names=NULL)
colnames(kofamKO) <- c("gene_id","KO.ID","E_value","KO_definition")
norepeats_KO <- kofamKO %>% distinct(gene_id, .keep_all = TRUE) #Keep only the unique gene_id rows. Kept the first occurrence, which worked out in all the instances I looked at 

GO <- read.delim("/My Drive/ShortTermStress-IlluminaData/FullExp/Cryp-DE/CCMP332_GFF_GO.txt", header = TRUE, sep = '\t', fill = TRUE, row.names=NULL)
colnames(GO) <- c("gene_id","GO.ID")

uniprot_data <- read.delim("/My Drive/ShortTermStress-IlluminaData/FullExp/Cryp-DE/CCMP332_uniprot.csv", header = TRUE, sep = '\t', fill = TRUE, row.names=NULL)
rownames(uniprot_data) <- uniprot_data[,1]
uniprot_data<-  subset(uniprot_data, select=-c(Entry,Gene.names,Organism,KEGG,KO,Gene.ontology.IDs))
colnames(uniprot_data) <- c("protein_name","molecular_function","cellular_component","biological_process")

dmndDB <- read.delim("/My Drive/ShortTermStress-IlluminaData/FullExp/Cryp-DE/CCMP332-diamondbp-matches.m8", header = TRUE, sep = '\t', fill = TRUE)
dmndDB <-  subset(dmndDB, select=-c(pident,length,mismatch,gapopen,qstart,qend,sstart,send))
colnames(dmndDB) <- c("gene_id","UniprotID","E.value","bitscore")

gene_uniprot_anno <- uniprot_data %>%
  tibble::rownames_to_column(.,"UniprotID") %>% #Use full_join to make sure we keep all the expression data and don't limit the GO and KO terms
  full_join(., dmndDB, by = NULL, copy = FALSE, suffix = c(".x",".y"))
gene_uniprot_anno <- gene_uniprot_anno %>% distinct(gene_id, .keep_all = TRUE) #Keep only the unique gene_id rows. Kept the first occurrence, which worked out in all the instances I looked at 

DE_LT <- tibble::rownames_to_column(DE_nakov$table, "gene_id")

paralogs <- read.delim("/My Drive/ShortTermStress-IlluminaData/FullExp/Cryp-DE/data/Cryp_ortho/Results_Feb06/Ortho.txt", header =FALSE, sep = '\t', fill = TRUE, row.names = NULL) #There
#paralogs <- subset(paralogs, select=-c(C_nana))
colnames(paralogs) <- c("orthogroup","gene_id")
paralogs$gene_id <- str_split(paralogs$gene_id, ",") # separate desired column on ,
paralogs <- paralogs %>% unnest(cols = gene_id) # unnest list column to rows
paralogs$gene_id <- paralogs$gene_id %>% str_extract("CCRYP_\\d+")
paralogs <- (na.omit(paralogs))
paralogs_multi <- paralogs %>% distinct(gene_id, .keep_all = TRUE)  %>% group_by(orthogroup) %>% filter( n() > 2 )
paralogs_pairs <- paralogs %>% distinct(gene_id, .keep_all = TRUE)  %>% group_by(orthogroup) %>% filter( n() == 2 )
singletons <- paralogs %>% distinct(gene_id, .keep_all = TRUE)  %>% group_by(orthogroup) %>% filter( n() == 1 )
singletons$cnt_OG <- "1" 
paralogs_pairs$cnt_OG <- "2" 
paralogs_multi$cnt_OG <- "3+" 

clusters <- read.delim("/My Drive/ShortTermStress-IlluminaData/FullExp/Cryp-DE/data/Clusters_11-19/10clusters.txt", header = TRUE, sep = '\t', fill = TRUE)

clusters_GO_KO_2sigdf <- sign2LFC %>% 
  tibble::rownames_to_column(.,"gene_id") %>% #Use full_join to make sure we keep all the expression data and don't limit the GO and KO terms
  left_join(., clusters, by = NULL, copy = FALSE, suffix = c(".x",".y")) %>%
  left_join(., GO, by = NULL, copy = FALSE, suffix = c(".x",".y")) %>%
  left_join(., norepeats_KO, by = NULL, copy = FALSE, suffix = c(".x",".y")) %>%
  left_join(., gene_uniprot_anno, by = NULL, copy = FALSE, suffix = c(".x",".y"))  %>%
  left_join(., DE_LT, by = NULL, copy = FALSE, suffix = c(".x",".y"))
colnames(clusters_GO_KO_2sigdf) <- c("gene_id","t15m","t15m_Padj","t30m","t30m_Padj","t1h","t1h_Padj","t2h","t2h_Padj",
                                    "t4h","t4h_Padj","t8h","t8h_Padj","t10h","t10h_Padj","cluster","GO.ID",
                                    "KO.ID","E-value","KO_definition","UniprotID","protein_name",
                                    "MF","CC","BP","E-value","Bitscore","LT_logFC","LT_CPM","LT_PValue")

clusters_GO_KO_2Sig <- subset(clusters_GO_KO_2sigdf, select=c("gene_id","t15m","t30m","t1h","t2h","t4h","t8h","t10h","LT_logFC","LT_PValue",
                                                  "cluster","UniprotID","protein_name","KO.ID","KO_definition",
                                                  "GO.ID","MF","BP","CC"))

NS_LFCanno_df <- NS_LFC_df %>% 
  tibble::rownames_to_column(.,"gene_id") %>% #Use full_join to make sure we keep all the expression data and don't limit the GO and KO terms
  left_join(., clusters, by = NULL, copy = FALSE, suffix = c(".x",".y")) %>%
  left_join(., GO, by = NULL, copy = FALSE, suffix = c(".x",".y")) %>%
  left_join(., norepeats_KO, by = NULL, copy = FALSE, suffix = c(".x",".y")) %>%
  left_join(., gene_uniprot_anno, by = NULL, copy = FALSE, suffix = c(".x",".y")) %>%
  left_join(., DE_LT, by = NULL, copy = FALSE, suffix = c(".x",".y"))
colnames(NS_LFCanno_df) <- c("gene_id","t15m","t15m_Padj","t30m","t30m_Padj","t1h","t1h_Padj","t2h","t2h_Padj",
                                    "t4h","t4h_Padj","t8h","t8h_Padj","t10h","t10h_Padj","cluster", "GO.ID",
                                    "KO.ID","E-value","KO_definition","UniprotID","protein_name",
                                    "MF","CC","BP","E-value","Bitscore","LT_logFC","LT_CPM","LT_PValue")

NS_LFCanno_df$cluster <- "Not2Sig"


NS_LFCanno <- subset(NS_LFCanno_df, select=c("gene_id","t15m","t30m","t1h","t2h","t4h","t8h","t10h","LT_logFC","LT_PValue",
                                                  "cluster","UniprotID","protein_name","KO.ID","KO_definition",
                                                  "GO.ID","MF","BP","CC"))

not_sig_anno_df <- not_sig_genes %>% 
  tibble::rownames_to_column(.,"gene_id") %>% #Use full_join to make sure we keep all the expression data and don't limit the GO and KO terms
  left_join(., GO, by = NULL, copy = FALSE, suffix = c(".x",".y")) %>%
  left_join(., norepeats_KO, by = NULL, copy = FALSE, suffix = c(".x",".y")) %>%
  left_join(., gene_uniprot_anno, by = NULL, copy = FALSE, suffix = c(".x",".y")) %>%
  left_join(., DE_LT, by = NULL, copy = FALSE, suffix = c(".x",".y"))
colnames(not_sig_anno_df) <- c("gene_id","cluster", "GO.ID",
                                    "KO.ID","E-value","KO_definition","UniprotID","protein_name",
                                    "MF","CC","BP","E-value","Bitscore","LT_logFC","LT_CPM","LT_PValue")
not_sig_anno_df$t15m <- "0"
not_sig_anno_df$t30m <- "0"
not_sig_anno_df$t1h <- "0"
not_sig_anno_df$t2h <- "0"
not_sig_anno_df$t4h <- "0"
not_sig_anno_df$t8h <- "0"
not_sig_anno_df$t10h <- "0"
not_sig_anno <- subset(not_sig_anno_df, select=c("gene_id","t15m","t30m","t1h","t2h","t4h","t8h","t10h","LT_logFC","LT_PValue",
                                                  "cluster","UniprotID","protein_name","KO.ID","KO_definition",
                                                  "GO.ID","MF","BP","CC"))

all_anno <- rbind(clusters_GO_KO_2Sig,NS_LFCanno,not_sig_anno)
paralogs <- rbind(paralogs_multi,singletons,paralogs_pairs)

raw_data <- as.data.frame(gene_cnts)

padj <- padj_df %>% tibble::rownames_to_column(.,"gene_id")

raw_genes <- raw_data %>%
  tibble::rownames_to_column(.,"gene_id") %>%
  left_join(., all_anno, by = NULL, copy = FALSE, suffix = c(".x",".y")) %>%
  left_join(., paralogs, by = NULL, copy = FALSE, suffix = c(".x",".y")) %>%
  left_join(., padj, by = NULL, copy = FALSE, suffix = c(".x",".y"))
colnames(raw_genes) <- c("gene_id","0m_R1","15m_R1","30m_R1","60m_R1","2hr_R1","4hr_R1","8hr_R1","10hr_R1",
                                     "0m_R2","15m_R2","30m_R2","60m_R2","2hr_R2","4hr_R2","8hr_R2","10hr_R2",
                                     "0m_R3","15m_R3","30m_R3","60m_R3","2hr_R3","4hr_R3","8hr_R3","10hr_R3",
                                     "t15m","t30m","t1h","t2h","t4h","t8h","t10h","LT_logFC","LT_PValue","cluster","UniprotID",
                                     "protein_name","KO.ID","KO_definition","GO.ID","MF","BP","CC","orthogroup","cnt_OG",
                                     "padjScreen","15m_Padj","30m_Padj","1h_Padj","2h_Padj","4h_Padj","8h_Padj","10h_Padj")
raw_genes_df <- subset(raw_genes, select=c("gene_id","t15m","15m_Padj","t30m","30m_Padj","t1h","1h_Padj","t2h","2h_Padj",
                                        "t4h","4h_Padj","t8h","8h_Padj","t10h","10h_Padj","LT_logFC","LT_PValue","orthogroup","cnt_OG",
                                        "cluster","UniprotID","protein_name","KO.ID","KO_definition", "GO.ID","MF","BP","CC"))
raw_genes <- subset(raw_genes, select=c("gene_id","t15m","t30m","t1h","t2h","t4h","t8h","t10h","LT_logFC","LT_PValue","orthogroup","cnt_OG",
                                        "cluster","UniprotID","protein_name","KO.ID","KO_definition", "GO.ID","MF","BP","CC"))

raw_genes[,c(2:8)] <- sapply(raw_genes[,c(2:8)], as.numeric)
raw_genes[,c(10)] <- sapply(raw_genes[,c(10)], as.character)

clust_paralogs <- raw_genes %>% filter(str_detect(cluster,"clust"))
```

Diatom genomes are not always well-annotated. How many of our genes
lack annotation?

```
no_annotation_data <- subset(all_anno, select=-c(1:10)) 
no_annotation_data_df <- filter(no_annotation_data, rowSums(is.na(no_annotation_data)) != ncol(no_annotation_data))
```

### 6. Manuscript graphs

#### 6.1 Gene expression behavior in short- and long-term

Graph of short-term behavior of genes found to be significant in the
long-term

```
#Set up the downregulated variables
dr_LT <- all_anno[,c(1,2,3,4,5,6,7,8,9)] 
colnames(dr_LT) <- c("gene_id","1","2","3","4","5","6","7","8")
dr_LT <- (na.omit(dr_LT))
dr_LT[,c(2:9)] <- sapply(dr_LT[,c(2:9)], as.numeric)
dr_LT <- subset(dr_LT, (`8`) <= -1) 
dr_ST <- pivot_longer(dr_LT[,c(1,2,3,4,5,6,7,8)], -c(gene_id), names_to = "timept")
dr_ST_md = dr_ST %>% group_by(timept) %>% summarize(value = mean(value), ) %>% mutate(gene_id = "repressed")
dr_LT <- pivot_longer(dr_LT[,c(1,2,3,4,5,6,7,8,9)], -c(gene_id), names_to = "timept")
dr_LT_md = dr_LT %>% group_by(timept) %>% summarize(value = mean(value), ) %>% mutate(gene_id = "repressed") 
dr_LT <- filter(dr_LT, timept == "8") #Get just the long-term timepoint for the boxplot

#Set up the upregulated variables
ur_LT <- all_anno[,c(1,2,3,4,5,6,7,8,9)] 
colnames(ur_LT) <- c("gene_id","1","2","3","4","5","6","7","8")
ur_LT <- (na.omit(ur_LT))
ur_LT[,c(2:9)] <- sapply(ur_LT[,c(2:9)], as.numeric)
ur_LT <- subset(ur_LT, (`8`) >= 1) 
ur_ST <- pivot_longer(ur_LT[,c(1,2,3,4,5,6,7,8)], -c(gene_id), names_to = "timept")
ur_ST_md = ur_ST %>% group_by(timept) %>% summarize(sd = sd(value, na.rm=TRUE), value = mean(value), ) %>% mutate(gene_id = "induced")
ur_LT <- pivot_longer(ur_LT[,c(1,2,3,4,5,6,7,8,9)], -c(gene_id), names_to = "timept")
ur_LT_md = ur_LT %>% group_by(timept) %>% summarize(value = mean(value), ) %>% mutate(gene_id = "induced") 
ur_LT <- filter(ur_LT, timept == "8") #Get just the long-term timepoint for the boxplot

#Split out the outliers to jitter them in the plot for visibility
dr_LT_outliers <- dr_LT %>% group_by(timept) %>%
  filter(value > quantile(value, 0.75) + 1.5 * IQR(value) | value < quantile(value, 0.25) - 1.5 * IQR(value))
ur_LT_outliers <- ur_LT %>% group_by(timept) %>%
  filter(value > quantile(value, 0.75) + 1.5 * IQR(value) | value < quantile(value, 0.25) - 1.5 * IQR(value))

ggplot(NULL)+ 
    #geom_line(data = dr_ST, size = 1, alpha=1/50, aes(x = timept, y = value, group = gene_id))  +
    geom_line(data = dr_ST_md, size=2, alpha = 1,  color = "#a9acff",aes(x = timept, y = value, group = gene_id)) +
    #geom_line(data = ur_ST, size = 1, alpha=1/50, aes(x = timept, y = value, group = gene_id)) +
    geom_line(data = ur_ST_md, size=2, alpha = 1,  color = "#28db69",  aes(x = timept, y = value, group = gene_id)) +
    geom_boxplot(data = dr_LT, color = "black", fill = "#28db69", notch = TRUE, aes(x = timept, y = value), outlier.shape = NA) +
    geom_boxplot(data = ur_LT, color = "black", fill = "#a9acff", notch = TRUE, aes(x = timept, y = value), outlier.shape = NA) +
    geom_jitter(aes(x = timept, y = value), height = 0, width = 0.1, data = ur_LT_outliers) +
    geom_jitter(aes(x = timept, y = value), height = 0, width = 0.1, data = dr_LT_outliers) +
    geom_hline(yintercept = 0, linetype = "dashed") +
    theme_classic() +
    theme(axis.ticks.x=element_blank(), 
          axis.text=element_text(size=12), 
          axis.title=element_text(size=14,face="bold")) +
    xlab("Time exposed to 0 ppt ASW") + 
    ylab("Log2 fold change") +
    ggtitle("Median expression over time for genes induced/repressed at 4 months") +
    theme(plot.title = element_text(hjust = 0.5, face = "bold", size = 16))
```

#### 6.2 Graph of median expression of genes DE at 1h and 10 h

```
min_1h <- filter(logFCdown_0v1h , which.pmin(logFCdown_0v1h) == 5)
min_1h <- tibble::rownames_to_column(min_1h, var = "gene_id")
min_1h <- pivot_longer(min_1h[,c(1,2,4,6,8,10,12,14)], -c(gene_id), names_to = "timept")
min_1h$timept <- ordered(min_1h$timept, levels =c("0v15m","0v30m","0v1h","0v2h","0v4h","0v8h","0v10h"))
min_1h_md = min_1h %>% group_by(timept) %>% summarize(sd = sd(value, na.rm=TRUE), value = median(value), ) %>% mutate(gene_id = "induced")
#outlier_min_1h <-  min_1h %>% group_by(timept) %>%
#  filter(value > quantile(value, 0.75) + 1.5 * IQR(value) | value < quantile(value, 0.25) - 1.5 * IQR(value))

max_1h <- filter(logFCup_0v1h , which.pmax(logFCup_0v1h) == 5)
max_1h <- tibble::rownames_to_column(max_1h, var = "gene_id")
max_1h <- pivot_longer(max_1h[,c(1,2,4,6,8,10,12,14)], -c(gene_id), names_to = "timept")
max_1h$timept <- ordered(max_1h$timept, levels =c("0v15m","0v30m","0v1h","0v2h","0v4h","0v8h","0v10h"))
max_1h_md = max_1h %>% group_by(timept) %>% summarize(sd = sd(value, na.rm=TRUE), value = median(value), ) %>% mutate(gene_id = "induced")
#outlier_max_1h <-  max_1h %>% group_by(timept) %>%
#  filter(value > quantile(value, 0.75) + 1.5 * IQR(value) | value < quantile(value, 0.25) - 1.5 * IQR(value))

min_10h <- filter(logFCdown_0v10h, which.pmin(logFCdown_0v10h) == 13)
min_10h <- tibble::rownames_to_column(min_10h, var = "gene_id")
min_10h <- pivot_longer(min_10h[,c(1,2,4,6,8,10,12,14)], -c(gene_id), names_to = "timept")
min_10h$timept <- ordered(min_10h$timept, levels =c("0v15m","0v30m","0v1h","0v2h","0v4h","0v8h","0v10h"))
min_10h_md = min_10h %>% group_by(timept) %>% summarize(sd = sd(value, na.rm=TRUE), value = median(value), ) %>% mutate(gene_id = "repressed")
#outlier_min_10h <-  min_10h %>% group_by(timept) %>%
#  filter(value > quantile(value, 0.75) + 1.5 * IQR(value) | value < quantile(value, 0.25) - 1.5 * IQR(value))

max_10h <- filter(logFCup_0v10h, which.pmax(logFCup_0v10h) == 13)
max_10h <- tibble::rownames_to_column(max_10h, var = "gene_id")
max_10h <- pivot_longer(max_10h[,c(1,2,4,6,8,10,12,14)], -c(gene_id), names_to = "timept")
max_10h$timept <- ordered(max_10h$timept, levels =c("0v15m","0v30m","0v1h","0v2h","0v4h","0v8h","0v10h"))
max_10h_md = max_10h %>% group_by(timept) %>% summarize(sd = sd(value, na.rm=TRUE), value = median(value), ) %>% mutate(gene_id = "induced")
#outlier_max_10h <-  max_10h %>% group_by(timept) %>%
#  filter(value > quantile(value, 0.75) + 1.5 * IQR(value) | value < quantile(value, 0.25) - 1.5 * IQR(value))


ggplot(NULL)+ 
    geom_line(data = min_1h_md, size=2, alpha = 1, color = "#28db69",aes(x = timept, y = value, group = gene_id)) +
    geom_line(data = max_1h_md, size=2, alpha = 1, color = "#a9acff",aes(x = timept, y = value, group = gene_id)) +
    geom_boxplot(data = min_1h, notch = TRUE, aes(x = timept, y = value), fill= "#28db69", width = 0.15, outlier.shape = NA) +
    geom_boxplot(data = max_1h, notch = TRUE, aes(x = timept, y = value), fill= "#a9acff", width = 0.15, outlier.shape = NA) +
    geom_hline(yintercept = 0, linetype = "dashed") +
    ylim(-2.5,2.5) +
    theme_classic() +
    theme(axis.ticks.x=element_blank(), 
          axis.text=element_text(size=12), 
          axis.title=element_text(size=14,face="bold")) +
    xlab("Time exposed to 0 ppt ASW") + 
    ylab("Log2 fold change") +
    ggtitle("Median expression over time for genes induced/repressed at 1h") +
    theme(plot.title = element_text(hjust = 0.5, face = "bold", size = 16))
```

```
ggplot(NULL)+ 
    geom_line(data = min_10h_md, size=2, alpha = 1, color = "#28db69",aes(x = timept, y = value, group = gene_id)) +
    geom_line(data = max_10h_md, size=2, alpha = 1, color = "#a9acff",aes(x = timept, y = value, group = gene_id)) +
    geom_boxplot(data = min_10h, notch = TRUE, aes(x = timept, y = value), fill= "#28db69", width = 0.15, outlier.shape = NA) +
    geom_boxplot(data = max_10h, notch = TRUE, aes(x = timept, y = value), fill= "#a9acff", width = 0.15, outlier.shape = NA) +
    geom_hline(yintercept = 0, linetype = "dashed") +
    ylim(-2.5,2.5) +
    theme_classic() +
    theme(axis.ticks.x=element_blank(), 
          axis.text=element_text(size=12), 
          axis.title=element_text(size=14,face="bold")) +
    xlab("Time exposed to 0 ppt ASW") + 
    ylab("Log2 fold change") +
    ggtitle("Median expression over time for genes induced/repressed at 10 h") +
    theme(plot.title = element_text(hjust = 0.5, face = "bold", size = 16))
```

#### 6.3 Heatmaps for cycles of interest

Use this code to isolate genes of interest from the metadata tables
produced in Section 5.

```
colors = c("#28db69","white","purple")
colors=colorRampPalette(colors)(1000)
my_palette <- colorRampPalette(c("#28db69", "white", "#a9acff"))(n = 1000)

Chlorophyll_hmp <- as.matrix(clusters_GO_KO_2Sig %>% arrange(.,KO_definition) %>%
                            filter(str_detect(KO_definition,"EC:6.1.1.17|EC:1.2.1.70|EC:5.4.3.8|EC:4.2.1.24|EC:2.5.1.61|EC:4.2.1.75|EC:4.1.1.37|EC:2.1.1.107|EC:1.3.3.4|EC:1.3.3.3|EC:6.6.1.1|EC:2.1.1.11|EC:1.3.1.75|EC:1.3.1.33|EC:2.5.1.62|EC:2.5.1.133|EC:1.1.1.294"))%>%
                            dplyr::select(`gene_id`,`KO_definition`,`t15m`,`t30m`,`t1h`,`t2h`,`t4h`,`t8h`,`t10h`,`LT_logFC`) %>%
                            unite(.,names,c(`gene_id`,`KO_definition`))
                            %>% remove_rownames %>% column_to_rownames(var="names"))
heatmap.2(Chlorophyll_hmp,dendrogram = "none",trace="none",col=colors,main="Genes in chlorophyll biosynthesis", cexRow = 0.75, cexCol = 1, Colv = FALSE, Rowv = FALSE, offsetRow = -20, density.info="none")
```

```
Glycolysis_hmp <- as.matrix(clusters_GO_KO_2Sig %>% arrange(.,KO_definition) %>% 
                            filter(str_detect(KO_definition,"EC:1.2.1.12|EC:2.7.1.90|EC:2.7.1.11|EC:3.1.3.11|EC:4.1.2.13|EC:2.7.2.3|EC:2.7.1.40]|EC:5.3.1.9|EC:4.2.1.11]|EC:5.4.2.11]"))%>%
                            dplyr::select(`gene_id`,`t15m`,`t30m`,`t1h`,`t2h`,`t4h`,`t8h`,`t10h`,`LT_logFC`,`KO_definition`) %>%
                            unite(.,names,c(`gene_id`,`KO_definition`))
                            %>% remove_rownames %>% column_to_rownames(var="names"))

Chitin_hmp <- as.matrix(clusters_GO_KO_2Sig %>% arrange(.,KO_definition) %>% 
                            filter(str_detect(KO_definition,"2.6.1.16|3.5.1.25|5.4.2.3|2.7.7.23|2.7.7.83|2.4.1.16|3.5.2.5|3.2.1.14|3.2.1.52"))%>%
                            dplyr::select(`gene_id`,`KO_definition`,`t15m`,`t30m`,`t1h`,`t2h`,`t4h`,`t8h`,`t10h`,`LT_logFC`) %>%
                            unite(.,names,c(`gene_id`,`KO_definition`))
                            %>% remove_rownames %>% column_to_rownames(var="names"))

CalvinCycle_hmp <- as.matrix(clusters_GO_KO_2Sig %>% arrange(.,KO_definition) %>%
                            filter(str_detect(KO_definition,"EC:2.7.2.3|EC:1.2.1.12|EC:4.1.2.13|EC:3.1.3.11|EC:2.2.1.1|EC:3.1.3.37|EC:5.3.1.6"))%>%
                            dplyr::select(`gene_id`,`KO_definition`,`t15m`,`t30m`,`t1h`,`t2h`,`t4h`,`t8h`,`t10h`,`LT_logFC`) %>%
                            unite(.,names,c(`gene_id`,`KO_definition`))
                            %>% remove_rownames %>% column_to_rownames(var="names"))

Metablism_hmp <- rbind (Glycolysis_hmp,CalvinCycle_hmp,Chitin_hmp)

heatmap.2(CalvinCycle_hmp,dendrogram = "none",trace="none",col=colors,main="Genes in oxidative stress", cexRow = 0.75, cexCol = 1, Colv = FALSE, Rowv = FALSE, offsetRow = 0, density.info="none")
```

```
Proline_hmp <- as.matrix(clusters_GO_KO_2Sig  %>% arrange(.,KO_definition) %>%
                            filter(str_detect(KO_definition,"EC:2.7.2.11|EC:1.2.1.41|EC:3.4.11.5|EC:2.6.1.13|EC:1.5.1.2]|EC:4.3.1.12"))%>%
                            dplyr::select(`gene_id`,`KO_definition`,`t15m`,`t30m`,`t1h`,`t2h`,`t4h`,`t8h`,`t10h`,`LT_logFC`) %>%
                            unite(.,names,c(`gene_id`,`KO_definition`))
                            %>% remove_rownames %>% column_to_rownames(var="names"))

Taurine_hmp <- as.matrix(raw_genes  %>% arrange(.,KO_definition) %>%
                            filter(str_detect(protein_name,"L-cysteate sulfo-lyase|glutamate decarboxylase|Gamma-glutamyl cyclotransferase"))%>%
                            dplyr::select(`gene_id`,`KO_definition`,`t15m`,`t30m`,`t1h`,`t2h`,`t4h`,`t8h`,`t10h`,`LT_logFC`) %>%
                            unite(.,names,c(`gene_id`,`KO_definition`))
                            %>% remove_rownames %>% column_to_rownames(var="names"))

Aquaporin_hmp <- as.matrix(raw_genes  %>% arrange(.,KO_definition) %>%
                            filter(str_detect(protein_name,"Aquaporin AqpM"))%>%
                            dplyr::select(`gene_id`,`KO_definition`,`t15m`,`t30m`,`t1h`,`t2h`,`t4h`,`t8h`,`t10h`,`LT_logFC`) %>%
                            unite(.,names,c(`gene_id`,`KO_definition`))
                            %>% remove_rownames %>% column_to_rownames(var="names"))

Osmo_hmp <- rbind(Proline_hmp, Taurine_hmp, Aquaporin_hmp)

heatmap.2(Osmo_hmp,dendrogram = "none",trace="none",col=colors,main="Genes in proline metabolism",  cexRow = 0.75, cexCol = 1, Colv = FALSE, Rowv = FALSE, offsetRow = 0, density.info="none")
```

```
HeatShock_hmp_KO <- as.matrix(raw_genes %>% arrange(.,KO_definition) %>%
                            filter(str_detect(KO.ID,"K04077|K04078|K03686|K03687|K04082|K04083|K04079|K13993"))%>%
                            dplyr::select(`gene_id`,`cluster`,`t15m`,`t30m`,`t1h`,`t2h`,`t4h`,`t8h`,`t10h`,`LT_logFC`) %>%
                            unite(.,names,c(`gene_id`,`cluster`)) %>% 
                            remove_rownames %>% column_to_rownames(var="names"))
HeatShock_hmp_UP <- as.matrix(raw_genes %>% arrange(.,protein_name) %>%
                            filter(str_detect(protein_name,"Hsp70|HSP70"))%>%
                            dplyr::select(`gene_id`,`cluster`,`t15m`,`t30m`,`t1h`,`t2h`,`t4h`,`t8h`,`t10h`,`LT_logFC`) %>%
                            unite(.,names,c(`gene_id`,`cluster`)) %>% 
                            remove_rownames %>% column_to_rownames(var="names"))
HeatShock_hmp <- rbind(HeatShock_hmp_KO,HeatShock_hmp_UP)

heatmap.2(HeatShock_hmp,dendrogram = "none",trace="none",col=colors,main="Genes in oxidative stress", cexRow = 0.75, cexCol = 1, Colv = FALSE, Rowv = FALSE, offsetRow = -20, density.info="none")
```

```
Sig_Ox <- as.matrix(raw_genes %>% arrange(.,KO_definition) %>%
                            filter(str_detect(gene_id,"CCRYP_015211|CCRYP_020438|CCRYP_013781|CCRYP_016723|CCRYP_002248|CCRYP_002534|CCRYP_010295|CCRYP_016269|CCRYP_002901|CCRYP_003411|CCRYP_020615|CCRYP_008084"))%>%
                            dplyr::select(`gene_id`,`KO_definition`,`t15m`,`t30m`,`t1h`,`t2h`,`t4h`,`t8h`,`t10h`,`LT_logFC`) %>%
                            unite(.,names,c(`gene_id`,`KO_definition`)) %>% 
                            remove_rownames %>% column_to_rownames(var="names"))

heatmap.2(Sig_Ox,dendrogram = "none",trace="none",col=colors,main="Genes in oxidative stress", cexRow = 0.75, cexCol = 1, Colv = FALSE, Rowv = FALSE, offsetRow = -30, density.info="none")
```

#### 6.4 Cluster plots

##### 6.4.1 Summary of the averages of all the clusters

```
cluster_LF10C <- dplyr::filter(clusters_GO_KO_2Sig) 
cluster_LFC10C <- pivot_longer(cluster_LF10C[ ,c(1:8,11)], -c(gene_id, cluster), names_to = "timept")
cluster_LFC10C$timept <- ordered(cluster_LFC10C$timept, levels=c("t15m","t30m","t1h","t2h","t4h","t8h","t10h"))
cluster1 <- dplyr::filter(cluster_LFC10C, (cluster == 'clust1')) 
cluster2 <- dplyr::filter(cluster_LFC10C, (cluster == 'clust2'))
cluster3 <- dplyr::filter(cluster_LFC10C, (cluster == 'clust3'))  
cluster4 <- dplyr::filter(cluster_LFC10C, (cluster == 'clust4')) 
cluster5 <- dplyr::filter(cluster_LFC10C, (cluster == 'clust5')) 
cluster6 <- dplyr::filter(cluster_LFC10C, (cluster == 'clust6'))  
cluster7 <- dplyr::filter(cluster_LFC10C, (cluster == 'clust7'))   
cluster8 <- dplyr::filter(cluster_LFC10C, (cluster == 'clust8'))  
cluster9 <- dplyr::filter(cluster_LFC10C, (cluster == 'clust9')) 
cluster10 <- dplyr::filter(cluster_LFC10C, (cluster == 'clust10'))  

data1 = cluster1 %>% group_by(timept,cluster) %>% summarize(sd = sd(value, na.rm = TRUE),value = mean(value), ) %>% mutate(gene_id = "1")
```

```
## `summarise()` has grouped output by 'timept'. You can override using the
## `.groups` argument.
```

```
data2 = cluster2 %>% group_by(timept,cluster) %>% summarize(sd = sd(value, na.rm = TRUE),value = mean(value), ) %>% mutate(gene_id = "2")
```

```
## `summarise()` has grouped output by 'timept'. You can override using the
## `.groups` argument.
```

```
data3 = cluster3 %>% group_by(timept,cluster) %>% summarize(sd = sd(value, na.rm = TRUE),value = mean(value), ) %>% mutate(gene_id = "3")
```

```
## `summarise()` has grouped output by 'timept'. You can override using the
## `.groups` argument.
```

```
data4 = cluster4 %>% group_by(timept,cluster) %>% summarize(sd = sd(value, na.rm = TRUE),value = mean(value), ) %>% mutate(gene_id = "4")
```

```
## `summarise()` has grouped output by 'timept'. You can override using the
## `.groups` argument.
```

```
data5 = cluster5 %>% group_by(timept,cluster) %>% summarize(sd = sd(value, na.rm = TRUE),value = mean(value), ) %>% mutate(gene_id = "5")
```

```
## `summarise()` has grouped output by 'timept'. You can override using the
## `.groups` argument.
```

```
data6 = cluster6 %>% group_by(timept,cluster) %>% summarize(sd = sd(value, na.rm = TRUE),value = mean(value), ) %>% mutate(gene_id = "6")
```

```
## `summarise()` has grouped output by 'timept'. You can override using the
## `.groups` argument.
```

```
data7 = cluster7 %>% group_by(timept,cluster) %>% summarize(sd = sd(value, na.rm = TRUE),value = mean(value), ) %>% mutate(gene_id = "7")
```

```
## `summarise()` has grouped output by 'timept'. You can override using the
## `.groups` argument.
```

```
data8 = cluster8 %>% group_by(timept,cluster) %>% summarize(sd = sd(value, na.rm = TRUE),value = mean(value), ) %>% mutate(gene_id = "8")
```

```
## `summarise()` has grouped output by 'timept'. You can override using the
## `.groups` argument.
```

```
data9 = cluster9 %>% group_by(timept,cluster) %>% summarize(sd = sd(value, na.rm = TRUE),value = mean(value), ) %>% mutate(gene_id = "9")
```

```
## `summarise()` has grouped output by 'timept'. You can override using the
## `.groups` argument.
```

```
data10 = cluster10 %>% group_by(timept,cluster) %>% summarize(sd = sd(value, na.rm = TRUE),value = mean(value), ) %>% mutate(gene_id = "10")
```

```
## `summarise()` has grouped output by 'timept'. You can override using the
## `.groups` argument.
```

```
data = rbind(data1, data2, data3, data4, data5, data6, data7, data8, data9, data10)
data$cluster <- factor(data$cluster, levels=c("clust1","clust2","clust3","clust4","clust5","clust6","clust7","clust8","clust9","clust10"))

#All averages
ggplot(data,  aes(x = timept, y = value, group = cluster, color = cluster)) + 
    geom_line(size = 1.5) +
    ylim(-3.5,3.5) +
    geom_hline(yintercept = 0, linetype = "dashed") +
    theme(axis.text.x = element_text(vjust=115, face = "bold"),
          axis.ticks.x=element_blank()) +
    xlab("Time exposed to 0 ppt ASW") + 
    ylab("Average fold change (log 2)") +
    scale_color_viridis(discrete = TRUE, option = "D") +
    ggtitle("Average expression over time for each cluster") +
    theme(plot.title = element_text(hjust = 0.5, face = "bold"))
```

##### 6.4.2 Individual graphs showing cluster averages with error bars

```
#Individual cluster averages with background data
ggplot(data1, aes(x = timept, y = value, group = gene_id, ymin = value-sd, ymax = value+sd))  + 
    geom_line(color = "#450755", size = 4) +
    geom_errorbar(width = 0.2) +
    geom_hline(yintercept = 0, linetype = "dashed",size=1) +
    ylim(-4,3) +
    theme(axis.title.x = element_blank()) +
    theme(axis.title.y = element_text(face="bold")) +
    theme(legend.position = "none") +
    ylab("Average fold change (log 2)") +
    theme_classic() +
    theme(plot.title = element_text(hjust = 0.5, face = "bold"))
```

```
ggplot(data2, aes(x = timept, y = value, group = gene_id, ymin = value-sd, ymax = value+sd))  + 
    geom_line(color = "#48247a", size = 4) +
    geom_errorbar(width = 0.2) +
    geom_hline(yintercept = 0, linetype = "dashed",size=1) +
    ylim(-4,3) +
    theme(axis.title.x = element_text(face="bold")) +
    theme(axis.title.y = element_text(face="bold")) +
    theme(legend.position = "none") +
    ylab("Average fold change (log 2)") +
    theme_classic() +
    theme(plot.title = element_text(hjust = 0.5, face = "bold"))
```

```
ggplot(data3, aes(x = timept, y = value, group = gene_id, ymin = value-sd, ymax = value+sd))  + 
    geom_line(color = "#3d488b", size = 4) +
    geom_errorbar(width = 0.2) +
    geom_hline(yintercept = 0, linetype = "dashed",size=1) +
    ylim(-4,3) +
    theme(axis.title.x = element_blank()) +
    theme(axis.title.y = element_text(face="bold")) +
    theme(legend.position = "none") +
    ylab("Average fold change (log 2)") +
    theme_classic() +
    theme(plot.title = element_text(hjust = 0.5, face = "bold"))
```

```
ggplot(data4, aes(x = timept, y = value, group = gene_id, ymin = value-sd, ymax = value+sd))  + 
    geom_line(color = "#2d6790", size = 4) +
    geom_errorbar(width = 0.2) +
    geom_hline(yintercept = 0, linetype = "dashed",size=1) +
    ylim(-4,3) +
    theme(axis.title.x = element_blank()) +
    theme(axis.title.y = element_text(face="bold")) +
    theme(legend.position = "none") +
    ylab("Average fold change (log 2)") +
    theme_classic() +
    theme(plot.title = element_text(hjust = 0.5, face = "bold"))
```

```
ggplot(data5, aes(x = timept, y = value, group = gene_id, ymin = value-sd, ymax = value+sd))  + 
    geom_line(color = "#1e828f", size = 4) +
    geom_errorbar(width = 0.2) +
    geom_hline(yintercept = 0, linetype = "dashed",size=1) +
    ylim(-4,3) +
    theme(axis.title.x = element_blank()) +
    theme(axis.title.y = element_text(face="bold")) +
    theme(legend.position = "none") +
    ylab("Average fold change (log 2)") +
    theme_classic() +
    theme(plot.title = element_text(hjust = 0.5, face = "bold"))
```

```
ggplot(data6, aes(x = timept, y = value, group = gene_id, ymin = value-sd, ymax = value+sd))  + 
    geom_line(color = "#129f89", size = 4) +
    geom_errorbar(width = 0.2) +
    geom_hline(yintercept = 0, linetype = "dashed",size=1) +
    ylim(-4,3) +
    theme(axis.title.x = element_blank()) +
    theme(axis.title.y = element_text(face="bold")) +
    theme(legend.position = "none") +
    ylab("Average fold change (log 2)") +
    theme_classic() +
    theme(plot.title = element_text(hjust = 0.5, face = "bold"))
```

```
ggplot(data7, aes(x = timept, y = value, group = gene_id, ymin = value-sd, ymax = value+sd))  + 
    geom_line(color = "#2cb877", size = 4) +
    geom_errorbar(width = 0.2) +
    geom_hline(yintercept = 0, linetype = "dashed",size=1) +
    ylim(-4,3) +
    theme(axis.title.x = element_text(face="bold")) +
    theme(axis.title.y = element_text(face="bold")) +
    theme(legend.position = "none") +
    ylab("Average fold change (log 2)") +
    theme_classic() +
    theme(plot.title = element_text(hjust = 0.5, face = "bold"))
```

```
ggplot(data8, aes(x = timept, y = value, group = gene_id, ymin = value-sd, ymax = value+sd))  + 
    geom_line(color = "#69cf52", size = 4) +
    geom_errorbar(width = 0.2) +
    geom_hline(yintercept = 0, linetype = "dashed",size=1) +
    ylim(-4,3) +
    theme(axis.title.x = element_text(face="bold")) +
    theme(axis.title.y = element_text(face="bold")) +
    theme(legend.position = "none") +
    ylab("Average fold change (log 2)") +
    theme_classic() +
    theme(plot.title = element_text(hjust = 0.5, face = "bold"))
```

```
ggplot(data9, aes(x = timept, y = value, group = gene_id, ymin = value-sd, ymax = value+sd))  + 
    geom_line(color = "#b3e000", size = 4) +
    geom_errorbar(width = 0.2) +
    geom_hline(yintercept = 0, linetype = "dashed",size=1) +
    ylim(-4,3) +
    theme(axis.title.x = element_text(face="bold")) +
    theme(axis.title.y = element_text(face="bold")) +
    theme(legend.position = "none") +
    ylab("Average fold change (log 2)") +
    theme_classic() +
    theme(plot.title = element_text(hjust = 0.5, face = "bold"))
```

```
ggplot(data10, aes(x = timept, y = value, group = gene_id, ymin = value-sd, ymax = value+sd))  + 
    geom_line(color = "#fee900", size = 4) +
    geom_errorbar(width = 0.2) +
    geom_hline(yintercept = 0, linetype = "dashed",size=1) +
    ylim(-4,3) +
    theme(axis.title.x = element_text(face="bold")) +
    theme(axis.title.y = element_text(face="bold")) +
    theme(legend.position = "none") +
    ylab("Average fold change (log 2)") +
    theme_classic() +
    theme(plot.title = element_text(hjust = 0.5, face = "bold"))
```

### 7. Paralogs simulation

We developed a function in an attempt to determine if the
distribution of paralogs was significant or not

```
paralog_sim_df <- subset(raw_genes, select=c("gene_id","orthogroup","cnt_OG","cluster"))

singles_sim_df <- paralog_sim_df %>%distinct(gene_id, .keep_all = TRUE) %>% group_by(orthogroup) %>% filter( n() == 1 )
singles_sim_df[,c(4)] <- sapply(singles_sim_df[,c(4)], as.character) 
singles_sim_df <- (na.omit(singles_sim_df))

doubles_sim_df <- paralog_sim_df %>% distinct(gene_id, .keep_all = TRUE) %>% group_by(orthogroup) %>% filter( n() == 2 )
doubles_sim_df[,c(4)] <- sapply(doubles_sim_df[,c(4)], as.character)
doubles_sim_df <- (na.omit(doubles_sim_df))

# 'notin' for lazy exclusion of filter searches
`%notin%` <- Negate(`%in%`) # a suprise tool to help us later

# function specifying a loop, X cycles, df as user input to function
paralog_sim <- function(x = 10, df) { # current default 10 for testing, change to 1000
  
  # create empty vector of length X for data storage
  simres <- rep(0, x)
  
  # create empty df for row info storage (for testing)
  simstore <- as.data.frame(matrix(0, x, 2))
  # simstore <- matrix(0,x,2) # uncomment for uniformity test
  
  # get length of df outside loop, for random draw boundary (0 : N)
  N <- nrow(df)
  # (loop body)
  for(n in 1:x) {
    # random draw of numbers (in vector slots [1] and [2]), without replacement
    random <- sample(1:N, 2, replace = F) # cannot draw same # twice
    
    # store values in simstore, for testing
    simstore[n,1] <- df[random[1],4]
    simstore[n,2] <- df[random[2],4]
    
    # store just the drawn nums, for uniformity test
    # simstore[n, ] <- random

    # IF statement to store 1 if one random row is not sig and other is sig
    if ((df[random[1],4] %in% c("Not2Sig","NotSig") & 
        df[random[2],4] %notin% c("Not2Sig", "NotSig")) |
       (df[random[1],4] %notin% c("Not2Sig", "NotSig") & 
        df[random[2],4] %in% c("Not2Sig", "NotSig"))) {
      
      # store 1 in vector of length X made at top of function     
        simres[n] <- 1
       }
  }


  # after loop complete, get mean of vector X (1s and 0s only, mean is proportion that match criterion: 1 row sig, other not)
  simmean <- mean(simres) 
  
  # return this as function output
  #return(simmean) # comment this out for troubleshooting
  
  # if above return (ends function) is commented out, diagnostic info returned: keep the x small to ensure results are reasonably human-readable
  testlist <- list(themean = simmean, resvector = simres, allstore = simstore)
  return(testlist)
}

simulation_test1 <- paralog_sim(1000, singles_sim_df)
simulation_test2 <- paralog_sim(1000, singles_sim_df)
simulation_test3 <- paralog_sim(1000, singles_sim_df)
simulation_test4 <- paralog_sim(1000, singles_sim_df)
simulation_test5 <- paralog_sim(1000, singles_sim_df)
simulation_test6 <- paralog_sim(1000, singles_sim_df)
simulation_test7 <- paralog_sim(1000, singles_sim_df)
simulation_test8 <- paralog_sim(1000, singles_sim_df)
simulation_test9 <- paralog_sim(1000, singles_sim_df)
simulation_test10 <- paralog_sim(1000, singles_sim_df)
simulation_test11 <- paralog_sim(1000, singles_sim_df)
simulation_test12 <- paralog_sim(1000, singles_sim_df)
simulation_test13 <- paralog_sim(1000, singles_sim_df)
simulation_test14 <- paralog_sim(1000, singles_sim_df)
simulation_test15 <- paralog_sim(1000, singles_sim_df)
simulation_test16 <- paralog_sim(1000, singles_sim_df)
simulation_test17 <- paralog_sim(1000, singles_sim_df)
simulation_test18 <- paralog_sim(1000, singles_sim_df)
simulation_test19 <- paralog_sim(1000, singles_sim_df)
simulation_test20 <- paralog_sim(1000, singles_sim_df)
simulation_test21 <- paralog_sim(1000, singles_sim_df)
simulation_test22 <- paralog_sim(1000, singles_sim_df)
simulation_test23 <- paralog_sim(1000, singles_sim_df)
simulation_test24 <- paralog_sim(1000, singles_sim_df)
simulation_test25 <- paralog_sim(1000, singles_sim_df)
simulation_test26 <- paralog_sim(1000, singles_sim_df)
simulation_test27 <- paralog_sim(1000, singles_sim_df)

paralog_sample <- function(x = 10, df) { # current default 10 for testing, change to 1000
  
  # create empty vector of length X for data storage
  simres <- rep(0, x)
  
  # create empty df for row info storage (for testing)
  simstore <- as.data.frame(matrix(0, x, 2))
  # simstore <- matrix(0,x,2) # uncomment for uniformity test
  
  # get length of df outside loop, for random draw boundary (0 : N)
  N <- nrow(df)
  # (loop body)
  for(n in 1:x) {
    # random draw of numbers (in vector slots [1] and [2]), without replacement
    lines1 <- sample(1:N, 2, replace = F) # cannot draw same # twice
    
    # store values in simstore, for testing
    simstore[n,1] <- df[lines1[1],4]
    simstore[n,2] <- df[lines1[2],4]
    
    # store just the drawn nums, for uniformity test
    # simstore[n, ] <- lines1

    # IF statement to store 1 if one lines1 row is not sig and other is sig
    if ((df[lines1[1],4] %in% c("Not2Sig","NotSig") & 
        df[lines1[2],4] %notin% c("Not2Sig", "NotSig")) |
       (df[lines1[1],4] %notin% c("Not2Sig", "NotSig") & 
        df[lines1[2],4] %in% c("Not2Sig", "NotSig"))) {
      
      # store 1 in vector of length X made at top of function     
        simres[n] <- 1
       }
  }


  # after loop complete, get mean of vector X (1s and 0s only, mean is proportion that match criterion: 1 row sig, other not)
  simmean <- mean(simres) 
  
  # return this as function output
  return(simmean) # comment this out for troubleshooting
  
  # if above return (ends function) is commented out, diagnostic info returned: keep the x small to ensure results are reasonably human-readable
  testlist <- list(themean = simmean, resvector = simres, allstore = simstore)
  return(testlist)
}

paralog_1st <- paralog_sample(, paralog_sim_df)
```
