## Appendix S2 for "The dynamic response to hypoosmotic stress reveals distinct stages of freshwater acclimation by a euryhaline diatom": Appendix 2.rmd

````
---
title: '**Downey et al. - The dynamic response to hypoosmotic stress reveals distinct stages of freshwater acclimation by a euryhaline diatom**'
output:
  html_document:
    toc: true
    theme: united
  pdf_document:
    toc: true
    theme: united
---

This document gives an overview of the data analysis associated with the manuscript rapid hypo-osmotic stress response of the euryhaline diatom *Cyclotella cryptica* to freshwater.

# 1. Depiction of C. crytpica growth data 

## 1.1 Experimental design and methods

* Species: *Cyclotella cryptica*, strain CCMP332, from National Center for Marine Algae and Microbiota
* Culture maintained in 1 L artificial seawater (ASW) at 24 ppt in 15 C incubator, ~20 mol photons/m2/s irradiance
* Experimental design
  * Replicates: three 2 mL aliquots taken from source to inoculate flasks of 500 mL ASW 24 ppt, grown until exponential growth was reached
  * Inoculate volumes: 3 x 10^6 cells, estimated using 500 uL of replicates in FlowCam 
  * Treatments
    * Stress media: 38 mL freshwater (0 ppt) with ASW nutrients in a 50 mL falcon tube
    * Time-series exposures: 0 minutes, 15 minutes, 30 minutes, 60 minutes, 120 minutes, 240 minutes, 480 minutes, 600 minutes 
    * Mock/Control of ASW 24 ppt treated in exactly the same way to confirm no substantial stress occurred at 0 minutes
*Time series collection
  * Collected volume required for inoculation in two 50 mL falcon tubes
    * Concentrated in centrifuge 4 C for 3 minutes at 3000 rpm
    * Supernatant decanted and cells combined in 18 mL of ASW 0 ppt (2 mL for each time point)
    * Mock/Control suspended in 2 mL ASW 24 ppt
    * All volumes transferred to treatment vessels in 15 C incubator, gentle agitation on rocker
  * Collection
    * Mock/Control and 0 minute collected immediately
    * Spun down at 1500 rpm for 3 minutes at 4 C
    * Supernatant decanted and pellet flash-frozen in liquid nitrogen 
* RNA extraction: RNeasy kit, quality check with Qubit and Tapestation
* Library development: KAPA mRNA HyperPrep kit, libraries multiplexed for sequencing on Illumina HiSeq 4000 
  * Pooled with *Skeletonema marinoi* sequences from a similar short-term stress experiment conducted simultaneously

## 1.2 Packages used in analyses

```{r message=FALSE, warning=FALSE, results='hide'}
setwd ("/My Drive/ShortTermStress-IlluminaData/FullExp/Cryp-DE/")

#Load libraries to be used in this project into a variable.

libraries <- c("DESeq2", "edgeR", "shiny", "shinythemes", "tidyverse", "lubridate",  "viridis", "ggthemr", "ggrepel", "rlang", "reshape", "naniar", "pheatmap","rhdf5", "cgwtools", "NMF", "RColorBrewer" ,"topGO", "genefilter", "PoiClaClu", "gplots", "statmod", "stageR", "data.table", "radiant.data", "modelr", "here")

#Command to load libraries all at once.  
lapply(libraries, require, character.only = TRUE)

#You can save yourself some time and heartache by making an RData file that saves all your variables. If you ever accidentally wipe your environment, or something crashes, loading the RData will import any variables you've saved along the way. Establish an RData file with ---save("directory/file_name.RData")---. Use the `resave` function that I have in a few sections to save new variables to the file without overwriting anything. `load` will import the variables into your environment.

#load("/My Drive/ShortTermStress-IlluminaData/FullExp/Cryp-DE/data/import_rmd_vars.RData")
```

## 1.3 Growth data for short term cultures

We monitored growth of C. cryptica in freshwater over 7 days via chlorophyll-a fluorescence and cell counts. 

```{r}
#Import data
growth_cells <- read.delim("/My Drive/Downey_Manuscripts/Ccryptica_STS/Ccryptica_STS_files/Growth_cells.txt", header = TRUE, sep = '\t', fill = TRUE, row.names=NULL)
growth_RFU <- read.delim("/My Drive/Downey_Manuscripts/Ccryptica_STS/Ccryptica_STS_files/Growth_RFU.txt",header = TRUE, sep = '\t', fill = TRUE, row.names=NULL)

#Growth data from cell counts obtained using FlowCam
growth_cells_df <- pivot_longer(growth_cells[ ,c(1:13)], -c(Replicate,Trt), names_to = "timept")
growth_cells_df$timept <- ordered(growth_cells_df$timept, levels=c("t0m","t15m","t30m","t1h","t2h","t4h","t8h","t10h","t12h","t24h","t7d"))

growth_cells_df <- growth_cells_df %>% mutate(timept = case_when(
  growth_cells_df$timept == "t0m"  ~ 0,
  growth_cells_df$timept == "t15m" ~ 15,
  growth_cells_df$timept == "t30m" ~ 30,
  growth_cells_df$timept == "t1h" ~ 60,
  growth_cells_df$timept == "t2h" ~ 120,
  growth_cells_df$timept == "t4h" ~ 240,
  growth_cells_df$timept == "t8h" ~ 480,
  growth_cells_df$timept == "t10h" ~ 600,
  growth_cells_df$timept == "t12h" ~ 720,
  growth_cells_df$timept == "t24h" ~ 1440,
  growth_cells_df$timept == "t7d" ~ 10080
))

growth_cells_data <- growth_cells_df %>% group_by(timept,Trt) %>% summarize(sd = sd(value, na.rm = TRUE),value = mean(value), )

#Visualize
ggplot(growth_cells_data,  aes(x = timept, y = value, group = Trt, color = Trt, ymin = value-sd, ymax = value+sd))  + 
    geom_errorbar(width = 0.5) +
    geom_line(size = 1) +
    theme(axis.text.x = element_text(vjust=0, face = "bold"),
          axis.ticks.x=element_blank()) +
    coord_trans(y = "log") +
    xlab("Time in minutes") + 
    ylab("Cells per mL") +
    theme_classic() + 
    scale_color_manual(values=c("#112943", "#51b0fa")) +
    theme(plot.title = element_text(hjust = 0.5, face = "bold"))

#Growth data from relative fluorescent units measured with Trilogy
growth_RFU_df <- pivot_longer(growth_RFU[ ,c(1:13)], -c(Replicate,Trt), names_to = "timept")
growth_RFU_df$timept <- ordered(growth_RFU_df$timept, levels=c("t0m","t15m","t30m","t1h","t2h","t4h","t8h","t10h","t12h","t24h","t7d"))

growth_RFU_df <- growth_RFU_df %>% mutate(timept = case_when(
  growth_RFU_df$timept == "t0m" ~ 0,
  growth_RFU_df$timept == "t15m" ~ 15,
  growth_RFU_df$timept == "t30m" ~ 30,
  growth_RFU_df$timept == "t1h" ~ 60,
  growth_RFU_df$timept == "t2h" ~ 120,
  growth_RFU_df$timept == "t4h" ~ 240,
  growth_RFU_df$timept == "t8h" ~ 480,
  growth_RFU_df$timept == "t10h" ~ 600,
  growth_RFU_df$timept == "t12h" ~ 720,
  growth_RFU_df$timept == "t24h" ~ 1440,
  growth_RFU_df$timept == "t7d" ~ 10080
))

growth_RFU_data <- growth_RFU_df %>% group_by(timept,Trt) %>% summarize(sd = sd(value, na.rm = TRUE),value = mean(value), )

#Visualize
ggplot(growth_RFU_data,  aes(x = timept, y = value, group = Trt, color = Trt, ymin = value-sd, ymax = value+sd))  + 
    geom_errorbar(width = 0.5) +
    geom_line(size = 1) +
    theme(axis.text.x = element_text(vjust=0, face = "bold"),
          axis.ticks.x=element_blank()) +
    coord_trans(y = "log") +
    xlab("Time in minutes") + 
    ylab("Average growth (log)") +
    theme_classic() + 
    scale_color_manual(values=c("#112943", "#51b0fa")) +
    theme(plot.title = element_text(hjust = 0.5, face = "bold"))

#resave(growth_cells, growth_RFU, file = "/My Drive/ShortTermStress-IlluminaData/FullExp/Cryp-DE/data/import_rmd_vars.RData")
```

# 2. Bioinformatic analysis of RNA-seq reads
445,128,534 100bp paired-end reads recovered for C. cryptica

##  2.1 Quality control, adapter removal & trimming

1. We ran [FastQC](https://github.com/s-andrews/FastQC) on all fastq files:

```{r, engine = 'bash', eval = FALSE}
# run FastQC v0.11.5
fastqc *.fastq
```

2. Adapter removal and trimming of sequences was done using [Ktrim](https://github.com/hellosunking/Ktrim):

```{r, engine = 'bash', eval = FALSE}


for i in $(cat original_names.txt); do ktrim \
  -1 $i_L004_R1_001.fastq \
  -2 $i_L004_R2_001.fastq \
  -o $i\_ -t 15 -p 33 -q 20 -s 36\
  -a TACACTCTTTCCCTACACGACGCTCTTCCGATCT \
  -a GTGACTGGAGTTCAGACGTGTGCTCTTCCGATCT \
  -a TACACTCTTTCCCTACACGACGCTCTTCCGATCT \
  -a AGATCGGAAGAGCGTCGTSGTAGGGAAAGAGTGTA \
  -a GTGACTGGAGTTCAGACGTGTGCTCTTCCGATCT \
  -a AGATCGGAAGAGCACACGTCTGAACTCCAGTCAC \
  -m 0.5; done
```

The output files of the Ktrim run were checked again using FastQC:

```{r, engine = 'bash', eval = FALSE}
# run Fastqc v0.11.5
fastqc *_L004_R1_001.fastq
fastqc *_L004_R2_001.fastq
```

##2.2 STAR/2.7.3a genome mapping

```{r, engine = 'bash', eval = FALSE}
STAR --runThreadN 8 --runMode genomeGenerate \
  --genomeDir original_names/CCMP332_STARIndex \
  --genomeFastaFiles CCMP332_data/CCMP332_renamed.fasta \
  --sjdbGTFfile CCMP332_data/cryptica_pilon5.all.functional_ipr.functional_blast.renamed.gff \
  --sjdbGTFfeatureExon exon \
  --sjdbGTFtagExonParentTranscript Parent \
  --sjdbOverhang 99 --genomeSAindexNbases 12
#the --genomeSAindexNbases = 12 parameter was suggested after running STAR without this parameter. It is based on: min(1, log2(152305664)/2 - 1) with 152305664 being the length of the genome in bases
#--sjdbOverhang =  max read length - 1 = 100 - 1 = 99
```

We ran STAR

```{r, engine = 'bash', eval = FALSE}
  for i in $(cat original_names.txt); do STAR  \
  --runThreadN 15 --genomeDir CCMP332_STARIndex \
  --outSAMtype BAM SortedByCoordinate --outReadsUnmapped Fastx \
  --alignIntronMin 1 --alignIntronMax 22618 \
  --readFilesIn trimmed_CCMP332_seqs/$i\.read1.fq\
   trimmed_CCMP332_seqs/$i\.read2.fq \
  --outFileNamePrefix CCMP332_STAR_output/$i\_; done

```

##2.3 Read quantification with HTSeq on gene-level

We ran [HTSeq](https://htseq.readthedocs.io/en/master/count.html) as follows on a gff file without the fasta included at the end:

```{r, engine = 'bash', eval = FALSE}
for i in $(cat original_names.txt);do /share/apps/python/anaconda3-python-3.7.3/lib/python3.7/site-packages/HTSeq-0.11.3-py3.7-linux-x86_64.egg-info/scripts/htseq-count \
--format=bam \
--order=pos \
--stranded=reverse \
--minaqual=10 \
--type=gene \
--idattr=ID \
--mode=union \
--nonunique=none \
--samout=$i\_HTSeq.gene-lvl.out \
CCMP332_STAR_output/$i\_Aligned.sortedByCoord.out.bam \
CCMP332_noFASTA.gff \
>> $i\_HTSeq.gene-lvl.STDOUT;done
```

# 3. Statistical analysis of RNA-seq data in R

## 3.1 Model fitting in EdgeR

###3.1.1 Data import 

Import the count data for Downey et al. 2022 and Nakov et al. 2020 (the long-term hypo-osmotic growth study for *C. cryptica*). The Nakov et al. 2020 required re-analysis because the initial mapping used a de-novo transcriptome, rather than the annotated genome from Wade et al. 2021. We utilized only the CCMP332 data from the long-term study.

```{r}
#Data Import 
x_cnts <- read.table("/My Drive/ShortTermStress-IlluminaData/FullExp/Cryp-DE/CCMP332_HTSeq_output.txt", header = TRUE, row.names = 1)

nakov_cnts <- read.table("/My Drive/ShortTermStress-IlluminaData/FullExp/Cryp-DE/nakov_data/nakov_HTSeq_output.txt", header = FALSE, row.names = 1)

colnames(x_cnts) <- c("Cntl_R1","0m_R1","15m_R1","30m_R1","60m_R1","2hr_R1","4hr_R1","8hr_R1","10hr_R1",
                      "Cntl_R2","0m_R2","15m_R2","30m_R2","60m_R2","2hr_R2","4hr_R2","8hr_R2","10hr_R2",
                      "Cntl_R3","0m_R3","15m_R3","30m_R3","60m_R3","2hr_R3","4hr_R3","8hr_R3","10hr_R3")

colnames(nakov_cnts) <- c("R1_0","R1_24","R2_0","R2_24","R3_0","R3_24")

x_cnts = x_cnts[-c(21251:21254), ] #drop the last lines that do not contain information pertinent to gene counts

total_gene_number = nrow(x_cnts) #21250

#resave(x_cnts,nakov_cnts, file = "/My Drive/ShortTermStress-IlluminaData/FullExp/Cryp-DE/data/import_rmd_vars.RData")
```

EdgeR works with a DGEList data class object. This needed to be created using the count data and a group object that contains information on the different groups. We also developed multidimensional scale plot including the mock control ASW 24 treatment to look at data similarity. 

```{r}
#----All replicates without Control----#
# We removed the  because initial comparisons showed no significant difference
group_cntl <- c("Cntl_R1","0m","15m","30m","60m","2hr","4hr","8hr","10hr",
           "Cntl_R2","0m","15m","30m","60m","2hr","4hr","8hr","10hr",
           "Cntl_R3","0m","15m","30m","60m","2hr","4hr","8hr","10hr")

y_list_cntl <- DGEList(counts=as.matrix(x_cnts),group=group_cntl)
```

### 3.1.2 Data filtering 

Here, the genes that have very low counts across all the libraries were removed. Filtering was done using the CPM (count per million). Here, we retained all the genes that have least one CPM in at least three samples:

```{r}
keep <- rowSums(cpm(y_list_cntl)>1)>=3 #keep genes that have a least one count per million in at least three samples
y_list_cntl <- y_list_cntl[keep,]
y_list_cntl[,]
y_list_cntl$samples$lib.size <- colSums(y_list_cntl$counts)
num_kept_gene = nrow(y_list_cntl$counts)
```

### 3.1.3 Data normalization

Next, we calculated a set of normalization factors (one for each sample) to eliminate composition biases between libraries:

```{r}
y_list_cntl <- calcNormFactors(y_list_cntl, method = 'TMM') #normalizes for RNA composition (highly expressed genes)
head(y_list_cntl$samples)
plotMDS(y_list_cntl, top = 500)
```

Based on similarity of the mock control counts to that of the 0 minute control, we removed the "Cntl" samples and repeated the above steps. Further analyses used this subset. 

```{r}
#MDS plot without mock control data
gene_cnts <-  subset(x_cnts, select=-c(Cntl_R1,Cntl_R2,Cntl_R3))
group <- c("0m","15m","30m","60m","2hr","4hr","8hr","10hr",
          "0m","15m","30m","60m","2hr","4hr","8hr","10hr",
          "0m","15m","30m","60m","2hr","4hr","8hr","10hr")

y_list <- DGEList(counts=as.matrix(gene_cnts),group=group)
keep <- rowSums(cpm(y_list)>1)>=3 #keep genes that have a least one count per million in at least three samples
y_list <- y_list[keep,]
y_toss <- y_list[,]
y_list$samples$lib.size <- colSums(y_list$counts)
num_kept_gene = nrow(y_list$counts)
y_list <- calcNormFactors(y_list, method = 'TMM') #normalizes for RNA composition (highly expressed genes)
head(y_list$samples)
plotMDS(y_list, top = 500)

#resave(gene_cnts,y_list, file = "/My Drive/ShortTermStress-IlluminaData/FullExp/Cryp-DE/data/import_rmd_vars.RData")
```

We repeated the process with the Nakov et al. 2020 data

```{r}
#Setting it all up with the C. cryptica long term data
nakov_332 <- subset(nakov_cnts, select=c(R2_0,R2_24))
group_nakov <- c("0", "24","0", "24","0", "24")
group_332 <- c("0", "24")
y_list_nakov <- DGEList(counts=as.matrix(nakov_332),group=group_332)
keep_nakov <- rowSums(cpm(y_list_nakov)>1)>=2 #keep genes that have a least one count per million in at least three samples
y_list_nakov <- y_list_nakov[keep_nakov,]
dim(y_list_nakov)
y_list_nakov$samples$lib.size <- colSums(y_list_nakov$counts)
num_kept_gene = nrow(y_list_nakov$counts)
y_list_nakov <- calcNormFactors(y_list_nakov, method = 'TMM') #normalizes for RNA composition (highly expressed genes)
head(y_list_nakov$samples)

#resave(nakov_332,y_list_nakov, file = "/My Drive/ShortTermStress-IlluminaData/FullExp/Cryp-DE/data/import_rmd_vars.RData")
```

We calculated Poisson distances for both sets of data and visualized with pheatmap

```{r fig.width=10, fig.height=8, fig.align="center"}
poisd <- PoissonDistance(t(y_list$counts))
poisd_nakov <- PoissonDistance(t(y_list_nakov$counts))

#define colors to be used in the plot
colors_heatmap <- colorRampPalette(rev(brewer.pal(9,"Blues")))(255)

#plot a heatmap
samplePoisDistMatrix <- as.matrix(poisd$dd)
rownames(samplePoisDistMatrix) <- paste(y_list$samples$group)
colnames(samplePoisDistMatrix) <- paste(y_list$samples$group)
pheatmap(samplePoisDistMatrix,
         clustering_distance_rows=poisd$dd,
         clustering_distance_cols=poisd$dd,
         col=colors_heatmap,cex=1)#calculate poisson distances for the normalized count data
#C. cryptica longterm data
samplePoisDistMatrix <- as.matrix(poisd_nakov$dd)
rownames(samplePoisDistMatrix) <- paste(y_list_nakov$samples$group)
colnames(samplePoisDistMatrix) <- paste(y_list_nakov$samples$group)
pheatmap(samplePoisDistMatrix,
         clustering_distance_rows=poisd_nakov$dd,
         clustering_distance_cols=poisd_nakov$dd,
         col=colors_heatmap,cex=1)#calculate poisson distances for the normalized count data

```

### 3.1.4 Design matrix and dispersion estimation

Prior to differential expression analysis, dispersion needs to be estimated from a data matrix. edgeR uses the negative binomial (NB) distribution to model the read counts for each gene in each sample. The dispersion parameter of the NB distribution accounts for variability between biological replicates. edgeR estimates an empirical Bayes moderated dispersion for each individual gene. It also estimates a common dispersion, which is a global dispersion estimate averaged over all genes, and a trended dispersion where the dispersion of a gene is predicted from its abundance.

```{r fig.width=10, fig.height=5, fig.align="center"}
#Created a design matrix for future data contrasts

design <- model.matrix(~0+group, data = y_list$samples)
colnames(design) <- levels(y_list$samples$group)

#Dispersion estimation & model fitting
y_list <- estimateDisp(y_list, design)
plotBCV(y_list)

#Design matrix for long-term data

design_nakov <- model.matrix(~0+group_nakov, data = y_list_nakov$samples)
colnames(design_nakov) <- levels(y_list_nakov$samples$group)

#resave(design, design_nakov, file = "/My Drive/ShortTermStress-IlluminaData/FullExp/Cryp-DE/data/import_rmd_vars.RData")
```

### 3.1.5 Model fitting

glmQLFit is a quasi-likelihood (QL) method that accounts for gene-specific variability from both biological and technical sources. Under the QL framework, the NB dispersion trend is used to describe the overall biological variability across all genes, and gene-specific variability above and below the overall level is picked up by the QL dispersion. In the QL approach, the individual (tagwise) NB dispersions are not used.

```{r fig.width=10, fig.height=5, fig.align="center"}
fit_group_model <- glmQLFit(y_list, design, robust = TRUE)
head(fit_group_model$coefficients)
plotQLDisp(fit_group_model)

#C. cryptica longterm data
DE_nakov <- exactTest(y_list_nakov,dispersion = 0.16) #dispersion assigned based on exactTest author recommendation
#For future comparison, we used a subset of the long-term data. Due to the last replication for the CCMP332 expression data, we imposed a strenuous LFC filter. 
nakov_DE_dr <- subset(DE_nakov$table, (`logFC`) <= -1) #downregulated genes
nakov_DE_ur <- subset(DE_nakov$table, (`logFC`) >= 1) #upregulated genes

nakov_DE <- rbind(nakov_DE_dr,nakov_DE_ur)

#resave(nakov_DE, fit_group_model, file = "/My Drive/ShortTermStress-IlluminaData/FullExp/Cryp-DE/data/import_rmd_vars.RData")
```

## 3.2 Testing for differential expression using stage-wise analysis:

### 3.2.1 Defining contrasts to test

We contrasted all timepoints against the 0 minute timepoint
```{r}
Contrasts=matrix(0,nrow = ncol(fit_group_model$coefficients),ncol=7)
rownames(Contrasts)=colnames(fit_group_model$coefficients)
colnames(Contrasts)=c("C0m-C15m", "C0m-C30m", "C0m-C60m", "C0m-C2hr", "C0m-C4hr", "C0m-C8hr", "C0m-C10hr")

Contrasts[c("0m", "15m"), "C0m-C15m"]=c(1,-1)
Contrasts[c("0m", "30m"), "C0m-C30m"]=c(1,-1)
Contrasts[c("0m", "60m"), "C0m-C60m"]=c(1,-1)
Contrasts[c("0m", "2hr"), "C0m-C2hr"]=c(1,-1)
Contrasts[c("0m", "4hr"), "C0m-C4hr"]=c(1,-1)
Contrasts[c("0m", "8hr"), "C0m-C8hr"]=c(1,-1)
Contrasts[c("0m", "10hr"), "C0m-C10hr"]=c(1,-1)

#resave(Contrasts, file = "/My Drive/ShortTermStress-IlluminaData/FullExp/Cryp-DE/data/import_rmd_vars.RData")
```

### 3.2.2 Stage-wise testing 

We performed the stage-wise testing procedure in [stageR](https://bioconductor.org/packages/release/bioc/html/stageR.html). StageR allows for simultaneous FDR control in all the contrasts, and consists of two steps: the screening stage, and the confirmation stage.

The screening stage tested whether any of the contrasts were significant. The screening stage gives P-values as output, but these are not yet FDR-controlled so should not be used in downstream analyses.

```{r}
#screening stage
alpha=0.01 
screenTest <- glmQLFTest(fit_group_model, contrast=Contrasts)
pScreen <- screenTest$table$PValue
names(pScreen) <- rownames(screenTest$table)
```

The screening stage was followed by the confirmation stage. In the confirmation stage, every contrast was assessed separately. The confirmation stage P-values were adjusted to control the FWER across the hypotheses within a gene and are subsequently corrected to the BH-adjusted significance level of the screening stage. This allowed for a direct comparison of the adjusted P-values to the provided significance level alpha for both screening and confirmation stage adjusted P-values. Here, we used the holm method for correction of the P-values.

```{r}
#confirmation stage
confirmationResults <- sapply(1:ncol(Contrasts),function(i) glmQLFTest(fit_group_model, 
                              contrast = Contrasts[,i]), simplify=FALSE) #calculates F-test for each contrast
confirmationPList <- lapply(confirmationResults, 
                            function(x) x$table$PValue) #takes the p-values from all genes for each contrast and puts them in a list
confirmationP <- as.matrix(Reduce(f=cbind,confirmationPList)) 
rownames(confirmationP) <- rownames(confirmationResults[[1]]$table)
colnames(confirmationP) <- colnames(Contrasts)
stageRObj <- stageR(pScreen=pScreen, pConfirmation=confirmationP) #constructs an object
stageRAdj <- stageWiseAdjustment(object=stageRObj, 
                                 method="holm", alpha=0.01) #adjusts the P-values using FWER correction using the holm method
res <- getResults(stageRAdj)

#Number of differentially expressed genes in every contrast
SignifGenes <- colSums(res)

#Establish adjusted p-values
adjusted_p <- getAdjustedPValues(stageRAdj, onlySignificantGenes = FALSE, order = FALSE)
padj_df <- as.data.frame(adjusted_p) #establish as a dataframe for use downstream
```

Upon finishing the stage-wise testing procedure, we checked the number of significant genes:

```{r}
#visualize the number of significant genes
res_df = as.data.frame(res) #res is a matrix, to be able to subset, change it to a dataframe
res_df2 = res_df
res_df2$gene = rownames(res_df2)
OnlySignGenes = res_df2[res_df2$padjScreen == 1,] #removes rows for which global test was non significant
dim(OnlySignGenes) #10569 significant genes based on stringent alpha = 0.01

genesSI <- rownames(adjusted_p)[adjusted_p[,"padjScreen"]<=0.01] 
length(genesSI) #10569 unique genes
genesNotFoundStageII <- genesSI[genesSI %in% rownames(res)[rowSums(res==0)==8]]
length(genesNotFoundStageII) #There are 0 genes found to be significant in screening, but not confirmation

#resave(adjusted_p, OnlySignGenes, genesNotFoundStageII, file = "/My Drive/ShortTermStress-IlluminaData/FullExp/Cryp-DE/data/import_rmd_vars.RData")
```

We only included genes that were significant after the confirmation stage
```{r}
OnlySignGenes_ConStage <- OnlySignGenes [!rownames(OnlySignGenes) %in% genesNotFoundStageII, ]
nrow(OnlySignGenes_ConStage) #10569 rows
```

### 3.2.3 Summarize the results for downstream analyses

```{r}
#get information on total number of expressed genes = 12939
expressed_genes = nrow(fit_group_model$fitted.values)
expressed_genes

#relative number of expressed genes in the genome of C. cryptica = 60.89 %
rel_exp_genes = expressed_genes / total_gene_number
rel_exp_genes

#get information on totaly number of DE expressed genes = 10569
DE_expressed_genes = nrow(OnlySignGenes_ConStage)
DE_expressed_genes

#relative number of DE expressed genes in the genome of C. cryptica = 81.68% %
rel_DEexp_genes = DE_expressed_genes  / expressed_genes 
rel_DEexp_genes
```

Before we continued with the downstream analyses, we created a single data object that contains some key-information of the statistical pipeline outlined above. This included information on logFC, logCPM and P-values for each gene for each contrast.

First, we selected the FDR adjusted P-values for each contrast using the output of the stageR screening stage:
```{r}
colnames(adjusted_p)[1] = "padjScreen"
colnames(adjusted_p)[2] = "C0mvsC15m_Padj"
colnames(adjusted_p)[3] = "C0mvsC30m_Padj"
colnames(adjusted_p)[4] = "C0mvsC60m_Padj"
colnames(adjusted_p)[5] = "C0mvsC2hr_Padj"
colnames(adjusted_p)[6] = "C0mvsC4hr_Padj"
colnames(adjusted_p)[7] = "C0mvsC8hr_Padj"
colnames(adjusted_p)[8] = "C0mvsC10hr_Padj"

#resave(OnlySignGenes_ConStage, adjusted_p_onlysign, file = "/My Drive/ShortTermStress-IlluminaData/FullExp/Cryp-DE/data/import_rmd_vars.RData")
```

Second, we extracted the information on logFC, logCPM, F value and non-adjusted P-values from the *confirmationResults* object:

```{r}
#create empty list to hold the data values
datalist = list()

#loop over the confirmationResults object to obtain the relevant information (table)
for (contrast in c(1,2,3,4,5,6,7)){
  table = confirmationResults[[contrast]]$table
  datalist[[contrast]] <- table
}

#turn list into data frame
confirmationResults_total_dataset = cbind(datalist[[1]],datalist[[2]],datalist[[3]],datalist[[4]]
                                               ,datalist[[5]],datalist[[6]],datalist[[7]])

#rename column names for tracability
#logCPM tells us about the overall presence of that gene in the entire dataset, which is why it's identical across the board. Has nothing to do with contrasts
colnames(confirmationResults_total_dataset)=c("C0mvsC15m_logFC","C0mvsC15m_logCPM","C0mvsC15m_F","C0mvsC15m_nonadj_PValue","C0mvsC30m_logFC","C0mvsC30m_logCPM","C0mvsC30m_F","C0mvsC30m_nonadj_PValue","C0mvsC60m_logFC","C0mvsC60m_logCPM","C0mvsC60m_F","C0mvsC60m_nonadj_PValue","C0mvsC2hr_logFC","C0mvsC2hr_logCPM","C0mvsC2hr_F","C0mvsC2hr_nonadj_PValue","C0mvsC4hr_logFC","C0mvsC4hr_logCPM","C0mvsC4hr_F","C0mvsC4hr_nonadj_PValue","C0mvsC8hr_logFC","C0mvsC8hr_logCPM","C0mvsC8hr_F","C0mvsC8hr_nonadj_PValue","C0mvsC10hr_logFC","C0mvsC10hr_logCPM","C0mvsC10hr_F","C0mvsC10hr_nonadj_PValue")
```

Then we combined the FDR-adjusted P-values and the table with information on logFC etc. into a single data frame:

```{r}
#merge the data frames
table = merge(confirmationResults_total_dataset,adjusted_p, by = 0, all = TRUE)
all_results <- table[,-1]

#use the first column (gene names) for the row names
rownames(all_results) <- table[,1]
all_results <- (na.omit(all_results))
head(all_results)
#resave(OnlySignGenes_ConStage, confirmationResults_total_dataset, table, file = "/My Drive/ShortTermStress-IlluminaData/FullExp/Cryp-DE/data/import_rmd_vars.RData")
```

The resulting data frame *all_results* was used in multiple analyses downstream to access basic stats on each gene for each contrast.


### 3.2.4 Subset to genes DE at 2 timepoints

In order to focus on genes most likely to play a significant role in the short-term hypo-osmotic stress response, we limited our dataset to those genes that were significant at two consecutive timepoints in the same direction (i.e., both up/downregulated).

```{r fig.width=20, fig.height= 15}
df <- res_df2[,-c(1,9)]
colSums(df)
keep_2_SignDEG <- df[rowSums(df[-1] & df[-ncol(df)]) > 0,]
keep2_colSums <- colSums(keep_2_SignDEG)
DEG2sig_names = rownames(keep_2_SignDEG)

DEG2sig_results <- (na.omit(all_results[DEG2sig_names,]))

sign2LFC <- subset(DEG2sig_results, select=c("C0mvsC15m_logFC","C0mvsC15m_Padj","C0mvsC30m_logFC","C0mvsC30m_Padj",
                                                            "C0mvsC60m_logFC","C0mvsC60m_Padj","C0mvsC2hr_logFC","C0mvsC2hr_Padj",
                                                            "C0mvsC4hr_logFC","C0mvsC4hr_Padj","C0mvsC8hr_logFC","C0mvsC8hr_Padj",
                                                            "C0mvsC10hr_logFC","C0mvsC10hr_Padj"))
colnames(sign2LFC) <- c("0v15m","0v15m_Padj","0v30m","0v30m_Padj","0v1h","0v1h_Padj","0v2h","0v2h_Padj",
                             "0v4h","0v4h_Padj","0v8h","0v8h_Padj","0v10h","0v10h_Padj")
```

Plotted the heatmap of the significant data with a gene expression data matrix

```{r fig.width=20, fig.height= 15}
sign2LFC_mat <- as.matrix(sign2LFC %>% dplyr::select(`0v15m`,`0v30m`,`0v1h`,`0v2h`,`0v4h`,`0v8h`,`0v10h`))
min(sign2LFC_mat) #-9.470548
max(sign2LFC_mat) #7.635282

paletteLength <- 500
myColor <- colorRampPalette(c("blue", "snow", "red"))(paletteLength)
sign2LFC_hmp <- heatmap.2(sign2LFC_mat,dendrogram = "row",trace="none",col=myColor, labRow=TRUE, cexCol = 1, Colv = FALSE)

#resave(all_results, sign2LFC, file = "/My Drive/ShortTermStress-IlluminaData/FullExp/Cryp-DE/data/import_rmd_vars.RData")
```

We also looked at the non-significant genes as a quick verification

```{r fig.width=20, fig.height= 15}
#The genes that are not differentially expressed
all_sig_names <- rownames(all_results)
pScreen_df <- as.data.frame(pScreen)
pScreen_df$cluster <- "NotSig"
not_sig_genes <- (na.omit(pScreen_df[!row.names(pScreen_df)%in%all_sig_names,]))
not_sig_genes <- subset(not_sig_genes, select=c("cluster"))

NS_LFC_results <- (na.omit(all_results[!row.names(all_results)%in%DEG2sig_names,]))
NS_LFC_df <- subset(NS_LFC_results, select=c("C0mvsC15m_logFC","C0mvsC15m_Padj","C0mvsC30m_logFC","C0mvsC30m_Padj",
                                                "C0mvsC60m_logFC","C0mvsC60m_Padj","C0mvsC2hr_logFC","C0mvsC2hr_Padj",
                                                "C0mvsC4hr_logFC","C0mvsC4hr_Padj","C0mvsC8hr_logFC","C0mvsC8hr_Padj",
                                                "C0mvsC10hr_logFC","C0mvsC10hr_Padj"))
colnames(NS_LFC_df) <- c("0v15m","0v15m_Padj","0v30m","0v30m_Padj","0v1h","0v1h_Padj","0v2h","0v2h_Padj",
                       "0v4h","0v4h_Padj","0v8h","0v8h_Padj","0v10h","0v10h_Padj")

#Rows of significant and non-significant dataframes add up to 11528 as expected. No genes missing

NS_LFC_mat <- as.matrix(NS_LFC_df %>% dplyr::select(`0v15m`,`0v30m`,`0v1h`,`0v2h`,`0v4h`,`0v8h`,`0v10h`))

min(NS_LFC_mat) #-9.452026
max(NS_LFC_mat) #7.762832

paletteLength <- 500
myColor <- colorRampPalette(c("blue", "snow", "red"))(paletteLength)
NS_LFC_hmp <- heatmap.2(NS_LFC_mat,dendrogram = "row",trace="none",col=myColor, labRow=TRUE, cexCol = 1, Colv = FALSE)

```

### 3.2.5 Graph of genes up and downregulated at each timepoint

```{r fig.width=10, fig.height=6}

sign_0v15m <- rownames(keep_2_SignDEG)[keep_2_SignDEG[,"C0m-C15m"]==1] 
length(sign_0v15m) 
sign2LFC_0v15m <- sign2LFC [rownames(sign2LFC) %in% sign_0v15m, ]
logFCup_0v15m <- subset(sign2LFC_0v15m, (`0v15m`) > 0) 
logFCdown_0v15m <- subset(sign2LFC_0v15m, (`0v15m`) < 0)

sign_0v30m <- rownames(keep_2_SignDEG)[keep_2_SignDEG[,"C0m-C30m"]==1] 
length(sign_0v30m) 
sign2LFC_0v30m <- sign2LFC [rownames(sign2LFC) %in% sign_0v30m, ]
logFCup_0v30m <- subset(sign2LFC_0v30m, (`0v30m`) > 0) 
logFCdown_0v30m <- subset(sign2LFC_0v30m, (`0v30m`) < 0)

sign_0v1h <- rownames(keep_2_SignDEG)[keep_2_SignDEG[,"C0m-C60m"]==1] 
length(sign_0v1h) 
sign2LFC_0v1h <- sign2LFC [rownames(sign2LFC) %in% sign_0v1h, ]
logFCup_0v1h <- subset(sign2LFC_0v1h, (`0v1h`) > 0) 
logFCdown_0v1h <- subset(sign2LFC_0v1h, (`0v1h`) < 0)

sign_0v2h <- rownames(keep_2_SignDEG)[keep_2_SignDEG[,"C0m-C2hr"]==1] 
length(sign_0v2h) 
sign2LFC_0v2h <- sign2LFC [rownames(sign2LFC) %in% sign_0v2h, ]
logFCup_0v2h <- subset(sign2LFC_0v2h, (`0v2h`) > 0) 
logFCdown_0v2h <- subset(sign2LFC_0v2h,  (`0v2h`) < 0)

sign_0v4h <- rownames(keep_2_SignDEG)[keep_2_SignDEG[,"C0m-C4hr"]==1] 
length(sign_0v4h) 
sign2LFC_0v4h <- sign2LFC [rownames(sign2LFC) %in% sign_0v4h, ]
logFCup_0v4h <- subset(sign2LFC_0v4h, (`0v4h`) > 0) 
logFCdown_0v4h <- subset(sign2LFC_0v4h,  (`0v4h`) < 0)

sign_0v8h <- rownames(keep_2_SignDEG)[keep_2_SignDEG[,"C0m-C8hr"]==1] 
length(sign_0v8h) 
sign2LFC_0v8h <- sign2LFC [rownames(sign2LFC) %in% sign_0v8h, ]
logFCup_0v8h <- subset(sign2LFC_0v8h, (`0v8h`) > 0) 
logFCdown_0v8h <- subset(sign2LFC_0v8h, (`0v8h`) < 0)

sign_0v10h <- rownames(keep_2_SignDEG)[keep_2_SignDEG[,"C0m-C10hr"]==1] 
length(sign_0v10h) 
sign2LFC_0v10h <- sign2LFC [rownames(sign2LFC) %in% sign_0v10h, ]
logFCup_0v10h <- subset(sign2LFC_0v10h, (`0v10h`) > 0) 
logFCdown_0v10h <- subset(sign2LFC_0v10h, (`0v10h`) < 0)

up <- c(584, 1555, 1364, 664, 540, 763, 659)
down <- c(-1029, -1942, -2048, -1498, -883, -561, -772)
timept <- c("t15m","t30m","t1h","t2h","t4h","t8h","t10h","t15m","t30m","t1h","t2h","t4h","t8h","t10h")
level_order = c("t15m","t30m","t1h","t2h","t4h","t8h","t10h")
dat <- data.frame(
  group = rep(c("upregulated", "downregulated"), each=7),
  x = timept,
  y = c(up, down)
)

dat$x <- as.character(dat$x)
dat$x <- factor(dat$x, levels=(c("t15m","t30m","t1h","t2h","t4h","t8h","t10h")))
ggplot(dat, aes(x=x, y=y, fill=group)) + 
  geom_bar(stat="identity", position="identity", width=0.5) +
  scale_fill_manual(values=c("#28db69","#a9acff")) +
  scale_x_discrete(timept) +
  scale_y_continuous(breaks=seq(-2500,2500,500), limits=c(-2200,2200)) +
  theme_classic() +
  theme(axis.text.x=element_text(size=14), axis.text.y=element_text(size=14), legend.text=element_text(size=14)) +
  ylab("Total number of DE genes") +
  xlab("Time exposed to 0 ppt ASW") 
```

### 3.2.6 Unique and shared DE between timepoints

We created variables for the shared and unique sets of DE genes

```{r}
#calculate number of DE genes for each contrast
allDE_C0mvsC15m = length(c(rownames (subset (OnlySignGenes_ConStage, OnlySignGenes_ConStage$`C0m-C15m`== 1)))) #2442
allDE_C0mvsC30m = length(c(rownames (subset (OnlySignGenes_ConStage, OnlySignGenes_ConStage$`C0m-C30m`== 1)))) #4900
allDE_C0mvsC60m = length(c(rownames (subset (OnlySignGenes_ConStage, OnlySignGenes_ConStage$`C0m-C60m`== 1)))) #4453
allDE_C0mvsC2hr = length(c(rownames (subset (OnlySignGenes_ConStage, OnlySignGenes_ConStage$`C0m-C2hr`== 1)))) #2533
allDE_C0mvsC4hr = length(c(rownames (subset (OnlySignGenes_ConStage, OnlySignGenes_ConStage$`C0m-C4hr`== 1)))) #1848
allDE_C0mvsC8hr = length(c(rownames (subset (OnlySignGenes_ConStage, OnlySignGenes_ConStage$`C0m-C8hr`== 1)))) #1963
allDE_C0mvsC10hr = length(c(rownames (subset (OnlySignGenes_ConStage, OnlySignGenes_ConStage$`C0m-C10hr`== 1)))) #2031

#calculate number of DE genes unique within in a contrast within a genotype/average response
UniqueSubDE_C0mvsC15m = length(c(rownames (subset (OnlySignGenes_ConStage, OnlySignGenes_ConStage$`C0m-C15m` == 1 
                                                   & OnlySignGenes_ConStage$`C0m-C30m` == 0 & OnlySignGenes_ConStage$`C0m-C60m` == 0 
                                                   & OnlySignGenes_ConStage$`C0m-C2hr` == 0 & OnlySignGenes_ConStage$`C0m-C4hr` == 0 
                                                   & OnlySignGenes_ConStage$`C0m-C8hr` == 0 & OnlySignGenes_ConStage$`C0m-C10hr` == 0)))) #334
UniqueSubDE_C0mvsC30m = length(c(rownames (subset (OnlySignGenes_ConStage, OnlySignGenes_ConStage$`C0m-C15m` == 0 
                                                   & OnlySignGenes_ConStage$`C0m-C30m` == 1 & OnlySignGenes_ConStage$`C0m-C60m` == 0 
                                                   & OnlySignGenes_ConStage$`C0m-C2hr` == 0 & OnlySignGenes_ConStage$`C0m-C4hr` == 0 
                                                   & OnlySignGenes_ConStage$`C0m-C8hr` == 0 & OnlySignGenes_ConStage$`C0m-C10hr` == 0)))) #1014
UniqueSubDE_C0mvsC60m = length(c(rownames (subset (OnlySignGenes_ConStage, OnlySignGenes_ConStage$`C0m-C15m` == 0 
                                                   & OnlySignGenes_ConStage$`C0m-C30m` == 0 & OnlySignGenes_ConStage$`C0m-C60m` == 1 
                                                   & OnlySignGenes_ConStage$`C0m-C2hr` == 0 & OnlySignGenes_ConStage$`C0m-C4hr` == 0 
                                                   & OnlySignGenes_ConStage$`C0m-C8hr` == 0 & OnlySignGenes_ConStage$`C0m-C10hr` == 0)))) #740
UniqueSubDE_C0mvsC2hr = length(c(rownames (subset (OnlySignGenes_ConStage, OnlySignGenes_ConStage$`C0m-C15m` == 0 
                                                   & OnlySignGenes_ConStage$`C0m-C30m` == 0 & OnlySignGenes_ConStage$`C0m-C60m` == 0 
                                                   & OnlySignGenes_ConStage$`C0m-C2hr` == 1 & OnlySignGenes_ConStage$`C0m-C4hr` == 0 
                                                   & OnlySignGenes_ConStage$`C0m-C8hr` == 0 & OnlySignGenes_ConStage$`C0m-C10hr` == 0)))) #110
UniqueSubDE_C0mvsC4hr = length(c(rownames (subset (OnlySignGenes_ConStage, OnlySignGenes_ConStage$`C0m-C15m` == 0 
                                                   & OnlySignGenes_ConStage$`C0m-C30m` == 0 & OnlySignGenes_ConStage$`C0m-C60m` == 0 
                                                   & OnlySignGenes_ConStage$`C0m-C2hr` == 0 & OnlySignGenes_ConStage$`C0m-C4hr` == 1 
                                                   & OnlySignGenes_ConStage$`C0m-C8hr` == 0 & OnlySignGenes_ConStage$`C0m-C10hr` == 0)))) #169
UniqueSubDE_C0mvsC8hr = length(c(rownames (subset (OnlySignGenes_ConStage, OnlySignGenes_ConStage$`C0m-C15m` == 0 
                                                   & OnlySignGenes_ConStage$`C0m-C30m` == 0 & OnlySignGenes_ConStage$`C0m-C60m` == 0 
                                                   & OnlySignGenes_ConStage$`C0m-C2hr` == 0 & OnlySignGenes_ConStage$`C0m-C4hr` == 0 
                                                   & OnlySignGenes_ConStage$`C0m-C8hr` == 1 & OnlySignGenes_ConStage$`C0m-C10hr` == 0)))) #303
UniqueSubDE_C0mvsC10hr = length(c(rownames (subset (OnlySignGenes_ConStage, OnlySignGenes_ConStage$`C0m-C15m` == 0 
                                                    & OnlySignGenes_ConStage$`C0m-C30m` == 0 & OnlySignGenes_ConStage$`C0m-C60m` == 0 
                                                    & OnlySignGenes_ConStage$`C0m-C2hr` == 0 & OnlySignGenes_ConStage$`C0m-C4hr` == 0 
                                                    & OnlySignGenes_ConStage$`C0m-C8hr` == 0 & OnlySignGenes_ConStage$`C0m-C10hr` == 1)))) #200


DE_info = as.vector(cbind(allDE_C0mvsC15m,UniqueSubDE_C0mvsC15m,
                               allDE_C0mvsC30m,UniqueSubDE_C0mvsC30m,
                               allDE_C0mvsC60m,UniqueSubDE_C0mvsC60m,
                               allDE_C0mvsC2hr,UniqueSubDE_C0mvsC2hr,
                               allDE_C0mvsC4hr,UniqueSubDE_C0mvsC4hr,
                               allDE_C0mvsC8hr,UniqueSubDE_C0mvsC8hr,
                               allDE_C0mvsC10hr,UniqueSubDE_C0mvsC10hr))


rownames = as.vector(cbind("allDE_C0mvsC15m","UniqueSubDE_C0mvsC15m",
                                "allDE_C0mvsC30m","UniqueSubDE_C0mvsC30m",
                                "allDE_C0mvsC60m","UniqueSubDE_C0mvsC60m",
                                "allDE_C0mvsC2hr","UniqueSubDE_C0mvsC2hr",
                                "allDE_C0mvsC4hr","UniqueSubDE_C0mvsC4hr",
                                "allDE_C0mvsC8hr","UniqueSubDE_C0mvsC8hr",
                                "allDE_C0mvs10hr","UniqueSubDE_C0mvsC10hr"))

DE_colors = rep(c("slategray1","steelblue4","gray80","gray55","sandybrown","chocolate4",
                       "slategray1","steelblue4","gray80","gray55","sandybrown","chocolate4",
                       "slategray1","steelblue4"))
table  <- as.data.frame(cbind(rownames,DE_info,DE_colors ))                        
DE_info = table[,-1]
rownames(DE_info) = table[,1]
DE_info$DE_info_rel  <- (as.numeric(as.character(DE_info$DE_info)) / DE_expressed_genes) * 100 

```

Then we plotted the total and unique DE genes at each timepoint

``` {r fig.width=10, fig.height=5, fig.align="center"}
par(mfrow=c(1,2))

colsums = colSums(subset(keep_2_SignDEG, select=c("C0m-C15m", "C0m-C30m", "C0m-C60m", "C0m-C2hr", "C0m-C4hr", "C0m-C8hr", "C0m-C10hr")))
par(las=2)
par(mar = c(6, 5, 3, 1), xpd = TRUE)
barplot(colsums,col = "black",
        main = "# DEG in contrasts", ylab = "Total number of DE genes", 
        border=NA, cex.axis=0.9, cex.names = 0.9, cex.lab=1, font.lab=2, ylim=c(0,4000)) #plot barplot


par(mar = c(5, 4, 4, 8), xpd = TRUE)
space<-c(0,0,1,0,1,0,1,0,1,0,1,0,1,0) #create spaces between bars

barplot(DE_info$DE_info_rel,col = as.vector(DE_info$DE_colors), space=space, xaxt='n', 
        main = "% DEG over time", ylab = "Percentage of DE genes (%)", xlab = "Exposure time points",
        border=NA, cex.axis=0.9, cex.lab=1, font.lab=1,ylim=c(0,50)) #plot barplot

legend("topleft",inset=c(1,0),
       c(rows), c( "Total DE genes [0-15m]", "Unique DE genes [0-15m]",
                   "Total DE genes [0-30m]", "Unique DE genes [0-30m]", 
                   "Total DE genes [0-60m]", "Unique DE genes [0-60m]", 
                   "Total DE genes [0-2hr]", "Unique DE genes [0-2hr]",
                   "Total DE genes [0-4hr]", "Unique DE genes [0-4hr]",
                   "Total DE genes [0-8hr]", "Unique DE genes [0-8hr]",
                   "Total DE genes [0-10hr]", "Unique DE genes [0-10hr]"),
       fill = c("slategray1","steelblue4","gray80","gray55","sandybrown","chocolate4"), 
       bty = "n", border=NA, cex=0.5) 
mtext('15 min', side=1, line=0.45, at=1, cex = 0.8)
mtext('30 min', side=1, line=0.45, at=4, cex = 0.8)
mtext('1 hour', side=1, line=0.45, at=7, cex = 0.8)
mtext('2 hour', side=1, line=0.45, at=10, cex = 0.8)
mtext('4 hour', side=1, line=0.45, at=13, cex = 0.8)
mtext('8 hour', side=1, line=0.45, at=16, cex = 0.8)
mtext('10 hour', side=1, line=0.45, at=19, cex = 0.8)
```

# 4. Gene enrichment analyses of: 
## 4.1 Full dataset

Gene enrichment analyses for the entire dataset with GO terms from the GFF file. This does not account for up or down regulation, it's just a bulk overview.

```{r, echo=TRUE, message=FALSE, warning=FALSE, results='hide'}

#GO enrichment analyses: ORA (over-represetntation analysis) [topGO] for all genes
geneID2GO = readMappings(file = "/My Drive/ShortTermStress-IlluminaData/FullExp/Cryp-DE/CCMP332_GFF_GO.txt")
geneUniverse <- names(geneID2GO)
length(geneUniverse) #6309
#Example search
GOtoMatch = c("GO:0033036")
head(names(geneID2GO)[sapply(geneID2GO,function(x) all(GOtoMatch %in% x))])

number_of_sign_genes = nrow(getAdjustedPValues(stageRAdj, onlySignificantGenes = TRUE, order = FALSE)) #11,759
Padjscreen_sorted = DEG2sig_results[with(DEG2sig_results, 
                                    order(DEG2sig_results$padjScreen)),] #rank genes based on padjScreen
genesOfInterest = rownames(subset(Padjscreen_sorted, rownames(Padjscreen_sorted)%in%geneUniverse))
length(genesOfInterest)

#create gene list for input in topGO
geneList = factor(as.integer(geneUniverse %in% genesOfInterest))
names(geneList) = geneUniverse
str(geneList)

##create a topGO object (for biological process GOs)
GOdata_BP = new("topGOdata", ontology="BP", allGenes=geneList, 
                              annot = annFUN.gene2GO, gene2GO = geneID2GO)

##create a topGO object (for molecular function GOs)
GOdata_MF = new("topGOdata", ontology="MF", allGenes=geneList, 
                              annot = annFUN.gene2GO, gene2GO = geneID2GO)

#run Fisher's exact test
resultFisher_BP <- runTest(GOdata_BP, algorithm = "elim", statistic = "fisher")
resultFisher_MF <- runTest(GOdata_MF, algorithm = "elim", statistic = "fisher")

#extract the significant GO terms, trim the topNodes down to whatever you want (i.e., classic < 0.05)
allRes_BP <- GenTable(GOdata_BP, classic = resultFisher_BP, 
                                    orderBy = "weight", ranksOf = "weight", topNodes = 48)
allRes_MF <- GenTable(GOdata_MF, classic = resultFisher_MF, 
                                    orderBy = "weight", ranksOf = "weight", topNodes = 43)
```

## 4.2 Significantly differentially expressed genes

Gene enrichment analyses of the significantly DE genes without accounting for up/down regulation.

```{r echo = TRUE, message=FALSE, warning=FALSE, results='hide'}
#select the set of significant DE genes with GO terms
genesOfInterest_allSign  = rownames(subset(OnlySignGenes_ConStage, rownames(OnlySignGenes_ConStage)%in%geneUniverse))
length(genesOfInterest_allSign) 

#create gene list for input in topGO
geneList_allDE = factor(as.integer(geneUniverse %in% genesOfInterest_allSign))
names(geneList_allDE) = geneUniverse
str(geneList_allDE)

##create a topGO object (for biological process GOs)
GOdata_BP_allDE = new("topGOdata", ontology="BP", allGenes=geneList_allDE, 
                           annot = annFUN.gene2GO, gene2GO = geneID2GO)
##create a topGO object (for molecular function GOs)
GOdata_MF_allDE = new("topGOdata", ontology="MF", allGenes=geneList_allDE, 
                           annot = annFUN.gene2GO, gene2GO = geneID2GO)

#run Fisher's exact test
resultFisher_BP_DEG_elim <- runTest(GOdata_BP_allDE, algorithm = "elim", statistic = "fisher")
resultFisher_MF_DEG_elim <- runTest(GOdata_MF_allDE, algorithm = "elim", statistic = "fisher")

#extract the significant GO terms, trim the topNodes down to whatever you want (i.e., classic < 0.05)
DEG_BP_elim <- GenTable(GOdata_BP_allDE, classic = resultFisher_BP_DEG_elim, 
                                        orderBy = "weight", ranksOf = "weight", topNodes = 50)
DEG_MF_elim <- GenTable(GOdata_MF_allDE, classic = resultFisher_MF_DEG_elim, 
                                        orderBy = "weight", ranksOf = "weight", topNodes = 50)


#Generate the files for REVIGO input, which only needs a GO.ID and the p-value 

#import results from REVIGO back in if you want visualize them or use them in future analyses. I don't use this set, so I won't import them again.

```

## 4.3 Long-term data

Gene enrichment analyses for the long-term Nakov et al. 2020 subset, separated by regulation direction

```{r echo = TRUE, message=FALSE, warning=FALSE, results='hide'}

#C.cryptica longterm data, by FC direction (up or downregulated)

nakov_dr_GOEnrich = rownames(subset(nakov_DE_dr, rownames(nakov_DE_dr)%in%geneUniverse))
nakov_ur_GOEnrich = rownames(subset(nakov_DE_ur, rownames(nakov_DE_ur)%in%geneUniverse))

geneList_nakov_dr = factor(as.integer(geneUniverse %in% nakov_dr_GOEnrich))
names(geneList_nakov_dr) = geneUniverse
str(geneList_nakov_dr)
geneList_nakov_ur = factor(as.integer(geneUniverse %in% nakov_ur_GOEnrich))
names(geneList_nakov_ur) = geneUniverse
str(geneList_nakov_ur)

##create a topGO object (for up GOs)
GOdata_BP_nakov_dr = new("topGOdata", ontology="BP", allGenes=geneList_nakov_dr, 
                        annot = annFUN.gene2GO, gene2GO = geneID2GO)
GOdata_MF_nakov_dr = new("topGOdata", ontology="MF", allGenes=geneList_nakov_dr, 
                        annot = annFUN.gene2GO, gene2GO = geneID2GO)
GOdata_CC_nakov_dr = new("topGOdata", ontology="CC", allGenes=geneList_nakov_dr, 
                        annot = annFUN.gene2GO, gene2GO = geneID2GO)
##create a topGO object (for down GOs)
GOdata_BP_nakov_ur  = new("topGOdata", ontology="BP", allGenes=geneList_nakov_ur , 
                           annot = annFUN.gene2GO, gene2GO = geneID2GO)
GOdata_MF_nakov_ur  = new("topGOdata", ontology="MF", allGenes=geneList_nakov_ur , 
                           annot = annFUN.gene2GO, gene2GO = geneID2GO)
GOdata_CC_nakov_ur  = new("topGOdata", ontology="CC", allGenes=geneList_nakov_ur , 
                           annot = annFUN.gene2GO, gene2GO = geneID2GO)

#run Fisher's exact test
resultFisher_BP_nakov_dr_elim <- runTest(GOdata_BP_nakov_dr, algorithm = "elim", statistic = "fisher")
resultFisher_MF_nakov_dr_elim <- runTest(GOdata_MF_nakov_dr, algorithm = "elim", statistic = "fisher")
resultFisher_CC_nakov_dr_elim <- runTest(GOdata_CC_nakov_dr, algorithm = "elim", statistic = "fisher")
resultFisher_BP_nakov_ur_elim <- runTest(GOdata_BP_nakov_ur, algorithm = "elim", statistic = "fisher")
resultFisher_MF_nakov_ur_elim <- runTest(GOdata_MF_nakov_ur, algorithm = "elim", statistic = "fisher")
resultFisher_CC_nakov_ur_elim <- runTest(GOdata_CC_nakov_ur, algorithm = "elim", statistic = "fisher")

#extract the significant GO terms
nakov_dr_BP_elim <- GenTable(GOdata_BP_nakov_dr, classic = resultFisher_BP_nakov_dr_elim, 
                            orderBy = "weight", ranksOf = "weight", topNodes = 50)
nakov_dr_MF_elim <- GenTable(GOdata_MF_nakov_dr, classic = resultFisher_MF_nakov_dr_elim, 
                            orderBy = "weight", ranksOf = "weight", topNodes = 50)
nakov_dr_CC_elim <- GenTable(GOdata_CC_nakov_dr, classic = resultFisher_CC_nakov_dr_elim, 
                            orderBy = "weight", ranksOf = "weight", topNodes = 50)
nakov_ur_BP_elim <- GenTable(GOdata_BP_nakov_ur, classic = resultFisher_BP_nakov_ur_elim, 
                              orderBy = "weight", ranksOf = "weight", topNodes = 50)
nakov_ur_MF_elim <- GenTable(GOdata_MF_nakov_ur, classic = resultFisher_MF_nakov_ur_elim, 
                              orderBy = "weight", ranksOf = "weight", topNodes = 50)
nakov_ur_CC_elim <- GenTable(GOdata_CC_nakov_ur, classic = resultFisher_CC_nakov_ur_elim, 
                              orderBy = "weight", ranksOf = "weight", topNodes = 50)

#Generate the files for REVIGO input, which only needs a GO.ID and the p-value 
#Write the tables to you desired location to run through revigo using code below as a template.
#write.table(nakov_ur_BP_elim,
#            "/My Drive/ShortTermStress-IlluminaData/FullExp/Cryp-DE/data/ReviGo_files/BP_upLT_elim.txt")
#write.table(nakov_ur_MF_elim,
#            "/My Drive/ShortTermStress-IlluminaData/FullExp/Cryp-DE/data/ReviGo_files/MF_upLT_elim.txt")
#write.table(nakov_ur_CC_elim,
#            "/My Drive/ShortTermStress-IlluminaData/FullExp/Cryp-DE/data/ReviGo_files/CC_upLT_elim.txt")
#write.table(nakov_dr_BP_elim,
#            "/My Drive/ShortTermStress-IlluminaData/FullExp/Cryp-DE/data/ReviGo_files/BP_dnLT_elim.txt")
#write.table(nakov_dr_MF_elim,
#            "/My Drive/ShortTermStress-IlluminaData/FullExp/Cryp-DE/data/ReviGo_files/MF_dnLT_elim.txt")
#write.table(nakov_dr_CC_elim,
#            "/My Drive/ShortTermStress-IlluminaData/FullExp/Cryp-DE/data/ReviGo_files/CC_dnLT_elim.txt")

#import results from REVIGO back in if you want visualize them or use them in future analyses. The results were so small, no reduction was necessary

```

## 4.4 Comparisons between peak stress, acclimation, and post-acclimation 

Gene enrichment analyses for only the significant genes at 60 min and 10 hour timepoints, which we identified as the "peak" stress response and the beginning of acclimation, separated by direction of regulation. We then compile tables and figures to compare against long-term data

15 minute timepoint

```{r}
#Select only the genes significant at a timepoint from the reduced subset of 2 consecutive gene. This can be done for all time points, but is limited here to the ones discussed extensively in the manuscript
sign_0v15m <- rownames(keep_2_SignDEG)[keep_2_SignDEG[,"C0m-C60m"]==1] 

length(sign_0v15m) #3497 and this matches the output from SignifGenes
sign2LFC_0v15m <- sign2LFC [rownames(sign2LFC) %in% sign_0v15m, ] #3019 rows, which means one row had NAs and got omitted. 
logFCup_0v15m <- subset(sign2LFC_0v15m, (`0v15m_Padj`) <= 0.01 & (`0v15m`) > 0)
logFCdown_0v15m <- subset(sign2LFC_0v15m, (`0v15m_Padj`) <= 0.01 & (`0v15m`) < 0)

logFCup_0v15m_GOEnrich  = rownames(subset(logFCup_0v15m, rownames(logFCup_0v15m)%in%genesOfInterest_allSign))
logFCdown_0v15m_GOEnrich  = rownames(subset(logFCdown_0v15m, rownames(logFCdown_0v15m)%in%genesOfInterest_allSign))


#create gene list for input in topGO for 30 minutes
geneList_up0v15m = factor(as.integer(geneUniverse %in% logFCup_0v15m_GOEnrich))
names(geneList_up0v15m) = geneUniverse
str(geneList_up0v15m)
geneList_down0v15m = factor(as.integer(geneUniverse %in% logFCdown_0v15m_GOEnrich))
names(geneList_down0v15m) = geneUniverse
str(geneList_down0v15m)

##create a topGO object (for up GOs)
GOdata_BP_up0v15m = new("topGOdata", ontology="BP", allGenes=geneList_up0v15m, 
                        annot = annFUN.gene2GO, gene2GO = geneID2GO)
GOdata_MF_up0v15m = new("topGOdata", ontology="MF", allGenes=geneList_up0v15m, 
                        annot = annFUN.gene2GO, gene2GO = geneID2GO)
GOdata_CC_up0v15m = new("topGOdata", ontology="CC", allGenes=geneList_up0v15m, 
                        annot = annFUN.gene2GO, gene2GO = geneID2GO)
##create a topGO object (for down GOs)
GOdata_BP_down0v15m  = new("topGOdata", ontology="BP", allGenes=geneList_down0v15m , 
                           annot = annFUN.gene2GO, gene2GO = geneID2GO)
GOdata_MF_down0v15m  = new("topGOdata", ontology="MF", allGenes=geneList_down0v15m , 
                           annot = annFUN.gene2GO, gene2GO = geneID2GO)
GOdata_CC_down0v15m  = new("topGOdata", ontology="CC", allGenes=geneList_down0v15m , 
                           annot = annFUN.gene2GO, gene2GO = geneID2GO)

#run Fisher's exact test
resultFisher_BP_up0v15m_elim <- runTest(GOdata_BP_up0v15m, algorithm = "elim", statistic = "fisher")
resultFisher_MF_up0v15m_elim <- runTest(GOdata_MF_up0v15m, algorithm = "elim", statistic = "fisher")
resultFisher_CC_up0v15m_elim <- runTest(GOdata_CC_up0v15m, algorithm = "elim", statistic = "fisher")
resultFisher_BP_down0v15m_elim <- runTest(GOdata_BP_down0v15m, algorithm = "elim", statistic = "fisher")
resultFisher_MF_down0v15m_elim <- runTest(GOdata_MF_down0v15m, algorithm = "elim", statistic = "fisher")
resultFisher_CC_down0v15m_elim <- runTest(GOdata_CC_down0v15m, algorithm = "elim", statistic = "fisher")

#extract the significant GO terms
up0v15m_BP_elim <- GenTable(GOdata_BP_up0v15m, classic = resultFisher_BP_up0v15m_elim, 
                            orderBy = "weight", ranksOf = "weight", topNodes = 50)
up0v15m_MF_elim <- GenTable(GOdata_MF_up0v15m, classic = resultFisher_MF_up0v15m_elim, 
                            orderBy = "weight", ranksOf = "weight", topNodes = 50)
up0v15m_CC_elim <- GenTable(GOdata_CC_up0v15m, classic = resultFisher_CC_up0v15m_elim, 
                            orderBy = "weight", ranksOf = "weight", topNodes = 50)
down0v15m_BP_elim <- GenTable(GOdata_BP_down0v15m, classic = resultFisher_BP_down0v15m_elim, 
                              orderBy = "weight", ranksOf = "weight", topNodes = 50)
down0v15m_MF_elim <- GenTable(GOdata_MF_down0v15m, classic = resultFisher_MF_down0v15m_elim, 
                              orderBy = "weight", ranksOf = "weight", topNodes = 50)
down0v15m_CC_elim <- GenTable(GOdata_CC_down0v15m, classic = resultFisher_CC_down0v15m_elim, 
                              orderBy = "weight", ranksOf = "weight", topNodes = 50)

#Write the tables to you desired location to run through revigo using code below as a template.
#write.table(up0v15m_BP_elim,
#            "/My Drive/ShortTermStress-IlluminaData/FullExp/Cryp-DE/data/ReviGo_files/BP_up15m_elim.txt")
#write.table(up0v15m_MF_elim,
#            "/My Drive/ShortTermStress-IlluminaData/FullExp/Cryp-DE/data/ReviGo_files/MF_up15m_elim.txt")
#write.table(up0v15m_CC_elim,
#            "/My Drive/ShortTermStress-IlluminaData/FullExp/Cryp-DE/data/ReviGo_files/CC_up15m_elim.txt")
#write.table(down0v15m_BP_elim,
#            "/My Drive/ShortTermStress-IlluminaData/FullExp/Cryp-DE/data/ReviGo_files/BP_dn15m_elim.txt")
#write.table(down0v15m_MF_elim,
#            "/My Drive/ShortTermStress-IlluminaData/FullExp/Cryp-DE/data/ReviGo_files/MF_dn15m_elim.txt")
#write.table(down0v15m_CC_elim,
#            "/My Drive/ShortTermStress-IlluminaData/FullExp/Cryp-DE/data/ReviGo_files/CC_dn15m_elim.txt")
```

30 minute timepoint

```{r}
#Select only the genes significant at a timepoint from the reduced subset of 2 consecutive gene. This can be done for all time points, but is limited here to the ones discussed extensively in the manuscript
sign_0v30m <- rownames(keep_2_SignDEG)[keep_2_SignDEG[,"C0m-C60m"]==1] 

length(sign_0v30m) #3497 and this matches the output from SignifGenes
sign2LFC_0v30m <- sign2LFC [rownames(sign2LFC) %in% sign_0v30m, ] #3019 rows, which means one row had NAs and got omitted. 
logFCup_0v30m <- subset(sign2LFC_0v30m, (`0v30m_Padj`) <= 0.01 & (`0v30m`) > 0)
logFCdown_0v30m <- subset(sign2LFC_0v30m, (`0v30m_Padj`) <= 0.01 & (`0v30m`) < 0)

logFCup_0v30m_GOEnrich  = rownames(subset(logFCup_0v30m, rownames(logFCup_0v30m)%in%genesOfInterest_allSign))
logFCdown_0v30m_GOEnrich  = rownames(subset(logFCdown_0v30m, rownames(logFCdown_0v30m)%in%genesOfInterest_allSign))


#create gene list for input in topGO for 30 minutes
geneList_up0v30m = factor(as.integer(geneUniverse %in% logFCup_0v30m_GOEnrich))
names(geneList_up0v30m) = geneUniverse
str(geneList_up0v30m)
geneList_down0v30m = factor(as.integer(geneUniverse %in% logFCdown_0v30m_GOEnrich))
names(geneList_down0v30m) = geneUniverse
str(geneList_down0v30m)

##create a topGO object (for up GOs)
GOdata_BP_up0v30m = new("topGOdata", ontology="BP", allGenes=geneList_up0v30m, 
                        annot = annFUN.gene2GO, gene2GO = geneID2GO)
GOdata_MF_up0v30m = new("topGOdata", ontology="MF", allGenes=geneList_up0v30m, 
                        annot = annFUN.gene2GO, gene2GO = geneID2GO)
GOdata_CC_up0v30m = new("topGOdata", ontology="CC", allGenes=geneList_up0v30m, 
                        annot = annFUN.gene2GO, gene2GO = geneID2GO)
##create a topGO object (for down GOs)
GOdata_BP_down0v30m  = new("topGOdata", ontology="BP", allGenes=geneList_down0v30m , 
                           annot = annFUN.gene2GO, gene2GO = geneID2GO)
GOdata_MF_down0v30m  = new("topGOdata", ontology="MF", allGenes=geneList_down0v30m , 
                           annot = annFUN.gene2GO, gene2GO = geneID2GO)
GOdata_CC_down0v30m  = new("topGOdata", ontology="CC", allGenes=geneList_down0v30m , 
                           annot = annFUN.gene2GO, gene2GO = geneID2GO)

#run Fisher's exact test
resultFisher_BP_up0v30m_elim <- runTest(GOdata_BP_up0v30m, algorithm = "elim", statistic = "fisher")
resultFisher_MF_up0v30m_elim <- runTest(GOdata_MF_up0v30m, algorithm = "elim", statistic = "fisher")
resultFisher_CC_up0v30m_elim <- runTest(GOdata_CC_up0v30m, algorithm = "elim", statistic = "fisher")
resultFisher_BP_down0v30m_elim <- runTest(GOdata_BP_down0v30m, algorithm = "elim", statistic = "fisher")
resultFisher_MF_down0v30m_elim <- runTest(GOdata_MF_down0v30m, algorithm = "elim", statistic = "fisher")
resultFisher_CC_down0v30m_elim <- runTest(GOdata_CC_down0v30m, algorithm = "elim", statistic = "fisher")

#extract the significant GO terms
up0v30m_BP_elim <- GenTable(GOdata_BP_up0v30m, classic = resultFisher_BP_up0v30m_elim, 
                            orderBy = "weight", ranksOf = "weight", topNodes = 50)
up0v30m_MF_elim <- GenTable(GOdata_MF_up0v30m, classic = resultFisher_MF_up0v30m_elim, 
                            orderBy = "weight", ranksOf = "weight", topNodes = 50)
up0v30m_CC_elim <- GenTable(GOdata_CC_up0v30m, classic = resultFisher_CC_up0v30m_elim, 
                            orderBy = "weight", ranksOf = "weight", topNodes = 50)
down0v30m_BP_elim <- GenTable(GOdata_BP_down0v30m, classic = resultFisher_BP_down0v30m_elim, 
                              orderBy = "weight", ranksOf = "weight", topNodes = 50)
down0v30m_MF_elim <- GenTable(GOdata_MF_down0v30m, classic = resultFisher_MF_down0v30m_elim, 
                              orderBy = "weight", ranksOf = "weight", topNodes = 50)
down0v30m_CC_elim <- GenTable(GOdata_CC_down0v30m, classic = resultFisher_CC_down0v30m_elim, 
                              orderBy = "weight", ranksOf = "weight", topNodes = 50)

#Write the tables to you desired location to run through revigo using code below as a template.
#write.table(up0v30m_BP_elim,
#            "/My Drive/ShortTermStress-IlluminaData/FullExp/Cryp-DE/data/ReviGo_files/BP_up30m_elim.txt")
#write.table(up0v30m_MF_elim,
#            "/My Drive/ShortTermStress-IlluminaData/FullExp/Cryp-DE/data/ReviGo_files/MF_up30m_elim.txt")
#write.table(up0v30m_CC_elim,
#            "/My Drive/ShortTermStress-IlluminaData/FullExp/Cryp-DE/data/ReviGo_files/CC_up30m_elim.txt")
#write.table(down0v30m_BP_elim,
#            "/My Drive/ShortTermStress-IlluminaData/FullExp/Cryp-DE/data/ReviGo_files/BP_dn30m_elim.txt")
#write.table(down0v30m_MF_elim,
#            "/My Drive/ShortTermStress-IlluminaData/FullExp/Cryp-DE/data/ReviGo_files/MF_dn30m_elim.txt")
#write.table(down0v30m_CC_elim,
#            "/My Drive/ShortTermStress-IlluminaData/FullExp/Cryp-DE/data/ReviGo_files/CC_dn30m_elim.txt")
```

1 hour timepoint; peak period of low salinity stress

```{r}
#Select only the genes significant at a timepoint from the reduced subset of 2 consecutive gene. This can be done for all time points, but is limited here to the ones discussed extensively in the manuscript
sign_0v1h <- rownames(keep_2_SignDEG)[keep_2_SignDEG[,"C0m-C60m"]==1] 

length(sign_0v1h) #3497 and this matches the output from SignifGenes
sign2LFC_0v1h <- sign2LFC [rownames(sign2LFC) %in% sign_0v1h, ] #3019 rows, which means one row had NAs and got omitted. 
logFCup_0v1h <- subset(sign2LFC_0v1h, (`0v1h_Padj`) <= 0.01 & (`0v1h`) > 0)
logFCdown_0v1h <- subset(sign2LFC_0v1h, (`0v1h_Padj`) <= 0.01 & (`0v1h`) < 0)

logFCup_0v1h_GOEnrich  = rownames(subset(logFCup_0v1h, rownames(logFCup_0v1h)%in%genesOfInterest_allSign))
logFCdown_0v1h_GOEnrich  = rownames(subset(logFCdown_0v1h, rownames(logFCdown_0v1h)%in%genesOfInterest_allSign))


#create gene list for input in topGO for 30 minutes
geneList_up0v1h = factor(as.integer(geneUniverse %in% logFCup_0v1h_GOEnrich))
names(geneList_up0v1h) = geneUniverse
str(geneList_up0v1h)
geneList_down0v1h = factor(as.integer(geneUniverse %in% logFCdown_0v1h_GOEnrich))
names(geneList_down0v1h) = geneUniverse
str(geneList_down0v1h)

##create a topGO object (for up GOs)
GOdata_BP_up0v1h = new("topGOdata", ontology="BP", allGenes=geneList_up0v1h, 
                        annot = annFUN.gene2GO, gene2GO = geneID2GO)
GOdata_MF_up0v1h = new("topGOdata", ontology="MF", allGenes=geneList_up0v1h, 
                        annot = annFUN.gene2GO, gene2GO = geneID2GO)
GOdata_CC_up0v1h = new("topGOdata", ontology="CC", allGenes=geneList_up0v1h, 
                        annot = annFUN.gene2GO, gene2GO = geneID2GO)
##create a topGO object (for down GOs)
GOdata_BP_down0v1h  = new("topGOdata", ontology="BP", allGenes=geneList_down0v1h , 
                           annot = annFUN.gene2GO, gene2GO = geneID2GO)
GOdata_MF_down0v1h  = new("topGOdata", ontology="MF", allGenes=geneList_down0v1h , 
                           annot = annFUN.gene2GO, gene2GO = geneID2GO)
GOdata_CC_down0v1h  = new("topGOdata", ontology="CC", allGenes=geneList_down0v1h , 
                           annot = annFUN.gene2GO, gene2GO = geneID2GO)

#run Fisher's exact test
resultFisher_BP_up0v1h_elim <- runTest(GOdata_BP_up0v1h, algorithm = "elim", statistic = "fisher")
resultFisher_MF_up0v1h_elim <- runTest(GOdata_MF_up0v1h, algorithm = "elim", statistic = "fisher")
resultFisher_CC_up0v1h_elim <- runTest(GOdata_CC_up0v1h, algorithm = "elim", statistic = "fisher")
resultFisher_BP_down0v1h_elim <- runTest(GOdata_BP_down0v1h, algorithm = "elim", statistic = "fisher")
resultFisher_MF_down0v1h_elim <- runTest(GOdata_MF_down0v1h, algorithm = "elim", statistic = "fisher")
resultFisher_CC_down0v1h_elim <- runTest(GOdata_CC_down0v1h, algorithm = "elim", statistic = "fisher")

#extract the significant GO terms
up0v1h_BP_elim <- GenTable(GOdata_BP_up0v1h, classic = resultFisher_BP_up0v1h_elim, 
                            orderBy = "weight", ranksOf = "weight", topNodes = 50)
up0v1h_MF_elim <- GenTable(GOdata_MF_up0v1h, classic = resultFisher_MF_up0v1h_elim, 
                            orderBy = "weight", ranksOf = "weight", topNodes = 50)
up0v1h_CC_elim <- GenTable(GOdata_CC_up0v1h, classic = resultFisher_CC_up0v1h_elim, 
                            orderBy = "weight", ranksOf = "weight", topNodes = 50)
down0v1h_BP_elim <- GenTable(GOdata_BP_down0v1h, classic = resultFisher_BP_down0v1h_elim, 
                              orderBy = "weight", ranksOf = "weight", topNodes = 50)
down0v1h_MF_elim <- GenTable(GOdata_MF_down0v1h, classic = resultFisher_MF_down0v1h_elim, 
                              orderBy = "weight", ranksOf = "weight", topNodes = 50)
down0v1h_CC_elim <- GenTable(GOdata_CC_down0v1h, classic = resultFisher_CC_down0v1h_elim, 
                              orderBy = "weight", ranksOf = "weight", topNodes = 50)

#Write the tables to you desired location to run through revigo using code below as a template.
#write.table(up0v1h_BP_elim,
#            "/My Drive/ShortTermStress-IlluminaData/FullExp/Cryp-DE/data/ReviGo_files/BP_up1h_elim.txt")
#write.table(up0v1h_MF_elim,
#            "/My Drive/ShortTermStress-IlluminaData/FullExp/Cryp-DE/data/ReviGo_files/MF_up1h_elim.txt")
#write.table(up0v1h_CC_elim,
#            "/My Drive/ShortTermStress-IlluminaData/FullExp/Cryp-DE/data/ReviGo_files/CC_up1h_elim.txt")
#write.table(down0v1h_BP_elim,
#            "/My Drive/ShortTermStress-IlluminaData/FullExp/Cryp-DE/data/ReviGo_files/BP_dn1h_elim.txt")
#write.table(down0v1h_MF_elim,
#            "/My Drive/ShortTermStress-IlluminaData/FullExp/Cryp-DE/data/ReviGo_files/MF_dn1h_elim.txt")
#write.table(down0v1h_CC_elim,
#            "/My Drive/ShortTermStress-IlluminaData/FullExp/Cryp-DE/data/ReviGo_files/CC_dn1h_elim.txt")
```

2 hour timepoint

```{r}
#Select only the genes significant at a timepoint from the reduced subset of 2 consecutive gene. This can be done for all time points, but is limited here to the ones discussed extensively in the manuscript
sign_0v2h <- rownames(keep_2_SignDEG)[keep_2_SignDEG[,"C0m-C60m"]==1] 

length(sign_0v2h) #3497 and this matches the output from SignifGenes
sign2LFC_0v2h <- sign2LFC [rownames(sign2LFC) %in% sign_0v2h, ] #3019 rows, which means one row had NAs and got omitted. 
logFCup_0v2h <- subset(sign2LFC_0v2h, (`0v2h_Padj`) <= 0.01 & (`0v2h`) > 0)
logFCdown_0v2h <- subset(sign2LFC_0v2h, (`0v2h_Padj`) <= 0.01 & (`0v2h`) < 0)

logFCup_0v2h_GOEnrich  = rownames(subset(logFCup_0v2h, rownames(logFCup_0v2h)%in%genesOfInterest_allSign))
logFCdown_0v2h_GOEnrich  = rownames(subset(logFCdown_0v2h, rownames(logFCdown_0v2h)%in%genesOfInterest_allSign))


#create gene list for input in topGO for 30 minutes
geneList_up0v2h = factor(as.integer(geneUniverse %in% logFCup_0v2h_GOEnrich))
names(geneList_up0v2h) = geneUniverse
str(geneList_up0v2h)
geneList_down0v2h = factor(as.integer(geneUniverse %in% logFCdown_0v2h_GOEnrich))
names(geneList_down0v2h) = geneUniverse
str(geneList_down0v2h)

##create a topGO object (for up GOs)
GOdata_BP_up0v2h = new("topGOdata", ontology="BP", allGenes=geneList_up0v2h, 
                        annot = annFUN.gene2GO, gene2GO = geneID2GO)
GOdata_MF_up0v2h = new("topGOdata", ontology="MF", allGenes=geneList_up0v2h, 
                        annot = annFUN.gene2GO, gene2GO = geneID2GO)
GOdata_CC_up0v2h = new("topGOdata", ontology="CC", allGenes=geneList_up0v2h, 
                        annot = annFUN.gene2GO, gene2GO = geneID2GO)
##create a topGO object (for down GOs)
GOdata_BP_down0v2h  = new("topGOdata", ontology="BP", allGenes=geneList_down0v2h , 
                           annot = annFUN.gene2GO, gene2GO = geneID2GO)
GOdata_MF_down0v2h  = new("topGOdata", ontology="MF", allGenes=geneList_down0v2h , 
                           annot = annFUN.gene2GO, gene2GO = geneID2GO)
GOdata_CC_down0v2h  = new("topGOdata", ontology="CC", allGenes=geneList_down0v2h , 
                           annot = annFUN.gene2GO, gene2GO = geneID2GO)

#run Fisher's exact test
resultFisher_BP_up0v2h_elim <- runTest(GOdata_BP_up0v2h, algorithm = "elim", statistic = "fisher")
resultFisher_MF_up0v2h_elim <- runTest(GOdata_MF_up0v2h, algorithm = "elim", statistic = "fisher")
resultFisher_CC_up0v2h_elim <- runTest(GOdata_CC_up0v2h, algorithm = "elim", statistic = "fisher")
resultFisher_BP_down0v2h_elim <- runTest(GOdata_BP_down0v2h, algorithm = "elim", statistic = "fisher")
resultFisher_MF_down0v2h_elim <- runTest(GOdata_MF_down0v2h, algorithm = "elim", statistic = "fisher")
resultFisher_CC_down0v2h_elim <- runTest(GOdata_CC_down0v2h, algorithm = "elim", statistic = "fisher")

#extract the significant GO terms
up0v2h_BP_elim <- GenTable(GOdata_BP_up0v2h, classic = resultFisher_BP_up0v2h_elim, 
                            orderBy = "weight", ranksOf = "weight", topNodes = 50)
up0v2h_MF_elim <- GenTable(GOdata_MF_up0v2h, classic = resultFisher_MF_up0v2h_elim, 
                            orderBy = "weight", ranksOf = "weight", topNodes = 50)
up0v2h_CC_elim <- GenTable(GOdata_CC_up0v2h, classic = resultFisher_CC_up0v2h_elim, 
                            orderBy = "weight", ranksOf = "weight", topNodes = 50)
down0v2h_BP_elim <- GenTable(GOdata_BP_down0v2h, classic = resultFisher_BP_down0v2h_elim, 
                              orderBy = "weight", ranksOf = "weight", topNodes = 50)
down0v2h_MF_elim <- GenTable(GOdata_MF_down0v2h, classic = resultFisher_MF_down0v2h_elim, 
                              orderBy = "weight", ranksOf = "weight", topNodes = 50)
down0v2h_CC_elim <- GenTable(GOdata_CC_down0v2h, classic = resultFisher_CC_down0v2h_elim, 
                              orderBy = "weight", ranksOf = "weight", topNodes = 50)

#Write the tables to you desired location to run through revigo using code below as a template.
#write.table(up0v2h_BP_elim,
#            "/My Drive/ShortTermStress-IlluminaData/FullExp/Cryp-DE/data/ReviGo_files/BP_up2h_elim.txt")
#write.table(up0v2h_MF_elim,
#            "/My Drive/ShortTermStress-IlluminaData/FullExp/Cryp-DE/data/ReviGo_files/MF_up2h_elim.txt")
#write.table(up0v2h_CC_elim,
#            "/My Drive/ShortTermStress-IlluminaData/FullExp/Cryp-DE/data/ReviGo_files/CC_up2h_elim.txt")
#write.table(down0v2h_BP_elim,
#            "/My Drive/ShortTermStress-IlluminaData/FullExp/Cryp-DE/data/ReviGo_files/BP_dn2h_elim.txt")
#write.table(down0v2h_MF_elim,
#            "/My Drive/ShortTermStress-IlluminaData/FullExp/Cryp-DE/data/ReviGo_files/MF_dn2h_elim.txt")
#write.table(down0v2h_CC_elim,
#            "/My Drive/ShortTermStress-IlluminaData/FullExp/Cryp-DE/data/ReviGo_files/CC_dn2h_elim.txt")
```

4 hour timepoint

```{r}
#Select only the genes significant at a timepoint from the reduced subset of 2 consecutive gene. This can be done for all time points, but is limited here to the ones discussed extensively in the manuscript
sign_0v4h <- rownames(keep_2_SignDEG)[keep_2_SignDEG[,"C0m-C60m"]==1] 

length(sign_0v4h) #3497 and this matches the output from SignifGenes
sign2LFC_0v4h <- sign2LFC [rownames(sign2LFC) %in% sign_0v4h, ] #3019 rows, which means one row had NAs and got omitted. 
logFCup_0v4h <- subset(sign2LFC_0v4h, (`0v4h_Padj`) <= 0.01 & (`0v4h`) > 0)
logFCdown_0v4h <- subset(sign2LFC_0v4h, (`0v4h_Padj`) <= 0.01 & (`0v4h`) < 0)

logFCup_0v4h_GOEnrich  = rownames(subset(logFCup_0v4h, rownames(logFCup_0v4h)%in%genesOfInterest_allSign))
logFCdown_0v4h_GOEnrich  = rownames(subset(logFCdown_0v4h, rownames(logFCdown_0v4h)%in%genesOfInterest_allSign))


#create gene list for input in topGO for 30 minutes
geneList_up0v4h = factor(as.integer(geneUniverse %in% logFCup_0v4h_GOEnrich))
names(geneList_up0v4h) = geneUniverse
str(geneList_up0v4h)
geneList_down0v4h = factor(as.integer(geneUniverse %in% logFCdown_0v4h_GOEnrich))
names(geneList_down0v4h) = geneUniverse
str(geneList_down0v4h)

##create a topGO object (for up GOs)
GOdata_BP_up0v4h = new("topGOdata", ontology="BP", allGenes=geneList_up0v4h, 
                        annot = annFUN.gene2GO, gene2GO = geneID2GO)
GOdata_MF_up0v4h = new("topGOdata", ontology="MF", allGenes=geneList_up0v4h, 
                        annot = annFUN.gene2GO, gene2GO = geneID2GO)
GOdata_CC_up0v4h = new("topGOdata", ontology="CC", allGenes=geneList_up0v4h, 
                        annot = annFUN.gene2GO, gene2GO = geneID2GO)
##create a topGO object (for down GOs)
GOdata_BP_down0v4h  = new("topGOdata", ontology="BP", allGenes=geneList_down0v4h , 
                           annot = annFUN.gene2GO, gene2GO = geneID2GO)
GOdata_MF_down0v4h  = new("topGOdata", ontology="MF", allGenes=geneList_down0v4h , 
                           annot = annFUN.gene2GO, gene2GO = geneID2GO)
GOdata_CC_down0v4h  = new("topGOdata", ontology="CC", allGenes=geneList_down0v4h , 
                           annot = annFUN.gene2GO, gene2GO = geneID2GO)

#run Fisher's exact test
resultFisher_BP_up0v4h_elim <- runTest(GOdata_BP_up0v4h, algorithm = "elim", statistic = "fisher")
resultFisher_MF_up0v4h_elim <- runTest(GOdata_MF_up0v4h, algorithm = "elim", statistic = "fisher")
resultFisher_CC_up0v4h_elim <- runTest(GOdata_CC_up0v4h, algorithm = "elim", statistic = "fisher")
resultFisher_BP_down0v4h_elim <- runTest(GOdata_BP_down0v4h, algorithm = "elim", statistic = "fisher")
resultFisher_MF_down0v4h_elim <- runTest(GOdata_MF_down0v4h, algorithm = "elim", statistic = "fisher")
resultFisher_CC_down0v4h_elim <- runTest(GOdata_CC_down0v4h, algorithm = "elim", statistic = "fisher")

#extract the significant GO terms
up0v4h_BP_elim <- GenTable(GOdata_BP_up0v4h, classic = resultFisher_BP_up0v4h_elim, 
                            orderBy = "weight", ranksOf = "weight", topNodes = 50)
up0v4h_MF_elim <- GenTable(GOdata_MF_up0v4h, classic = resultFisher_MF_up0v4h_elim, 
                            orderBy = "weight", ranksOf = "weight", topNodes = 50)
up0v4h_CC_elim <- GenTable(GOdata_CC_up0v4h, classic = resultFisher_CC_up0v4h_elim, 
                            orderBy = "weight", ranksOf = "weight", topNodes = 50)
down0v4h_BP_elim <- GenTable(GOdata_BP_down0v4h, classic = resultFisher_BP_down0v4h_elim, 
                              orderBy = "weight", ranksOf = "weight", topNodes = 50)
down0v4h_MF_elim <- GenTable(GOdata_MF_down0v4h, classic = resultFisher_MF_down0v4h_elim, 
                              orderBy = "weight", ranksOf = "weight", topNodes = 50)
down0v4h_CC_elim <- GenTable(GOdata_CC_down0v4h, classic = resultFisher_CC_down0v4h_elim, 
                              orderBy = "weight", ranksOf = "weight", topNodes = 50)

#Write the tables to you desired location to run through revigo using code below as a template.
#write.table(up0v4h_BP_elim,
#            "/My Drive/ShortTermStress-IlluminaData/FullExp/Cryp-DE/data/ReviGo_files/BP_up4h_elim.txt")
#write.table(up0v4h_MF_elim,
#            "/My Drive/ShortTermStress-IlluminaData/FullExp/Cryp-DE/data/ReviGo_files/MF_up4h_elim.txt")
#write.table(up0v4h_CC_elim,
#            "/My Drive/ShortTermStress-IlluminaData/FullExp/Cryp-DE/data/ReviGo_files/CC_up4h_elim.txt")
#write.table(down0v4h_BP_elim,
#            "/My Drive/ShortTermStress-IlluminaData/FullExp/Cryp-DE/data/ReviGo_files/BP_dn4h_elim.txt")
#write.table(down0v4h_MF_elim,
#            "/My Drive/ShortTermStress-IlluminaData/FullExp/Cryp-DE/data/ReviGo_files/MF_dn4h_elim.txt")
#write.table(down0v4h_CC_elim,
#            "/My Drive/ShortTermStress-IlluminaData/FullExp/Cryp-DE/data/ReviGo_files/CC_dn4h_elim.txt")
```

8 hour timepoint

```{r}
#Select only the genes significant at a timepoint from the reduced subset of 2 consecutive gene. This can be done for all time points, but is limited here to the ones discussed extensively in the manuscript
sign_0v8h <- rownames(keep_2_SignDEG)[keep_2_SignDEG[,"C0m-C60m"]==1] 

length(sign_0v8h) #3497 and this matches the output from SignifGenes
sign2LFC_0v8h <- sign2LFC [rownames(sign2LFC) %in% sign_0v8h, ] #3019 rows, which means one row had NAs and got omitted. 
logFCup_0v8h <- subset(sign2LFC_0v8h, (`0v8h_Padj`) <= 0.01 & (`0v8h`) > 0)
logFCdown_0v8h <- subset(sign2LFC_0v8h, (`0v8h_Padj`) <= 0.01 & (`0v8h`) < 0)

logFCup_0v8h_GOEnrich  = rownames(subset(logFCup_0v8h, rownames(logFCup_0v8h)%in%genesOfInterest_allSign))
logFCdown_0v8h_GOEnrich  = rownames(subset(logFCdown_0v8h, rownames(logFCdown_0v8h)%in%genesOfInterest_allSign))


#create gene list for input in topGO for 30 minutes
geneList_up0v8h = factor(as.integer(geneUniverse %in% logFCup_0v8h_GOEnrich))
names(geneList_up0v8h) = geneUniverse
str(geneList_up0v8h)
geneList_down0v8h = factor(as.integer(geneUniverse %in% logFCdown_0v8h_GOEnrich))
names(geneList_down0v8h) = geneUniverse
str(geneList_down0v8h)

##create a topGO object (for up GOs)
GOdata_BP_up0v8h = new("topGOdata", ontology="BP", allGenes=geneList_up0v8h, 
                        annot = annFUN.gene2GO, gene2GO = geneID2GO)
GOdata_MF_up0v8h = new("topGOdata", ontology="MF", allGenes=geneList_up0v8h, 
                        annot = annFUN.gene2GO, gene2GO = geneID2GO)
GOdata_CC_up0v8h = new("topGOdata", ontology="CC", allGenes=geneList_up0v8h, 
                        annot = annFUN.gene2GO, gene2GO = geneID2GO)
##create a topGO object (for down GOs)
GOdata_BP_down0v8h  = new("topGOdata", ontology="BP", allGenes=geneList_down0v8h , 
                           annot = annFUN.gene2GO, gene2GO = geneID2GO)
GOdata_MF_down0v8h  = new("topGOdata", ontology="MF", allGenes=geneList_down0v8h , 
                           annot = annFUN.gene2GO, gene2GO = geneID2GO)
GOdata_CC_down0v8h  = new("topGOdata", ontology="CC", allGenes=geneList_down0v8h , 
                           annot = annFUN.gene2GO, gene2GO = geneID2GO)

#run Fisher's exact test
resultFisher_BP_up0v8h_elim <- runTest(GOdata_BP_up0v8h, algorithm = "elim", statistic = "fisher")
resultFisher_MF_up0v8h_elim <- runTest(GOdata_MF_up0v8h, algorithm = "elim", statistic = "fisher")
resultFisher_CC_up0v8h_elim <- runTest(GOdata_CC_up0v8h, algorithm = "elim", statistic = "fisher")
resultFisher_BP_down0v8h_elim <- runTest(GOdata_BP_down0v8h, algorithm = "elim", statistic = "fisher")
resultFisher_MF_down0v8h_elim <- runTest(GOdata_MF_down0v8h, algorithm = "elim", statistic = "fisher")
resultFisher_CC_down0v8h_elim <- runTest(GOdata_CC_down0v8h, algorithm = "elim", statistic = "fisher")

#extract the significant GO terms
up0v8h_BP_elim <- GenTable(GOdata_BP_up0v8h, classic = resultFisher_BP_up0v8h_elim, 
                            orderBy = "weight", ranksOf = "weight", topNodes = 50)
up0v8h_MF_elim <- GenTable(GOdata_MF_up0v8h, classic = resultFisher_MF_up0v8h_elim, 
                            orderBy = "weight", ranksOf = "weight", topNodes = 50)
up0v8h_CC_elim <- GenTable(GOdata_CC_up0v8h, classic = resultFisher_CC_up0v8h_elim, 
                            orderBy = "weight", ranksOf = "weight", topNodes = 50)
down0v8h_BP_elim <- GenTable(GOdata_BP_down0v8h, classic = resultFisher_BP_down0v8h_elim, 
                              orderBy = "weight", ranksOf = "weight", topNodes = 50)
down0v8h_MF_elim <- GenTable(GOdata_MF_down0v8h, classic = resultFisher_MF_down0v8h_elim, 
                              orderBy = "weight", ranksOf = "weight", topNodes = 50)
down0v8h_CC_elim <- GenTable(GOdata_CC_down0v8h, classic = resultFisher_CC_down0v8h_elim, 
                              orderBy = "weight", ranksOf = "weight", topNodes = 50)

#Write the tables to you desired location to run through revigo using code below as a template.
#write.table(up0v8h_BP_elim,
#            "/My Drive/ShortTermStress-IlluminaData/FullExp/Cryp-DE/data/ReviGo_files/BP_up8h_elim.txt")
#write.table(up0v8h_MF_elim,
#            "/My Drive/ShortTermStress-IlluminaData/FullExp/Cryp-DE/data/ReviGo_files/MF_up8h_elim.txt")
#write.table(up0v8h_CC_elim,
#            "/My Drive/ShortTermStress-IlluminaData/FullExp/Cryp-DE/data/ReviGo_files/CC_up8h_elim.txt")
#write.table(down0v8h_BP_elim,
#            "/My Drive/ShortTermStress-IlluminaData/FullExp/Cryp-DE/data/ReviGo_files/BP_dn8h_elim.txt")
#write.table(down0v8h_MF_elim,
#            "/My Drive/ShortTermStress-IlluminaData/FullExp/Cryp-DE/data/ReviGo_files/MF_dn8h_elim.txt")
#write.table(down0v8h_CC_elim,
#            "/My Drive/ShortTermStress-IlluminaData/FullExp/Cryp-DE/data/ReviGo_files/CC_dn8h_elim.txt")
```

10 hour timepoint; early acclimation

```{r}
#Time 0 vs 10 hour
sign_0v10h <- rownames(keep_2_SignDEG)[keep_2_SignDEG[,"C0m-C10hr"]==1] 

length(sign_0v10h) 
sign2LFC_0v10h <- sign2LFC [rownames(sign2LFC) %in% sign_0v10h, ]  
logFCup_0v10h <- subset(sign2LFC_0v10h, (`0v10h_Padj`) <= 0.01 & (`0v10h`) > 0)
logFCdown_0v10h <- subset(sign2LFC_0v10h, (`0v10h_Padj`) <= 0.01 & (`0v10h`) < 0)

logFCup_0v10h_GOEnrich  = rownames(subset(logFCup_0v10h, rownames(logFCup_0v10h)%in%genesOfInterest_allSign))
logFCdown_0v10h_GOEnrich  = rownames(subset(logFCdown_0v10h, rownames(logFCdown_0v10h)%in%genesOfInterest_allSign))


#create gene list for input in topGO for 10 hours
geneList_up0v10h = factor(as.integer(geneUniverse %in% logFCup_0v10h_GOEnrich))
names(geneList_up0v10h) = geneUniverse
str(geneList_up0v10h)
geneList_down0v10h = factor(as.integer(geneUniverse %in% logFCdown_0v10h_GOEnrich))
names(geneList_down0v10h) = geneUniverse
str(geneList_down0v10h)

##create a topGO object (for up GOs)
GOdata_BP_up0v10h = new("topGOdata", ontology="BP", allGenes=geneList_up0v10h, 
                        annot = annFUN.gene2GO, gene2GO = geneID2GO)
GOdata_MF_up0v10h = new("topGOdata", ontology="MF", allGenes=geneList_up0v10h, 
                        annot = annFUN.gene2GO, gene2GO = geneID2GO)
GOdata_CC_up0v10h = new("topGOdata", ontology="CC", allGenes=geneList_up0v10h, 
                        annot = annFUN.gene2GO, gene2GO = geneID2GO)
##create a topGO object (for down GOs)
GOdata_BP_down0v10h  = new("topGOdata", ontology="BP", allGenes=geneList_down0v10h , 
                           annot = annFUN.gene2GO, gene2GO = geneID2GO)
GOdata_MF_down0v10h  = new("topGOdata", ontology="MF", allGenes=geneList_down0v10h , 
                           annot = annFUN.gene2GO, gene2GO = geneID2GO)
GOdata_CC_down0v10h  = new("topGOdata", ontology="CC", allGenes=geneList_down0v10h , 
                           annot = annFUN.gene2GO, gene2GO = geneID2GO)

#run Fisher's exact test
resultFisher_BP_up0v10h_elim <- runTest(GOdata_BP_up0v10h, algorithm = "elim", statistic = "fisher")
resultFisher_MF_up0v10h_elim <- runTest(GOdata_MF_up0v10h, algorithm = "elim", statistic = "fisher")
resultFisher_CC_up0v10h_elim <- runTest(GOdata_CC_up0v10h, algorithm = "elim", statistic = "fisher")
resultFisher_BP_down0v10h_elim <- runTest(GOdata_BP_down0v10h, algorithm = "elim", statistic = "fisher")
resultFisher_MF_down0v10h_elim <- runTest(GOdata_MF_down0v10h, algorithm = "elim", statistic = "fisher")
resultFisher_CC_down0v10h_elim <- runTest(GOdata_CC_down0v10h, algorithm = "elim", statistic = "fisher")

#extract the significant GO terms
up0v10h_BP_elim <- GenTable(GOdata_BP_up0v10h, classic = resultFisher_BP_up0v10h_elim, 
                            orderBy = "weight", ranksOf = "weight", topNodes = 50)
up0v10h_MF_elim <- GenTable(GOdata_MF_up0v10h, classic = resultFisher_MF_up0v10h_elim, 
                            orderBy = "weight", ranksOf = "weight", topNodes = 50)
up0v10h_CC_elim <- GenTable(GOdata_CC_up0v10h, classic = resultFisher_CC_up0v10h_elim, 
                            orderBy = "weight", ranksOf = "weight", topNodes = 50)
down0v10h_BP_elim <- GenTable(GOdata_BP_down0v10h, classic = resultFisher_BP_down0v10h_elim, 
                              orderBy = "weight", ranksOf = "weight", topNodes = 50)
down0v10h_MF_elim <- GenTable(GOdata_MF_down0v10h, classic = resultFisher_MF_down0v10h_elim, 
                              orderBy = "weight", ranksOf = "weight", topNodes = 50)
down0v10h_CC_elim <- GenTable(GOdata_CC_down0v10h, classic = resultFisher_CC_down0v10h_elim, 
                              orderBy = "weight", ranksOf = "weight", topNodes = 50)

#Write the tables to you desired location to run through revigo using code below as a template.
#write.table(up0v10h_BP_elim,
#            "/My Drive/ShortTermStress-IlluminaData/FullExp/Cryp-DE/data/ReviGo_files/BP_up10h_elim.txt")
#write.table(up0v10h_MF_elim,
#            "/My Drive/ShortTermStress-IlluminaData/FullExp/Cryp-DE/data/ReviGo_files/MF_up10h_elim.txt")
#write.table(up0v10h_CC_elim,
#            "/My Drive/ShortTermStress-IlluminaData/FullExp/Cryp-DE/data/ReviGo_files/CC_up10h_elim.txt")
#write.table(down0v10h_BP_elim,
#            "/My Drive/ShortTermStress-IlluminaData/FullExp/Cryp-DE/data/ReviGo_files/BP_dn10h_elim.txt")
#write.table(down0v10h_MF_elim,
#            "/My Drive/ShortTermStress-IlluminaData/FullExp/Cryp-DE/data/ReviGo_files/MF_dn10h_elim.txt")
#write.table(down0v10h_CC_elim,
#            "/My Drive/ShortTermStress-IlluminaData/FullExp/Cryp-DE/data/ReviGo_files/CC_dn10h_elim.txt")
```

Import the necessary Revigo output columns back in. These files are not available because you'll want to run ReviGo yourself to pick your similarity values. 

```{r}
#dr_1h_bp = read.table("/My Drive/ShortTermStress-IlluminaData/FullExp/Cryp-DE/data/ReviGo_files/BP_dn1h_Revigo.tsv",
#                       colClasses = c(NA,"NULL","NULL","NULL","NULL","NULL","NULL","NULL","NULL","NULL","NULL","NULL","NULL",NA),
#                       header=TRUE) #biological process results
#ur_1h_bp = read.table("/My Drive/ShortTermStress-IlluminaData/FullExp/Cryp-DE/data/ReviGo_files/BP_up1h_Revigo.tsv",
#                     colClasses = c(NA,"NULL","NULL","NULL","NULL","NULL","NULL","NULL","NULL","NULL","NULL","NULL","NULL",NA),
#                     header=TRUE) #biological process results
#dr_1h_cc = read.table("/My Drive/ShortTermStress-IlluminaData/FullExp/Cryp-DE/data/ReviGo_files/CC_dn1h_Revigo.tsv",
#                     colClasses = c(NA,"NULL","NULL","NULL","NULL","NULL","NULL","NULL","NULL","NULL","NULL","NULL","NULL",NA),
#                     header=TRUE) #cellular component results
#ur_1h_cc = read.table("/My Drive/ShortTermStress-IlluminaData/FullExp/Cryp-DE/data/ReviGo_files/CC_up1h_Revigo.tsv",
#                     colClasses = c(NA,"NULL","NULL","NULL","NULL","NULL","NULL","NULL","NULL","NULL","NULL","NULL","NULL",NA),
#                     header=TRUE) #cellular component results
#dr_1h_mf = read.table("/My Drive/ShortTermStress-IlluminaData/FullExp/Cryp-DE/data/ReviGo_files/MF_dn1h_Revigo.tsv",
#                     colClasses = c(NA,"NULL","NULL","NULL","NULL","NULL","NULL","NULL","NULL","NULL","NULL","NULL","NULL",NA),
#                     header=TRUE) #molecular function results
#ur_1h_mf = read.table("/My Drive/ShortTermStress-IlluminaData/FullExp/Cryp-DE/data/ReviGo_files/MF_up1h_Revigo.tsv",
#                     colClasses = c(NA,"NULL","NULL","NULL","NULL","NULL","NULL","NULL","NULL","NULL","NULL","NULL","NULL",NA),
#                     header=TRUE) #molecular function results
##
#dr_10h_bp = read.table("/My Drive/ShortTermStress-IlluminaData/FullExp/Cryp-DE/data/ReviGo_files/BP_dn10h_Revigo.tsv",
#                     colClasses = c(NA,"NULL","NULL","NULL","NULL","NULL","NULL","NULL","NULL","NULL","NULL","NULL","NULL",NA),
#                     header=TRUE) #biological process results
#ur_10h_bp = read.table("/My Drive/ShortTermStress-IlluminaData/FullExp/Cryp-DE/data/ReviGo_files/BP_up10h_Revigo.tsv",
#                     colClasses = c(NA,"NULL","NULL","NULL","NULL","NULL","NULL","NULL","NULL","NULL","NULL","NULL","NULL",NA),
#                     header=TRUE) #biological process results
#dr_10h_cc = read.table("/My Drive/ShortTermStress-IlluminaData/FullExp/Cryp-DE/data/ReviGo_files/CC_dn10h_Revigo.tsv",
#                     colClasses = c(NA,"NULL","NULL","NULL","NULL","NULL","NULL","NULL","NULL","NULL","NULL","NULL","NULL",NA),
#                     header=TRUE) #cellular component results
#ur_10h_cc = read.table("/My Drive/ShortTermStress-IlluminaData/FullExp/Cryp-DE/data/ReviGo_files/CC_up10h_Revigo.tsv",
#                     colClasses = c(NA,"NULL","NULL","NULL","NULL","NULL","NULL","NULL","NULL","NULL","NULL","NULL","NULL",NA),
#                     header=TRUE) #cellular component results
#dr_10h_mf = read.table("/My Drive/ShortTermStress-IlluminaData/FullExp/Cryp-DE/data/ReviGo_files/MF_dn10h_Revigo.tsv",
#                     colClasses = c(NA,"NULL","NULL","NULL","NULL","NULL","NULL","NULL","NULL","NULL","NULL","NULL","NULL",NA),
#                     header=TRUE) #molecular function results
#ur_10h_mf = read.table("/My Drive/ShortTermStress-IlluminaData/FullExp/Cryp-DE/data/ReviGo_files/MF_up10h_Revigo.tsv",
#                     colClasses = c(NA,"NULL","NULL","NULL","NULL","NULL","NULL","NULL","NULL","NULL","NULL","NULL","NULL",NA),
#                     header=TRUE) #molecular function results
#
#names(dr_1h_bp) <- c('GO.ID','Eliminated')
#names(dr_1h_cc) <- c('GO.ID','Eliminated')
#names(dr_1h_mf) <- c('GO.ID','Eliminated')
#names(ur_1h_bp) <- c('GO.ID','Eliminated')
#names(ur_1h_cc) <- c('GO.ID','Eliminated')
#names(ur_1h_mf) <- c('GO.ID','Eliminated')
#
#names(dr_10h_bp) <- c('GO.ID','Eliminated')
#names(dr_10h_cc) <- c('GO.ID','Eliminated')
#names(dr_10h_mf) <- c('GO.ID','Eliminated')
#names(ur_10h_bp) <- c('GO.ID','Eliminated')
#names(ur_10h_cc) <- c('GO.ID','Eliminated')
#names(ur_10h_mf) <- c('GO.ID','Eliminated')
#
#resave(dr_1h_bp,dr_1h_cc,dr_1h_mf,ur_1h_bp,ur_1h_cc,ur_1h_mf,
#       dr_10h_bp,dr_10h_cc,dr_10h_mf,ur_10h_bp,ur_10h_cc,ur_10h_mf,
#       file = "/My Drive/ShortTermStress-IlluminaData/FullExp/Cryp-DE/data/import_rmd_vars.RData")
```

### 4.4.1. Comparing 60 min and 10 h enrichments

#### 4.4.1.1 For a quick glance, we compiled it all into a single table. 

```{r}
ur_10h_bp <- up0v10h_BP_elim %>% 
  full_join(., ur_10h_bp, by = "GO.ID", copy = FALSE, suffix = c(".x",".y"))
ur_10h_bp$timept <- '10h'
ur_10h_bp$category <- 'biological process'
ur_10h_bp$degree <- 'upregulated'
ur_10h_cc <- up0v10h_CC_elim %>% 
  full_join(., ur_10h_cc, by = "GO.ID", copy = FALSE, suffix = c(".x",".y"))
ur_10h_cc$timept <- '10h'
ur_10h_cc$category <- 'cellular component'
ur_10h_cc$degree <- 'upregulated'
ur_10h_mf <- up0v10h_MF_elim %>% 
  full_join(., ur_10h_mf, by = "GO.ID", copy = FALSE, suffix = c(".x",".y"))
ur_10h_mf$timept <- '10h'
ur_10h_mf$category <- 'molecular function'
ur_10h_mf$degree <- 'upregulated'
dr_10h_bp <- down0v10h_BP_elim %>% 
  full_join(., dr_10h_bp, by = "GO.ID", copy = FALSE, suffix = c(".x",".y"))
dr_10h_bp$timept <- '10h'
dr_10h_bp$category <- 'biological process'
dr_10h_bp$degree <- 'downregulated'
dr_10h_cc <- down0v10h_CC_elim %>% 
  full_join(., dr_10h_cc, by = "GO.ID", copy = FALSE, suffix = c(".x",".y"))
dr_10h_cc$timept <- '10h'
dr_10h_cc$category <- 'cellular component'
dr_10h_cc$degree <- 'downregulated'
dr_10h_mf <- down0v10h_MF_elim %>% 
  full_join(., dr_10h_mf, by = "GO.ID", copy = FALSE, suffix = c(".x",".y"))
dr_10h_mf$timept <- '10h'
dr_10h_mf$category <- 'molecular function'
dr_10h_mf$degree <- 'downregulated'

ur_1h_bp <- up0v1h_BP_elim %>% 
  full_join(., ur_1h_bp, by = "GO.ID", copy = FALSE, suffix = c(".x",".y"))
ur_1h_bp$timept <- '1h'
ur_1h_bp$category <- 'biological process'
ur_1h_bp$degree <- 'upregulated'
ur_1h_cc <- up0v1h_CC_elim %>% 
  full_join(., ur_1h_cc, by = "GO.ID", copy = FALSE, suffix = c(".x",".y"))
ur_1h_cc$timept <- '1h'
ur_1h_cc$category <- 'cellular component'
ur_1h_cc$degree <- 'upregulated'
ur_1h_mf <- up0v1h_MF_elim %>% 
  full_join(., ur_1h_mf, by = "GO.ID", copy = FALSE, suffix = c(".x",".y"))
ur_1h_mf$timept <- '1h'
ur_1h_mf$category <- 'molecular function'
ur_1h_mf$degree <- 'upregulated'
dr_1h_bp <- down0v1h_BP_elim %>% 
  full_join(., dr_1h_bp, by = "GO.ID", copy = FALSE, suffix = c(".x",".y"))
dr_1h_bp$timept <- '1h'
dr_1h_bp$category <- 'biological process'
dr_1h_bp$degree <- 'downregulated'
dr_1h_cc <- down0v1h_CC_elim %>% 
  full_join(., dr_1h_cc, by = "GO.ID", copy = FALSE, suffix = c(".x",".y"))
dr_1h_cc$timept <- '1h'
dr_1h_cc$category <- 'cellular component'
dr_1h_cc$degree <- 'downregulated'
dr_1h_mf <- down0v1h_MF_elim %>% 
  full_join(., dr_1h_mf, by = "GO.ID", copy = FALSE, suffix = c(".x",".y"))
dr_1h_mf$timept <- '1h'
dr_1h_mf$category <- 'molecular function'
dr_1h_mf$degree <- 'downregulated'

#After reviewing the cellular component data, we opted not to include it in the comparison between timepoints since it didn't provide much additional depth.
go_data <- rbind(ur_10h_bp,ur_10h_mf,dr_10h_bp,dr_10h_mf,ur_1h_bp,ur_1h_mf,dr_1h_bp,dr_1h_mf)

names(go_data) <- c("GO.ID","Term","Annotated","Sign","Exp","Pvalue","Eliminated","timept","category","degree")
go_data$GO.Term <- paste(go_data$Term, go_data$GO.ID, sep=" ")
go_data_reduced <- filter(go_data, Eliminated == "False") 
go_data_reduced <- subset(go_data_reduced,select=c("GO.Term","Annotated","Pvalue","timept","category","degree"))
go_data_reduced <- transform(go_data_reduced, Percentage = (Annotated / 3193) * 100)
go_data_reduced <- subset(go_data_reduced, (`Percentage`) > 1) 
go_data_reduced <- subset(go_data_reduced, (`Pvalue`) < 0.05) 
go_data_reduced <- go_data_reduced %>%
  mutate(Percentage = ifelse(degree == "upregulated",
                             Percentage,
                             -1*Percentage))
go_data_reduced$Pvalue<-gsub("<","",as.character(go_data_reduced$Pvalue))
```

#### 4.4.1.2 Create the variables for the graphs and plot.

```{r fig.width=10, fig.height=8}
#Data for graph

data_1h <- filter(go_data_reduced, timept == "1h")
data_1h <- data_1h[
    order( data_1h[,5], data_1h[,6], data_1h[,7]),
]
data_1h$GO.Term <- factor(data_1h$GO.Term, levels = data_1h$GO.Term)
data_1h$Pvalue <- as.numeric(data_1h$Pvalue)

data_10h <- filter(go_data_reduced, timept == "10h")
data_10h <- data_10h[
    order( data_10h[,5], data_10h[,6], data_10h[,7]),
]
data_10h$GO.Term <- factor(data_10h$GO.Term, levels = data_10h$GO.Term)
data_10h$Pvalue <- as.numeric(data_10h$Pvalue)
y_breaks <- pretty(go_data_reduced$Percentage)

GO_plot_1h <- ggplot(data_1h,
                      aes(x=GO.Term,
                         y=Percentage,
                         fill=Pvalue)) +
  geom_bar(stat = "identity") +
  coord_flip()+
  ylim(-25,25) +
  xlab("Enriched Gene Ontology Terms") + 
  ylab("Percentage of Significant DEG") +
  ggtitle("0m vs 1h") +
  theme_classic() +
  theme(plot.title = element_text(hjust = 0.4))

GO_plot_10h <- ggplot(data_10h,
                      aes(x=GO.Term,
                         y=Percentage,
                         fill=Pvalue)) +
  geom_bar(stat = "identity") +
  coord_flip() +
  ylim(-25,25) +
  xlab("Enriched Gene Ontology Terms") + 
  ylab("Percentage of Significant DEG") +
  ggtitle("0m vs 10h") +
  theme_classic() +
  theme(plot.title = element_text(hjust = 0.4))

#Figure plotting
(GO_plot_1h / GO_plot_10h)
```


#### 4.4.1.3 Hypergeometric tests

We also looked at the significance of shared DE genes between 60 min, 10 h, and the long-term datasets.

```{r}
#Hypergeometric assessment of overlap of DE genes between basically everything we can think of.
genes_1h_up <- rownames(logFCup_0v1h)
genes_1h_down <- rownames(logFCdown_0v1h)
genes_10h_up <- rownames(logFCup_0v10h)
genes_10h_down <- rownames(logFCdown_0v10h)
genes_LT_up <- rownames(nakov_DE_ur)
genes_LT_down <- rownames(nakov_DE_dr)
genes_1h <- rownames(sign2LFC_0v1h)
genes_10h <- rownames(sign2LFC_0v10h)
genes_LT <- rownames(nakov_DE)
genes_all_signDE <- rownames(Padjscreen_sorted)
genes_all <- rownames(OnlySignGenes_ConStage)
genes_2 <- rownames(DEG2sig_results)

#ID the overlapping genes

int_up <- intersect (genes_1h_up, genes_10h_up)
int_down <- intersect (genes_1h_down, genes_10h_down)
int_updown <- intersect (genes_1h_up, genes_10h_down)
int_downup <- intersect (genes_1h_down, genes_10h_up)
int_1v10 <- intersect (genes_1h, genes_10h)
int_LTv1_up <- intersect (genes_1h_up, genes_LT_up)
int_LTv10_up <- intersect (genes_10h_up, genes_LT_up)
int_LTv1_dn <- intersect (genes_1h_down, genes_LT_down)
int_LTv10_dn <- intersect (genes_10h_down, genes_LT_down)
int_LTv1_updown <- intersect (genes_1h_up, genes_LT_down)
int_LTv1_downup <- intersect (genes_1h_down, genes_LT_up)
int_LTv10_updown <- intersect (genes_LT_up, genes_10h_down)
int_LTv10_downup <- intersect (genes_LT_down, genes_10h_up)
int_LTv10 <- intersect (genes_10h, genes_LT)
int_LTv1h <- intersect (genes_1h, genes_LT)
int_LTvAll <- intersect (genes_all_signDE, genes_LT)
int_All <- intersect (genes_all, genes_LT)
int_LTv2sig <- intersect (genes_2, genes_LT)

#Execute the hypergeometric test to determine if that number of overlapping genes is significant.

phyper(416,417,(2048-417),772,lower.tail = FALSE) # 1h vs 10h repressed (8.349016e-186)
phyper(243,244,(1364-244),660,lower.tail = FALSE) #1h vs 10h induced (1.496532e-113)
phyper(56,57,(1364-57),772,lower.tail = FALSE) #10h repressed vs 1h induced (8.057105e-42)
phyper(92,93,(2048-93),660,lower.tail = FALSE) #1h repressed vs 10h induced (7.854042e-106)

phyper(969,970,(3497-970),1431,lower.tail = FALSE) #1h signDE vs 10h signDE (0?????)
phyper(258,259,(4298-259),1220,lower.tail = FALSE) #All short-term signDE vs long-term signDE (9.145136e-152)

phyper(79,80,(1431-80),1220,lower.tail = FALSE) #All 10h signDE vs LT signDE (1.926452e-06)
phyper(205,206,(3497-206),1220,lower.tail = FALSE) #All 1h signDE vs LT signDE (2.592496e-103)
phyper(50,51,(1942-51),695,lower.tail = FALSE)#1h vs LT repressed (5.145085e-24)
phyper(22,23,(772-23),695,lower.tail = FALSE)#10h vs LT repressed 0.0859712)
phyper(42,43,(1555-43),525,lower.tail = FALSE)#1h vs LT induced (1.617967e-21)
phyper(17,18,(659-18),525,lower.tail = FALSE)#10h vs LT induced (0.01572783)
phyper(53,54,(1942-54),525,lower.tail = FALSE)#1h repressed vs LT induced (2.630185e-32)
phyper(16,17,(772-17),525,lower.tail = FALSE)#10h repressed vs LT induced (0.001307894
phyper(63,64,(1555-64),695,lower.tail = FALSE)#1h induced vs LT repressed (7.711969e-24)
phyper(21,22,(695-22),659,lower.tail = FALSE)#10h induced vs LT repressed (0.3046189)
```

### 4.4.2 GO enrichment for different LFC cutoffs for 1h and 10 h. 

We started with the genes at 1h that had an LFC less than 0.585 to look at the primary processes enriched in lowly expressed genes during peak hyposalinity stress.

```{r}
##This is for looking at LFC 0.585 in 1 h 
LFCupless0.585_0v1h <- subset(sign2LFC_0v1h, (`0v1h_Padj`) <= 0.01 & (`0v1h`) < 0.585 & (`0v1h`) > 0 )
LFCdnless0.585_0v1h <- subset(sign2LFC_0v1h, (`0v1h_Padj`) <= 0.01 & (`0v1h`) > -0.585 & (`0v1h`) < 0)

LFCupless0.585_0v1h_GOEnrich  = rownames(subset(LFCupless0.585_0v1h, rownames(LFCupless0.585_0v1h)%in%genesOfInterest_allSign))
LFCdnless0.585_0v1h_GOEnrich  = rownames(subset(LFCdnless0.585_0v1h, rownames(LFCdnless0.585_0v1h)%in%genesOfInterest_allSign))


#create gene list for input in topGO
geneList_up1hLFCless0.585 = factor(as.integer(geneUniverse %in% LFCupless0.585_0v1h_GOEnrich))
names(geneList_up1hLFCless0.585) = geneUniverse
str(geneList_up1hLFCless0.585)
geneList_dn1hLFCless0.585 = factor(as.integer(geneUniverse %in% LFCdnless0.585_0v1h_GOEnrich))
names(geneList_dn1hLFCless0.585) = geneUniverse
str(geneList_dn1hLFCless0.585)

##create a topGO object (for up GOs)
GOdata_BP_up1hLFCless0.585 = new("topGOdata", ontology="BP", allGenes=geneList_up1hLFCless0.585, 
                        annot = annFUN.gene2GO, gene2GO = geneID2GO)
GOdata_MF_up1hLFCless0.585 = new("topGOdata", ontology="MF", allGenes=geneList_up1hLFCless0.585, 
                        annot = annFUN.gene2GO, gene2GO = geneID2GO)
GOdata_CC_up1hLFCless0.585 = new("topGOdata", ontology="CC", allGenes=geneList_up1hLFCless0.585, 
                        annot = annFUN.gene2GO, gene2GO = geneID2GO)
##create a topGO object (for down GOs)
GOdata_BP_dn1hLFCless0.585  = new("topGOdata", ontology="BP", allGenes=geneList_dn1hLFCless0.585, 
                           annot = annFUN.gene2GO, gene2GO = geneID2GO)
GOdata_MF_dn1hLFCless0.585  = new("topGOdata", ontology="MF", allGenes=geneList_dn1hLFCless0.585, 
                           annot = annFUN.gene2GO, gene2GO = geneID2GO)
GOdata_CC_dn1hLFCless0.585  = new("topGOdata", ontology="CC", allGenes=geneList_dn1hLFCless0.585, 
                           annot = annFUN.gene2GO, gene2GO = geneID2GO)

#run Fisher's exact test
resultFisher_BP_up1hLFCless0.585_elim <- runTest(GOdata_BP_up1hLFCless0.585, algorithm = "elim", statistic = "fisher")
resultFisher_MF_up1hLFCless0.585_elim <- runTest(GOdata_MF_up1hLFCless0.585, algorithm = "elim", statistic = "fisher")
resultFisher_CC_up1hLFCless0.585_elim <- runTest(GOdata_CC_up1hLFCless0.585, algorithm = "elim", statistic = "fisher")
resultFisher_BP_dn1hLFCless0.585_elim <- runTest(GOdata_BP_dn1hLFCless0.585, algorithm = "elim", statistic = "fisher")
resultFisher_MF_dn1hLFCless0.585_elim <- runTest(GOdata_MF_dn1hLFCless0.585, algorithm = "elim", statistic = "fisher")
resultFisher_CC_dn1hLFCless0.585_elim <- runTest(GOdata_CC_dn1hLFCless0.585, algorithm = "elim", statistic = "fisher")

#extract the significant GO terms
up1hLFCless0.585_BP_elim <- GenTable(GOdata_BP_up1hLFCless0.585, classic = resultFisher_BP_up1hLFCless0.585_elim, 
                            orderBy = "weight", ranksOf = "weight", topNodes = 50)
up1hLFCless0.585_MF_elim <- GenTable(GOdata_MF_up1hLFCless0.585, classic = resultFisher_MF_up1hLFCless0.585_elim, 
                            orderBy = "weight", ranksOf = "weight", topNodes = 50)
up1hLFCless0.585_CC_elim <- GenTable(GOdata_CC_up1hLFCless0.585, classic = resultFisher_CC_up1hLFCless0.585_elim, 
                            orderBy = "weight", ranksOf = "weight", topNodes = 50)
dn1hLFCless0.585_BP_elim <- GenTable(GOdata_BP_dn1hLFCless0.585, classic = resultFisher_BP_dn1hLFCless0.585_elim, 
                              orderBy = "weight", ranksOf = "weight", topNodes = 50)
dn1hLFCless0.585_MF_elim <- GenTable(GOdata_MF_dn1hLFCless0.585, classic = resultFisher_MF_dn1hLFCless0.585_elim, 
                              orderBy = "weight", ranksOf = "weight", topNodes = 50)
dn1hLFCless0.585_CC_elim <- GenTable(GOdata_CC_dn1hLFCless0.585, classic = resultFisher_CC_dn1hLFCless0.585_elim, 
                              orderBy = "weight", ranksOf = "weight", topNodes = 50)

#write.table(up1hLFCless0.585_BP_elim,
#            "/My Drive/ShortTermStress-IlluminaData/FullExp/Cryp-DE/data/ReviGo_files/BP_up0.585_elim.txt")
#write.table(up1hLFCless0.585_MF_elim,
#            "/My Drive/ShortTermStress-IlluminaData/FullExp/Cryp-DE/data/ReviGo_files/MF_up0.585_elim.txt")
#write.table(up1hLFCless0.585_CC_elim,
#            "/My Drive/ShortTermStress-IlluminaData/FullExp/Cryp-DE/data/ReviGo_files/CC_up0.585_elim.txt")
#write.table(dn1hLFCless0.585_BP_elim,
#            "/My Drive/ShortTermStress-IlluminaData/FullExp/Cryp-DE/data/ReviGo_files/BP_dn0.585_elim.txt")
#write.table(dn1hLFCless0.585_MF_elim,
#            "/My Drive/ShortTermStress-IlluminaData/FullExp/Cryp-DE/data/ReviGo_files/MF_dn0.585_elim.txt")
#write.table(dn1hLFCless0.585_CC_elim,
#            "/My Drive/ShortTermStress-IlluminaData/FullExp/Cryp-DE/data/ReviGo_files/CC_dn0.585_elim.txt")

#Not used in further anayses, so we didn't import it back into the code after revigo.
```

Enrichment analyses on timepoints 1 h and 10 h for genes with an LFC < -1 and > 1

```{r}
##This is for looking at LFC 1 in 1 h and 10 h time points
LFCup1_0v1h <- subset(sign2LFC_0v1h, (`0v1h_Padj`) <= 0.01 & (`0v1h`) > 1)
LFCdn1_0v1h <- subset(sign2LFC_0v1h, (`0v1h_Padj`) <= 0.01 & (`0v1h`) < -1)

LFCup1_0v1h_GOEnrich  = rownames(subset(LFCup1_0v1h, rownames(LFCup1_0v1h)%in%genesOfInterest_allSign))
LFCdn1_0v1h_GOEnrich  = rownames(subset(LFCdn1_0v1h, rownames(LFCdn1_0v1h)%in%genesOfInterest_allSign))


#create gene list for input in topGO for cluster 1
geneList_up1hLFC1 = factor(as.integer(geneUniverse %in% LFCup1_0v1h_GOEnrich))
names(geneList_up1hLFC1) = geneUniverse
str(geneList_up1hLFC1)
geneList_dn1hLFC1 = factor(as.integer(geneUniverse %in% LFCdn1_0v1h_GOEnrich))
names(geneList_dn1hLFC1) = geneUniverse
str(geneList_dn1hLFC1)

##create a topGO object (for up GOs)
GOdata_BP_up1hLFC1 = new("topGOdata", ontology="BP", allGenes=geneList_up1hLFC1, 
                        annot = annFUN.gene2GO, gene2GO = geneID2GO)
GOdata_MF_up1hLFC1 = new("topGOdata", ontology="MF", allGenes=geneList_up1hLFC1, 
                        annot = annFUN.gene2GO, gene2GO = geneID2GO)
GOdata_CC_up1hLFC1 = new("topGOdata", ontology="CC", allGenes=geneList_up1hLFC1, 
                        annot = annFUN.gene2GO, gene2GO = geneID2GO)
##create a topGO object (for down GOs)
GOdata_BP_dn1hLFC1  = new("topGOdata", ontology="BP", allGenes=geneList_dn1hLFC1 , 
                           annot = annFUN.gene2GO, gene2GO = geneID2GO)
GOdata_MF_dn1hLFC1  = new("topGOdata", ontology="MF", allGenes=geneList_dn1hLFC1 , 
                           annot = annFUN.gene2GO, gene2GO = geneID2GO)
GOdata_CC_dn1hLFC1  = new("topGOdata", ontology="CC", allGenes=geneList_dn1hLFC1 , 
                           annot = annFUN.gene2GO, gene2GO = geneID2GO)

#run Fisher's exact test
resultFisher_BP_up1hLFC1_elim <- runTest(GOdata_BP_up1hLFC1, algorithm = "elim", statistic = "fisher")
resultFisher_MF_up1hLFC1_elim <- runTest(GOdata_MF_up1hLFC1, algorithm = "elim", statistic = "fisher")
resultFisher_CC_up1hLFC1_elim <- runTest(GOdata_CC_up1hLFC1, algorithm = "elim", statistic = "fisher")
resultFisher_BP_dn1hLFC1_elim <- runTest(GOdata_BP_dn1hLFC1, algorithm = "elim", statistic = "fisher")
resultFisher_MF_dn1hLFC1_elim <- runTest(GOdata_MF_dn1hLFC1, algorithm = "elim", statistic = "fisher")
resultFisher_CC_dn1hLFC1_elim <- runTest(GOdata_CC_dn1hLFC1, algorithm = "elim", statistic = "fisher")

#extract the significant GO terms
up1hLFC1_BP_elim <- GenTable(GOdata_BP_up1hLFC1, classic = resultFisher_BP_up1hLFC1_elim, 
                            orderBy = "weight", ranksOf = "weight", topNodes = 50)
up1hLFC1_MF_elim <- GenTable(GOdata_MF_up1hLFC1, classic = resultFisher_MF_up1hLFC1_elim, 
                            orderBy = "weight", ranksOf = "weight", topNodes = 50)
up1hLFC1_CC_elim <- GenTable(GOdata_CC_up1hLFC1, classic = resultFisher_CC_up1hLFC1_elim, 
                            orderBy = "weight", ranksOf = "weight", topNodes = 50)
dn1hLFC1_BP_elim <- GenTable(GOdata_BP_dn1hLFC1, classic = resultFisher_BP_dn1hLFC1_elim, 
                              orderBy = "weight", ranksOf = "weight", topNodes = 50)
dn1hLFC1_MF_elim <- GenTable(GOdata_MF_dn1hLFC1, classic = resultFisher_MF_dn1hLFC1_elim, 
                              orderBy = "weight", ranksOf = "weight", topNodes = 50)
dn1hLFC1_CC_elim <- GenTable(GOdata_CC_dn1hLFC1, classic = resultFisher_CC_dn1hLFC1_elim, 
                              orderBy = "weight", ranksOf = "weight", topNodes = 50)

##Time 0 vs 10 hour
LFCup1_0v10h <- subset(sign2LFC_0v10h, (`0v10h_Padj`) <= 0.01 & (`0v10h`) > 1)
LFCdn1_0v10h <- subset(sign2LFC_0v10h, (`0v10h_Padj`) <= 0.01 & (`0v10h`) < -1)

LFCup1_0v10h_GOEnrich  = rownames(subset(LFCup1_0v10h, rownames(LFCup1_0v10h)%in%genesOfInterest_allSign))
LFCdn1_0v10h_GOEnrich  = rownames(subset(LFCdn1_0v10h, rownames(LFCdn1_0v10h)%in%genesOfInterest_allSign))


#create gene list for input in topGO for cluster 1
geneList_up10hLFC1 = factor(as.integer(geneUniverse %in% LFCup1_0v10h_GOEnrich))
names(geneList_up10hLFC1) = geneUniverse
str(geneList_up10hLFC1)
geneList_dn10hLFC1 = factor(as.integer(geneUniverse %in% LFCdn1_0v10h_GOEnrich))
names(geneList_dn10hLFC1) = geneUniverse
str(geneList_dn10hLFC1)

##create a topGO object (for up GOs)
GOdata_BP_up10hLFC1 = new("topGOdata", ontology="BP", allGenes=geneList_up10hLFC1, 
                        annot = annFUN.gene2GO, gene2GO = geneID2GO)
GOdata_MF_up10hLFC1 = new("topGOdata", ontology="MF", allGenes=geneList_up10hLFC1, 
                        annot = annFUN.gene2GO, gene2GO = geneID2GO)
GOdata_CC_up10hLFC1 = new("topGOdata", ontology="CC", allGenes=geneList_up10hLFC1, 
                        annot = annFUN.gene2GO, gene2GO = geneID2GO)
##create a topGO object (for down GOs)
GOdata_BP_dn10hLFC1  = new("topGOdata", ontology="BP", allGenes=geneList_dn10hLFC1 , 
                           annot = annFUN.gene2GO, gene2GO = geneID2GO)
GOdata_MF_dn10hLFC1  = new("topGOdata", ontology="MF", allGenes=geneList_dn10hLFC1 , 
                           annot = annFUN.gene2GO, gene2GO = geneID2GO)
GOdata_CC_dn10hLFC1  = new("topGOdata", ontology="CC", allGenes=geneList_dn10hLFC1 , 
                           annot = annFUN.gene2GO, gene2GO = geneID2GO)

#run Fisher's exact test
resultFisher_BP_up10hLFC1_elim <- runTest(GOdata_BP_up10hLFC1, algorithm = "elim", statistic = "fisher")
resultFisher_MF_up10hLFC1_elim <- runTest(GOdata_MF_up10hLFC1, algorithm = "elim", statistic = "fisher")
resultFisher_CC_up10hLFC1_elim <- runTest(GOdata_CC_up10hLFC1, algorithm = "elim", statistic = "fisher")
resultFisher_BP_dn10hLFC1_elim <- runTest(GOdata_BP_dn10hLFC1, algorithm = "elim", statistic = "fisher")
resultFisher_MF_dn10hLFC1_elim <- runTest(GOdata_MF_dn10hLFC1, algorithm = "elim", statistic = "fisher")
resultFisher_CC_dn10hLFC1_elim <- runTest(GOdata_CC_dn10hLFC1, algorithm = "elim", statistic = "fisher")

#extract the significant GO terms
up10hLFC1_BP_elim <- GenTable(GOdata_BP_up10hLFC1, classic = resultFisher_BP_up10hLFC1_elim, 
                            orderBy = "weight", ranksOf = "weight", topNodes = 50)
up10hLFC1_MF_elim <- GenTable(GOdata_MF_up10hLFC1, classic = resultFisher_MF_up10hLFC1_elim, 
                            orderBy = "weight", ranksOf = "weight", topNodes = 50)
up10hLFC1_CC_elim <- GenTable(GOdata_CC_up10hLFC1, classic = resultFisher_CC_up10hLFC1_elim, 
                            orderBy = "weight", ranksOf = "weight", topNodes = 50)
dn10hLFC1_BP_elim <- GenTable(GOdata_BP_dn10hLFC1, classic = resultFisher_BP_dn10hLFC1_elim, 
                              orderBy = "weight", ranksOf = "weight", topNodes = 50)
dn10hLFC1_MF_elim <- GenTable(GOdata_MF_dn10hLFC1, classic = resultFisher_MF_dn10hLFC1_elim, 
                              orderBy = "weight", ranksOf = "weight", topNodes = 50)
dn10hLFC1_CC_elim <- GenTable(GOdata_CC_dn10hLFC1, classic = resultFisher_CC_dn10hLFC1_elim, 
                              orderBy = "weight", ranksOf = "weight", topNodes = 50)

#write.table(up1hLFC1_BP_elim,
#            "/My Drive/ShortTermStress-IlluminaData/FullExp/Cryp-DE/data/ReviGo_files/BP_up1hLFC1_elim.txt")
#write.table(up1hLFC1_MF_elim,
#            "/My Drive/ShortTermStress-IlluminaData/FullExp/Cryp-DE/data/ReviGo_files/MF_up1hLFC1_elim.txt")
#write.table(up1hLFC1_CC_elim,
#            "/My Drive/ShortTermStress-IlluminaData/FullExp/Cryp-DE/data/ReviGo_files/CC_up1hLFC1_elim.txt")
#write.table(dn1hLFC1_BP_elim,
#            "/My Drive/ShortTermStress-IlluminaData/FullExp/Cryp-DE/data/ReviGo_files/BP_dn1hLFC1_elim.txt")
#write.table(dn1hLFC1_MF_elim,
#            "/My Drive/ShortTermStress-IlluminaData/FullExp/Cryp-DE/data/ReviGo_files/MF_dn1hLFC1_elim.txt")
#write.table(dn1hLFC1_CC_elim,
#            "/My Drive/ShortTermStress-IlluminaData/FullExp/Cryp-DE/data/ReviGo_files/CC_dn1hLFC1_elim.txt")

#write.table(up10hLFC1_BP_elim,
#            "/My Drive/ShortTermStress-IlluminaData/FullExp/Cryp-DE/data/ReviGo_files/BP_up10hLFC1_elim.txt")
#write.table(up10hLFC1_MF_elim,
#            "/My Drive/ShortTermStress-IlluminaData/FullExp/Cryp-DE/data/ReviGo_files/MF_up10hLFC1_elim.txt")
#write.table(up10hLFC1_CC_elim,
#            "/My Drive/ShortTermStress-IlluminaData/FullExp/Cryp-DE/data/ReviGo_files/CC_up10hLFC1_elim.txt")
#write.table(dn10hLFC1_BP_elim,
#            "/My Drive/ShortTermStress-IlluminaData/FullExp/Cryp-DE/data/ReviGo_files/BP_dn10hLFC1_elim.txt")
#write.table(dn10hLFC1_MF_elim,
#            "/My Drive/ShortTermStress-IlluminaData/FullExp/Cryp-DE/data/ReviGo_files/MF_dn10hLFC1_elim.txt")
#write.table(dn10hLFC1_CC_elim,
#            "/My Drive/ShortTermStress-IlluminaData/FullExp/Cryp-DE/data/ReviGo_files/CC_dn10hLFC1_elim.txt")
```

Import the data back in from ReviGO

```{r}
dr_1hLFC1_bp = read.table("/My Drive/ShortTermStress-IlluminaData/FullExp/Cryp-DE/data/ReviGo_files/BP_dn1hLFC1.tsv",
                     colClasses = c(NA,"NULL","NULL","NULL","NULL","NULL","NULL","NULL","NULL","NULL","NULL","NULL","NULL",NA),
                     header=TRUE) #biological process results
ur_1hLFC1_bp = read.table("/My Drive/ShortTermStress-IlluminaData/FullExp/Cryp-DE/data/ReviGo_files/BP_up1hLFC1.tsv",
                     colClasses = c(NA,"NULL","NULL","NULL","NULL","NULL","NULL","NULL","NULL","NULL","NULL","NULL","NULL",NA),
                     header=TRUE) #biological process results
dr_1hLFC1_mf = read.table("/My Drive/ShortTermStress-IlluminaData/FullExp/Cryp-DE/data/ReviGo_files/MF_dn1hLFC1.tsv",
                     colClasses = c(NA,"NULL","NULL","NULL","NULL","NULL","NULL","NULL","NULL","NULL","NULL","NULL","NULL",NA),
                     header=TRUE) #molecular function results
ur_1hLFC1_mf = read.table("/My Drive/ShortTermStress-IlluminaData/FullExp/Cryp-DE/data/ReviGo_files/MF_up1hLFC1.tsv",
                     colClasses = c(NA,"NULL","NULL","NULL","NULL","NULL","NULL","NULL","NULL","NULL","NULL","NULL","NULL",NA),
                     header=TRUE) #molecular function results

dr_10hLFC1_bp = read.table("/My Drive/ShortTermStress-IlluminaData/FullExp/Cryp-DE/data/ReviGo_files/BP_dn10hLFC1.tsv",
                     colClasses = c(NA,"NULL","NULL","NULL","NULL","NULL","NULL","NULL","NULL","NULL","NULL","NULL","NULL",NA),
                     header=TRUE) #biological process results
ur_10hLFC1_bp = read.table("/My Drive/ShortTermStress-IlluminaData/FullExp/Cryp-DE/data/ReviGo_files/BP_up10hLFC1.tsv",
                     colClasses = c(NA,"NULL","NULL","NULL","NULL","NULL","NULL","NULL","NULL","NULL","NULL","NULL","NULL",NA),
                     header=TRUE) #biological process results
dr_10hLFC1_mf = read.table("/My Drive/ShortTermStress-IlluminaData/FullExp/Cryp-DE/data/ReviGo_files/MF_dn10hLFC1.tsv",
                     colClasses = c(NA,"NULL","NULL","NULL","NULL","NULL","NULL","NULL","NULL","NULL","NULL","NULL","NULL",NA),
                     header=TRUE) #molecular function results
ur_10hLFC1_mf = read.table("/My Drive/ShortTermStress-IlluminaData/FullExp/Cryp-DE/data/ReviGo_files/MF_up10hLFC1.tsv",
                     colClasses = c(NA,"NULL","NULL","NULL","NULL","NULL","NULL","NULL","NULL","NULL","NULL","NULL","NULL",NA),
                     header=TRUE) #molecular function results

#
names(dr_1hLFC1_bp) <- c('GO.ID','Eliminated')
names(dr_1hLFC1_mf) <- c('GO.ID','Eliminated')
names(ur_1hLFC1_bp) <- c('GO.ID','Eliminated')
names(ur_1hLFC1_mf) <- c('GO.ID','Eliminated')

names(dr_10hLFC1_bp) <- c('GO.ID','Eliminated')
names(dr_10hLFC1_mf) <- c('GO.ID','Eliminated')
names(ur_10hLFC1_bp) <- c('GO.ID','Eliminated')
names(ur_10hLFC1_mf) <- c('GO.ID','Eliminated')
#

#resave(dr_1hLFC1_bp,dr_1hLFC1_mf,ur_1hLFC1_bp,ur_1hLFC1_mf,
#       dr_10hLFC1_bp,dr_10hLFC1_mf,ur_10hLFC1_bp,ur_10hLFC1_mf,
#       file = "/My Drive/ShortTermStress-IlluminaData/FullExp/Cryp-DE/data/import_rmd_vars.RData")
```

Make a table for quick comparison of the genes that meet the LFC cutoff for 1 h and 10 h 

```{r}
ur_10hLFC1_bp <- up10hLFC1_BP_elim %>% 
  full_join(., ur_10hLFC1_bp, by = "GO.ID", copy = FALSE, suffix = c(".x",".y"))
ur_10hLFC1_bp$timept <- '10h'
ur_10hLFC1_bp$category <- 'biological process'
ur_10hLFC1_bp$degree <- 'upregulated'
ur_10hLFC1_mf <- up10hLFC1_MF_elim %>% 
  full_join(., ur_10hLFC1_mf, by = "GO.ID", copy = FALSE, suffix = c(".x",".y"))
ur_10hLFC1_mf$timept <- '10h'
ur_10hLFC1_mf$category <- 'molecular function'
ur_10hLFC1_mf$degree <- 'upregulated'
dr_10hLFC1_bp <- dn10hLFC1_BP_elim %>% 
  full_join(., dr_10hLFC1_bp, by = "GO.ID", copy = FALSE, suffix = c(".x",".y"))
dr_10hLFC1_bp$timept <- '10h'
dr_10hLFC1_bp$category <- 'biological process'
dr_10hLFC1_bp$degree <- 'downregulated'
dr_10hLFC1_mf <- dn10hLFC1_MF_elim %>% 
  full_join(., dr_10hLFC1_mf, by = "GO.ID", copy = FALSE, suffix = c(".x",".y"))
dr_10hLFC1_mf$timept <- '10h'
dr_10hLFC1_mf$category <- 'molecular function'
dr_10hLFC1_mf$degree <- 'downregulated'

ur_1hLFC1_bp <- up1hLFC1_BP_elim %>% 
  full_join(., ur_1hLFC1_bp, by = "GO.ID", copy = FALSE, suffix = c(".x",".y"))
ur_1hLFC1_bp$timept <- '1h'
ur_1hLFC1_bp$category <- 'biological process'
ur_1hLFC1_bp$degree <- 'upregulated'
ur_1hLFC1_mf <- up1hLFC1_MF_elim %>% 
  full_join(., ur_1hLFC1_mf, by = "GO.ID", copy = FALSE, suffix = c(".x",".y"))
ur_1hLFC1_mf$timept <- '1h'
ur_1hLFC1_mf$category <- 'molecular function'
ur_1hLFC1_mf$degree <- 'upregulated'
dr_1hLFC1_bp <- dn1hLFC1_BP_elim %>% 
  full_join(., dr_1hLFC1_bp, by = "GO.ID", copy = FALSE, suffix = c(".x",".y"))
dr_1hLFC1_bp$timept <- '1h'
dr_1hLFC1_bp$category <- 'biological process'
dr_1hLFC1_bp$degree <- 'downregulated'
dr_1hLFC1_mf <- dn1hLFC1_MF_elim %>% 
  full_join(., dr_1hLFC1_mf, by = "GO.ID", copy = FALSE, suffix = c(".x",".y"))
dr_1hLFC1_mf$timept <- '1h'
dr_1hLFC1_mf$category <- 'molecular function'
dr_1hLFC1_mf$degree <- 'downregulated'

go_data_LFC1 <- rbind(ur_10hLFC1_bp,ur_10hLFC1_mf,dr_10hLFC1_bp,dr_10hLFC1_mf)

names(go_data_LFC1) <- c("GO.ID","Term","Annotated","Sign","Exp","Pvalue","Eliminated","timept","category","degree")
go_data_LFC1$GO.Term <- paste(go_data_LFC1$Term, go_data_LFC1$GO.ID, sep=" ")
go_data_LFC1_reduced <- filter(go_data_LFC1, Eliminated == " False") 
go_data_LFC1_reduced <- subset(go_data_LFC1_reduced,select=c("GO.Term","Annotated","Pvalue","timept","category","degree"))
go_data_LFC1_reduced <- transform(go_data_LFC1_reduced, Percentage = (Annotated / 3193) * 100)
go_data_LFC1_reduced <- subset(go_data_LFC1_reduced, (`Percentage`) > 0.5 & (`Percentage`) < 23) 
go_data_LFC1_reduced <- subset(go_data_LFC1_reduced, (`Pvalue`) < 0.05 |(`Pvalue`) > 0.06) #Necessary to remove > 0.05 but keep e- data
go_data_LFC1_reduced <- go_data_LFC1_reduced %>%
  mutate(Percentage = ifelse(degree == "upregulated",
                             Percentage,
                             -1*Percentage))
```


## 4.5 Clusters identified from TreeView (centroid linkage)

Cluster 1

```{r}
#Cluster 1 
clust1_genes = read.delim("/My Drive/ShortTermStress-IlluminaData/FullExp/Cryp-DE/data/Clusters_11-19/sign2LFC_Holm_C1.txt",header = TRUE,row.names = 1)
clust1_genes = clust1_genes[-1, ]

clust1_GOEnrich = rownames(clust1_genes)

#create gene list for input in topGO for cluster 1
geneList_clust1 = factor(as.integer(geneUniverse %in% clust1_GOEnrich))
names(geneList_clust1) = geneUniverse
str(geneList_clust1)

##create a topGO object cluster1
GOdata_BP_clust1 = new("topGOdata", ontology="BP", allGenes=geneList_clust1, 
                           annot = annFUN.gene2GO, gene2GO = geneID2GO)
GOdata_MF_clust1 = new("topGOdata", ontology="MF", allGenes=geneList_clust1, 
                           annot = annFUN.gene2GO, gene2GO = geneID2GO)
GOdata_CC_clust1 = new("topGOdata", ontology="CC", allGenes=geneList_clust1, 
                           annot = annFUN.gene2GO, gene2GO = geneID2GO)

#run Fisher's exact test
resultFisher_BP_clust1_elim <- runTest(GOdata_BP_clust1, algorithm = "elim", statistic = "fisher")
resultFisher_MF_clust1_elim <- runTest(GOdata_MF_clust1, algorithm = "elim", statistic = "fisher")
resultFisher_CC_clust1_elim <- runTest(GOdata_CC_clust1, algorithm = "elim", statistic = "fisher")

#extract the significant GO terms
clust1_BP_elim <- GenTable(GOdata_BP_clust1, classic = resultFisher_BP_clust1_elim, 
                                        orderBy = "weight", ranksOf = "weight", topNodes = 50)
clust1_MF_elim <- GenTable(GOdata_MF_clust1, classic = resultFisher_MF_clust1_elim, 
                                        orderBy = "weight", ranksOf = "weight", topNodes = 50)
clust1_CC_elim <- GenTable(GOdata_CC_clust1, classic = resultFisher_CC_clust1_elim, 
                                        orderBy = "weight", ranksOf = "weight", topNodes = 50)

#files for REVIGO input
#write.table(clust1_BP_elim,
#            "/My Drive/ShortTermStress-IlluminaData/FullExp/Cryp-DE/data/ReviGo_files/Clusters/clust1_BP_elim.txt")
#write.table(clust1_MF_elim,
#            "/My Drive/ShortTermStress-IlluminaData/FullExp/Cryp-DE/data/ReviGo_files/Clusters/clust1_MF_elim.txt")
#write.table(clust1_CC_elim,
#            "/My Drive/ShortTermStress-IlluminaData/FullExp/Cryp-DE/data/ReviGo_files/Clusters/clust1_CC_elim.txt")
```

Cluster 2

```{r}
clust2_genes = read.delim("/My Drive/ShortTermStress-IlluminaData/FullExp/Cryp-DE/data/Clusters_11-19/sign2LFC_Holm_C2.txt",header = TRUE,row.names = 1)
clust2_genes = clust2_genes[-1, ]

clust2_GOEnrich = rownames(clust2_genes)

#create gene list for input in topGO for cluster 1
geneList_clust2 = factor(as.integer(geneUniverse %in% clust2_GOEnrich))
names(geneList_clust2) = geneUniverse


##create a topGO object cluster1
GOdata_BP_clust2 = new("topGOdata", ontology="BP", allGenes=geneList_clust2, 
                           annot = annFUN.gene2GO, gene2GO = geneID2GO)
GOdata_MF_clust2 = new("topGOdata", ontology="MF", allGenes=geneList_clust2, 
                           annot = annFUN.gene2GO, gene2GO = geneID2GO)
GOdata_CC_clust2 = new("topGOdata", ontology="CC", allGenes=geneList_clust2, 
                           annot = annFUN.gene2GO, gene2GO = geneID2GO)

#run Fisher's exact test
resultFisher_BP_clust2_elim <- runTest(GOdata_BP_clust2, algorithm = "elim", statistic = "fisher")
resultFisher_MF_clust2_elim <- runTest(GOdata_MF_clust2, algorithm = "elim", statistic = "fisher")
resultFisher_CC_clust2_elim <- runTest(GOdata_CC_clust2, algorithm = "elim", statistic = "fisher")


#extract the significant GO terms
clust2_BP_elim <- GenTable(GOdata_BP_clust2, classic = resultFisher_BP_clust2_elim, 
                                        orderBy = "weight", ranksOf = "weight", topNodes = 50)
clust2_MF_elim <- GenTable(GOdata_MF_clust2, classic = resultFisher_MF_clust2_elim, 
                                        orderBy = "weight", ranksOf = "weight", topNodes = 50)
clust2_CC_elim <- GenTable(GOdata_CC_clust2, classic = resultFisher_CC_clust2_elim, 
                                        orderBy = "weight", ranksOf = "weight", topNodes = 50)


#files for REVIGO input2.
#write.table(clust2_BP_elim,
#            "/My Drive/ShortTermStress-IlluminaData/FullExp/Cryp-DE/data/ReviGo_files/Clusters/clust2_BP_elim.txt")
#write.table(clust2_MF_elim,
#            "/My Drive/ShortTermStress-IlluminaData/FullExp/Cryp-DE/data/ReviGo_files/Clusters/clust2_MF_elim.txt")
#write.table(clust2_CC_elim,
#            "/My Drive/ShortTermStress-IlluminaData/FullExp/Cryp-DE/data/ReviGo_files/Clusters/clust2_CC_elim.txt")
```

Cluster 3

```{r}
clust3_genes = read.delim("/My Drive/ShortTermStress-IlluminaData/FullExp/Cryp-DE/data/Clusters_11-19/sign2LFC_Holm_C3.txt",header = TRUE,row.names = 1)
clust3_genes = clust3_genes[-1, ]

clust3_GOEnrich = rownames(clust3_genes)

#create gene list for input in topGO for cluster 1
geneList_clust3 = factor(as.integer(geneUniverse %in% clust3_GOEnrich))
names(geneList_clust3) = geneUniverse

##create a topGO object cluster1
GOdata_BP_clust3 = new("topGOdata", ontology="BP", allGenes=geneList_clust3, 
                           annot = annFUN.gene2GO, gene2GO = geneID2GO)
GOdata_MF_clust3 = new("topGOdata", ontology="MF", allGenes=geneList_clust3, 
                           annot = annFUN.gene2GO, gene2GO = geneID2GO)
GOdata_CC_clust3 = new("topGOdata", ontology="CC", allGenes=geneList_clust3, 
                           annot = annFUN.gene2GO, gene2GO = geneID2GO)

#run Fisher's exact test
resultFisher_BP_clust3_elim <- runTest(GOdata_BP_clust3, algorithm = "elim", statistic = "fisher")
resultFisher_MF_clust3_elim <- runTest(GOdata_MF_clust3, algorithm = "elim", statistic = "fisher")
resultFisher_CC_clust3_elim <- runTest(GOdata_CC_clust3, algorithm = "elim", statistic = "fisher")


#extract the significant GO terms
clust3_BP_elim <- GenTable(GOdata_BP_clust3, classic = resultFisher_BP_clust3_elim, 
                                        orderBy = "weight", ranksOf = "weight", topNodes = 50)
clust3_MF_elim <- GenTable(GOdata_MF_clust3, classic = resultFisher_MF_clust3_elim, 
                                        orderBy = "weight", ranksOf = "weight", topNodes = 50)
clust3_CC_elim <- GenTable(GOdata_CC_clust3, classic = resultFisher_CC_clust3_elim, 
                                        orderBy = "weight", ranksOf = "weight", topNodes = 50)

#files for REVIGO input2.
#write.table(clust3_BP_elim,
#            "/My Drive/ShortTermStress-IlluminaData/FullExp/Cryp-DE/data/ReviGo_files/Clusters/clust3_BP_elim.txt")
#write.table(clust3_MF_elim,
#            "/My Drive/ShortTermStress-IlluminaData/FullExp/Cryp-DE/data/ReviGo_files/Clusters/clust3_MF_elim.txt")
#write.table(clust3_CC_elim,
#            "/My Drive/ShortTermStress-IlluminaData/FullExp/Cryp-DE/data/ReviGo_files/Clusters/clust3_CC_elim.txt")
```

Cluster 4

```{r}
clust4_genes = read.delim("/My Drive/ShortTermStress-IlluminaData/FullExp/Cryp-DE/data/Clusters_11-19/sign2LFC_Holm_C4.txt",header = TRUE,row.names = 1)
clust4_genes = clust4_genes[-1, ]

clust4_GOEnrich = rownames(clust4_genes)

#create gene list for input in topGO for cluster 1
geneList_clust4 = factor(as.integer(geneUniverse %in% clust4_GOEnrich))
names(geneList_clust4) = geneUniverse

##create a topGO object cluster1
GOdata_BP_clust4 = new("topGOdata", ontology="BP", allGenes=geneList_clust4, 
                           annot = annFUN.gene2GO, gene2GO = geneID2GO)
GOdata_MF_clust4 = new("topGOdata", ontology="MF", allGenes=geneList_clust4, 
                           annot = annFUN.gene2GO, gene2GO = geneID2GO)
GOdata_CC_clust4 = new("topGOdata", ontology="CC", allGenes=geneList_clust4, 
                           annot = annFUN.gene2GO, gene2GO = geneID2GO)

#run Fisher's exact test
resultFisher_BP_clust4_elim <- runTest(GOdata_BP_clust4, algorithm = "elim", statistic = "fisher")
resultFisher_MF_clust4_elim <- runTest(GOdata_MF_clust4, algorithm = "elim", statistic = "fisher")
resultFisher_CC_clust4_elim <- runTest(GOdata_CC_clust4, algorithm = "elim", statistic = "fisher")


#extract the significant GO terms
clust4_BP_elim <- GenTable(GOdata_BP_clust4, classic = resultFisher_BP_clust4_elim, 
                                        orderBy = "weight", ranksOf = "weight", topNodes = 50)
clust4_MF_elim <- GenTable(GOdata_MF_clust4, classic = resultFisher_MF_clust4_elim, 
                                        orderBy = "weight", ranksOf = "weight", topNodes = 50)
clust4_CC_elim <- GenTable(GOdata_CC_clust4, classic = resultFisher_CC_clust4_elim, 
                                        orderBy = "weight", ranksOf = "weight", topNodes = 50)

#files for REVIGO input2.
#write.table(clust4_BP_elim,
#            "/My Drive/ShortTermStress-IlluminaData/FullExp/Cryp-DE/data/ReviGo_files/Clusters/clust4_BP_elim.txt")
#write.table(clust4_MF_elim,
#            "/My Drive/ShortTermStress-IlluminaData/FullExp/Cryp-DE/data/ReviGo_files/Clusters/clust4_MF_elim.txt")
#write.table(clust4_CC_elim,
#            "/My Drive/ShortTermStress-IlluminaData/FullExp/Cryp-DE/data/ReviGo_files/Clusters/clust4_CC_elim.txt")
```

Cluster 5

```{r}
clust5_genes = read.delim("/My Drive/ShortTermStress-IlluminaData/FullExp/Cryp-DE/data/Clusters_11-19/sign2LFC_Holm_C5.txt",header = TRUE,row.names = 1)
clust5_genes = clust5_genes[-1, ]

clust5_GOEnrich = rownames(clust5_genes)

#create gene list for input in topGO for cluster 1
geneList_clust5 = factor(as.integer(geneUniverse %in% clust5_GOEnrich))
names(geneList_clust5) = geneUniverse

##create a topGO object cluster1
GOdata_BP_clust5 = new("topGOdata", ontology="BP", allGenes=geneList_clust5, 
                           annot = annFUN.gene2GO, gene2GO = geneID2GO)
GOdata_MF_clust5 = new("topGOdata", ontology="MF", allGenes=geneList_clust5, 
                           annot = annFUN.gene2GO, gene2GO = geneID2GO)
GOdata_CC_clust5 = new("topGOdata", ontology="CC", allGenes=geneList_clust5, 
                           annot = annFUN.gene2GO, gene2GO = geneID2GO)

#run Fisher's exact test
resultFisher_BP_clust5_elim <- runTest(GOdata_BP_clust5, algorithm = "elim", statistic = "fisher")
resultFisher_MF_clust5_elim <- runTest(GOdata_MF_clust5, algorithm = "elim", statistic = "fisher")
resultFisher_CC_clust5_elim <- runTest(GOdata_CC_clust5, algorithm = "elim", statistic = "fisher")


#extract the significant GO terms
clust5_BP_elim <- GenTable(GOdata_BP_clust5, classic = resultFisher_BP_clust5_elim, 
                                        orderBy = "weight", ranksOf = "weight", topNodes = 50)
clust5_MF_elim <- GenTable(GOdata_MF_clust5, classic = resultFisher_MF_clust5_elim, 
                                        orderBy = "weight", ranksOf = "weight", topNodes = 50)
clust5_CC_elim <- GenTable(GOdata_CC_clust5, classic = resultFisher_CC_clust5_elim, 
                                        orderBy = "weight", ranksOf = "weight", topNodes = 50)

#files for REVIGO input.
#write.table(clust5_BP_elim,
#            "/My Drive/ShortTermStress-IlluminaData/FullExp/Cryp-DE/data/ReviGo_files/Clusters/clust5_BP_elim.txt")
#write.table(clust5_MF_elim,
#            "/My Drive/ShortTermStress-IlluminaData/FullExp/Cryp-DE/data/ReviGo_files/Clusters/clust5_MF_elim.txt")
#write.table(clust5_CC_elim,
#            "/My Drive/ShortTermStress-IlluminaData/FullExp/Cryp-DE/data/ReviGo_files/Clusters/clust5_CC_elim.txt")
```

Cluster 6

```{r}
clust6_genes = read.delim("/My Drive/ShortTermStress-IlluminaData/FullExp/Cryp-DE/data/Clusters_11-19/sign2LFC_Holm_C6.txt",header = TRUE,row.names = 1)
clust6_genes = clust6_genes[-1, ]

clust6_GOEnrich = rownames(clust6_genes)

#create gene list for input in topGO for cluster 1
geneList_clust6 = factor(as.integer(geneUniverse %in% clust6_GOEnrich))
names(geneList_clust6) = geneUniverse

##create a topGO object cluster1
GOdata_BP_clust6 = new("topGOdata", ontology="BP", allGenes=geneList_clust6, 
                           annot = annFUN.gene2GO, gene2GO = geneID2GO)
GOdata_MF_clust6 = new("topGOdata", ontology="MF", allGenes=geneList_clust6, 
                           annot = annFUN.gene2GO, gene2GO = geneID2GO)
GOdata_CC_clust6 = new("topGOdata", ontology="CC", allGenes=geneList_clust6, 
                           annot = annFUN.gene2GO, gene2GO = geneID2GO)

#run Fisher's exact test
resultFisher_BP_clust6_elim <- runTest(GOdata_BP_clust6, algorithm = "elim", statistic = "fisher")
resultFisher_MF_clust6_elim <- runTest(GOdata_MF_clust6, algorithm = "elim", statistic = "fisher")
resultFisher_CC_clust6_elim <- runTest(GOdata_CC_clust6, algorithm = "elim", statistic = "fisher")


#extract the significant GO terms
clust6_BP_elim <- GenTable(GOdata_BP_clust6, classic = resultFisher_BP_clust6_elim, 
                                        orderBy = "weight", ranksOf = "weight", topNodes = 50)
clust6_MF_elim <- GenTable(GOdata_MF_clust6, classic = resultFisher_MF_clust6_elim, 
                                        orderBy = "weight", ranksOf = "weight", topNodes = 50)
clust6_CC_elim <- GenTable(GOdata_CC_clust6, classic = resultFisher_CC_clust6_elim, 
                                        orderBy = "weight", ranksOf = "weight", topNodes = 50)

#files for REVIGO input2.
#write.table(clust6_BP_elim,
#            "/My Drive/ShortTermStress-IlluminaData/FullExp/Cryp-DE/data/ReviGo_files/Clusters/clust6_BP_elim.txt")
#write.table(clust6_MF_elim,
#            "/My Drive/ShortTermStress-IlluminaData/FullExp/Cryp-DE/data/ReviGo_files/Clusters/clust6_MF_elim.txt")
#write.table(clust6_CC_elim,
#            "/My Drive/ShortTermStress-IlluminaData/FullExp/Cryp-DE/data/ReviGo_files/Clusters/clust6_CC_elim.txt")
```

Cluster 7

```{r}
clust7_genes = read.delim("/My Drive/ShortTermStress-IlluminaData/FullExp/Cryp-DE/data/Clusters_11-19/sign2LFC_Holm_C7.txt",header = TRUE,row.names = 1)
clust7_genes = clust7_genes[-1, ]

clust7_GOEnrich = rownames(clust7_genes)

#create gene list for input in topGO for cluster 1
geneList_clust7 = factor(as.integer(geneUniverse %in% clust7_GOEnrich))
names(geneList_clust7) = geneUniverse

##create a topGO object cluster1
GOdata_BP_clust7 = new("topGOdata", ontology="BP", allGenes=geneList_clust7, 
                           annot = annFUN.gene2GO, gene2GO = geneID2GO)
GOdata_MF_clust7 = new("topGOdata", ontology="MF", allGenes=geneList_clust7, 
                           annot = annFUN.gene2GO, gene2GO = geneID2GO)
GOdata_CC_clust7 = new("topGOdata", ontology="CC", allGenes=geneList_clust7, 
                           annot = annFUN.gene2GO, gene2GO = geneID2GO)

#run Fisher's exact test
resultFisher_BP_clust7_elim <- runTest(GOdata_BP_clust7, algorithm = "elim", statistic = "fisher")
resultFisher_MF_clust7_elim <- runTest(GOdata_MF_clust7, algorithm = "elim", statistic = "fisher")
resultFisher_CC_clust7_elim <- runTest(GOdata_CC_clust7, algorithm = "elim", statistic = "fisher")


#extract the significant GO terms
clust7_BP_elim <- GenTable(GOdata_BP_clust7, classic = resultFisher_BP_clust7_elim, 
                                        orderBy = "weight", ranksOf = "weight", topNodes = 50)
clust7_MF_elim <- GenTable(GOdata_MF_clust7, classic = resultFisher_MF_clust7_elim, 
                                        orderBy = "weight", ranksOf = "weight", topNodes = 50)
clust7_CC_elim <- GenTable(GOdata_CC_clust7, classic = resultFisher_CC_clust7_elim, 
                                        orderBy = "weight", ranksOf = "weight", topNodes = 50)

#files for REVIGO input.
#write.table(clust7_BP_elim,
#            "/My Drive/ShortTermStress-IlluminaData/FullExp/Cryp-DE/data/ReviGo_files/Clusters/clust7_BP_elim.txt")
#write.table(clust7_MF_elim,
#            "/My Drive/ShortTermStress-IlluminaData/FullExp/Cryp-DE/data/ReviGo_files/Clusters/clust7_MF_elim.txt")
#write.table(clust7_CC_elim,
#            "/My Drive/ShortTermStress-IlluminaData/FullExp/Cryp-DE/data/ReviGo_files/Clusters/clust7_CC_elim.txt")
```

Cluster 8

```{r}
clust8_genes = read.delim("/My Drive/ShortTermStress-IlluminaData/FullExp/Cryp-DE/data/Clusters_11-19/sign2LFC_Holm_C8.txt",header = TRUE,row.names = 1)
clust8_genes = clust8_genes[-1, ]

clust8_GOEnrich = rownames(clust8_genes)

#create gene list for input in topGO for cluster 1
geneList_clust8 = factor(as.integer(geneUniverse %in% clust8_GOEnrich))
names(geneList_clust8) = geneUniverse

##create a topGO object cluster1
GOdata_BP_clust8 = new("topGOdata", ontology="BP", allGenes=geneList_clust8, 
                           annot = annFUN.gene2GO, gene2GO = geneID2GO)
GOdata_MF_clust8 = new("topGOdata", ontology="MF", allGenes=geneList_clust8, 
                           annot = annFUN.gene2GO, gene2GO = geneID2GO)
GOdata_CC_clust8 = new("topGOdata", ontology="CC", allGenes=geneList_clust8, 
                           annot = annFUN.gene2GO, gene2GO = geneID2GO)

#run Fisher's exact test
resultFisher_BP_clust8_elim <- runTest(GOdata_BP_clust8, algorithm = "elim", statistic = "fisher")
resultFisher_MF_clust8_elim <- runTest(GOdata_MF_clust8, algorithm = "elim", statistic = "fisher")
resultFisher_CC_clust8_elim <- runTest(GOdata_CC_clust8, algorithm = "elim", statistic = "fisher")


#extract the significant GO terms
clust8_BP_elim <- GenTable(GOdata_BP_clust8, classic = resultFisher_BP_clust8_elim, 
                                        orderBy = "weight", ranksOf = "weight", topNodes = 50)
clust8_MF_elim <- GenTable(GOdata_MF_clust8, classic = resultFisher_MF_clust8_elim, 
                                        orderBy = "weight", ranksOf = "weight", topNodes = 50)
clust8_CC_elim <- GenTable(GOdata_CC_clust8, classic = resultFisher_CC_clust8_elim, 
                                        orderBy = "weight", ranksOf = "weight", topNodes = 50)

#files for REVIGO input.
#write.table(clust8_BP_elim,
#            "/My Drive/ShortTermStress-IlluminaData/FullExp/Cryp-DE/data/ReviGo_files/Clusters/clust8_BP_elim.txt")
#write.table(clust8_MF_elim,
#            "/My Drive/ShortTermStress-IlluminaData/FullExp/Cryp-DE/data/ReviGo_files/Clusters/clust8_MF_elim.txt")
#write.table(clust8_CC_elim,
#            "/My Drive/ShortTermStress-IlluminaData/FullExp/Cryp-DE/data/ReviGo_files/Clusters/clust8_CC_elim.txt")
```

Cluster 9

```{r}
clust9_genes = read.delim("/My Drive/ShortTermStress-IlluminaData/FullExp/Cryp-DE/data/Clusters_11-19/sign2LFC_Holm_C9.txt",header = TRUE,row.names = 1)
clust9_genes = clust9_genes[-1, ]

clust9_GOEnrich = rownames(clust9_genes)

#create gene list for input in topGO for cluster 1
geneList_clust9 = factor(as.integer(geneUniverse %in% clust9_GOEnrich))
names(geneList_clust9) = geneUniverse

##create a topGO object cluster1
GOdata_BP_clust9 = new("topGOdata", ontology="BP", allGenes=geneList_clust9, 
                           annot = annFUN.gene2GO, gene2GO = geneID2GO)
GOdata_MF_clust9 = new("topGOdata", ontology="MF", allGenes=geneList_clust9, 
                           annot = annFUN.gene2GO, gene2GO = geneID2GO)
GOdata_CC_clust9 = new("topGOdata", ontology="CC", allGenes=geneList_clust9, 
                           annot = annFUN.gene2GO, gene2GO = geneID2GO)

#run Fisher's exact test
resultFisher_BP_clust9_elim <- runTest(GOdata_BP_clust9, algorithm = "elim", statistic = "fisher")
resultFisher_MF_clust9_elim <- runTest(GOdata_MF_clust9, algorithm = "elim", statistic = "fisher")
resultFisher_CC_clust9_elim <- runTest(GOdata_CC_clust9, algorithm = "elim", statistic = "fisher")


#extract the significant GO terms
clust9_BP_elim <- GenTable(GOdata_BP_clust9, classic = resultFisher_BP_clust9_elim, 
                                        orderBy = "weight", ranksOf = "weight", topNodes = 50)
clust9_MF_elim <- GenTable(GOdata_MF_clust9, classic = resultFisher_MF_clust9_elim, 
                                        orderBy = "weight", ranksOf = "weight", topNodes = 50)
clust9_CC_elim <- GenTable(GOdata_CC_clust9, classic = resultFisher_CC_clust9_elim, 
                                        orderBy = "weight", ranksOf = "weight", topNodes = 50)

#files for REVIGO input2.
#write.table(clust9_BP_elim,
#            "/My Drive/ShortTermStress-IlluminaData/FullExp/Cryp-DE/data/ReviGo_files/Clusters/clust9_BP_elim.txt")
#write.table(clust9_MF_elim,
#            "/My Drive/ShortTermStress-IlluminaData/FullExp/Cryp-DE/data/ReviGo_files/Clusters/clust9_MF_elim.txt")
#write.table(clust9_CC_elim,
#            "/My Drive/ShortTermStress-IlluminaData/FullExp/Cryp-DE/data/ReviGo_files/Clusters/clust9_CC_elim.txt")
```

Cluster 10

```{r}
clust10_genes = read.delim("/My Drive/ShortTermStress-IlluminaData/FullExp/Cryp-DE/data/Clusters_11-19/sign2LFC_Holm_C10.txt",header = TRUE,row.names = 1)
clust10_genes = clust10_genes[-1, ]

clust10_GOEnrich = rownames(clust10_genes)

#create gene list for input in topGO for cluster 1
geneList_clust10 = factor(as.integer(geneUniverse %in% clust10_GOEnrich))
names(geneList_clust10) = geneUniverse

##create a topGO object cluster1
GOdata_BP_clust10 = new("topGOdata", ontology="BP", allGenes=geneList_clust10, 
                           annot = annFUN.gene2GO, gene2GO = geneID2GO)
GOdata_MF_clust10 = new("topGOdata", ontology="MF", allGenes=geneList_clust10, 
                           annot = annFUN.gene2GO, gene2GO = geneID2GO)
GOdata_CC_clust10 = new("topGOdata", ontology="CC", allGenes=geneList_clust10, 
                           annot = annFUN.gene2GO, gene2GO = geneID2GO)

#run Fisher's exact test
resultFisher_BP_clust10_elim <- runTest(GOdata_BP_clust10, algorithm = "elim", statistic = "fisher")
resultFisher_MF_clust10_elim <- runTest(GOdata_MF_clust10, algorithm = "elim", statistic = "fisher")
resultFisher_CC_clust10_elim <- runTest(GOdata_CC_clust10, algorithm = "elim", statistic = "fisher")


#extract the significant GO terms
clust10_BP_elim <- GenTable(GOdata_BP_clust10, classic = resultFisher_BP_clust10_elim, 
                                        orderBy = "weight", ranksOf = "weight", topNodes = 50)
clust10_MF_elim <- GenTable(GOdata_MF_clust10, classic = resultFisher_MF_clust10_elim, 
                                        orderBy = "weight", ranksOf = "weight", topNodes = 50)
clust10_CC_elim <- GenTable(GOdata_CC_clust10, classic = resultFisher_CC_clust10_elim, 
                                        orderBy = "weight", ranksOf = "weight", topNodes = 50)

#files for REVIGO input
#write.table(clust10_BP_elim,
#            "/My Drive/ShortTermStress-IlluminaData/FullExp/Cryp-DE/data/ReviGo_files/Clusters/clust10_BP_elim.txt")
#write.table(clust10_MF_elim,
#            "/My Drive/ShortTermStress-IlluminaData/FullExp/Cryp-DE/data/ReviGo_files/Clusters/clust10_MF_elim.txt")
#write.table(clust10_CC_elim,
#            "/My Drive/ShortTermStress-IlluminaData/FullExp/Cryp-DE/data/ReviGo_files/Clusters/clust10_CC_elim.txt")
```

# 5. Combine data for KO, GO, uniprot, and cluster information with the LFC data for perusal. 

Here we create a couple metadata tables to reference for a wholistic view of the data

```{r, message=FALSE, warning=FALSE, results='hide'}
kofamKO <- read.delim("/My Drive/ShortTermStress-IlluminaData/FullExp/Cryp-DE/CCMP332_KOtoMerge.txt", header = TRUE, sep = '\t', fill = TRUE, row.names=NULL)
colnames(kofamKO) <- c("gene_id","KO.ID","E_value","KO_definition")
norepeats_KO <- kofamKO %>% distinct(gene_id, .keep_all = TRUE) #Keep only the unique gene_id rows. Kept the first occurrence, which worked out in all the instances I looked at 

GO <- read.delim("/My Drive/ShortTermStress-IlluminaData/FullExp/Cryp-DE/CCMP332_GFF_GO.txt", header = TRUE, sep = '\t', fill = TRUE, row.names=NULL)
colnames(GO) <- c("gene_id","GO.ID")

uniprot_data <- read.delim("/My Drive/ShortTermStress-IlluminaData/FullExp/Cryp-DE/CCMP332_uniprot.csv", header = TRUE, sep = '\t', fill = TRUE, row.names=NULL)
rownames(uniprot_data) <- uniprot_data[,1]
uniprot_data<-  subset(uniprot_data, select=-c(Entry,Gene.names,Organism,KEGG,KO,Gene.ontology.IDs))
colnames(uniprot_data) <- c("protein_name","molecular_function","cellular_component","biological_process")

dmndDB <- read.delim("/My Drive/ShortTermStress-IlluminaData/FullExp/Cryp-DE/CCMP332-diamondbp-matches.m8", header = TRUE, sep = '\t', fill = TRUE)
dmndDB <-  subset(dmndDB, select=-c(pident,length,mismatch,gapopen,qstart,qend,sstart,send))
colnames(dmndDB) <- c("gene_id","UniprotID","E.value","bitscore")

gene_uniprot_anno <- uniprot_data %>%
  tibble::rownames_to_column(.,"UniprotID") %>% #Use full_join to make sure we keep all the expression data and don't limit the GO and KO terms
  full_join(., dmndDB, by = NULL, copy = FALSE, suffix = c(".x",".y"))
gene_uniprot_anno <- gene_uniprot_anno %>% distinct(gene_id, .keep_all = TRUE) #Keep only the unique gene_id rows. Kept the first occurrence, which worked out in all the instances I looked at 

DE_LT <- tibble::rownames_to_column(DE_nakov$table, "gene_id")

paralogs <- read.delim("/My Drive/ShortTermStress-IlluminaData/FullExp/Cryp-DE/data/Cryp_ortho/Results_Feb06/Ortho.txt", header =FALSE, sep = '\t', fill = TRUE, row.names = NULL) #There
#paralogs <- subset(paralogs, select=-c(C_nana))
colnames(paralogs) <- c("orthogroup","gene_id")
paralogs$gene_id <- str_split(paralogs$gene_id, ",") # separate desired column on ,
paralogs <- paralogs %>% unnest(cols = gene_id) # unnest list column to rows
paralogs$gene_id <- paralogs$gene_id %>% str_extract("CCRYP_\\d+")
paralogs <- (na.omit(paralogs))
paralogs_multi <- paralogs %>% distinct(gene_id, .keep_all = TRUE)  %>% group_by(orthogroup) %>% filter( n() > 2 )
paralogs_pairs <- paralogs %>% distinct(gene_id, .keep_all = TRUE)  %>% group_by(orthogroup) %>% filter( n() == 2 )
singletons <- paralogs %>% distinct(gene_id, .keep_all = TRUE)  %>% group_by(orthogroup) %>% filter( n() == 1 )
singletons$cnt_OG <- "1" 
paralogs_pairs$cnt_OG <- "2" 
paralogs_multi$cnt_OG <- "3+" 

clusters <- read.delim("/My Drive/ShortTermStress-IlluminaData/FullExp/Cryp-DE/data/Clusters_11-19/10clusters.txt", header = TRUE, sep = '\t', fill = TRUE)

clusters_GO_KO_2sigdf <- sign2LFC %>% 
  tibble::rownames_to_column(.,"gene_id") %>% #Use full_join to make sure we keep all the expression data and don't limit the GO and KO terms
  left_join(., clusters, by = NULL, copy = FALSE, suffix = c(".x",".y")) %>%
  left_join(., GO, by = NULL, copy = FALSE, suffix = c(".x",".y")) %>%
  left_join(., norepeats_KO, by = NULL, copy = FALSE, suffix = c(".x",".y")) %>%
  left_join(., gene_uniprot_anno, by = NULL, copy = FALSE, suffix = c(".x",".y"))  %>%
  left_join(., DE_LT, by = NULL, copy = FALSE, suffix = c(".x",".y"))
colnames(clusters_GO_KO_2sigdf) <- c("gene_id","t15m","t15m_Padj","t30m","t30m_Padj","t1h","t1h_Padj","t2h","t2h_Padj",
                                    "t4h","t4h_Padj","t8h","t8h_Padj","t10h","t10h_Padj","cluster","GO.ID",
                                    "KO.ID","E-value","KO_definition","UniprotID","protein_name",
                                    "MF","CC","BP","E-value","Bitscore","LT_logFC","LT_CPM","LT_PValue")

clusters_GO_KO_2Sig <- subset(clusters_GO_KO_2sigdf, select=c("gene_id","t15m","t30m","t1h","t2h","t4h","t8h","t10h","LT_logFC","LT_PValue",
                                                  "cluster","UniprotID","protein_name","KO.ID","KO_definition",
                                                  "GO.ID","MF","BP","CC"))

NS_LFCanno_df <- NS_LFC_df %>% 
  tibble::rownames_to_column(.,"gene_id") %>% #Use full_join to make sure we keep all the expression data and don't limit the GO and KO terms
  left_join(., clusters, by = NULL, copy = FALSE, suffix = c(".x",".y")) %>%
  left_join(., GO, by = NULL, copy = FALSE, suffix = c(".x",".y")) %>%
  left_join(., norepeats_KO, by = NULL, copy = FALSE, suffix = c(".x",".y")) %>%
  left_join(., gene_uniprot_anno, by = NULL, copy = FALSE, suffix = c(".x",".y")) %>%
  left_join(., DE_LT, by = NULL, copy = FALSE, suffix = c(".x",".y"))
colnames(NS_LFCanno_df) <- c("gene_id","t15m","t15m_Padj","t30m","t30m_Padj","t1h","t1h_Padj","t2h","t2h_Padj",
                                    "t4h","t4h_Padj","t8h","t8h_Padj","t10h","t10h_Padj","cluster", "GO.ID",
                                    "KO.ID","E-value","KO_definition","UniprotID","protein_name",
                                    "MF","CC","BP","E-value","Bitscore","LT_logFC","LT_CPM","LT_PValue")

NS_LFCanno_df$cluster <- "Not2Sig"


NS_LFCanno <- subset(NS_LFCanno_df, select=c("gene_id","t15m","t30m","t1h","t2h","t4h","t8h","t10h","LT_logFC","LT_PValue",
                                                  "cluster","UniprotID","protein_name","KO.ID","KO_definition",
                                                  "GO.ID","MF","BP","CC"))

not_sig_anno_df <- not_sig_genes %>% 
  tibble::rownames_to_column(.,"gene_id") %>% #Use full_join to make sure we keep all the expression data and don't limit the GO and KO terms
  left_join(., GO, by = NULL, copy = FALSE, suffix = c(".x",".y")) %>%
  left_join(., norepeats_KO, by = NULL, copy = FALSE, suffix = c(".x",".y")) %>%
  left_join(., gene_uniprot_anno, by = NULL, copy = FALSE, suffix = c(".x",".y")) %>%
  left_join(., DE_LT, by = NULL, copy = FALSE, suffix = c(".x",".y"))
colnames(not_sig_anno_df) <- c("gene_id","cluster", "GO.ID",
                                    "KO.ID","E-value","KO_definition","UniprotID","protein_name",
                                    "MF","CC","BP","E-value","Bitscore","LT_logFC","LT_CPM","LT_PValue")
not_sig_anno_df$t15m <- "0"
not_sig_anno_df$t30m <- "0"
not_sig_anno_df$t1h <- "0"
not_sig_anno_df$t2h <- "0"
not_sig_anno_df$t4h <- "0"
not_sig_anno_df$t8h <- "0"
not_sig_anno_df$t10h <- "0"
not_sig_anno <- subset(not_sig_anno_df, select=c("gene_id","t15m","t30m","t1h","t2h","t4h","t8h","t10h","LT_logFC","LT_PValue",
                                                  "cluster","UniprotID","protein_name","KO.ID","KO_definition",
                                                  "GO.ID","MF","BP","CC"))

all_anno <- rbind(clusters_GO_KO_2Sig,NS_LFCanno,not_sig_anno)
paralogs <- rbind(paralogs_multi,singletons,paralogs_pairs)

raw_data <- as.data.frame(gene_cnts)

padj <- padj_df %>% tibble::rownames_to_column(.,"gene_id")

raw_genes <- raw_data %>%
  tibble::rownames_to_column(.,"gene_id") %>%
  left_join(., all_anno, by = NULL, copy = FALSE, suffix = c(".x",".y")) %>%
  left_join(., paralogs, by = NULL, copy = FALSE, suffix = c(".x",".y")) %>%
  left_join(., padj, by = NULL, copy = FALSE, suffix = c(".x",".y"))
colnames(raw_genes) <- c("gene_id","0m_R1","15m_R1","30m_R1","60m_R1","2hr_R1","4hr_R1","8hr_R1","10hr_R1",
                                     "0m_R2","15m_R2","30m_R2","60m_R2","2hr_R2","4hr_R2","8hr_R2","10hr_R2",
                                     "0m_R3","15m_R3","30m_R3","60m_R3","2hr_R3","4hr_R3","8hr_R3","10hr_R3",
                                     "t15m","t30m","t1h","t2h","t4h","t8h","t10h","LT_logFC","LT_PValue","cluster","UniprotID",
                                     "protein_name","KO.ID","KO_definition","GO.ID","MF","BP","CC","orthogroup","cnt_OG",
                                     "padjScreen","15m_Padj","30m_Padj","1h_Padj","2h_Padj","4h_Padj","8h_Padj","10h_Padj")
raw_genes_df <- subset(raw_genes, select=c("gene_id","t15m","15m_Padj","t30m","30m_Padj","t1h","1h_Padj","t2h","2h_Padj",
                                        "t4h","4h_Padj","t8h","8h_Padj","t10h","10h_Padj","LT_logFC","LT_PValue","orthogroup","cnt_OG",
                                        "cluster","UniprotID","protein_name","KO.ID","KO_definition", "GO.ID","MF","BP","CC"))
raw_genes <- subset(raw_genes, select=c("gene_id","t15m","t30m","t1h","t2h","t4h","t8h","t10h","LT_logFC","LT_PValue","orthogroup","cnt_OG",
                                        "cluster","UniprotID","protein_name","KO.ID","KO_definition", "GO.ID","MF","BP","CC"))

raw_genes[,c(2:8)] <- sapply(raw_genes[,c(2:8)], as.numeric)
raw_genes[,c(10)] <- sapply(raw_genes[,c(10)], as.character)

clust_paralogs <- raw_genes %>% filter(str_detect(cluster,"clust"))

```

Diatom genomes are not always well-annotated. How many of our genes lack annotation?

```{r}
no_annotation_data <- subset(all_anno, select=-c(1:10)) 
no_annotation_data_df <- filter(no_annotation_data, rowSums(is.na(no_annotation_data)) != ncol(no_annotation_data))
```

# 6. Manuscript graphs 

## 6.1 Gene expression behavior in short- and long-term

Graph of short-term behavior of genes found to be significant in the long-term

```{r fig.height=6, fig.width=10, warning=FALSE, `copy` = TRUE}
#Set up the downregulated variables
dr_LT <- all_anno[,c(1,2,3,4,5,6,7,8,9)] 
colnames(dr_LT) <- c("gene_id","1","2","3","4","5","6","7","8")
dr_LT <- (na.omit(dr_LT))
dr_LT[,c(2:9)] <- sapply(dr_LT[,c(2:9)], as.numeric)
dr_LT <- subset(dr_LT, (`8`) <= -1) 
dr_ST <- pivot_longer(dr_LT[,c(1,2,3,4,5,6,7,8)], -c(gene_id), names_to = "timept")
dr_ST_md = dr_ST %>% group_by(timept) %>% summarize(value = mean(value), ) %>% mutate(gene_id = "repressed")
dr_LT <- pivot_longer(dr_LT[,c(1,2,3,4,5,6,7,8,9)], -c(gene_id), names_to = "timept")
dr_LT_md = dr_LT %>% group_by(timept) %>% summarize(value = mean(value), ) %>% mutate(gene_id = "repressed") 
dr_LT <- filter(dr_LT, timept == "8") #Get just the long-term timepoint for the boxplot

#Set up the upregulated variables
ur_LT <- all_anno[,c(1,2,3,4,5,6,7,8,9)] 
colnames(ur_LT) <- c("gene_id","1","2","3","4","5","6","7","8")
ur_LT <- (na.omit(ur_LT))
ur_LT[,c(2:9)] <- sapply(ur_LT[,c(2:9)], as.numeric)
ur_LT <- subset(ur_LT, (`8`) >= 1) 
ur_ST <- pivot_longer(ur_LT[,c(1,2,3,4,5,6,7,8)], -c(gene_id), names_to = "timept")
ur_ST_md = ur_ST %>% group_by(timept) %>% summarize(sd = sd(value, na.rm=TRUE), value = mean(value), ) %>% mutate(gene_id = "induced")
ur_LT <- pivot_longer(ur_LT[,c(1,2,3,4,5,6,7,8,9)], -c(gene_id), names_to = "timept")
ur_LT_md = ur_LT %>% group_by(timept) %>% summarize(value = mean(value), ) %>% mutate(gene_id = "induced") 
ur_LT <- filter(ur_LT, timept == "8") #Get just the long-term timepoint for the boxplot

#Split out the outliers to jitter them in the plot for visibility
dr_LT_outliers <- dr_LT %>% group_by(timept) %>%
  filter(value > quantile(value, 0.75) + 1.5 * IQR(value) | value < quantile(value, 0.25) - 1.5 * IQR(value))
ur_LT_outliers <- ur_LT %>% group_by(timept) %>%
  filter(value > quantile(value, 0.75) + 1.5 * IQR(value) | value < quantile(value, 0.25) - 1.5 * IQR(value))

ggplot(NULL)+ 
    #geom_line(data = dr_ST, size = 1, alpha=1/50, aes(x = timept, y = value, group = gene_id))  +
    geom_line(data = dr_ST_md, size=2, alpha = 1,  color = "#a9acff",aes(x = timept, y = value, group = gene_id)) +
    #geom_line(data = ur_ST, size = 1, alpha=1/50, aes(x = timept, y = value, group = gene_id)) +
    geom_line(data = ur_ST_md, size=2, alpha = 1,  color = "#28db69",  aes(x = timept, y = value, group = gene_id)) +
    geom_boxplot(data = dr_LT, color = "black", fill = "#28db69", notch = TRUE, aes(x = timept, y = value), outlier.shape = NA) +
    geom_boxplot(data = ur_LT, color = "black", fill = "#a9acff", notch = TRUE, aes(x = timept, y = value), outlier.shape = NA) +
    geom_jitter(aes(x = timept, y = value), height = 0, width = 0.1, data = ur_LT_outliers) +
    geom_jitter(aes(x = timept, y = value), height = 0, width = 0.1, data = dr_LT_outliers) +
    geom_hline(yintercept = 0, linetype = "dashed") +
    theme_classic() +
    theme(axis.ticks.x=element_blank(), 
          axis.text=element_text(size=12), 
          axis.title=element_text(size=14,face="bold")) +
    xlab("Time exposed to 0 ppt ASW") + 
    ylab("Log2 fold change") +
    ggtitle("Median expression over time for genes induced/repressed at 4 months") +
    theme(plot.title = element_text(hjust = 0.5, face = "bold", size = 16)) 

```

## 6.2 Graph of median expression of genes DE at 1h and 10 h

```{r fig.height=6, fig.width=10, warning=FALSE, `copy` = TRUE}
min_1h <- filter(logFCdown_0v1h , which.pmin(logFCdown_0v1h) == 5)
min_1h <- tibble::rownames_to_column(min_1h, var = "gene_id")
min_1h <- pivot_longer(min_1h[,c(1,2,4,6,8,10,12,14)], -c(gene_id), names_to = "timept")
min_1h$timept <- ordered(min_1h$timept, levels =c("0v15m","0v30m","0v1h","0v2h","0v4h","0v8h","0v10h"))
min_1h_md = min_1h %>% group_by(timept) %>% summarize(sd = sd(value, na.rm=TRUE), value = median(value), ) %>% mutate(gene_id = "induced")
#outlier_min_1h <-  min_1h %>% group_by(timept) %>%
#  filter(value > quantile(value, 0.75) + 1.5 * IQR(value) | value < quantile(value, 0.25) - 1.5 * IQR(value))

max_1h <- filter(logFCup_0v1h , which.pmax(logFCup_0v1h) == 5)
max_1h <- tibble::rownames_to_column(max_1h, var = "gene_id")
max_1h <- pivot_longer(max_1h[,c(1,2,4,6,8,10,12,14)], -c(gene_id), names_to = "timept")
max_1h$timept <- ordered(max_1h$timept, levels =c("0v15m","0v30m","0v1h","0v2h","0v4h","0v8h","0v10h"))
max_1h_md = max_1h %>% group_by(timept) %>% summarize(sd = sd(value, na.rm=TRUE), value = median(value), ) %>% mutate(gene_id = "induced")
#outlier_max_1h <-  max_1h %>% group_by(timept) %>%
#  filter(value > quantile(value, 0.75) + 1.5 * IQR(value) | value < quantile(value, 0.25) - 1.5 * IQR(value))

min_10h <- filter(logFCdown_0v10h, which.pmin(logFCdown_0v10h) == 13)
min_10h <- tibble::rownames_to_column(min_10h, var = "gene_id")
min_10h <- pivot_longer(min_10h[,c(1,2,4,6,8,10,12,14)], -c(gene_id), names_to = "timept")
min_10h$timept <- ordered(min_10h$timept, levels =c("0v15m","0v30m","0v1h","0v2h","0v4h","0v8h","0v10h"))
min_10h_md = min_10h %>% group_by(timept) %>% summarize(sd = sd(value, na.rm=TRUE), value = median(value), ) %>% mutate(gene_id = "repressed")
#outlier_min_10h <-  min_10h %>% group_by(timept) %>%
#  filter(value > quantile(value, 0.75) + 1.5 * IQR(value) | value < quantile(value, 0.25) - 1.5 * IQR(value))

max_10h <- filter(logFCup_0v10h, which.pmax(logFCup_0v10h) == 13)
max_10h <- tibble::rownames_to_column(max_10h, var = "gene_id")
max_10h <- pivot_longer(max_10h[,c(1,2,4,6,8,10,12,14)], -c(gene_id), names_to = "timept")
max_10h$timept <- ordered(max_10h$timept, levels =c("0v15m","0v30m","0v1h","0v2h","0v4h","0v8h","0v10h"))
max_10h_md = max_10h %>% group_by(timept) %>% summarize(sd = sd(value, na.rm=TRUE), value = median(value), ) %>% mutate(gene_id = "induced")
#outlier_max_10h <-  max_10h %>% group_by(timept) %>%
#  filter(value > quantile(value, 0.75) + 1.5 * IQR(value) | value < quantile(value, 0.25) - 1.5 * IQR(value))


ggplot(NULL)+ 
    geom_line(data = min_1h_md, size=2, alpha = 1, color = "#28db69",aes(x = timept, y = value, group = gene_id)) +
    geom_line(data = max_1h_md, size=2, alpha = 1, color = "#a9acff",aes(x = timept, y = value, group = gene_id)) +
    geom_boxplot(data = min_1h, notch = TRUE, aes(x = timept, y = value), fill= "#28db69", width = 0.15, outlier.shape = NA) +
    geom_boxplot(data = max_1h, notch = TRUE, aes(x = timept, y = value), fill= "#a9acff", width = 0.15, outlier.shape = NA) +
    geom_hline(yintercept = 0, linetype = "dashed") +
    ylim(-2.5,2.5) +
    theme_classic() +
    theme(axis.ticks.x=element_blank(), 
          axis.text=element_text(size=12), 
          axis.title=element_text(size=14,face="bold")) +
    xlab("Time exposed to 0 ppt ASW") + 
    ylab("Log2 fold change") +
    ggtitle("Median expression over time for genes induced/repressed at 1h") +
    theme(plot.title = element_text(hjust = 0.5, face = "bold", size = 16))


ggplot(NULL)+ 
    geom_line(data = min_10h_md, size=2, alpha = 1, color = "#28db69",aes(x = timept, y = value, group = gene_id)) +
    geom_line(data = max_10h_md, size=2, alpha = 1, color = "#a9acff",aes(x = timept, y = value, group = gene_id)) +
    geom_boxplot(data = min_10h, notch = TRUE, aes(x = timept, y = value), fill= "#28db69", width = 0.15, outlier.shape = NA) +
    geom_boxplot(data = max_10h, notch = TRUE, aes(x = timept, y = value), fill= "#a9acff", width = 0.15, outlier.shape = NA) +
    geom_hline(yintercept = 0, linetype = "dashed") +
    ylim(-2.5,2.5) +
    theme_classic() +
    theme(axis.ticks.x=element_blank(), 
          axis.text=element_text(size=12), 
          axis.title=element_text(size=14,face="bold")) +
    xlab("Time exposed to 0 ppt ASW") + 
    ylab("Log2 fold change") +
    ggtitle("Median expression over time for genes induced/repressed at 10 h") +
    theme(plot.title = element_text(hjust = 0.5, face = "bold", size = 16))

```

## 6.3 Heatmaps for cycles of interest

Use this code to isolate genes of interest from the metadata tables produced in Section 5. 

```{r, message=FALSE, warning=FALSE, results='hide', fig.width=8, fig.height=10}
colors = c("#28db69","white","purple")
colors=colorRampPalette(colors)(1000)
my_palette <- colorRampPalette(c("#28db69", "white", "#a9acff"))(n = 1000)

Chlorophyll_hmp <- as.matrix(clusters_GO_KO_2Sig %>% arrange(.,KO_definition) %>%
                            filter(str_detect(KO_definition,"EC:6.1.1.17|EC:1.2.1.70|EC:5.4.3.8|EC:4.2.1.24|EC:2.5.1.61|EC:4.2.1.75|EC:4.1.1.37|EC:2.1.1.107|EC:1.3.3.4|EC:1.3.3.3|EC:6.6.1.1|EC:2.1.1.11|EC:1.3.1.75|EC:1.3.1.33|EC:2.5.1.62|EC:2.5.1.133|EC:1.1.1.294"))%>%
                            dplyr::select(`gene_id`,`KO_definition`,`t15m`,`t30m`,`t1h`,`t2h`,`t4h`,`t8h`,`t10h`,`LT_logFC`) %>%
                            unite(.,names,c(`gene_id`,`KO_definition`))
                            %>% remove_rownames %>% column_to_rownames(var="names"))
heatmap.2(Chlorophyll_hmp,dendrogram = "none",trace="none",col=colors,main="Genes in chlorophyll biosynthesis", cexRow = 0.75, cexCol = 1, Colv = FALSE, Rowv = FALSE, offsetRow = -20, density.info="none")

Glycolysis_hmp <- as.matrix(clusters_GO_KO_2Sig %>% arrange(.,KO_definition) %>% 
                            filter(str_detect(KO_definition,"EC:1.2.1.12|EC:2.7.1.90|EC:2.7.1.11|EC:3.1.3.11|EC:4.1.2.13|EC:2.7.2.3|EC:2.7.1.40]|EC:5.3.1.9|EC:4.2.1.11]|EC:5.4.2.11]"))%>%
                            dplyr::select(`gene_id`,`t15m`,`t30m`,`t1h`,`t2h`,`t4h`,`t8h`,`t10h`,`LT_logFC`,`KO_definition`) %>%
                            unite(.,names,c(`gene_id`,`KO_definition`))
                            %>% remove_rownames %>% column_to_rownames(var="names"))

Chitin_hmp <- as.matrix(clusters_GO_KO_2Sig %>% arrange(.,KO_definition) %>% 
                            filter(str_detect(KO_definition,"2.6.1.16|3.5.1.25|5.4.2.3|2.7.7.23|2.7.7.83|2.4.1.16|3.5.2.5|3.2.1.14|3.2.1.52"))%>%
                            dplyr::select(`gene_id`,`KO_definition`,`t15m`,`t30m`,`t1h`,`t2h`,`t4h`,`t8h`,`t10h`,`LT_logFC`) %>%
                            unite(.,names,c(`gene_id`,`KO_definition`))
                            %>% remove_rownames %>% column_to_rownames(var="names"))

CalvinCycle_hmp <- as.matrix(clusters_GO_KO_2Sig %>% arrange(.,KO_definition) %>%
                            filter(str_detect(KO_definition,"EC:2.7.2.3|EC:1.2.1.12|EC:4.1.2.13|EC:3.1.3.11|EC:2.2.1.1|EC:3.1.3.37|EC:5.3.1.6"))%>%
                            dplyr::select(`gene_id`,`KO_definition`,`t15m`,`t30m`,`t1h`,`t2h`,`t4h`,`t8h`,`t10h`,`LT_logFC`) %>%
                            unite(.,names,c(`gene_id`,`KO_definition`))
                            %>% remove_rownames %>% column_to_rownames(var="names"))

Metablism_hmp <- rbind (Glycolysis_hmp,CalvinCycle_hmp,Chitin_hmp)

heatmap.2(CalvinCycle_hmp,dendrogram = "none",trace="none",col=colors,main="Genes in oxidative stress", cexRow = 0.75, cexCol = 1, Colv = FALSE, Rowv = FALSE, offsetRow = 0, density.info="none")


Proline_hmp <- as.matrix(clusters_GO_KO_2Sig  %>% arrange(.,KO_definition) %>%
                            filter(str_detect(KO_definition,"EC:2.7.2.11|EC:1.2.1.41|EC:3.4.11.5|EC:2.6.1.13|EC:1.5.1.2]|EC:4.3.1.12"))%>%
                            dplyr::select(`gene_id`,`KO_definition`,`t15m`,`t30m`,`t1h`,`t2h`,`t4h`,`t8h`,`t10h`,`LT_logFC`) %>%
                            unite(.,names,c(`gene_id`,`KO_definition`))
                            %>% remove_rownames %>% column_to_rownames(var="names"))

Taurine_hmp <- as.matrix(raw_genes  %>% arrange(.,KO_definition) %>%
                            filter(str_detect(protein_name,"L-cysteate sulfo-lyase|glutamate decarboxylase|Gamma-glutamyl cyclotransferase"))%>%
                            dplyr::select(`gene_id`,`KO_definition`,`t15m`,`t30m`,`t1h`,`t2h`,`t4h`,`t8h`,`t10h`,`LT_logFC`) %>%
                            unite(.,names,c(`gene_id`,`KO_definition`))
                            %>% remove_rownames %>% column_to_rownames(var="names"))

Aquaporin_hmp <- as.matrix(raw_genes  %>% arrange(.,KO_definition) %>%
                            filter(str_detect(protein_name,"Aquaporin AqpM"))%>%
                            dplyr::select(`gene_id`,`KO_definition`,`t15m`,`t30m`,`t1h`,`t2h`,`t4h`,`t8h`,`t10h`,`LT_logFC`) %>%
                            unite(.,names,c(`gene_id`,`KO_definition`))
                            %>% remove_rownames %>% column_to_rownames(var="names"))

Osmo_hmp <- rbind(Proline_hmp, Taurine_hmp, Aquaporin_hmp)

heatmap.2(Osmo_hmp,dendrogram = "none",trace="none",col=colors,main="Genes in proline metabolism",  cexRow = 0.75, cexCol = 1, Colv = FALSE, Rowv = FALSE, offsetRow = 0, density.info="none")


HeatShock_hmp_KO <- as.matrix(raw_genes %>% arrange(.,KO_definition) %>%
                            filter(str_detect(KO.ID,"K04077|K04078|K03686|K03687|K04082|K04083|K04079|K13993"))%>%
                            dplyr::select(`gene_id`,`cluster`,`t15m`,`t30m`,`t1h`,`t2h`,`t4h`,`t8h`,`t10h`,`LT_logFC`) %>%
                            unite(.,names,c(`gene_id`,`cluster`)) %>% 
                            remove_rownames %>% column_to_rownames(var="names"))
HeatShock_hmp_UP <- as.matrix(raw_genes %>% arrange(.,protein_name) %>%
                            filter(str_detect(protein_name,"Hsp70|HSP70"))%>%
                            dplyr::select(`gene_id`,`cluster`,`t15m`,`t30m`,`t1h`,`t2h`,`t4h`,`t8h`,`t10h`,`LT_logFC`) %>%
                            unite(.,names,c(`gene_id`,`cluster`)) %>% 
                            remove_rownames %>% column_to_rownames(var="names"))
HeatShock_hmp <- rbind(HeatShock_hmp_KO,HeatShock_hmp_UP)

heatmap.2(HeatShock_hmp,dendrogram = "none",trace="none",col=colors,main="Genes in oxidative stress", cexRow = 0.75, cexCol = 1, Colv = FALSE, Rowv = FALSE, offsetRow = -20, density.info="none")


Sig_Ox <- as.matrix(raw_genes %>% arrange(.,KO_definition) %>%
                            filter(str_detect(gene_id,"CCRYP_015211|CCRYP_020438|CCRYP_013781|CCRYP_016723|CCRYP_002248|CCRYP_002534|CCRYP_010295|CCRYP_016269|CCRYP_002901|CCRYP_003411|CCRYP_020615|CCRYP_008084"))%>%
                            dplyr::select(`gene_id`,`KO_definition`,`t15m`,`t30m`,`t1h`,`t2h`,`t4h`,`t8h`,`t10h`,`LT_logFC`) %>%
                            unite(.,names,c(`gene_id`,`KO_definition`)) %>% 
                            remove_rownames %>% column_to_rownames(var="names"))

heatmap.2(Sig_Ox,dendrogram = "none",trace="none",col=colors,main="Genes in oxidative stress", cexRow = 0.75, cexCol = 1, Colv = FALSE, Rowv = FALSE, offsetRow = -30, density.info="none")
```

## 6.4 Cluster plots

### 6.4.1 Summary of the averages of all the clusters

```{r fig.height=6, fig.width=10, warning=FALSE, copy = TRUE}
cluster_LF10C <- dplyr::filter(clusters_GO_KO_2Sig) 
cluster_LFC10C <- pivot_longer(cluster_LF10C[ ,c(1:8,11)], -c(gene_id, cluster), names_to = "timept")
cluster_LFC10C$timept <- ordered(cluster_LFC10C$timept, levels=c("t15m","t30m","t1h","t2h","t4h","t8h","t10h"))
cluster1 <- dplyr::filter(cluster_LFC10C, (cluster == 'clust1')) 
cluster2 <- dplyr::filter(cluster_LFC10C, (cluster == 'clust2'))
cluster3 <- dplyr::filter(cluster_LFC10C, (cluster == 'clust3'))  
cluster4 <- dplyr::filter(cluster_LFC10C, (cluster == 'clust4')) 
cluster5 <- dplyr::filter(cluster_LFC10C, (cluster == 'clust5')) 
cluster6 <- dplyr::filter(cluster_LFC10C, (cluster == 'clust6'))  
cluster7 <- dplyr::filter(cluster_LFC10C, (cluster == 'clust7'))   
cluster8 <- dplyr::filter(cluster_LFC10C, (cluster == 'clust8'))  
cluster9 <- dplyr::filter(cluster_LFC10C, (cluster == 'clust9')) 
cluster10 <- dplyr::filter(cluster_LFC10C, (cluster == 'clust10'))  

data1 = cluster1 %>% group_by(timept,cluster) %>% summarize(sd = sd(value, na.rm = TRUE),value = mean(value), ) %>% mutate(gene_id = "1")
data2 = cluster2 %>% group_by(timept,cluster) %>% summarize(sd = sd(value, na.rm = TRUE),value = mean(value), ) %>% mutate(gene_id = "2")
data3 = cluster3 %>% group_by(timept,cluster) %>% summarize(sd = sd(value, na.rm = TRUE),value = mean(value), ) %>% mutate(gene_id = "3")
data4 = cluster4 %>% group_by(timept,cluster) %>% summarize(sd = sd(value, na.rm = TRUE),value = mean(value), ) %>% mutate(gene_id = "4")
data5 = cluster5 %>% group_by(timept,cluster) %>% summarize(sd = sd(value, na.rm = TRUE),value = mean(value), ) %>% mutate(gene_id = "5")
data6 = cluster6 %>% group_by(timept,cluster) %>% summarize(sd = sd(value, na.rm = TRUE),value = mean(value), ) %>% mutate(gene_id = "6")
data7 = cluster7 %>% group_by(timept,cluster) %>% summarize(sd = sd(value, na.rm = TRUE),value = mean(value), ) %>% mutate(gene_id = "7")
data8 = cluster8 %>% group_by(timept,cluster) %>% summarize(sd = sd(value, na.rm = TRUE),value = mean(value), ) %>% mutate(gene_id = "8")
data9 = cluster9 %>% group_by(timept,cluster) %>% summarize(sd = sd(value, na.rm = TRUE),value = mean(value), ) %>% mutate(gene_id = "9")
data10 = cluster10 %>% group_by(timept,cluster) %>% summarize(sd = sd(value, na.rm = TRUE),value = mean(value), ) %>% mutate(gene_id = "10")

data = rbind(data1, data2, data3, data4, data5, data6, data7, data8, data9, data10)
data$cluster <- factor(data$cluster, levels=c("clust1","clust2","clust3","clust4","clust5","clust6","clust7","clust8","clust9","clust10"))

#All averages
ggplot(data,  aes(x = timept, y = value, group = cluster, color = cluster)) + 
    geom_line(size = 1.5) +
    ylim(-3.5,3.5) +
    geom_hline(yintercept = 0, linetype = "dashed") +
    theme(axis.text.x = element_text(vjust=115, face = "bold"),
          axis.ticks.x=element_blank()) +
    xlab("Time exposed to 0 ppt ASW") + 
    ylab("Average fold change (log 2)") +
    scale_color_viridis(discrete = TRUE, option = "D") +
    ggtitle("Average expression over time for each cluster") +
    theme(plot.title = element_text(hjust = 0.5, face = "bold"))
```

### 6.4.2 Individual graphs showing cluster averages with error bars

```{r fig.width=10, fig.height=6}
#Individual cluster averages with background data
ggplot(data1, aes(x = timept, y = value, group = gene_id, ymin = value-sd, ymax = value+sd))  + 
    geom_line(color = "#450755", size = 4) +
    geom_errorbar(width = 0.2) +
    geom_hline(yintercept = 0, linetype = "dashed",size=1) +
    ylim(-4,3) +
    theme(axis.title.x = element_blank()) +
    theme(axis.title.y = element_text(face="bold")) +
    theme(legend.position = "none") +
    ylab("Average fold change (log 2)") +
    theme_classic() +
    theme(plot.title = element_text(hjust = 0.5, face = "bold"))

ggplot(data2, aes(x = timept, y = value, group = gene_id, ymin = value-sd, ymax = value+sd))  + 
    geom_line(color = "#48247a", size = 4) +
    geom_errorbar(width = 0.2) +
    geom_hline(yintercept = 0, linetype = "dashed",size=1) +
    ylim(-4,3) +
    theme(axis.title.x = element_text(face="bold")) +
    theme(axis.title.y = element_text(face="bold")) +
    theme(legend.position = "none") +
    ylab("Average fold change (log 2)") +
    theme_classic() +
    theme(plot.title = element_text(hjust = 0.5, face = "bold"))

ggplot(data3, aes(x = timept, y = value, group = gene_id, ymin = value-sd, ymax = value+sd))  + 
    geom_line(color = "#3d488b", size = 4) +
    geom_errorbar(width = 0.2) +
    geom_hline(yintercept = 0, linetype = "dashed",size=1) +
    ylim(-4,3) +
    theme(axis.title.x = element_blank()) +
    theme(axis.title.y = element_text(face="bold")) +
    theme(legend.position = "none") +
    ylab("Average fold change (log 2)") +
    theme_classic() +
    theme(plot.title = element_text(hjust = 0.5, face = "bold"))

ggplot(data4, aes(x = timept, y = value, group = gene_id, ymin = value-sd, ymax = value+sd))  + 
    geom_line(color = "#2d6790", size = 4) +
    geom_errorbar(width = 0.2) +
    geom_hline(yintercept = 0, linetype = "dashed",size=1) +
    ylim(-4,3) +
    theme(axis.title.x = element_blank()) +
    theme(axis.title.y = element_text(face="bold")) +
    theme(legend.position = "none") +
    ylab("Average fold change (log 2)") +
    theme_classic() +
    theme(plot.title = element_text(hjust = 0.5, face = "bold"))

ggplot(data5, aes(x = timept, y = value, group = gene_id, ymin = value-sd, ymax = value+sd))  + 
    geom_line(color = "#1e828f", size = 4) +
    geom_errorbar(width = 0.2) +
    geom_hline(yintercept = 0, linetype = "dashed",size=1) +
    ylim(-4,3) +
    theme(axis.title.x = element_blank()) +
    theme(axis.title.y = element_text(face="bold")) +
    theme(legend.position = "none") +
    ylab("Average fold change (log 2)") +
    theme_classic() +
    theme(plot.title = element_text(hjust = 0.5, face = "bold"))

ggplot(data6, aes(x = timept, y = value, group = gene_id, ymin = value-sd, ymax = value+sd))  + 
    geom_line(color = "#129f89", size = 4) +
    geom_errorbar(width = 0.2) +
    geom_hline(yintercept = 0, linetype = "dashed",size=1) +
    ylim(-4,3) +
    theme(axis.title.x = element_blank()) +
    theme(axis.title.y = element_text(face="bold")) +
    theme(legend.position = "none") +
    ylab("Average fold change (log 2)") +
    theme_classic() +
    theme(plot.title = element_text(hjust = 0.5, face = "bold"))

ggplot(data7, aes(x = timept, y = value, group = gene_id, ymin = value-sd, ymax = value+sd))  + 
    geom_line(color = "#2cb877", size = 4) +
    geom_errorbar(width = 0.2) +
    geom_hline(yintercept = 0, linetype = "dashed",size=1) +
    ylim(-4,3) +
    theme(axis.title.x = element_text(face="bold")) +
    theme(axis.title.y = element_text(face="bold")) +
    theme(legend.position = "none") +
    ylab("Average fold change (log 2)") +
    theme_classic() +
    theme(plot.title = element_text(hjust = 0.5, face = "bold"))

ggplot(data8, aes(x = timept, y = value, group = gene_id, ymin = value-sd, ymax = value+sd))  + 
    geom_line(color = "#69cf52", size = 4) +
    geom_errorbar(width = 0.2) +
    geom_hline(yintercept = 0, linetype = "dashed",size=1) +
    ylim(-4,3) +
    theme(axis.title.x = element_text(face="bold")) +
    theme(axis.title.y = element_text(face="bold")) +
    theme(legend.position = "none") +
    ylab("Average fold change (log 2)") +
    theme_classic() +
    theme(plot.title = element_text(hjust = 0.5, face = "bold"))

ggplot(data9, aes(x = timept, y = value, group = gene_id, ymin = value-sd, ymax = value+sd))  + 
    geom_line(color = "#b3e000", size = 4) +
    geom_errorbar(width = 0.2) +
    geom_hline(yintercept = 0, linetype = "dashed",size=1) +
    ylim(-4,3) +
    theme(axis.title.x = element_text(face="bold")) +
    theme(axis.title.y = element_text(face="bold")) +
    theme(legend.position = "none") +
    ylab("Average fold change (log 2)") +
    theme_classic() +
    theme(plot.title = element_text(hjust = 0.5, face = "bold"))

ggplot(data10, aes(x = timept, y = value, group = gene_id, ymin = value-sd, ymax = value+sd))  + 
    geom_line(color = "#fee900", size = 4) +
    geom_errorbar(width = 0.2) +
    geom_hline(yintercept = 0, linetype = "dashed",size=1) +
    ylim(-4,3) +
    theme(axis.title.x = element_text(face="bold")) +
    theme(axis.title.y = element_text(face="bold")) +
    theme(legend.position = "none") +
    ylab("Average fold change (log 2)") +
    theme_classic() +
    theme(plot.title = element_text(hjust = 0.5, face = "bold"))


```


# 7. Paralogs simulation

We developed a function in an attempt to determine if the distribution of paralogs was significant or not 

```{r loop1}
paralog_sim_df <- subset(raw_genes, select=c("gene_id","orthogroup","cnt_OG","cluster"))

singles_sim_df <- paralog_sim_df %>%distinct(gene_id, .keep_all = TRUE) %>% group_by(orthogroup) %>% filter( n() == 1 )
singles_sim_df[,c(4)] <- sapply(singles_sim_df[,c(4)], as.character) 
singles_sim_df <- (na.omit(singles_sim_df))

doubles_sim_df <- paralog_sim_df %>% distinct(gene_id, .keep_all = TRUE) %>% group_by(orthogroup) %>% filter( n() == 2 )
doubles_sim_df[,c(4)] <- sapply(doubles_sim_df[,c(4)], as.character)
doubles_sim_df <- (na.omit(doubles_sim_df))

# 'notin' for lazy exclusion of filter searches
`%notin%` <- Negate(`%in%`) # a suprise tool to help us later

# function specifying a loop, X cycles, df as user input to function
paralog_sim <- function(x = 10, df) { # current default 10 for testing, change to 1000
  
  # create empty vector of length X for data storage
  simres <- rep(0, x)
  
  # create empty df for row info storage (for testing)
  simstore <- as.data.frame(matrix(0, x, 2))
  # simstore <- matrix(0,x,2) # uncomment for uniformity test
  
  # get length of df outside loop, for random draw boundary (0 : N)
  N <- nrow(df)
  # (loop body)
  for(n in 1:x) {
    # random draw of numbers (in vector slots [1] and [2]), without replacement
    random <- sample(1:N, 2, replace = F) # cannot draw same # twice
    
    # store values in simstore, for testing
    simstore[n,1] <- df[random[1],4]
    simstore[n,2] <- df[random[2],4]
    
    # store just the drawn nums, for uniformity test
    # simstore[n, ] <- random

    # IF statement to store 1 if one random row is not sig and other is sig
    if ((df[random[1],4] %in% c("Not2Sig","NotSig") & 
        df[random[2],4] %notin% c("Not2Sig", "NotSig")) |
       (df[random[1],4] %notin% c("Not2Sig", "NotSig") & 
        df[random[2],4] %in% c("Not2Sig", "NotSig"))) {
      
      # store 1 in vector of length X made at top of function     
        simres[n] <- 1
       }
  }


  # after loop complete, get mean of vector X (1s and 0s only, mean is proportion that match criterion: 1 row sig, other not)
  simmean <- mean(simres) 
  
  # return this as function output
  #return(simmean) # comment this out for troubleshooting
  
  # if above return (ends function) is commented out, diagnostic info returned: keep the x small to ensure results are reasonably human-readable
  testlist <- list(themean = simmean, resvector = simres, allstore = simstore)
  return(testlist)
}

simulation_test1 <- paralog_sim(1000, singles_sim_df)
simulation_test2 <- paralog_sim(1000, singles_sim_df)
simulation_test3 <- paralog_sim(1000, singles_sim_df)
simulation_test4 <- paralog_sim(1000, singles_sim_df)
simulation_test5 <- paralog_sim(1000, singles_sim_df)
simulation_test6 <- paralog_sim(1000, singles_sim_df)
simulation_test7 <- paralog_sim(1000, singles_sim_df)
simulation_test8 <- paralog_sim(1000, singles_sim_df)
simulation_test9 <- paralog_sim(1000, singles_sim_df)
simulation_test10 <- paralog_sim(1000, singles_sim_df)
simulation_test11 <- paralog_sim(1000, singles_sim_df)
simulation_test12 <- paralog_sim(1000, singles_sim_df)
simulation_test13 <- paralog_sim(1000, singles_sim_df)
simulation_test14 <- paralog_sim(1000, singles_sim_df)
simulation_test15 <- paralog_sim(1000, singles_sim_df)
simulation_test16 <- paralog_sim(1000, singles_sim_df)
simulation_test17 <- paralog_sim(1000, singles_sim_df)
simulation_test18 <- paralog_sim(1000, singles_sim_df)
simulation_test19 <- paralog_sim(1000, singles_sim_df)
simulation_test20 <- paralog_sim(1000, singles_sim_df)
simulation_test21 <- paralog_sim(1000, singles_sim_df)
simulation_test22 <- paralog_sim(1000, singles_sim_df)
simulation_test23 <- paralog_sim(1000, singles_sim_df)
simulation_test24 <- paralog_sim(1000, singles_sim_df)
simulation_test25 <- paralog_sim(1000, singles_sim_df)
simulation_test26 <- paralog_sim(1000, singles_sim_df)
simulation_test27 <- paralog_sim(1000, singles_sim_df)

paralog_sample <- function(x = 10, df) { # current default 10 for testing, change to 1000
  
  # create empty vector of length X for data storage
  simres <- rep(0, x)
  
  # create empty df for row info storage (for testing)
  simstore <- as.data.frame(matrix(0, x, 2))
  # simstore <- matrix(0,x,2) # uncomment for uniformity test
  
  # get length of df outside loop, for random draw boundary (0 : N)
  N <- nrow(df)
  # (loop body)
  for(n in 1:x) {
    # random draw of numbers (in vector slots [1] and [2]), without replacement
    lines1 <- sample(1:N, 2, replace = F) # cannot draw same # twice
    
    # store values in simstore, for testing
    simstore[n,1] <- df[lines1[1],4]
    simstore[n,2] <- df[lines1[2],4]
    
    # store just the drawn nums, for uniformity test
    # simstore[n, ] <- lines1

    # IF statement to store 1 if one lines1 row is not sig and other is sig
    if ((df[lines1[1],4] %in% c("Not2Sig","NotSig") & 
        df[lines1[2],4] %notin% c("Not2Sig", "NotSig")) |
       (df[lines1[1],4] %notin% c("Not2Sig", "NotSig") & 
        df[lines1[2],4] %in% c("Not2Sig", "NotSig"))) {
      
      # store 1 in vector of length X made at top of function     
        simres[n] <- 1
       }
  }


  # after loop complete, get mean of vector X (1s and 0s only, mean is proportion that match criterion: 1 row sig, other not)
  simmean <- mean(simres) 
  
  # return this as function output
  return(simmean) # comment this out for troubleshooting
  
  # if above return (ends function) is commented out, diagnostic info returned: keep the x small to ensure results are reasonably human-readable
  testlist <- list(themean = simmean, resvector = simres, allstore = simstore)
  return(testlist)
}

paralog_1st <- paralog_sample(, paralog_sim_df)

```
````
